# 50,000 years of Evolutionary History of India: Insights from ∼2,700 Whole Genome Sequences

**DOI:** 10.1101/2024.02.15.580575

**Authors:** Elise Kerdoncuff, Laurits Skov, Nick Patterson, Wei Zhao, Yuk Yee Lueng, Gerard D. Schellenberg, Jennifer A. Smith, Sharmistha Dey, Andrea Ganna, AB Dey, Sharon L.R. Kardia, Jinkook Lee, Priya Moorjani

## Abstract

India has been underrepresented in whole genome sequencing studies. We generated 2,762 high coverage genomes from India––including individuals from most geographic regions, speakers of all major languages, and tribal and caste groups––providing a comprehensive survey of genetic variation in India. With these data, we reconstruct the evolutionary history of India through space and time at fine scales. We show that most Indians derive ancestry from three ancestral groups related to ancient Iranian farmers, Eurasian Steppe pastoralists and South Asian hunter-gatherers. We uncover a common source of Iranian-related ancestry from early Neolithic cultures of Central Asia into the ancestors of Ancestral South Indians (ASI), Ancestral North Indians (ANI), Austro-asiatic-related and East Asian-related groups in India. Following these admixtures, India experienced a major demographic shift towards endogamy, resulting in extensive homozygosity and identity-by-descent sharing among individuals. At deep time scales, Indians derive around 1-2% of their ancestry through gene flow from archaic hominins, Neanderthals and Denisovans. By assembling the surviving fragments of archaic ancestry in modern Indians, we recover ∼1.5 Gb (or 50%) of the introgressing Neanderthal and ∼0.6 Gb (or 20%) of the introgressing Denisovan genomes, more than any other previous archaic ancestry study. Moreover, Indians have the largest variation in Neanderthal ancestry, as well as the highest amount of population-specific Neanderthal segments among worldwide groups. Finally, we demonstrate that most of the genetic variation in Indians stems from a single major migration out of Africa that occurred around 50,000 years ago, with minimal contribution from earlier migration waves. Together, these analyses provide a detailed view of the population history of India and underscore the value of expanding genomic surveys to diverse groups outside Europe.

## Introduction

With more than 1.5 billion people and approximately 5,000 anthropologically well-defined ethno-linguistic and religious groups, India is a region of extraordinary diversity^1^. Yet, Indian populations are often underrepresented in genomic studies. Recent sequencing endeavors such as the 1000 Genomes Project (1000G)^2^, UK Biobank^3^, TopMed^4^, Simons Genome Diversity Panel^5^ and GenomeAsia^6,7^ have incorporated Indian populations. However, with the exception of GenomeAsia^6,7^, these efforts have either included very few individuals or primarily sampled expatriate communities outside of India, leading to a limited (and biased) representation of the genetic variation seen in India. As a result, many open questions remain about the population history of India: When did people first migrate to India from Africa––as part of the major migration out of Africa or at earlier times along the southern coastal route of migration? What is the contribution and legacy of archaic gene flow from Neanderthals and Denisovans to Indians? How have recent technological innovations like Neolithic farming and spread of languages impacted variation in India?

To obtain a more complete picture of human diversity in India, we generated deep coverage genome sequences of ∼2,700 individuals from 18 states in India. Our samples are part of the Longitudinal Aging Study in India - Diagnostic Assessment of Dementia (LASI-DAD)^8^ that is a population-based prospective cohort study that has collected nationally representative data of individuals that are 60 years or older. These data contain individuals from diverse geographic regions (including rural and urban areas), speakers for many language families (e.g., Indo-European, Dravidian and Tibeto-Burman languages) and various ethno-linguistic and caste groups (e.g., self-reported castes recognized by the Indian government), providing the most comprehensive snapshot of genetic diversity in India.

### Data and catalog of novel variants

A total of 2,762 LASI-DAD participants, including 22 trios (mother-father-child), were sequenced at MedGenome, Inc. (Bangalore, India) at an average read depth of 30x. Individuals were sampled from 18 different states across India (Fig 1A), with median sample size of 157 individuals per state (Supplementary Note S1). The raw whole genome sequences were sent to the Genome Center for Alzheimer’s Disease (GCAD) at the University of Pennsylvania for joint calling and quality control. A total of 2,679 samples and 73.2 million autosomal bi-allelic variants passed quality control filters, including 67.1 million single nucleotide variants (SNVs) and 6.04 million insertion-deletions (indels) (Supplementary Note S2). We identified 24 million novel SNVs and 2.2 million novel indels, underscoring the limitations of existing human genetic variation databases like the 1000G and Genome Aggregation Database (gnomAD)^9^ in representing diverse populations. The vast majority (>99%) of the newly identified variants are rare, including 68% of singletons and less than 1% common variants (with greater than 1% frequency) (Table S2.1). Genome phasing was conducted using SHAPEIT4^10^, and we estimated a low phase switch error rate of less than 1.15% in trios (Table S3.1).

**Figure 1.**
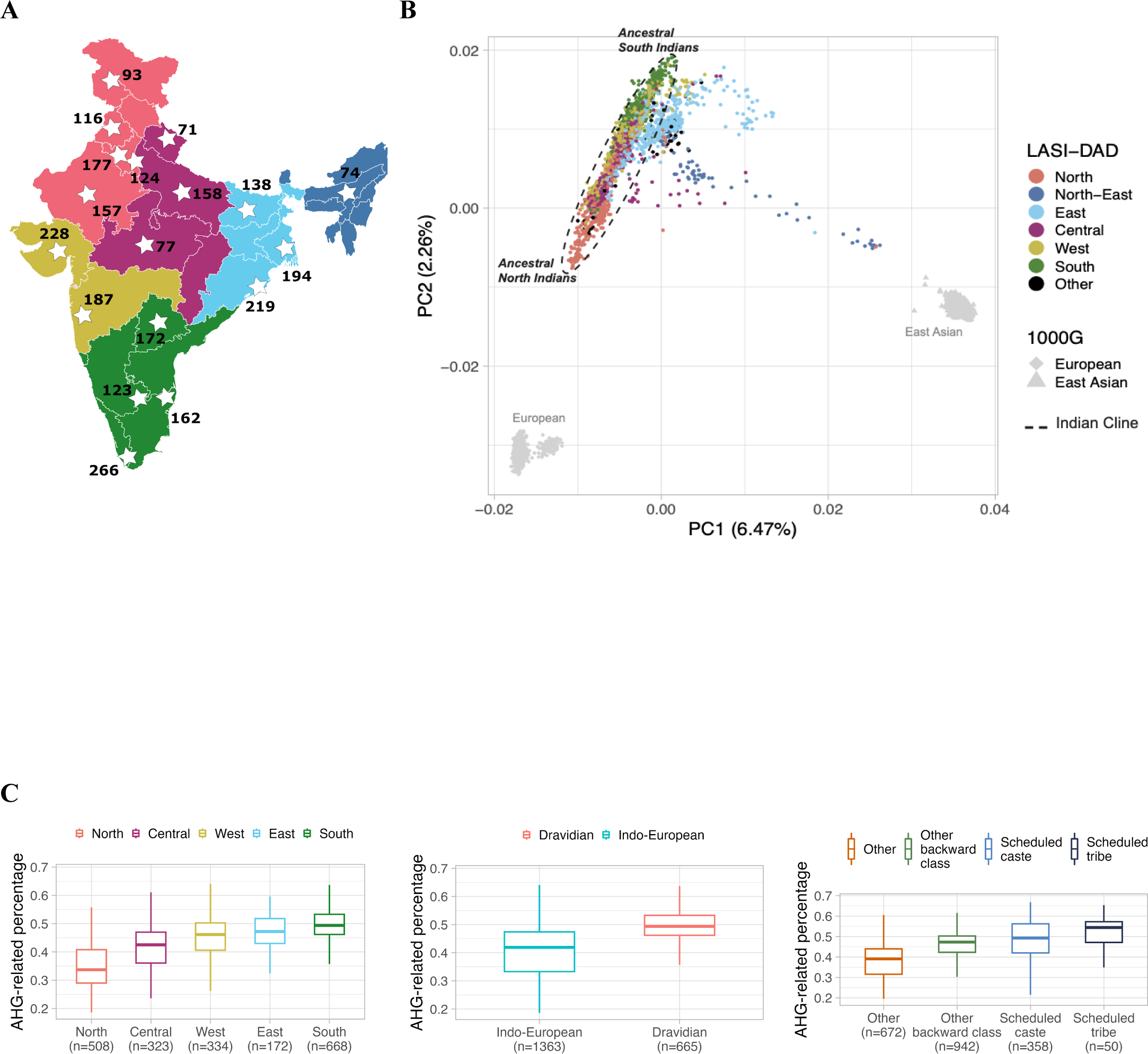
Population structure and admixture in India. (A) We show the sampling locations of individuals in the DAD study. States are colored by region (North, North-east, Central, South, East and West) used for analysis. (B) n Principal component analysis (PCA) for Indians in LASI-DAD and 1000G individuals of European (EUR), East (EAS) and South Asian (SAS) ancestry. We show the projection of the first two principal components, colored by of birth. (C) Using *qpAdm*, we inferred the ancestry proportions for each individual on the ‘Indian cline’ using *m_EN* as a proxy for Iranian farmer-related, *Central_Steppe_MLBA* as a proxy for Steppe pastoralist-related and (*Onge*) as a proxy for *AASI*-related ancestry. We compared *AHG-*related ancestry proportion by region (left), ge family (middle), and caste group (right) of each individual.

Our dataset is representative of the population diversity in India. It includes individuals born in 23 different states from both rural (63%) and urban (37%) areas. It comprises speakers of around 26 different languages that belong to diverse caste groups as recognized by the Indian government: 4% from Scheduled Tribes, 18% from Scheduled Castes, and 44% from other backward class (OBC). Nearly equal numbers of males and females were recruited in the study, with our dataset constituting 52% of females. For many analyses, we categorized individuals based on their birth location into six major geographic regions: North (*n*=555), West (*n*=385), Central (*n*=373), South (*n*=715), North-East (*n*=73), and East (*n*=530). After performing quality control checks and excluding first-degree relatives, we used a sample of 2,620 individuals for most of our analyses described below, unless specified otherwise (see Methods, Supplementary Note S1-2).

### Population structure and admixture

To study population relationships of Indians to other worldwide populations, we combined the LASI-DAD dataset with the 1000G^11^ and applied Principal component analysis (PCA)^12^, ADMIXTURE^13^ and *f*-statistics^14^. Consistent with previous reports^15,16^, we find that the population structure in India is related to individuals of West Eurasian-related ancestry (1000G EUR), with limited or no recent gene flow from populations related to sub-Saharan Africans (Fig 1B, Fig S4.1). The population structure in India is correlated to geography (state of birth) and linguistic affiliation, with three main clusters––one cluster that includes the majority of the individuals from North and South of India who speak Indo-European and Dravidian languages and represents varying relatedness to West Eurasians, referred to as ‘Indian cline’ (Fig 1B, Fig S4.2-3). The Indian cline has previously been shown to reflect variable proportions of ancestry from two ancestral groups: the *Ancestral North Indians* (*ANI*) who harbor large proportions of ancestry related to West Eurasians, and the *Ancestral South Indians* (*ASI*) who are distantly related to West Eurasians^15,16^. Recent ancient DNA analysis have shown that both *ANI* and *ASI* are admixed and in turn, have ancestry from groups related to ancient Iranian farmers, ancient Eurasian Steppe pastoralists, and unsampled indigenous South Asians (*Ancient Ancestral South Indians (AASI*)) distantly related to Andamanese hunter-gatherers (*AHG*)^17^.

Beyond the Indian cline, we find two primary clusters of individuals (*n*=494): a cluster towards the *ASI*-end of the cline, and another found closer to the center exhibiting clear relatedness to East Asian-related groups (1000G EAS) in PCA (Fig 1B). The former mainly includes individuals from Central and East India, with the majority from the state of Odisha where predominantly Indo-European and Austro-asiatic languages are spoken. The East Asian-related cluster includes individuals from East and North-East regions of India. West Bengal is the most representative state in this cluster, with almost 10% ancestry related to East Asians. Using ALDER^18^, we estimated the admixture related linkage disequilibrium related to EAS to infer that this gene flow occurred 50 generations ago or around 520 AD, possibly related to the invasions of the Huna people to India after the collapse of the Gupta Empire (Fig S4.11)^19,20^. Another predominant group in the East Asian-related cluster is from Assam. This group exhibits significant heterogeneity, as individuals have varying degrees of relatedness to EAS, indicative of the recent gene flow possibly related to the recent migration of East Asian tea plantation workers to India in the last two centuries^21^ (Fig 1B). Our ADMIXTURE^13^ analysis mirrors the patterns seen in PCA (Fig S4.6).

### Ancestry Composition and Sources

To model the ancestry in India, we used *qpAdm* that compares allele frequency correlations between a population of interest and a set of reference and outgroup populations^14,22^. First, we examined how well the three-way model with ancient Iranian farmer-related, Eurasian Steppe pastoralist-related, and *AHG*-related groups describes the ancestry of individuals on the Indian cline (Fig 1B). Following Narasimhan et al. 2019^17^, we used *Indus Periphery West* that is part of the *Indus Periphery Cline*––a heterogenous group of 11 outlier samples from Bronze Age cultures of Shahr-i-Sokhta and Bactria Margiana Archaeological Complex––as the proxy for Iranian farmer-related ancestry, Central Steppe Middle to late Bronze age (*Central_Steppe_MLBA*) as the source for Yamnaya Steppe pastoralist-derived ancestry and *AHG*-related individuals to represent *AASI* ancestry^17^. We find the three-way model provides a good fit for the majority (>90%) of the individuals on the Indian cline, with some exceptions (we define ‘good fit’ as models with *qpAdm p*-value > 0.01, see Methods). Notably, we find 22 individuals that can be fitted as a two-way mixture between ancient Iranian farmer-related and *AHG*-related ancestries without Steppe pastoralist-related ancestry (referred to as *ASI* henceforth).

The archaeological context of the *Indus Periphery Cline* and their relationship to ancient Indian civilizations (e.g., Indus Valley Civilization) is unclear as these were migrant samples from Bronze Age Central Asian cultures^17^. Thus, we examined fifteen ancient Iranian-related groups from the Neolithic to Iron Age as the potential source of the Iranian farmer-related ancestry for the 22 *ASI* individuals and *Indus Periphery West*. We obtain good fits for all 22 *ASI* individuals when the Iranian-related ancestry derives from early Neolithic and Copper Age individuals from Central Asian cultures of either *Sarazm_EN* or *Namazga_CA* or a group containing *Sarazm_EN* and *Parkhai_Anau_EN* that was previously suggested as the source for *Indus Periphery Cline*^17^. The latter two models also provide good fits for *Indus Periphery West*, though using *Sarazm_EN* alone as the source does not yield a good fit (Table S4.2). Furthermore, a model with *Sarazm_EN*, *AHG*-related and *Central_Steppe_MLBA* also provides a good fit for the vast majority (>95%) of individuals on the Indian cline (*p-*value in *qpAdm* > 0.01). In contrast, models with *Namazga_CA* fail for >15% of individuals on the Indian cline, contrary to previous claims based on fewer samples^23^. Similarly, models with *Sarazm_EN* and *Parkhai_Anau_EN* do not work well for modern Indians and yield negative coefficients for *Parkhai_Anau_EN* ancestry (Table S4.3).

Turning to the individuals that fall outside the Indian cline, we tried three models including *Sarazm_EN*, *AHG*-related, and either (*a*) Steppe pastoralist-related (as the Indian cline model), (*b*) Austro-asiatic-related (using *Nicobarese*), or (*c*) East Asian-related (using *EAS*) ancestries. We also tested four-way models with addition of *Central_Steppe_MLBA* if models (*b-c*) failed. We obtain good fits for 91% of the individuals that fall outside the cline (Table S4.4). Notably, there are 91 individuals that can be modeled without Steppe pastoralist-related ancestry, including ∼96% of the Austro-asiatic-related individuals (using model *b*). This suggests Iranian farmer-related ancestry likely did not come through Steppe pastoralist-related groups to India.

Archaeological studies have also documented trade connections between Sarazm and South Asia, including connections with agriculture sites of Mehrgarh and early Indus Valley Civilization^24^. Indeed, one of the two *Sarazm_EN* individuals (*Sarazm_EN_1*) was found with shell bangles that are identical to ones found at sites in Pakistan and India such as Shahi-Tump, Makran and Surkotada, Gujarat^25^ (*J. Mark Kenoyer*, personal communication). Surprisingly, when we applied *qpAdm*, we discovered that *Sarazm_EN_1* has substantial *AHG*-related ancestry (∼15%), unlike the other individual from the *Sarazm_EN* group (*Sarazm_EN_2*). Application of the three-way model with *Sarazm_EN_2*, *AHG*-related and *Central_Steppe_MLBA* continues to provide a good fit for most individuals (>96%) on the Indian cline, as well as off-cline individuals (Table S4.7-8). Moreover, the two-way model without Steppe Pastoralist-related ancestry works well for the 22 *ASI* individuals and *Indus Periphery West* (without need for additional ancestry from *Parkhai_Anau_EN*). Together, our data are consistent with a common source for the ancient Iranian-related ancestry in ANI, ASI, Austroasiatics-related and East Asian-related individuals in India, suggesting that the Iranian-related gene flow occurred well before the arrival of Steppe pastoralist-related ancestry in Bronze Age (∼1900–1500 BCE^17^).

Using *AHG*-related, *Sarazm_EN* and *Central_Steppe_MLBA* as reference populations, we inferred the genetic composition of individuals on the Indian cline. We find marked variation in ancestry proportions across India, with Iranian farmer-related ancestry varying between ∼27–68%, *AHG*-related between ∼19–69% and *Central_Steppe_MLBA* between ∼0–45%. Among the three ancestry components, variation in *AHG*-related shows the strongest correlation to the ANI-ASI cline in PCA (Fig S4.10). *AHG*-related ancestry proportion is significantly associated with geography (e.g., highest in South and lowest in North of India), language (i.e., higher in Dravidian vs. Indo-European language speakers) and caste affiliation (highest in Scheduled Castes, Scheduled Tribes and OBC compared to other groups) (Fig 1C, Extended Data Fig 1). This highlights that the ancient admixture events are related to the spread of languages and the history of the traditional caste system in India.

### Founder events increase homozygosity in India

Previous studies have shown that many Indian groups have a history of strong founder events, due to endogamous and consanguineous marriages^7,26,27^. Founder events reduce genetic variation and increase sharing of genomic regions that are inherited identical-by-descent (IBD) from a few common ancestors^28^. Descendants of consanguineous marriages (between close relatives) may inherit IBD segments from both parents, resulting in segments that are homozygous-by-descent (HBD). A founder event results in many, small HBD segments, while recent consanguinity results in fewer but longer HBD segments.

We identified IBD and HBD segments in LASI-DAD and 1000G datasets using a haplotype-based IBD detection method, *hap-IBD*^29^. To differentiate between the relative effects of founder events and recent consanguineous marriages, we stratified the HBD segments by length– long (> 8cM) indicative of consanguinity and short (< 8cM) mostly reflecting founder events. Indians, on average, have a larger fraction of their genome in HBD segments (∼29 cM) compared to 1000G EAS (∼6 cM), EUR (∼6 cM), and AFR (∼4 cM) (Fig 2A). Within India, individuals from South have significantly higher homozygosity, both in terms of the total amount of their genome in HBD segments (on average, ∼56 cM in South compared to ∼19 cM in other regions, *p*-value < 10^-16^) and the fraction of long HBD segments (8.4% vs. 4.3%, *p*-value < 10^-6^), reflecting the higher prevalence of consanguineous marriages in the South of India^30^ (Fig 2A, Fig S5.1-2). A majority (>90%) of the homozygosity stems from small HBD segments (rather than long HBD segments), suggesting a primary role of historical founder events rather than recent consanguinity as the source of homozygosity (Fig 2A, Fig S5.2). Similar results are obtained when we use a threshold of 20 cM to define long HBD segments (Fig S5.1, Fig S5.2B).

**Figure 2.**
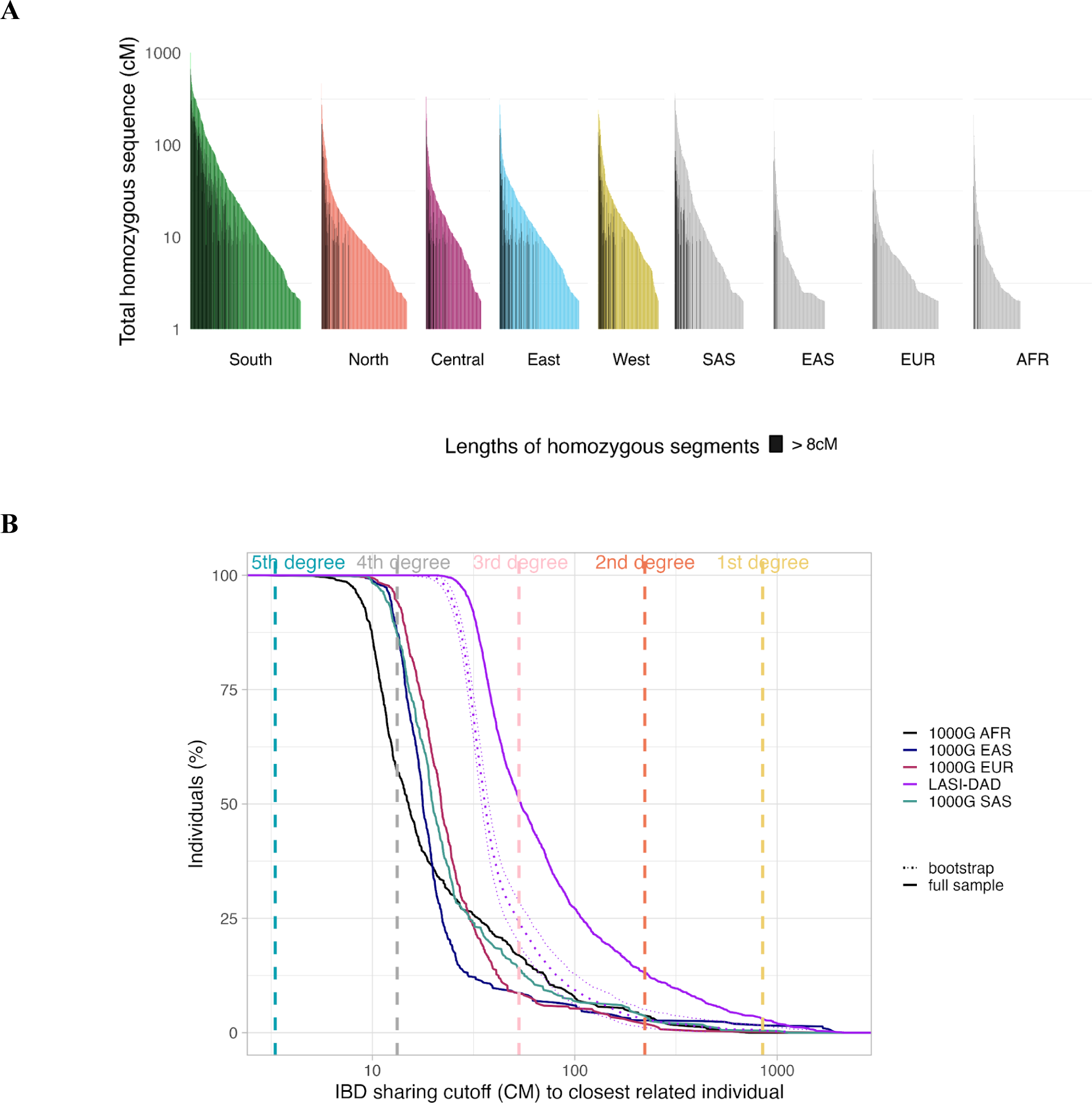
Founder events and consanguinity leads to high rates of homozygosity and relatedness in Indians. (A) plied hap-IBD to infer genome-wide homozygosity in LASI-DAD samples grouped per region and compared ther world-wide groups: East Asian, European, and South Asian populations from 1000G. Black lines show the amount of homozygous segments longer than 8cM per individual, and colored lines the total amount of zygous segments shorter than 8cM. (B) For each of the 2,620 Indian samples and AFR, EAS, EUR and SAS duals in 1000G, we detected the individual sharing the largest total amount (in cM) of genome IBD, referred to osest individual’. For each value *x* of total shared genome (in *cM*) on the *X*-axis, we report the percentage of es (*Y*-axis) that share *x* or more with their closest related individual. For LASI-DAD individuals, we also detect osest individuals while bootstrapping to 500 individuals (dashed lines representing mean and 95% CI). The ntal dashed lines indicate the expected value of the total IBD sharing for *k*th degree cousins. This figure was d from ^32^.

Next, we investigated genome-wide IBD-sharing across individuals. We computed the fraction of individuals who find at least one close genetic relative within LASI-DAD and compared this proportion across worldwide populations in 1000G (see Methods, Fig S5.3). We infer that ∼51.0% (38.4–59.2% across regions) of individuals in LASI-DAD find at least one genetic relative with expected IBD sharing equivalent to a 3rd degree cousin or closer relationship (∼53 cM) in LASI-DAD, which is markedly higher than 14.2% in SAS, 8.8% in EAS, 8.8% in EUR and 17.2% in AFR from 1000G (Fig 2B, Table S5.1) (note, a previous study identified ∼5–10% of individuals are first and second-degree relatives in Gambians from Mandinka (GWD) and Esan in Nigeria (ESN) contributing to higher relatedness in AFR^31^). The higher IBD sharing in LASI-DAD, especially compared to 1000G SAS may stem from: (a) larger sample size of LASI-DAD, or (b) ascertainment bias in selecting individuals in either study. We examined each of these hypotheses in turn. We performed bootstrap resampling of equal numbers of individuals (*n*=500) from LASI-DAD as 1000G SAS and inferred that the fraction of 3rd degree cousins decreased to 24.2% (95% CI: 19.4%–28.6%), yet significantly higher than 1000G SAS (Fig 2B, Table S5.1). In LASI-DAD, individuals were recruited using a stratified random sampling approach. First, Sampling Secondary Units (SSUs) (villages/urban census blocks) were chosen in each state and then within each SSU, individuals were selected randomly. To control for the impact of this ascertainment scheme, we considered pairwise cross-SSU comparisons among individuals (Supplementary Note S5). Using this approach and accounting for the sample size, we continue to find a significant shift in LASI-DAD compared to 1000G SAS, with ∼16.4–35.0% of individuals sharing IBD equivalent to 3rd degree cousins (Fig S5.4). This comparison highlights the limitations of the sampling of 1000G groups for representing genetic variation of India (with mainly a few groups from the subcontinent). Overall, we find that all individuals in LASI-DAD have at least one putative 4th degree cousin or closer relative (with IBD > 10 cM) in the dataset. The high level of relatedness in India is notable, as a similar level of IBD sharing is seen in Europeans with approximately 480,000 individuals (almost 200-fold higher sample size) in UK Biobank^32^.

The history of founder events predicts a high burden of deleterious variants and increased risk of recessive diseases, as seen in Finns and Ashkenazi Jews^28,33^. To assess the potential functional effects of founder events in India, we identified 385,985 missense and 20,319 putative loss of function (pLoF) variants (see Methods) (Table S5.2). Each individual carries ∼10,344 (range: 9,911–10,761) derived missense variants, and ∼67 (46–96) pLoF variants on autosomes. Most (>90%) of these variants are rare (frequency below 1%) or singletons (62%). As expected, we observe strong correlation between the homozygous deleterious mutation burden (measured as sum of homozygous missense and pLof variants carried by an individual) and the total sum of HBD per individual in India (Extended Data Fig 2). Among 18,451 protein-coding autosomal genes in the human genome (RefSeq database^34^), we find missense and pLoFs variants in 89.5% of the genes, ranging between 1–1,265 variants per gene. The top three genes with the highest number of pLoFs variants are mucin genes: MUC3A, MUC16 and MUC17, with respectively 52, 42 and 41 pLoFs, including homozygous pLoFs in MUC17. As there is partial redundancy in the function of mucin genes, there may be greater tolerance for loss of function variants^35^.

Among the 406,304 SNVs, we find about half are South Asian-specific and a large fraction (40%) are absent in gnomAD or 1000G (Table S5.2). We find that ∼4% of South-Asian specific non-ultra rare (frequency above 0.1%) missense/pLoF variants are present in the ClinVar database^36^, including 10 classified as ‘pathogenic’ variants (using ClinVar threshold of two-stars, Table S5.2). Among these, we find a South-Asian specific pathogenic variant in the *BHCE* gene that is present in 15 individuals (0.28%) in LASI-DAD (and not seen outside India). Patients with butyrylcholinesterase deficiency may experience prolonged neuromuscular blockade and muscle paralysis, in response to use of some muscle relaxants used during anesthesia. Previous studies have identified this variant in the founder community of Vysya from Andhra Pradesh where it has drifted to high frequency due to the history of founder events^27,37^. In LASI-DAD, 8 of the 15 individuals are from Telangana, the neighboring state of Andhra Pradesh. Local community doctors use the Vysya ancestry as a counter-indicator before administering anesthetic drugs, highlighting the potential of reducing disease burden by understanding and documenting the effects of founder events in India.

### Gene flow from archaic hominins in India

Most non-Africans, including Indians, derive ∼1-2% of their ancestry from gene flow from archaic hominins, Neanderthals and Denisovans^5,38^. The functional impact and regional variation in archaic ancestry in India, however, remains unclear. We applied a reference-free hidden Markov model, called *hmmix*^33^, to 2,679 phased individuals from India (to maximize our sample size, we retained first-degree relatives (except offspring of trios)). *hmmix* classifies genomic fragments into two states––‘modern human’ or ‘archaic’––by comparing the density of derived alleles that are not found in 490 sub-Saharan Africans (who have negligible amount of archaic ancestry^25^) (see Methods). We also applied *hmmix* to phased data from 2,309 individuals from 1000G, 825 individuals from Human Genome Diversity Panel (HGDP), and used the published results for 27,566 Icelanders from deCODE genetics that were also analyzed using the same method^26^. Unless stated otherwise, we retained archaic ancestry segments with a posterior probability greater than 0.8 for subsequent analysis that translates to <4% false positive rate in simulations^26^.

We inferred that Indians have an average of 102.98 Mb or 2.07% of the callable genome (95% percentile range: 1.84–2.34%) of archaic ancestry. By comparing the putative archaic segments to sequenced Neanderthal and Denisovan genomes^39,40,41,42^, we inferred the source of the archaic ancestry based on measuring the number of shared derived archaic variants (DAV) present on archaic segments. We find that each individual has ∼1.48% (95% percentile range: 1.30–1.69%) Neanderthal and ∼0.14% (95% percentile range: 0.07–0.21%) Denisovan ancestry. The Neanderthal ancestry proportion in India is similar to Europeans (1.3%) and Americans (1.4%), though significantly lower than East Asians (∼1.8%, Wilcoxon ranked test *p*-value < 10^-15,^). The highest Denisovan ancestry is inferred in Oceanians (∼1.8%), while Americans, East Asians and South Asians have similar amounts (∼0.1%) (Table S6.4-5).

By assembling non-overlapping archaic ancestry segments extracted from individuals in LASI-DAD, we reconstructed 1,524 Mb of the introgressing Neanderthal and 591 Mb of the introgressing Denisovan genome (Extended Data Fig 3). Notably, using individuals from all world-wide regions (from 1000G, HGDP and LASI-DAD), we reconstructed 1,679 Mb of the introgressed Neanderthal genome that is similar in size to the sequenced Neanderthal genomes (∼1,650 Mb, Fig S6.8, Supplementary Note 6). Despite higher per individual Neanderthal ancestry in East Asians, we recover more Neanderthal sequence from Indians than East Asians even after controlling for the sample size (as seen in ^38^, Table S6.5, S6.8). This is in part due to introgressed Neanderthal segments having a higher frequency in East Asia and thus being more likely to be shared across individuals (Fig S8.4)^38,43^. The largest study of archaic ancestry in 27,566 Icelanders recovered 978 Mb of the introgressed Neanderthal and 112 Mb of the introgressed Denisovan genome (using posterior probability >0.9 in *hmmix*)^44^. Even with the more stringent posterior probability threshold, we recover >50% more Neanderthal ancestry segments from Indians (LASI-DAD) than from Icelanders (Fig 3A). Using all world-wide regions, we reconstructed 1,080 Mb of the introgressing Denisovan genome. The largest amount of this is recovered from Indians, though this is not significant after downsampling to the sample size of Oceanians (*n*=28) (Fig S6.8).

**Figure 3.**
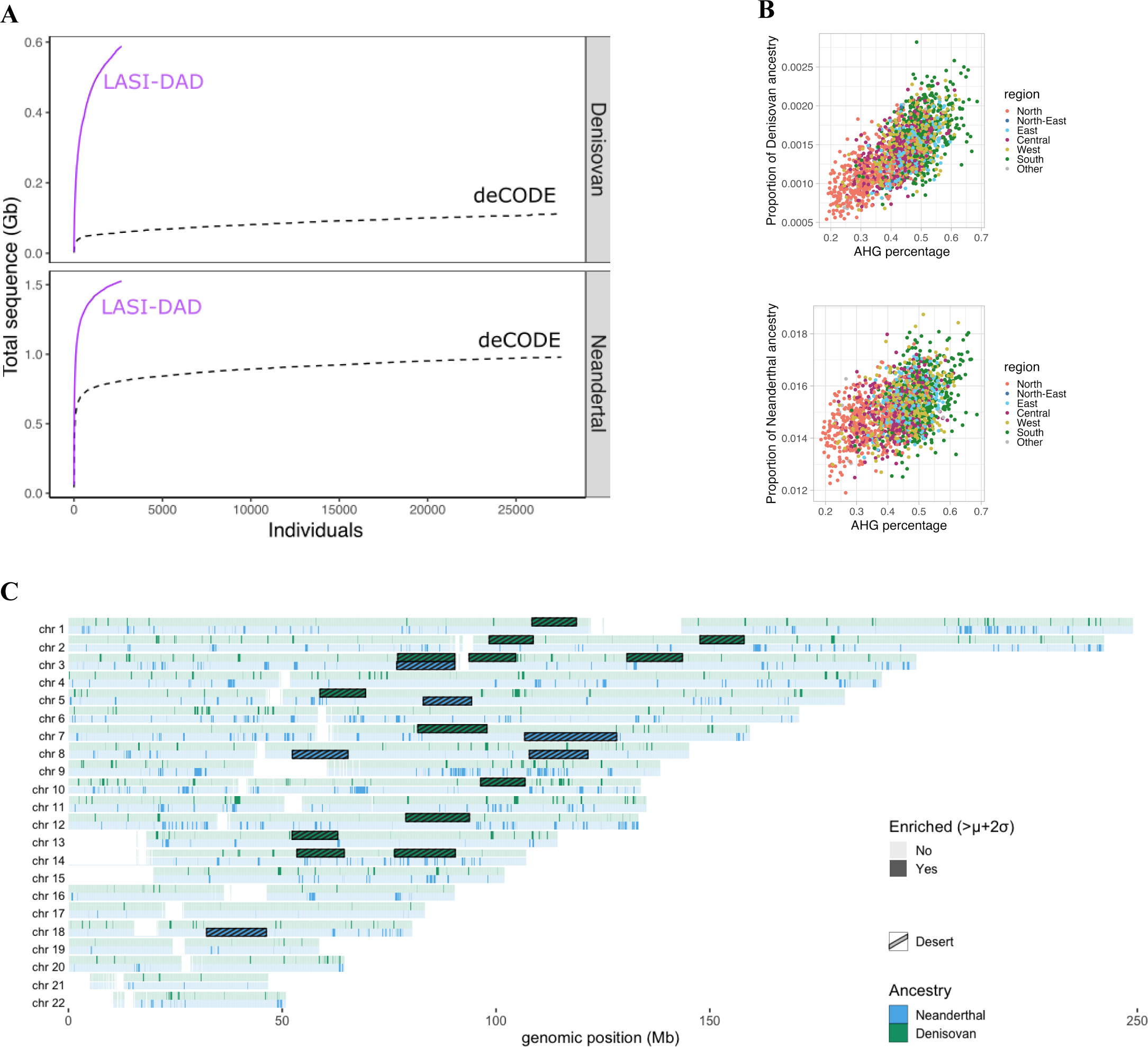
History of archaic gene flows in India. (A) Cumulative amount of unique sequence (in Gb) that is either an (top) or Neanderthal (bottom) as a function of number of individuals, in Indians from LASI-DAD (in purple) anders from deCODE (in black, dashed). (B) Correlation between *AHG-*related ancestry on the x-axis and total on of archaic sequence per individual. Individuals are colored according to which region of origin. We show the ion for Denisovan (top, r=0.49, *p*-value < 10^-15^) and Neanderthal (bottom, r=0.23, *p*-value < 10^-15^). (C) tion of archaic ancestry regions across the genome. We computed the mean archaic frequency along the genome I-DAD individuals and considered segments with an archaic frequency higher than the mean (*μ*) + two standard ns (σ) as enriched. We detected 117.28 Mb enriched in Neanderthal ancestry (in blue) and 61.52 Mb enriched in an ancestry (in green). We also show the location of archaic ancestry deserts: regions with < 0.1% archaic over 10 Mb (striped rectangles in bleu for Neanderthal and green for Denisovan).

Next, we calculated the amount of archaic sequence that is shared between Indians and other worldwide populations from 1000G and HGDP datasets. By sharing we refer to segments which overlap the same genomic regions. We find that 81.2% of Neanderthal ancestry is shared between at least two global regions (Extended Data Fig 4). We find a total of ∼11.7% (or 195.9 Mb out of 1,679 Mb) of uniquely India-specific Neanderthal sequences. Strikingly, ∼90.7% of worldwide Neanderthal sequences are seen in India (Extended Data Fig 5). Moreover, Oceanians and South Asians have large amounts of unique Denisovan ancestry sequences (Fig S6.6). Around 51% of Denisovan sequence (301.6 Mb out of 591 Mb) is unique to India (Fig S6.6). Even after downsampling to sample sizes to match the minimum sample sizes in 1000G (*n*=490) and HGDP (*n*=28), we find significant enrichment for unique Denisovan sequences in Indians (Fig S6.8).

To infer the relationship of the introgressed archaic population to the sequenced archaic genomes, we estimated DAV SNP match rates for each introgressed segment to sequenced Neanderthals and Denisovan genomes. We find on average the introgressed Neanderthal segments share 83% of the DAVs with one of the three sequenced Neanderthal genomes, with the highest sharing with the Vindija Neanderthal (Table S6.10 and Fig S6.11). In contrast, the introgressed Denisovan genome only shares 47% of DAVs with the sequenced Denisovan genome, indicating the Denisovan ancestry primarily derives from a group that is distantly related to the sequenced Altai Denisovan. Using a similar approach as Browning et al. 2018, we replicate the finding of a single pulse of Neanderthal gene flow in India (Supplementary Note 6)^45^. We find that a single Denisovan-related wave is consistent in most groups in India. Individuals in North-East and South of India, however, have evidence for two clusters of Denisovan-related sequences, one closely related to the sequenced Altai Denisovan genome (segments share on average 84% of DAV SNPs) and a more distantly related group (with 46–50% of shared DAV SNPs) (Fig S6.9). Individuals in North-East India derive a large fraction of ancestry from recent East Asian-related groups (Fig 1B) that have previously been shown to have two pulses of Denisovan ancestry^45^. Beyond Neanderthal and Denisovan ancestry, we inferred 0.42% (95% percentile range: 0.37–0.48%) of archaic ancestry from an unknown source in Indians (Table S6.4-5). This proportion is similar across all non-Africans and potentially related to the difference between the sequenced archaic genome and the introgressing archaic individuals (Fig S6.3). Consequently, this suggests that there is no clear evidence for additional contribution from other unknown archaic hominins to Indians (at least not more than other worldwide populations), contrary to previous claims^46^.

Archaic ancestry varies across regions in India, with the highest archaic ancestry in the North-East and East of India and lowest in North India (Figs 3B and S6.3, Tables S6.4 and S6.6). To investigate how recent gene flow events have shaped the distribution of archaic ancestry in India, we examined the relationship between Neanderthal and Denisovan ancestry as a function of the three main ancestry components in India. Focussing on individuals on the Indian cline (*n* = 2,126), we find the *AHG*-related ancestry is positively correlated with both Denisovan (*r* = 0.46, *p*-value < 10^-15^) and Neanderthal (*r* = 0.24, *p*-value < 10^-15^) ancestries (Fig 3B, Table S6.8). These results are robust to use of more stringent criteria for assigning archaic ancestry segments to Neanderthal and Denisovan origin, by focussing on sites where only one archaic group has a derived allele that matches modern humans (see Table S6.8). This suggests that a large amount of the archaic ancestry seen in present-day Indians is inherited through *AHG*-related ancestry and in turn, groups with higher *AHG*-related ancestry in the South have higher archaic ancestry.

### Functional legacy of archaic ancestry in India

Previous analyses have shown that archaic ancestry has played a major role in human adaptation and disease, however, few studies have evaluated its role in South Asian populations^38,47^. We examined the genome-wide distribution of archaic ancestry and identified regions of ‘high archaic frequency’ among Indians (defined as regions where the archaic frequency across individuals is two standard deviations above the genome-wide average) (Fig 3C). We identified 1,590 and 818 candidate regions with high frequency of Neanderthal and Denisovan ancestry respectively. For Neanderthals, we replicated genes such as FBP2 and FYCO1 previously identified in other studies^47–49^, as well as identified PCAT7 and CXCR6 as new candidates. For Denisovans, we replicated signals in WDFY2, CHD1L and HELZ2^47^ and identified several new candidates including LINC00708 and CDKN2B (Supplementary Note 7, Extended Data Table S3). Performing a gene ontology (GO) enrichment analysis, we find 14 pathways enriched for Neanderthal and 22 pathways for Denisovan ancestry primarily related to immune function (Extended Data Table S4).

Next, we searched for regions that have a high number of derived alleles that are shared between modern humans and archaic groups, a signature previously observed for *EPAS1* and Denisovan ancestry in Tibetans^50^. Interestingly, we find certain regions of the genome have a disproportionately elevated number of variants with derived alleles that are uniquely shared between Denisovans and Indians; though no similar enrichment is seen for uniquely Neanderthal shared variants (Supplementary Note 7). Notably, we find that the *BTNL2* gene, part of the major histocompatibility complex (MHC), contains 78 uniquely derived Denisovan variants within a 13.2-kilobase (kb) region with an exceptionally high Denisovan frequency in Indians of around 10% (> 99.9th percentile). There are two Denisovan haplotypes in this region: a *short* haplotype of 55–65 kb and a *long* one of ∼150 kb with 116.1 and 126.7 uniquely derived Denisovan variants respectively. The proportion of long haplotypes is lower in the North (*Z* = -2.26) and higher in the West of India (*Z* = 2.57) compared to all individuals in India (Fig S7.3-4). These Denisovan haplotypes are also present at high frequency in East Asians (∼11.8%, >99.8 percentile), but they are rare in Europeans (∼0.4%) and notably, absent in Oceanians (Table S7.2). The haplotype length and number of shared derived alleles between Indians and Denisovans suggests this region is likely a product of gene flow from Denisovan or Denisovan-related populations, rather than ancestral lineage sorting (*p*-value < 10^-6^ for the *long* haplotype; *p*-value=0.027 for the *short* haplotype). The MHC contains many genes associated with immune function and is most likely to be under balancing selection. Indeed, previous studies have identified *BTNL2* as a candidate for selection in East Asians^51^. Though simulations show that genetic drift generated by founder events alone can lead to high frequency of archaic ancestry in a region, thus caution is warranted when interpreting high frequency archaic regions as candidates for selection or adaptive introgression in modern humans (Supplementary Note S8).

To identify Indian-specific enriched archaic segments, we computed the population branch statistic (PBS)^52^. The PBS measures the increase in frequency at a given locus in a population, since its divergence from the two reference populations. To this end, we apply PBS using Indians as the population of interest and East Asians and Europeans as reference groups using archaic allele frequency vs. genotype frequencies to identify candidate archaic enriched regions in India (see Methods). We identified ∼10.7 Mb (or 235 genes) enriched for Neanderthal and ∼5.5 Mb (or 84 genes) for Denisovan ancestry (Extended Data Table S3). Denisovan ancestry regions are enriched for genes related to innate immune response, including several TRIM genes–TRIM26, TRIM31, TRIM15, TRIM10 and TRIM40– implicated in cellular processes related to entry (or exit) of virus into a host cell. Among the most significant candidate regions of Neanderthal ancestry is a gene cluster on chromosome 3 which has been previously associated to COVID susceptibility^53,54^ (PBS_Neanderthal_ > 0.118, in the 99.99% percentile of genome-wide PBS scores). In turn, it was discovered that there are two main haplotypes introgressed from Neanderthals containing the risk variant: a *core* haplotype of 49.4 kb and a *long* haplotype of 333.8 kb. In LASI-DAD, both of these haplotypes fall outside the 99% tail of our genome-wide distribution of Neanderthal ancestry, though there is large variation in Neanderthal haplotypes in this region including some very long haplotypes that are greater than 1 Mb (*p*-value for *core* haplotype = 0.00021, *p*-value for *long* haplotype = 0.0020, Fig S7.6A). Across India, the frequency of *core* haplotype ranges between 20.5% (in North-East) to 34.8% (in East India). The frequency of both the *core* and *long* haplotypes is significantly higher in the East of India compared to other regions (*core*: 34.8%, *Z* = 2.68, *long*: 23.2%, *Z* = 2.34).

We also examined regions of the genome devoid of archaic ancestry in modern humans, referred to as ‘archaic deserts’^44,48,55,56^. We identified six Neanderthal deserts spanning a total of 87.1 Mb including five that were previously reported (Fig 3C, Fig S7.8, Table S7.4). The location of these five Neanderthal deserts remains similar with around 70% overlap with previously identified deserts in Europeans and other populations (Table S7.4). Interestingly, among these deserts is a region that includes the FOXP2 gene that is associated with language development in humans^55^. We also identified 13 Denisovan deserts in Indians, including one that overlaps with previously reported Neanderthal deserts (Fig 3C, Fig S7.9, Table S7.5). Given the low genome-wide proportion of Denisovan ancestry in Indians, we likely miss Denisovan ancestry in some regions and thus, over-call Denisovan-related deserts.

### First arrival of modern humans to the Indian subcontinent

A central question in the peopling of India is when modern humans first arrived to the subcontinent from Africa. Archeological evidence suggests occupation in Northern India before and after the Toba eruption that occurred around 74,000 years ago^57^. It is unclear, however, if this group contributed to the ancestry of present-day peoples in India. In order to test this hypothesis, we computed the minimum coalescence time of present-day Indians, East Asians, Europeans and Americans to sub-Saharan Africans. If there is a substantial contribution from the population who lived in India before the Toba eruption, it should be detectable as an increase in coalescence time of Indians compared to individuals from other worldwide regions. To estimate the coalescent time for each non-African individual to sub-Saharan Africans, we used the rate of emission in the modern human state of *hmmix* after controlling for bioinformatics effects (phasing errors and depletion of triallelic sites) and excluding individuals with more than 1% sub-Saharan African-related ancestry (see Methods). Theoretically, the emission parameter should be proportional to the minimum coalescence time between the test individual and sub-Saharan Africans, human mutation rate (0.45×10^-9^ per base pair per year^58^, Fig S9.3) and the length of the genome surveyed.

We infer the minimum coalescence time between Indians and sub-Saharan Africans as 53,932 (95% percentile range: 53,190–54,644) years ago (Table S9.2, Fig 4). We obtain qualitatively similar results for Europeans, East Asians and South Asians in the HGDP dataset. Moreover, by performing simulations, we show the observed emission parameter in India is consistent with variation stemming from 0–3% of ancestry from an earlier migration that occurred around 74,000 years ago (Fig S9.5). Our results thus show that the majority of the ancestry of present-day Indians derives from a major migration event out of Africa that occurred 50,000 years ago.

**Figure 4.**
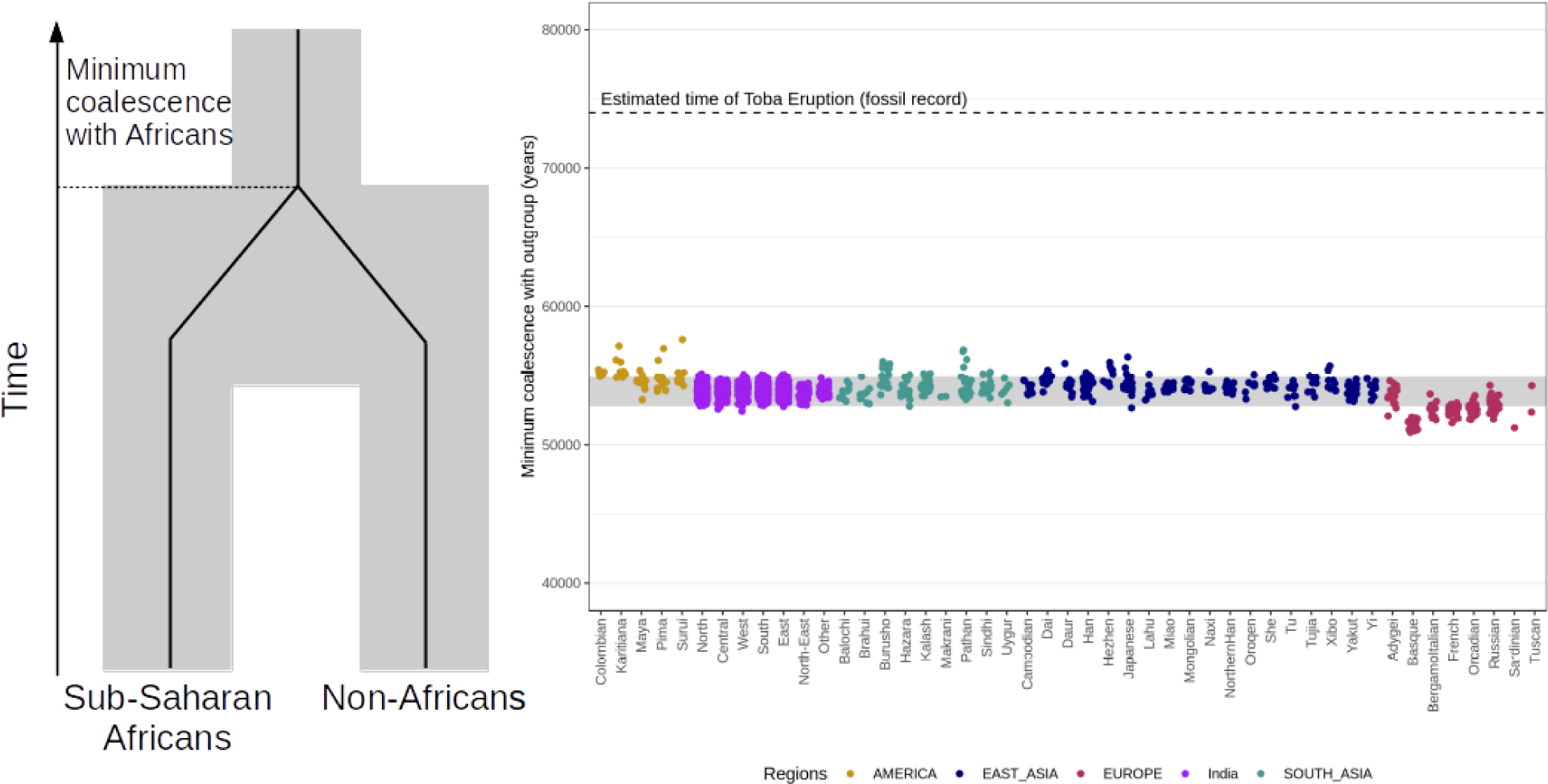
Minimum coalescence time with Sub-Saharan African populations. Each dot represents the minimum coalescence time with Sub-Saharan Africans estimated from the emission parameters of the human state using *hmmix*. The X-axis shows the population the individual belongs to and the color represents the region. The gray area represents 95% of the coalescence times for all non-African individuals. The dotted line shows the timing of the Toba eruption 74,000 years ago^57^ which provides a minimum bound for the Southern Dispersal out of Africa.

## Discussion

India is a region of extraordinary genetic diversity, including largest variation in archaic ancestry among modern humans. Notably, a majority of Neanderthal ancestry that exists today in present-day individuals is found in India, while other worldwide populations retain only a subset of this variation (Extended Fig 5). Indians also harbor the most Denisovan ancestry among Eurasian populations. Moreover, some of the deepest mtDNA and Y-chromosome lineages are seen in people from Andaman Islands^59^. Interestingly, such large diversity is also reflected in the early Middle Paleolithic stone tool culture that shows overlap of distinct cultures––Acheulean hand-axe and Levallois technologies––for over 200,000 years, unlike in other regions of the world^60,61^. These findings raise important questions about the dispersal and settlement of humans outside Africa: Did the range of Neanderthals and Denisovans extend to South Asia? Did modern humans encounter Neanderthals, and to some extent Denisovans, further east in Eurasia rather than the Middle East as widely believed? These observations call for a re-evaluation of models of human origins, for both modern human and archaic hominins, in light of the complex diversity in India.

## Methods

### Samples

We generated 2,762 high-coverage genomes as part of this project. These samples are a subset of the Longitudinal Aging Study in India (LASI) and are part of the Harmonized Diagnostic Assessment of Dementia of LASI (LASI-DAD)^8^ (https://lasi-dad.org, doi.org/10.25549/5hhx-s820). Participants consented to give venous blood samples (VBS) for genomics analysis. They also have consented to detailed cognitive assessment and informational interviews. Details on the sequenced individuals and metadata (i.e., sampling location, sex, language, caste etc.) can be found in Supplementary Note S1.

### Whole genome sequencing, variant calling and filtering

Whole-genome sequencing libraries were processed using a PCR-free library preparation and sequenced on Illumina HiSeq X Ten machines at Medgenome, Bangalore, India. The samples were sequenced using 100 base pair paired-end sequencing. The raw sequence reads (fastq) from Medgenome were sent to the Genome Center for Alzheimer’s Disease (GCAD) at the University of Pennsylvania for genome mapping to the human reference genome (build GRCh38/hg38). We used Variant Calling Pipeline and data management tool (VCPA) developed by GCAD in collaboration with Alzheimer’s Disease Sequencing Project (ADSP) to call variants in a uniform way across other studies that are part of ADSP. The pipeline uses best practices from Genome Analysis Took lit (GATK) to call variants. Details of the data processing are described in Supplementary Note S2. Overall, a total of 2,679 LASI-DAD samples passed sequencing metrics and quality control checks. Details of quality checks are described in Supplementary Note 2.

### Identification of first-degree relative pairs

We applied KING (v2.3.0)^62^ and the “--ibdseg” option to identify first degree relatives. Following software guidelines, we applied the following filters: sample pairs without any long IBD segments (>10Mb) were excluded and short IBD segments (<3Mb) were not utilized to estimate the proportion of IBD sharing between two individuals. Parent-offspring pairs share 50% of their genomes and siblings may share between 38-65% of their genome inherited IBD^63^. Thus, we use a minimum cutoff of 38% to identify first-degree relatives and consequently we flag 64 pairs of individuals. For each pair of first degree relatives, we removed the individual with the larger amount of missing data. In total, we removed 59 individuals (see details in Supplementary Note S2), leaving 2,620 individuals that were used for most downstream analyses.

### Population structure analysis

To learn about the population history of India and compare it to worldwide populations, we combined the LASI-DAD dataset with other published genomic datasets including present-day (1000G^11^, GenomeAsia^6^) and ancient DNA samples (Allen Ancient DNA Resource (AADR) v54 ^64^). GenomeAsia and AADR are available in hg19/GRCh37, we performed liftover to hg38/GRCH38 using liftOver (https://liftover.broadinstitute.org/). Then, we merged the datasets using *mergeit* (with ‘strandcheck: YES’) from the EIGENSOFT package (v7.2.1)^65,66^ which generates an intersection of the SNPs in the different datasets, keeping only variants present in all datasets. The number of individuals and variants for each merged dataset and the analyses they are used in are reported in Table S4.1.

### Principal component analysis (PCA) and ADMIXTURE

To perform PCA and *ADMIXTURE*, we excluded SNPs in linkage disequilibrium (LD) using PLINK with the option ‘--indep-pairwise 50 10 0.5’ that removes, one variant in each pair of SNPs in a window of 50 SNPs, if the LD is greater than 0.5. We further excluded variants with a MAF<0.05. We performed PCA using *smartpca* from the EIGENSOFT package (v7.2.1)^65,66^. We also applied unsupervised hierarchical clustering of individuals using the maximum likelihood method implemented in the ADMIXTURE software (v1.3.0)^13^. Following program documentation, we varied the number of clusters (K) between 2–6 and performed cross validation ten times (option: --cv=10). We stopped the algorithm when the change in log-likelihood between iterations was less than 0.1 (option: -C 0.1).

### qpAdm

We used the qpAdm^14,22^ package in ADMIXTOOLS (v7.0.2) to identify the best fitting model and estimate ancestry proportions in a population of interest that is modeled as a mixture of *n* ‘reference’ populations using a set of ‘Outgroup’ populations (reference (*left*) and outgroup (*right*)) populations for each analysis are listed in Supplementary Note S4). We set the parameters as ‘allsnps: NO’ and ‘details: YES’, which reports a normally distributed *Z* score for the fitted model. We computed coefficient estimations, standard deviations and p-values through block jackknife resampling. We considered a model to be a good fit if *p*-value > 0.01 and all coefficients are positive.

### ALDER

To infer the date of East Asian admixture and ancestry proportion in Bengalis (East of India), we used ALDER (v1.04)^18^. We used the ‘one-reference’ model (*runmode*: 1) with East Asians (*CHB.DG* from AADR v54) as the reference population with the following parameters: *binsize*: 0.001 Morgans; *maximum distance*: 1.0 Morgans; *zdipcorrmode*: YES; *jackknife*: YES. To convert the dates of admixture from generations to years, we assume the mean human generation time was 28 years^67^.

### IBD and HBD sharing

We identified IBD and HBD segments using hap-IBD^29^ with the following parameters: min-seed: 0.5; max-gap: 1000; min-extend: 0.5; min-output: 1.0; min-markers: 100; min-mac: 2; nthreads: 1. We used the HapMap genetic maps. To minimize false positives, we only considered shared segments with length greater at 2cM. Then, we filtered out segments that overlapped centromeres (using the GRCh38/hg38 annotation from genome.ucsc.edu/cgi-bin/hgTables). To infer the putative degree of relatedness between two individuals, we computed the total IBD sharing for kth degree cousins using *2G(1/2)*^2^(k+1), where *G* = 6,782cM is the total diploid autosomal genome size^68^ and *k* represents the degree of cousin relationship^69^. We note, however, the expected values assume a random mating population and a history of founder events could lead to increased genomic sharing and thus these values should be interpreted with caution.

### Loss of function (LoF)/missense variants

To quantify the mutational burden in India, we used the Variant Effect Predictor (VEP; version 105)^70^ and LOFTEE (v1.0.3)^9^ to identify missense and predicted loss-of-function (pLoF) single nucleotide variants (SNVs). VEP annotates each SNV according to its functional effect on gene transcripts. We used GENCODE^71^ as the transcript annotation reference and focused our analysis on the most severe functional effect per SNV across different transcripts. Besides the functional annotations directly obtained from VEP, we identified pLoF SNVs by coupling VEP with LOFTEE^9^. LOFTEE further assesses stop-gained, splice-site-disrupting, or frameshift SNVs identified by VEP and implements a set of filters to infer if a SNV should be considered a pLoF.We intersect the list of pLoF/missense variants with the RefSeq database^34^ and the ClinVar database^36^ (data release of 2023-12-17) to infer the nearest gene and any disease associations respectively. We consider ClinVar status for variants with a review of at least two stars. Information for each of the pLoF/missense variants is available in Extended Data Table S1.

### Inference of archaic ancestry

To learn about the genomic landscape and regional variation in archaic ancestry in Indians and compare it to worldwide populations, we applied *hmmix*^72^ to 2,679 phased individuals from India (we retain first-degree relatives (except offspring of trios) as they may have archaic ancestry in different positions). This method uses an outgroup who have negligible amount of archaic ancestry. We used 426 individuals from the 1000G^11^ including Yoruba in Ibadan, Nigeria, Mende in Sierra Leone (YRI), Esan in Nigeria (ESN) and 64 Africans from HGDP^73^, who have less than 1% West Eurasian admixture, including Bantu South Africa, Biaka Pygmy, Mbuti Pygmy, San and Yoruba. We estimated the number of callable sites, the single-nucleotide polymorphism density (as a proxy for per-window mutation rate) and the number of private variants with respect to the outgroup individuals in 1-kb windows across the genome. We obtained regions identified as ‘archaic’ and compared them to the four published high coverage archaic genomes––Altai Neanderthal^39^, Chagyrskaya Neanderthal^40^, Vindija Neanderthal^41^ and Altai Denisovan^42^to identify the source of the archaic ancestry (see details in Supplementary Note S6). We further compared archaic segments previously published for 27,566 individuals from Iceland^44^ that were also inferred using *hmmix*. The datasets and number of individuals per population used for the analysis of archaic ancestry in non-Africans are reported in Table S6.1.

### Inferring the timing of Out-of-African migration (OOA)

We infer the minimum coalescence time for non-African individuals with Sub-Saharan African individuals from the outgroup (*n=490*). Any systematic difference might indicate a difference in the timing of the out of Africa migration (OOA) for different populations. *hmmix* classifies the genome into ‘modern human’ and ‘archaic’ states. The emission parameters for the human state is informative about the minimum coalescence time between non-African individuals and Sub-Saharan African individuals.

We merge HGDP, 1000G and LASI-DAD dataset and subset to SNPs found in 1240K array^64^ and use ADMIXTURE (v1.3.0)^13^ in unsupervised-mode (*k=2*) to estimate Sub-Saharan ancestry. We remove all individuals with > 1% Sub-Saharan ancestry to minimize the effect of recent gene-flow on the minimum coalescence time estimate. To minimize the effect of archaic ancestry on the emission parameters for the human state we correct for the amount of high confidence archaic segments (posterior probability > 0.9). To compare coalescence times between HGDP and LASI-DAD we correct for phasing drop-out rate and the removal of multi-allelic sites. Assuming a mutation rate of 0.45e-9 ^58^ the emission parameter for the human state can be converted into a coalescence time.

### Ethics statement

The Longitudinal Aging Study in India (LASI, https://lasi-india.org) is a joint effort by the Harvard T.H. Chan School of Public Health (HSPH), the International Institute for Population Sciences (IIPS) in India, and the University of Southern California (USC). Longitudinal Aging Study in India - Diagnostic Assessment of Dementia (LASI-DAD) is an in-depth study of late-life cognition and dementia, drawing a subsample of the LASI. Principal Investigators teams are located at USC and All India Institute Of Medical Sciences (AIIMS). Interviews and sampling were conducted in collaboration with the Regional Geriatric Centers (RGCs) at the respondents homes or at the participating hospitals, reaching out to both rural and urban areas in 18 states across the country, representing the nation-wide diversity. The AIIMs in New Delhi, India coordinated field work across RGCs to recruit interviewers and provide training and logistical support to uniformly perform phenotyping across diverse regions across India. The lists of partner hospitals and field team members are accessible at https://lasi-dad.org/teams.

Ethics approval was obtained from the Indian Council of Medical Research and all collaborating institutions. The study was approved by Institutional Review Boards at the University of Southern California and the University of Michigan. Informed consent was obtained from all participants or their legal representative. As most individuals in this study are 60 years or older, some participants were cognitively impaired, in which case we obtained informed consent from a close family member, such as a spouse or adult child who was the legal representative of the participant. The consent materials were translated into as many local languages as necessary. Informed consent and interviews were collected and conducted in the respondent’s language. If the participant was unable to read the consent forms, the interviewer would verbally relay the information in the consent form. Participants who were unable to sign the consent forms had the option to use their thumb impression in place of a signature (a common practice in India).

DNA extraction and whole genome sequencing was performed at MedGenome, Bangalore, India. Anonymised data is available for the larger research community through a secured website hosted by the Gateway to Global Aging Data platform. Research findings from the LASI-DAD team are disseminated through journal publications and presentations at professional conferences.

## Data availability

All data is available through the National Institute on Aging Genetics of Alzheimer’s Disease Data Storage Site (NIAGADS) under the accession NG00148.v1. The post-qc vcf file is distributed by the Genome Center for Alzheimer’s Disease (GCAD) at the University of Pennsylvania and can be obtained by following the data request instructions available: https://dss.niagads.org/documentation/data-application-and-submission/application-instructions/

## Acknowledgements

We thank Vagheesh Narasimhan, Michael Frachetti and J. Mark Kenoyer for helpful discussion about the archaeological connections between Sarazm, Iran and South Asia. We also thank members of the LASI-DAD advising committee and Moorjani lab for helpful feedback on the analysis and results throughout the project. We thank Yulin Zhang, Elena Zavala, Monty Slatkin, Ben Peter, Vagheesh Narasimhan and Shai Carmi for helpful comments on the manuscript. The Longitudinal Aging Study in India, Diagnostic Assessment of Dementia data (https://lasi-dad.org, doi.org/10.25549/5hhx-s820) is sponsored by the National Institute on Aging (grant number R01AG051125, RF1AG055273, U01AG065958) and is conducted by the University of Southern California. PM was supported by U01AG065958 and NIH R35GM142978.

## Extended Data Figures

**Extended Data Figure 1.**
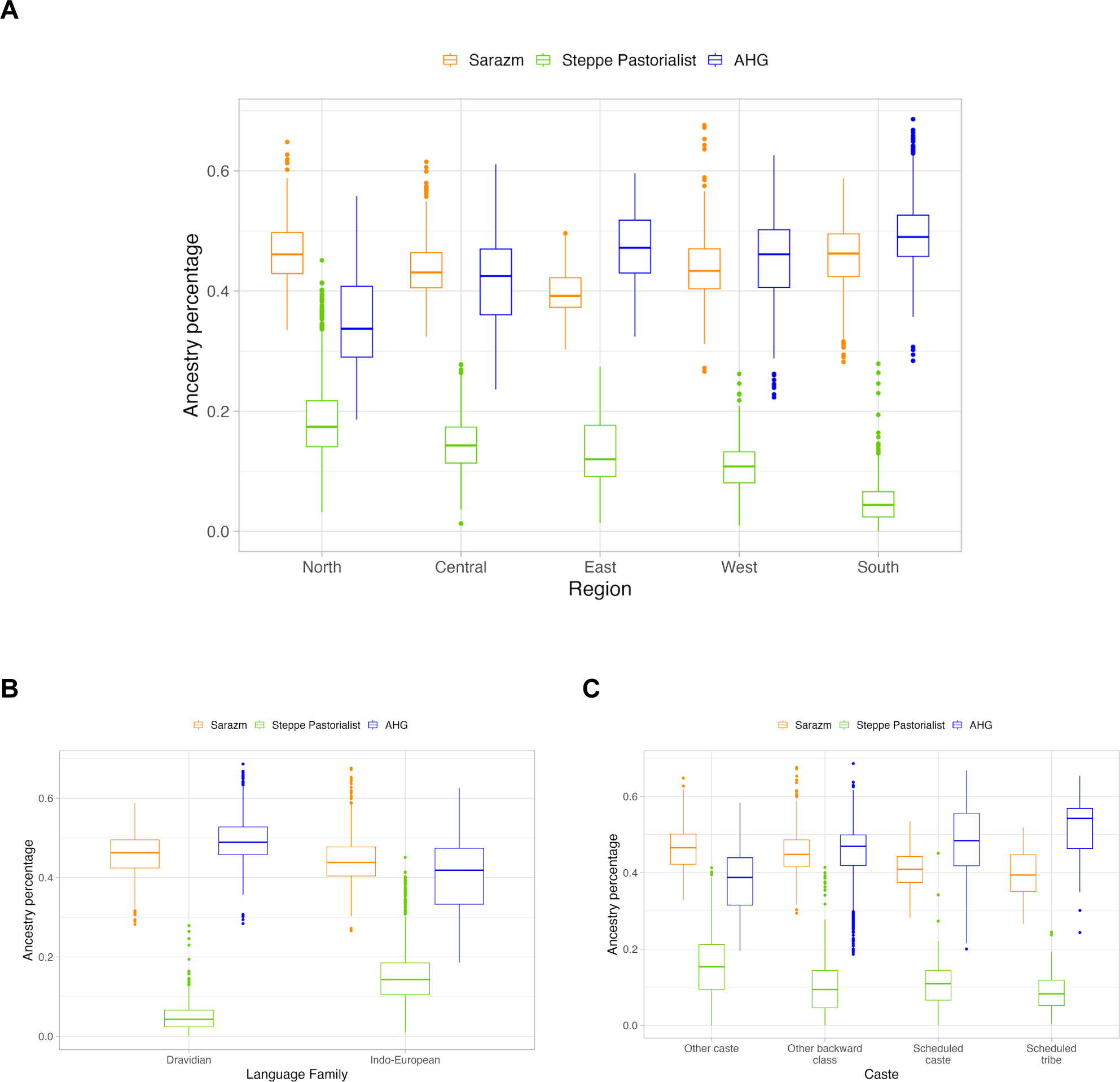
Ancestral population-related coefficients using the revised model. Inferred coefficients based on qpAdm using the three-way model with Sarazm_EN, Central_Steppe_MLBA and AHG-related groups shown by (A) region, (B) language family and (C) caste group. We show only results for 1,942 individuals for whom the three-way model was a good fit (*p*-value > 0.01 and inferred ancestry proportions were non-negative).

**Extended Data Figure 2.**
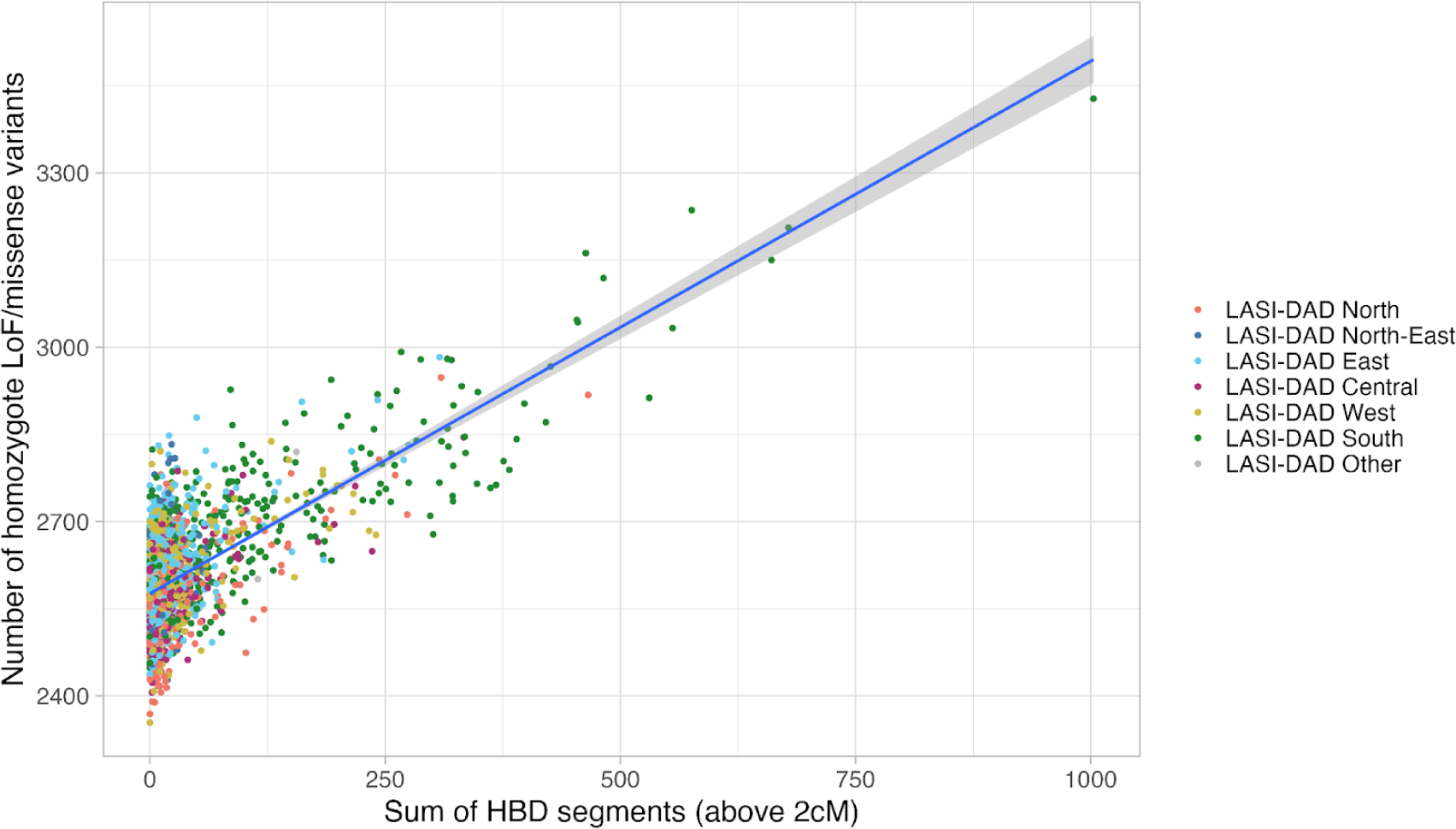
Relationship between the number of homozygous derived missense/pLoFs and the total sum of HBD segments per individual. Individuals are colored by region of birth. We fit a regression using generalized linear model (glm) and obtain the following fit: y = 2576 + 0.916*x.

**Extended Data Figure 3.**
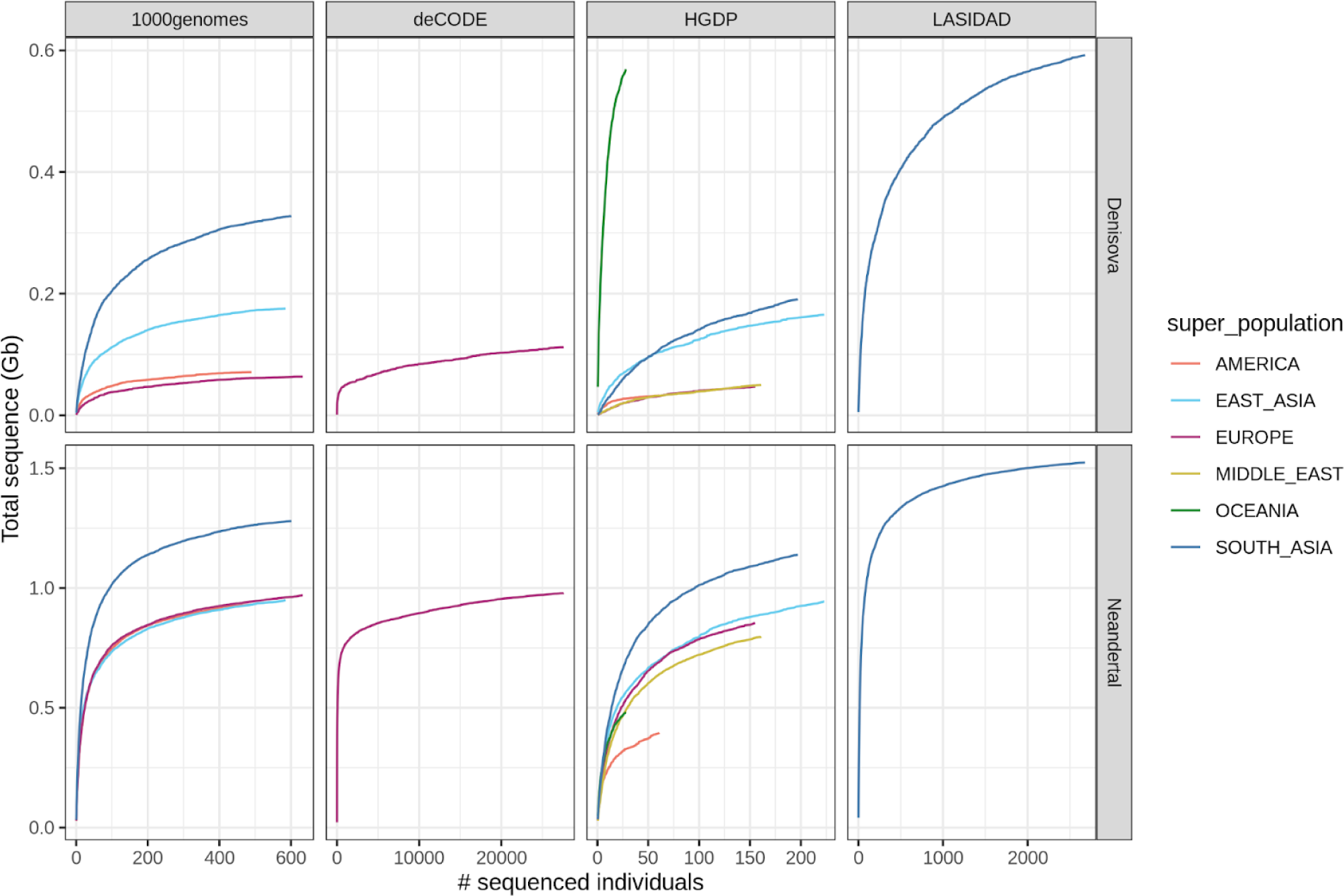
Amount of unique archaic sequence in worldwide populations. For Denisovan (top) and Neanderthal (bottom) as a function of the analyzed number of individuals in four different datasets (at a posterior cutoff of 0.9).

**Extended Data Figure 4.**
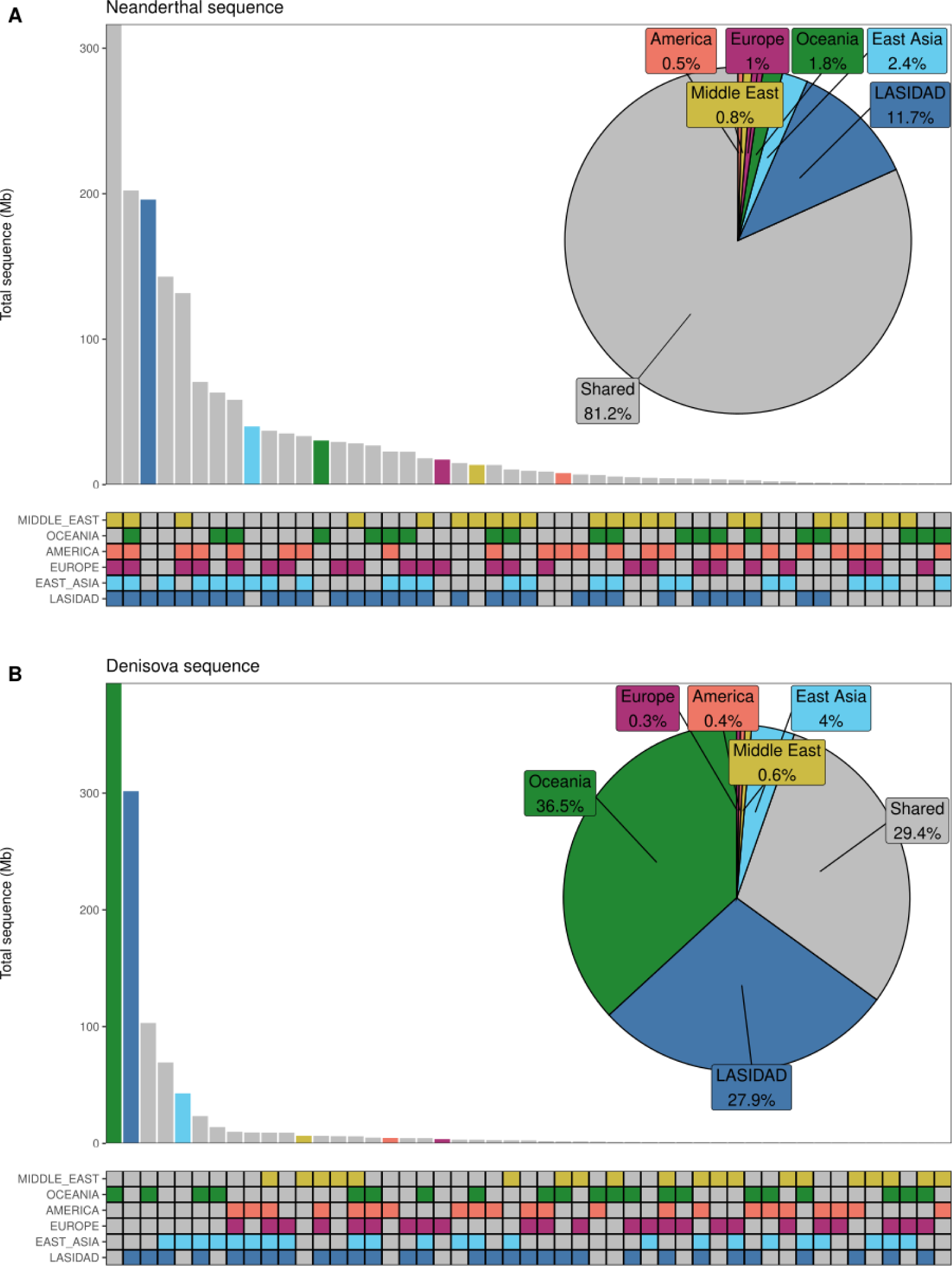
Sharing of Neanderthal and Denisova sequence. **A)** Upset plot of Neanderthal sequence found at a posterior probability cutoff at 0.8 (y-axis) that is shared between any combinations of regions (x-axis). Sequence that is unique to one region is colored according to which population it is found in while sequence that is found in at least 2 populations (shared) is colored in grey. In the pie chart the total amount of shared and unique are denoted in percent. **B)** same as **A)** but for Denisovan sequence.

**Extended Data Figure 5.**
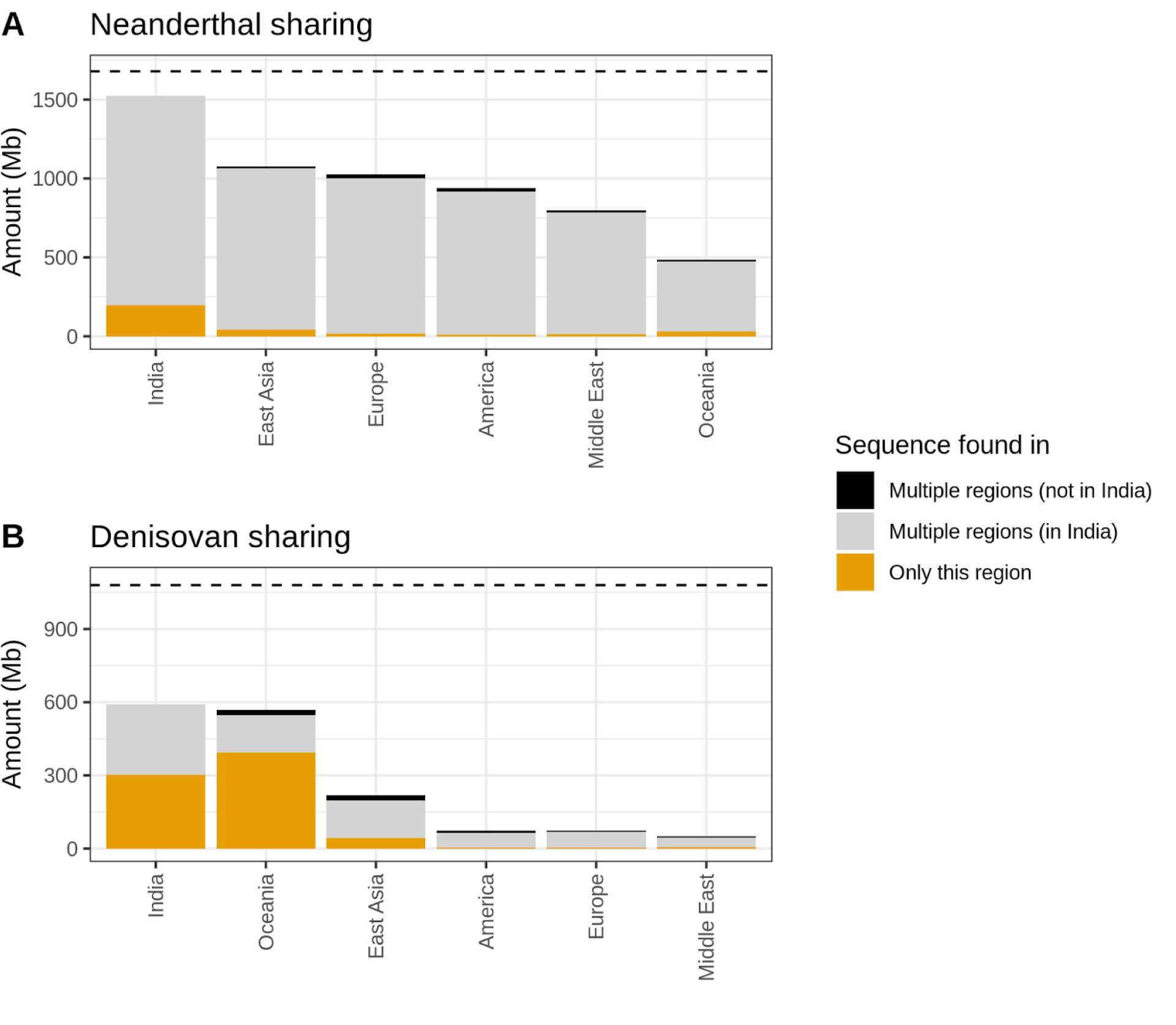
Neanderthal and Denisova sequence found in world-wide regions. **A)** Amount of Neanderthal sequence found at a posterior probability cutoff at 0.8 (y-axis) that is unique to any region, found in multiple regions (at least two) where one includes India (LASI-DAD dataset) or found in multiple regions (at least two) where India is not included. **B)** same as **A)** but for Denisovan sequence. Horizontal lines indicate the total length of the assembled Neanderthal and Denisova genome using LASI-DAD, HGDP and 1000G datasets.

