## Supplementary information for "50,000 years of Evolutionary History of India: Insights from ∼2,700 Whole Genome Sequences"

### Table of Contents

**Supplementary Information** 1
**Extended Data Tables** 3
**Supplementary Note 1: Samples, Sequencing and Variant Calling** 4
Sample Description 4
Whole genome sequencing 6
Genome Mapping and Variant Calling 6
Table S1.2. Genome mapping and variant calling tools / softwares. 7
**Supplementary Note 2: Data filtering and quality control** 8
WGS validation study 8
LASI-DAD VCF Quality Checks 9
VCF STEP 1 Quality Checks 9
VCF STEP 2 Quality Checks 10
Genotype-level 10
Variant-level 10
Sample-level 11
Comparison of summary statistics to published whole genome sequences 11
Per sample statistics 12
**Supplementary Note 3: Genome Phasing** 15
**Supplementary Note 4: Population structure and admixture** 18
Principal Component Analysis (PCA) 18
ADMIXTURE 22
Ancestry composition of India 25
Model of ancestry for individuals on the Indian cline 25
Source of Iranian farmer-related ancestry in India 26
Model of ancestry for individuals that fall outside the Indian cline 29
Population structure within the Sarazm_EN individuals 31
Estimating the genome-wide ancestry proportions 34
Population structure in West Bengal 37
**Supplementary Note 5: Founder events and consanguinity in India** 38
Distribution of HBD in India 38
Distribution of IBD sharing in LASI-DAD and 1000G SAS 41
**Supplementary Note 6: Inference of archaic ancestry** 45
HMM Training 47
HMM Decoding 49
Impact of recent demographic history on shaping patterns of archaic ancestry in India 62

Inferring the number of pulses of Neanderthal and Denisovan gene flow in India 64
Denisovan or Denisovan-related ancestry components 65
Neanderthal or Neanderthal-related ancestry components 67
**Supplementary Note 7: Genome-wide distribution of archaic ancestry in India 70**
Regions with high frequency of archaic ancestry segments 70
Gene Ontology (GO) analysis 71
Regions with Neanderthal-specific or Denisovan-specific variants 72
BTNL2 72
Population Branch Statistics Analysis 76
GO enrichment analysis of Indian-specific regions of archaic ancestry 77
Regions depleted for archaic ancestry (‘archaic ancestry deserts’) 81
**Supplementary Note 8: Simulations to evaluate the genomic distribution of archaic ancestry 84**
Simulation Scenarios 84
Distribution of archaic ancestry in simulations 87
Distribution of archaic ancestry in worldwide populations 89
**Supplementary Note 9: Revising the Timing the Out of Africa migration to India 92**
Model for estimating emission parameter for modern human state 92
**References 103**

### Extended Data Tables

**Extended Data Table S1: Summary loss of function (LOF) and missense variants in LASI-DAD.** (A) For each variant, we report their position, their frequency in LASI-DAD, the number of homozygous genotypes, the genes they are located in and the medical relevance of the genes, if available. (B) We report per gene, the number of LoF and missense variants overlapping the gene.

**Extended Data Table S2: Total amount of reconstructed Neanderthal and Denisovan genome.** Amount of reconstructed Neanderthal and Denisovan sequence for each population and dataset. We report the number of archaic overlapping clusters of segmetns and how much sequence they span. We report the numbers for phased and unphased data with posterior probability cutoff at 0.5, 0.8 and 0.9.

**Extended Data Table S3: List of Genes enriched in Neanderthal and Denisovan ancestry (High frequency and PBS).** We show genes enriched in Neanderthal and Densioan ancestry in South Asians, based on high frequency (A-B) or PBS analysis (C-D).

**Extended Data Table S4: GO enrichment analysis for archaic ancestry segments in India.** We show high frequency segments of Neanderthal (A) and Denisovan (B) ancestries and archaic enriched segments in Indians compared to European and East Asians (PBS analysis) for Neanderthal (C) and Denisovan (D) ancestries.

**Extended Data Table S5: Positions of archaic deserts in HGPD, 1000G and LASI-DAD.** Chromosome positions (in hg38) for Neanderthal ancestry deserts (A) and Denisovan ancestry deserts (B). For all populations except Oceania and the Middle East we combine the individuals from 1000G and HGDP.

### Supplementary Note 1: Samples, Sequencing and Variant Calling

#### Sample Description

The Longitudinal Aging Study in India (LASI) is a nationally representative sample of over 72,000 Indian adults of 45 years or older. A subset of ~4,000 individuals are part of the Harmonized Diagnostic Assessment of Dementia of LASI (LASI-DAD) (https://lasi-dad.org) that have consented to detailed cognitive assessment, informational interviews and genomics analysis. These samples were sampled from 18 states, with sample sizes approximately proportional to the census size of the state. A subset of 2,762 participants consented to give venous blood samples (VBS) for genomics analysis. The All India Institute of Medical Sciences (AIIMs) in New Delhi, India coordinated field work across 12 regional centers (RCs) to recruit interviewers and provide training and logistical support to uniformly perform phenotyping across diverse regions across India. The detailed sample collection protocol and list of collaborating institutions can be found in Lee et al. 2019[1].

Venous blood samples were collected by certified and trained phlebotomists from Metropolis laboratory, India who drew ~17 ml of VBS from each study participant. All specimens were shipped to the Metropolis laboratory in Delhi within approximately 24 hours via a cold chain (−20 °C for plasma and 4°C for other specimens) where the lab work was completed for 33 assays, including complete blood count, hemoglobin (HbA1c), Thyroid-stimulating hormone (TSH), lipid profile panel, lipoprotein A, Vitamin B12, Vitamin D, folic acid, homocysteine, high sensitivity C-Reactive Protein (CRP) and Natriuretic peptide tests (NT-proBNP). The blood samples were then shipped to MedGenome, Bangalore, India for DNA extraction and whole genome sequencing.

We list here the sample size for each region and state of origin, sex, mother tongue and caste affiliation. Sampling locations are represented on Fig 1A.

**Table S1.1 Description and sample size by Region, Sex, Linguistic and Caste Affiliation.**

| **A. Region** - including State / country if not India. | | | | | | |
|---|---|---|---|---|---|---|
| **Central** (*n=382*) | **East** (*n=542*) | **North** (*n=575*) | **North-East** (*n=75*) | **South** (*n=726*) | **West** (n=387) | **Other*** (*n=49*) |
| Uttar Pradesh (*n=239*) | Odisha (*n=212*) | Rajasthan (*n=173*) | Assam (*n=74*) | Kerala (*n=266*) | Gujarat (*n=223*) | Bangladesh (*n=29*) |
| Madhya Pradesh (*n=74*) | West Bengal (*n=171*) | Haryana (*n=150*) | Meghalaya (*n=1*) | Tamil Nadu (*n=170*) | Maharashtra (*n=164*) | Pakistan (*n=14*) |

| | | | | | | |
|---|---|---|---|---|---|---|
| Uttarakhand (*n=69*) | Bihar (*n=157*) | Punjab (*n=115*) | | Telangana (*n=162*) | | South Africa (*n=2*) |
| | Jharkhand (*n=2*) | Jammu & Kashmir (*n=93*) | | Karnataka (*n=117*) | | Burma/ Myanmar (*n=1*) |
| | | Delhi (*n= 42*) | | Andhra Pradesh (*n=11*) | | Missing (*n=3*) |
| | | Himachal Pradesh (*n=2*) | | | | |

Note: * includes country if other than India

| **B. Sex** | | |
|---|---|---|
| Female (*n=1443*) | Male (*n=1293*) | Not specified (*n=26*) |

| **C. Linguistic affiliation -** including language family/ native language (mother tongue) shown. | | | |
|---|---|---|---|
| **Dravidian** (*n=687*) | **Indo-European** (*n=2037*) | **Tibeto-Burman** (*n=8*) | **Not specified** (*n*=30) |
| Malayalam (*n=264*) | Hindi (*n=796*) | Bodo (*n=8*) | |
| Telugu (*n=183*) | Gujarati (*n=227*) | | |
| Tamil (*n=161*) | Oriya (*n=216*) | | |
| Kannada (*n=79*) | Bengali (*n=203*) | | |
| | Marathi (*n=141*) | | |
| | Punjabi (*n=129*) | | |
| | Rajasthani (*n=98*) | | |
| | Kashmiri (*n=93*) | | |
| | Assamese (*n=51*) | | |
| | Urdu (*n=42*) | | |
| | Bhojpuri (*n=12*) | | |

| | |
|---|---|
| | Adivasi (n=7) |
| | Konkani (*n=5*) |
| | Irani (*n=4*) |
| | Maithili (*n=3*) |
| | Sindhi (*n=3*) |
| | Bajjika (*n=2*) |
| | Haryanavi (*n=1*) |
| | Nepali (*n=2*) |
| | Marwari (*n=1*) |
| | English (n=1) |

| **D. Caste Affiliation** | | | | |
|---|---|---|---|---|
| Scheduled Caste (*n=498*) | Scheduled Tribe (*n=111*) | Other backward class (OBC) (*n=1187*) | Other caste* (*n=927*) | Not Specified (*n=39*) |

Note: *Other* implies that individuals self-reported as not belonging to either Scheduled Caste, Scheduled Tribe or OBC.

#### Whole genome sequencing

We performed high coverage (~30x) whole genome sequencing for 2,762 LASI-DAD individuals, including 22 trios, using Illumina HiSeq X Ten machines at Medgenome, Bangalore, India. All samples were processed using a PCR-free library preparation and sequencing protocol. The samples were sequenced using 100 base pair paired-end sequencing.

#### Genome Mapping and Variant Calling

The raw sequence reads (*fastq*) from Medgenome were sent to the Genome Center for Alzheimer's Disease (GCAD) at the University of Pennsylvania for genome mapping and variant calling. We used the sequencing processing pipeline and data management tool, called Variant Calling Pipeline and data management tool (VCPA), a workflow co-developed by the GCAD in collaboration with Alzheimer's Disease Sequencing Project (ADSP) to process FASTQ files to variant calls in a uniform way across studies that are part of ADSP. The pipeline is optimized in an Amazon cloud environment and includes all steps from aligning raw sequence reads to variant calling using recommendations by the Genome Analysis Took lit (GATK) best practices. VCPA also has a tracking database to store >100 quality metrics collected during data production, such as file size, mapping percentage, depth coverage, and quality and counts of called variants. To ensure high quality data and minimize false positives, various QC metrics are applied, and filtering details are tracked through the platform. VCPA has shown high concordance for variant calls (0.995) for WGS technical replicates sequenced from multiple different platforms at different sequencing centers. Details of the data processing pipeline are available at Leung et al. 2019[2], which we briefly summarize below.

Raw sequence reads were mapped to GRCh38/hg38 using BWA-mem (v0.7.15)[3] and duplicate reads were marked by BamUtil (v1.0.13)[4]. Next, BAM files were processed by Samblaster[5] (adding MC and MQ tags to pair-end reads) and sorted by genomic coordinates using SAMtools[6]. Finally, coverage statistics are computed using Sambamba[7].

We implemented the Genome Analysis Toolkit (GATK) Best Practices (https://software.broadinstitute.org/gatk/best-practices/) for variant calling and annotation for single nucleotide variants (SNVs) and indels, and generated genotype call files in the genomic Variant Call Format (gVCF) format for each sample individually[8,9]. Quality metrics of called variants were computed using GATK, including: (1) comparison of genotype concordance for SNV with previously generated single nucleotide polymorphism (SNP) genotypes from LASI-DAD[10]; (2) sex check using genetic data to identify possible sample swaps or misreporting; (3) contamination check for possible sample swaps; and (4) relatedness check to confirm known relationships, identify unknown duplicates, and assess potential cryptic relatedness. Overall, a total of 2,685 LASI-DAD samples passed sequencing metrics and quality control checks (including 6 technical replicates sequenced with LASI-DAD). Details of quality checks are described in Supplementary Note 2. The gVCFs were then combined into a VCF file via joint genotype calling using GATK4.1.1.

**Table S1.2. Genome mapping and variant calling tools / softwares.**

| Software | Link | Reference |
|---|---|---|
| BWA-mem (v0.7.15) | https://github.com/lh3/bwa | ([3]) |
| BamUtil (v1.0.13) | https://genome.sph.umich.edu/wiki/BamUtil | ([4]) |
| samtools | https://github.com/samtools/ | ([6]) |
| samblaster | https://github.com/GregoryFaust/samblaster | ([5]) |
| sambamba | https://lomereiter.github.io/sambamba/ | ([7]) |
| GATK4.1.1 | https://gatk.broadinstitute.org/hc/en-us/sections/360007336992-4-0-1-1 | ([8]) |

### Supplementary Note 2: Data filtering and quality control

#### WGS validation study

Before embarking on the whole genome sequencing for 2,762 samples that are part of LASI-DAD, we conducted a small pilot study to assess the quality of data generated by MedGenome, India. We performed three validation experiments, including

(a) comparison of whole genome sequences (WGS) of one European individual (NA12878) from the 1000 Genomes Project (1000G)[11] sequenced at MedGenome, Inc., the Broad Institute, Cambridge, USA (henceforth, Broad) and the platinum genome (gold-standard) provided by Illumina,

(b) comparison of WGS for a Finnish trio at both MedGenome and Broad, and

(c) comparison of our entire pipeline (sample to variant calling) for one South Asian individual. In these comparisons, we used a single individual or family as we reasoned that the variability across samples within the same sequencing center is typically small, while the variability across centers is expected to be higher[12].

All samples included in the analysis were sequenced using HiSeq X Ten machines at MedGenome or Broad, unless otherwise specified.

Raw sequences were processed using the same genome mapping and variant calling pipeline as recommended by Genome Analysis Toolkit (GATK) best practices (https://software.broadinstitute.org/gatk/best-practices/) (Supplementary Note 1). This implies that most of the observed discordances in coverage or variant calls between data generated at Medgenome or Broad stem from differences in quality of sequencing and not from the analysis pipelines. We applied 6 different quality control (QC) levels, from less to more stringent that includes

(i) **Base QC**: we removed low complexity and decoy regions (24,974,202 bp),

(ii) **Variant Quality Score Recalibration (VQSR) TR1**: Base QC + removed variants in VQSR tranche (TR) 99.80 to 99.90,

(iii) **VQSR TR2**: VQSR TR1+ removed variants in VQSR tranche 99.80 to 99.90 and 99.90 to 99.95,

(iv) **VQSR PASS**: Base QC + kept only PASS variants according to VQSR,

(v) **HIGH QC1**: VQSR PASS + set to missing genotypes with DP < 20 or GQ< 20 or allele balance greater than 0.2 or 0.8 and removed variants with call rate < 0.80, and

(vi) **HIGH QC2**: HIGH QC1 + removed variants with call rate < 0.95.

**In the first experiment**, we compared the genome sequences for the 1000G European individual, NA12878, generated at MedGenome, Broad to the platinum genome (gold-standard) provided by Illumina. We observed that the sensitivity (measured as the proportion of the variants with alternate alleles (heterozygous-alt or homozygous-alt) that match the gold standard) ranged between 88–91% depending on the QC filters applied and was virtually the same between the sample sequenced at MedGenome or Broad. The false discovery rate (measured as the proportion of variants with alternate alleles in the gold standard that were inaccurately called) was between 0.003–0.1%, slightly higher for the sample sequenced at Broad compared to MedGenome.

**In the second experiment**, we compared the quality of the genome sequences for a Finnish trio sequenced at MedGenome and Broad, by counting the number of Mendelian errors in the trio. We considered Mendelian errors as all those variants that were heterozygous in the proband, but not observed in either parent. We expect ~70-100 de novo mutations (dnms) per generation per trio[13], while the remaining are likely to be sequencing errors. We observed 165 putative dnms in the trio sequenced at Broad and 101 putative dnms in the trio sequenced at MedGenome, after applying stringent QC in both cases. The estimate from MedGenome is close to the theoretical expectation based on large-scale pedigree studies[13], indicating high data quality of the sequences generated at MedGenome.

**In the third experiment**, we sequenced DNA extracted from the same fresh venous blood sample from a South Asian individual both at MedGenome and Broad and evaluated the concordance in the variant calls, after sequencing and variant calling. Both samples were sequenced to 30x coverage. We obtained a high concordance in variant calls of 99.98% when strict QC procedures were used. Similar rate was seen even for variants outside the accessible genome (e.g., 1000G strict mask) typically used for whole genome sequence analysis.

Overall, the three experiments performed in this validation study support the high quality of sequence data generated at MedGenome, in terms of their low false discovery rate, high concordance with widely-used sequencing centers (like Broad) and low number of sequencing errors. Following these results, all LASI-DAD samples were sent to Medgenome, India for genome sequencing.

#### LASI-DAD VCF Quality Checks

After variant calling, we performed stringent checks to ensure the quality of the data in two main steps with increasing levels of stringency.

#### VCF STEP 1 Quality Checks

1. **Sample ID concordance check.** For a subset of LASI-DAD samples (n=960), we have both genotype array data (on Illumina Infinium Global Screening Array-24 v2.0 BeadChip)[10] and whole genome sequences (Supplementary Note S1). We compared the genotypes for the overlapping variants in the array data and WGS (determined by matching chromosomal positions in hg38). The goal of this analysis was to ensure that samples and IDs match throughout the data management and calling processes. Only 2 samples were dropped because of low concordance, the remaining samples have a concordance of > 0.91.
2. **Sex check:** Sex checks were performed using BCFtools[14], VCFtools[15], and PLINK[16] with the following steps:
    1. Use BCFtools to convert chrX vcf into plink format
    2. Filter out pseudoautosomal region (PAR)[17]

3. Filter out SNVs with MAF < 0.05 and run *impute-sex*
4. Run *sex-check* using PLINK for comparison

We find that the results of 'impute-sex' in BCFtools and '--sex-check' in PLINK were very similar, with or without filtering on minor allele frequencies (MAFs) or excluding variants in the pseudoautosomal region (PAR). 28 samples were excluded based on this analysis as they had discordant self-reported sex and inferred sex based on genomic data.

4. **Contamination check:** Sample-specific contamination was checked by using *VerifyBamID*[18] to calculate the concordance estimate between the genotype array data[10] and the hg38-mapped BAM file (Supplementary Note S1). This approach can help to identify potential sample contamination or swapping using the GWAS-BAM contamination estimate. The 'FREEMIX' modeling approach was used in this analysis. Following VerifyBamID recommendation, we considered a sample as potentially contaminated if the FREEMIX value is >0.05, 3 samples were removed based on this analysis.
5. **Relatedness check:** Relatedness check was performed in PLINK with the '--genome' command, which computes pairwise identity by state (IBS)/ identity by descent (IBD) allele sharing between all samples. Variant calls for all autosomes were converted to PLINK binary (BED/BIM/FAM) format files, combined, and then only SNVs common to both Affymetrix and Illumina genotyping arrays were retained (~21k SNVs in total). For every pair of individuals, we measured PI_HAT [PI_HAT = P(IBD=2) + 0.5*P(IBD=1)], a measure of relatedness. All pairs with PI_HAT > 0.4 were evaluated for known relatedness. 5 samples were unexpected duplicates and were removed from the analysis.

**VCF STEP 2 Quality Checks**

To ensure high reliability of the genotype calls, we applied additional filters and quality checks at each level (genotype, variant, and sample). All QC flags and filters were applied uniformly across all samples, regardless of sampling location or sex.

*Genotype-level*

Following STEP 1 Quality Checks, genotype-level QC was applied to individual genotypes. Each genotype was evaluated and set to missing ("./.") if either or both read depth ("DP") was less than 10 (DP < 10) or genotype quality ("GQ") score was less than 20 (GQ < 20). All these flagged genotypes were excluded from subsequent QC steps, except for estimation of variant-level averaged depth ("AverageReadDepth") in variant-level QC.

*Variant-level*

Variant-level QC was applied to all variants in the VCF. Filters were applied in the following order:

(i) Variants in GATK low sequence quality tranches [variants without a FILTER value of "PASS" that are above the 99.8% VQSR Tranche] were flagged;
(ii) Monomorphic variants were flagged;
(iii) Variants with high missing rate were flagged;

(iv) Variants with high read depth were flagged.

Variant-level QC criteria was applied to all variants with metrics (e.g., call rate, mean depth) computed across all samples per variant. Variant-level flags (“VFLAGS”) are enumerated and described below. Only variants with VLAGS=0 were retained for downstream analysis.

| VFLAGS | Meaning |
|---|---|
| 0 | Passed |
| 1 | *Variant failed preliminary QC:* GATK “FILTER” ≠ “PASS” or is in tranche ≥ 99.8% |
| 2 | *Variant failed preliminary QC:* All genotypes have DP<10 and/or GQ<20 |
| 3 | Monomorphic |
| 4 | Call Rate ≤ 80% |
| 5 | Average mean depth > 500 reads |

*Sample-level*

We removed 17 samples with genome-wide mean coverage < 20× and other sequencing quality issues (e.g., Mendelian inconsistencies). Sample-level QC metrics were computed across all variants for a specific subject. Mendelian inconsistency checking was performed at STEP 1 and STEP 2 QC. For STEP 1 QC, we inferred the Mendelian inconsistency (MI) rate of 0.12% among SNVs and 0.22% among indels. For STEP 2 QC, these rates shifted to 0.09% among SNVs and 0.20% among indels, demonstrating an overall improvement in data quality. MI checking was not performed on any pairs identified as cryptic relatives.

The 22 children of trios as well as 6 individuals with missing phenotypes (such as sampling information, etc.) were removed from the final dataset. A total of 2,679 individuals are considered in the rest of the analysis.

**Comparison of summary statistics to published whole genome sequences**

Our LASI-DAD filtered (after STEP 2 QC) variant call set includes 73.16 million (M) autosomal variants, including 67.11M SNVs and 6.04M indels. 53.5% of the SNVs are singletons (35.92M) and 34% (22.86M) are rare variants (allele count (AC) > 1 and allele frequency (AF) ≤ 1%). The Transition-to-Transversion ratio (Ti/Tv) for the SNVs is 2.13 for all variants, consistent with high quality and low error rate. Compared to other large scale sequencing datasets—the high coverage 1000G[19] and genome Aggregation Database (gnomAD, V3 (3.1.2))[20], LASI-DAD includes more than 26M novel autosomal SNVs (24 million novel SNVs and 2.2 million novel indels). The large majority of the new variants are rare, the 308,623 variants not present in

1000G have a frequency >=1% and 7,880 variants that are not present in gnomAD have a frequency ≥ 1% (Table S2.1).

**Table S2.1: Comparison of LASI-DAD autosomal variants with public datasets.** Number of variants corresponds to the number of SNVs + number of Indels.

| **Comparison set** | **#variants only seen in LASI-DAD** | **#variants overlapping comparison set** | **#variants unique to comparison set** | **#biallelic variants unique to comparison set but present in South Asians (in the comparison set)** |
|---|---|---|---|---|
| 1000G (phase 3) high coverage | 45,355,600 (singletons: 68%, MAF>1%: 431,861) | 27,801,012 | 95,773,594 | 16,132,139 |
| | SNVs: 41,387,839 | SNVs: 25,724,106 | SNVs: 86,136,390 | SNVs: 14,133,360 |
| gnomad v3 | 26,666,397 (singletons: 85%, MAF > 1% : 9,791) | 46,490,215 | 681,353,949 | 46,621,749 |
| | SNVs: 24,482,827 | SNVs: 42,629,118 | SNVs: 579,307,108 | SNVs: 23,831,475 |

*Per sample statistics*

We ran “bcftools stats -S” (*bcftools v1.6*) per sample to evaluate the quality of variant calls in LASI-DAD for both SNPs and indels. On average, the mean coverage per individual is 33.17x, with a minimum coverage of 25.6x. Summary statistics are presented in Figure S2.1-2 and Table S2.2.

**Table S2.2 Summary of the distributions of the number of variants per sample in the 2,679 LASI-DAD individuals.**

| **Mutation type** | **Median count** | **Mean count** | **25th percentile** | **75th percentile** |
|---|---|---|---|---|
| Substitutions | 3,211,043 | 3,206,584 | 3,199,559 | 3,220,356 |
| Singletons | 15,197 | 14,652.24 | 12,995.5 | 16,458 |
| Indels | 270,952 | 270,613.6 | 269,855.5 | 271,864 |
| Heterozygous to Non-reference homozygous ratio | 1.793 | 1.778 | 1.763 | 1.821 |

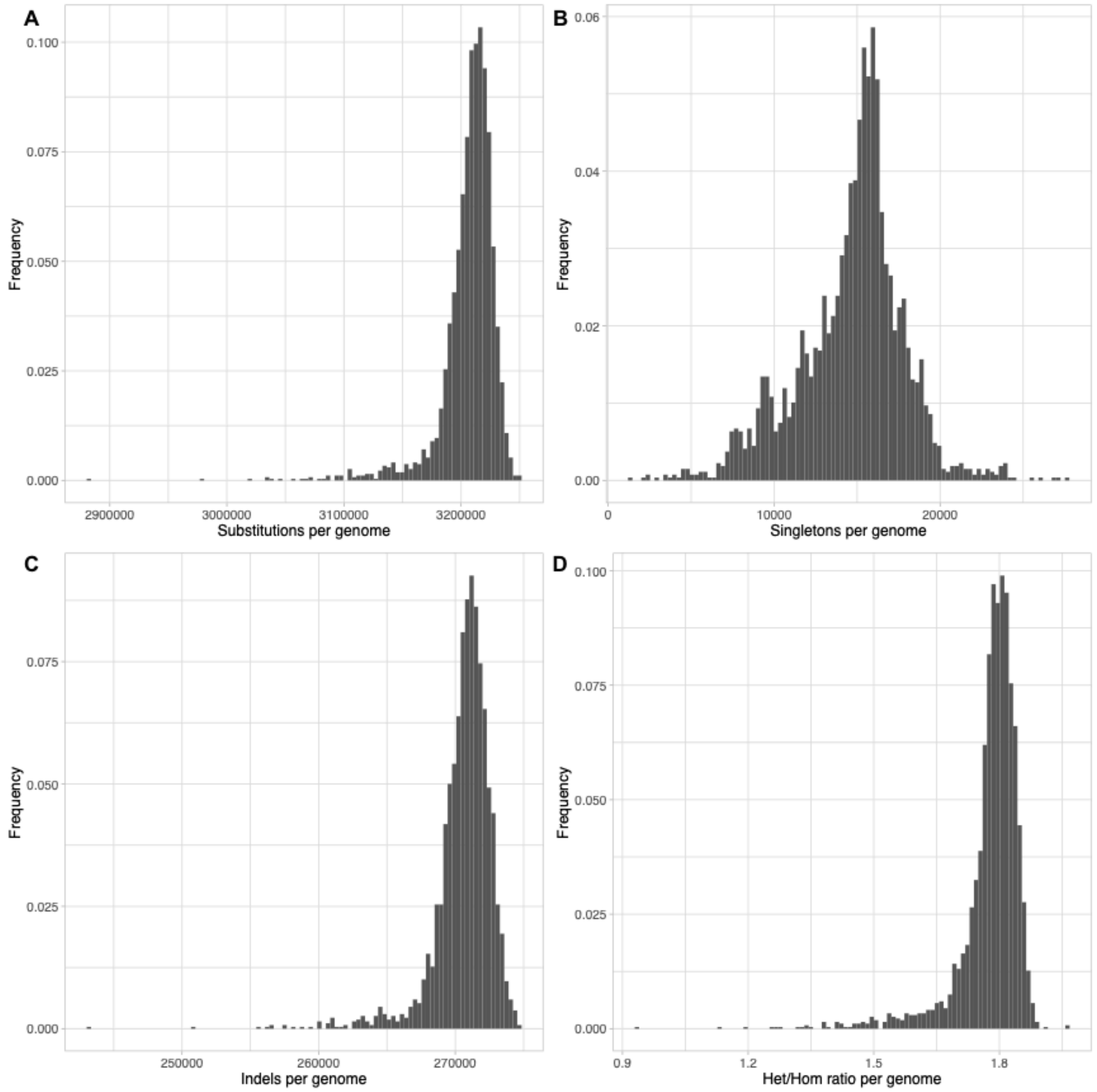

**Figure S2.1: Variant summary statistics per individual in LASI-DAD**. (A) Substitutions identified in LASI-DAD individuals. (B) Singletons identified in LASI-DAD individuals. (C) Indels identified in LASI-DAD individuals (D) The ratio of heterozygous to homozygous variants identified in LASI-DAD individuals.

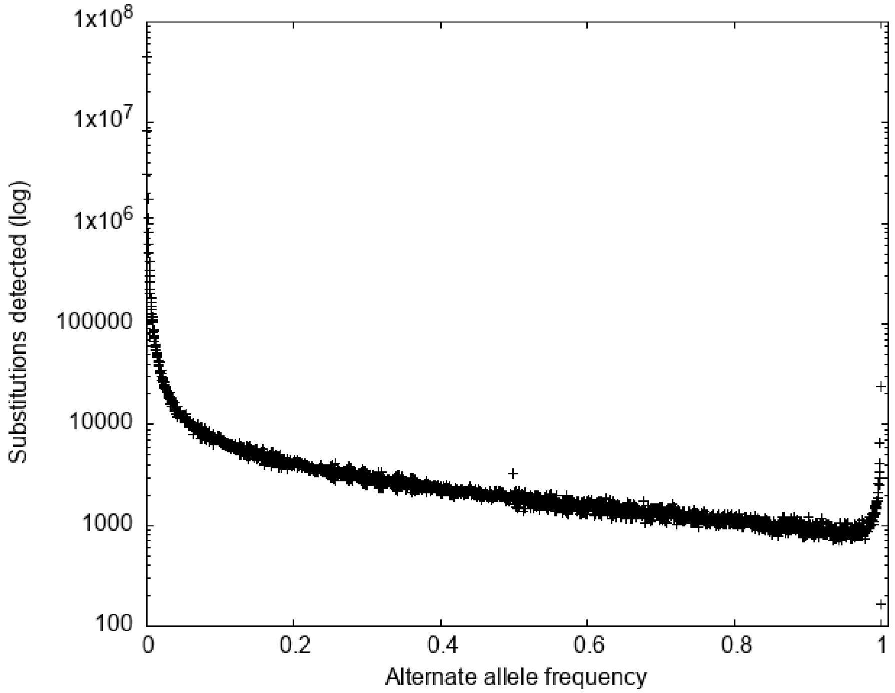

**Figure S2.2: Number of substitutions identified in function of the alternate allele count in LASI-DAD individuals**. Most of the alternate alleles are rare s (~90% of substitutions have a frequency <= 3%).

### Supplementary Note 3: Genome Phasing

For phasing, we used variants and genotypes that passed QC and included trios to examine the accuracy of computational phasing. This dataset comprises 2,762 individuals and 67,516,883 biallelic SNPs, including 22 parent-offspring trios.

To evaluate phasing accuracy, we applied two complementary approaches to 66 individuals from 22 trios (mother-father-child). First, we phased the 66 individuals using Beagle4[21] leveraging trio information for phasing. Second, we used computational phasing assuming all individuals were unrelated using SHAPEIT4[22], either without a reference panel or using two publicly available large datasets as reference panels: (i) The Human Genome Diversity Project (HGDP)[23] and (ii) 1000G[19]. We used the HapMap recombination map[24] for all three setups. We estimated the proportion of heterozygous variants that could be phased for each trio individual and the number of 'switch errors' that we define as the number of heterozygous sites where the genotype differs between the computational phasing and trio-based phasing for each individual.

The results for the comparison of trio-phased and computational phased heterozygous genotypes are shown in Table S3.1. We find the switch-error rates are similar across the 22 trios (Figure S3.1). Across reference panels, we find the switch error rate is lowest when using the HGDP reference panel; however, this setup only includes 19 million variants as we lose a large fraction of population-specific variants when combining the LASI-DAD samples with HGDP. The highest switch error rate (>1%) is obtained when we perform phasing without any reference haplotypes, though this setup allows phasing of almost all variants in the dataset (Table S3.1).

**Table S3.1**. **Estimated switch error rates for computational phasing with different reference panels.**

| Reference panel | Average switch error rate (percent per individual) | Percent of heterozygous variants which could be phased (percent per individual) | Number of variants in phased call set |
|---|---|---|---|
| HGDP | 0.614 | 95.6 | 19,204,945 |
| 1000G | 0.629 | 97.3 | 21,448,356 |
| No reference panel | 1.13 | 100 | 67,516,883 |

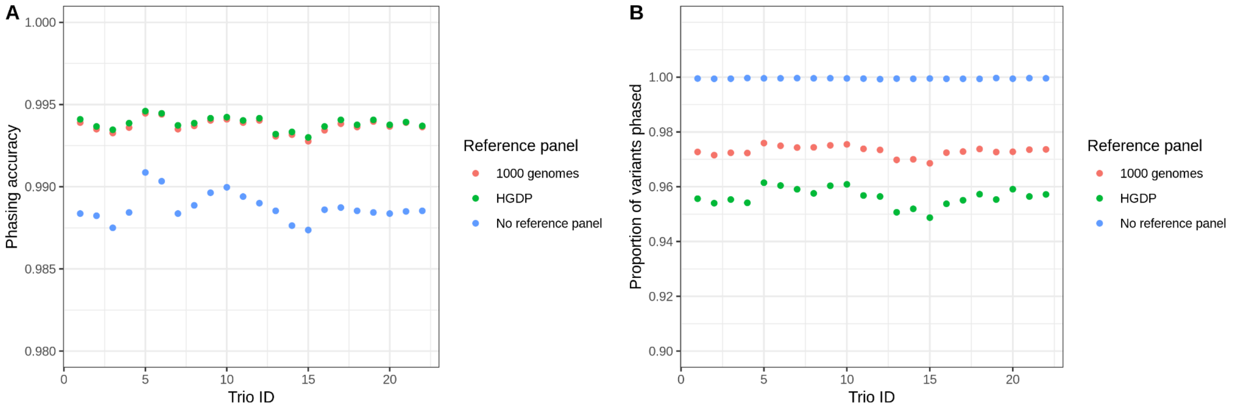

**Figure S3.1**. **Performance of genome phasing with different reference panels.** We applied trio-based (using Beagle4) and computational phasing (using Shapeit4) with or without reference panels for 22 trios and examined the switch error rates and proportion of variants phased. We show (A) The phasing accuracy (1 - switch error rate) for each trio, and (B) The average proportion of phased heterozygous variants for each trio.

To investigate the distribution of phasing errors by allele frequency, we grouped variants into 0.1% frequency bins (based on the frequency in 2,762 LASI-DAD individuals). For each bin, we computed the average switch error rate and the average proportion of phased variants. We inferred these summary statistics separately for each reference panel. We find that the phasing works reliably (>95% accuracy) for common variants (minor allele frequency >1%) under all three setups. For rare variants (<0.1% frequency), when we use reference data (HGDP or 1000G), we obtain >95% accuracy though most (~80%) of the rare variants are excluded. We obtain phase information for all variants when we do not use a reference panel, but the error rates for rare variants are substantially higher (~10-15%) (Figure S3.2). We also investigated the density of switch errors along the genome in 500 kb windows. We find that errors are primarily located adjacent to telomeres and centromeres (Figure S3.3).

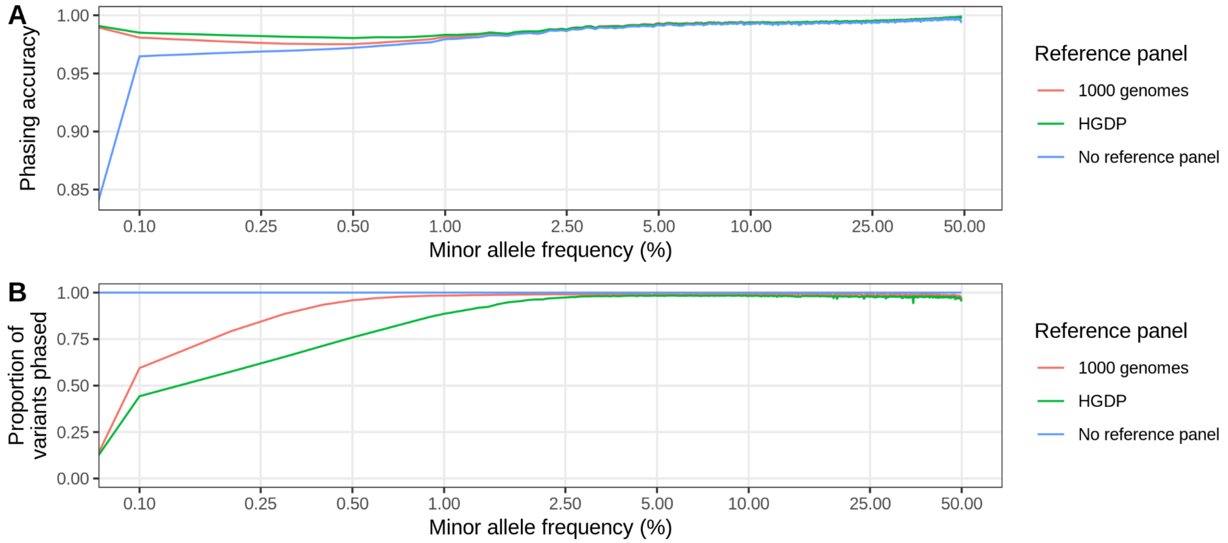

**Figure S3.2. Performance of genome phasing depending on the minor allele frequency.** (A) Phasing accuracy as a function of minor allele frequency (on a log scale) in the LASI-DAD dataset for all three reference panels. (B) Proportion of variants phased (amount of heterozygous sites that could be phased.

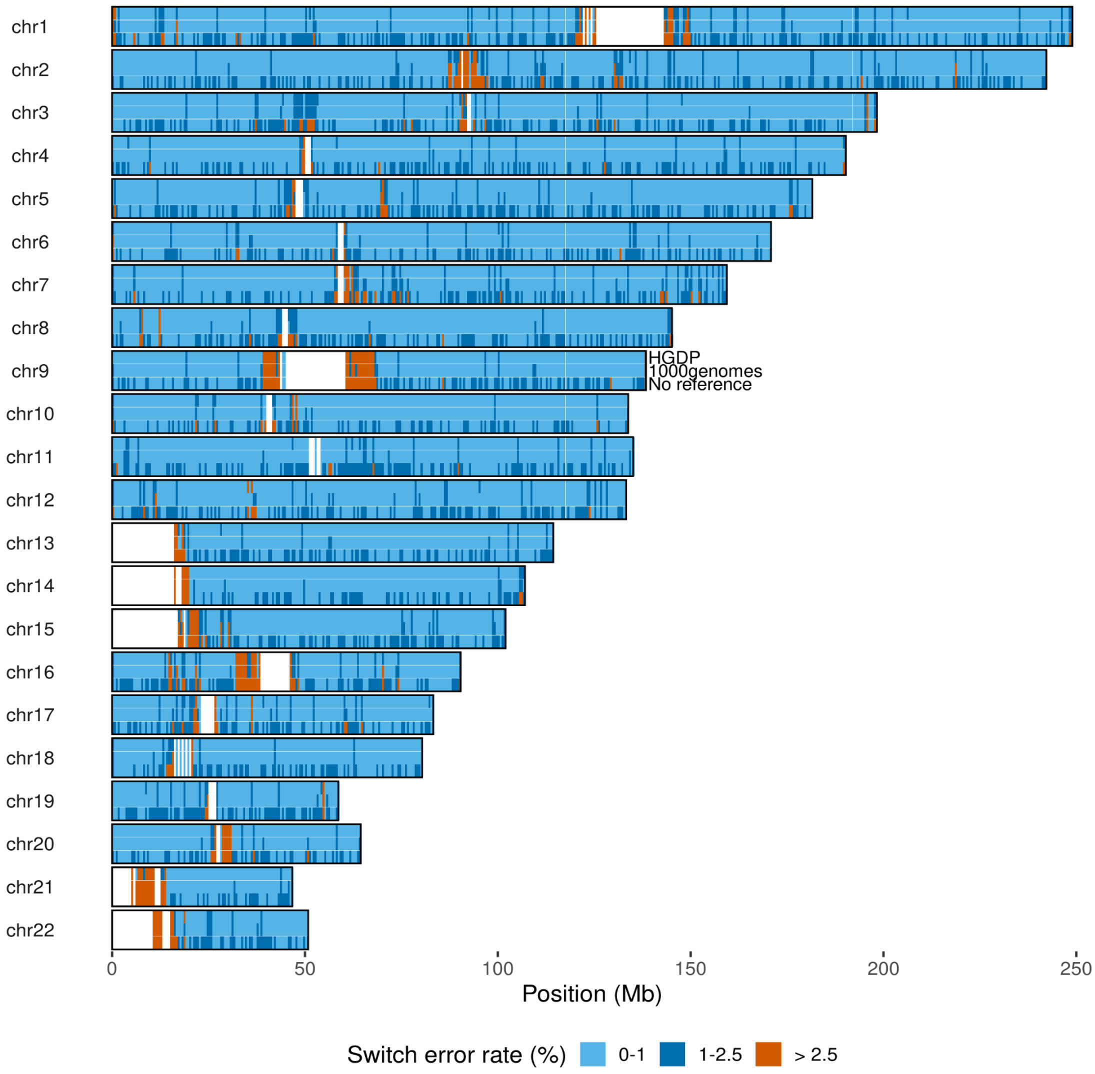

**Figure S3.3. Distribution of switch errors along the genome.** We measure switch errors in windows of 500 kb along the genome. Windows without any variants present are colored in white and mostly correspond to centromeres and telomeres. For each chromosome, we show the average switch error rate for three phasing setups: the first row includes the results with the HGDP reference panel, the second row includes the results with the 1000G reference panel, and the last row includes the phasing results without any reference panel.

### Supplementary Note 4: Population structure and admixture

To learn about the population history of India, we combined the LASI-DAD dataset with other published genomic datasets including present-day individuals from 1000G[19], GenomeAsia[25] and ancient DNA samples from the Allen Ancient DNA Resource (AADR)[26]. We performed PCA[27], ADMIXTURE[28] and *f*-statistics[29] to study the population relationships and ALDER[30] to infer the timing of admixture events.

**Table S4.1 Description of datasets used for the different analysis of this section, the number of individuals and the number of variants in the final datasets.**

| Dataset | #Individuals | #Variants | LD-pruned* - Y/N | Analysis used for |
|---|---|---|---|---|
| LASI-DAD + 1000G[19] | 5,332 | 1,654,184 (frequency >0.05) | Y | PCA, ADMIXTURE |
| LASI-DAD + AADR (v54)[26] + GenomeAsia[27] | 20,340 | 929,153 | N | *f*-statistics, *qpAdm*, ALDER |

Note: *For LD-pruning, we used PLINK with the option '--indep-pairwise 50 10 0.5' to remove, in a window of 50 SNPs, one variant of a pair of SNPs if the LD is greater than 0.5.

#### *Principal Component Analysis (PCA)*

To understand the population structure of Indians and their relationship to other worldwide populations, we performed Principal component analysis (PCA) using smartpca[27]. We used 2,620 unrelated individuals LASI-DAD and 2,712 individuals from 1000G including Europeans (EUR, 633 individuals from CEU, TSI, FIN, GBR and IBS), East Asians (EAS, 585 individuals from CHB, JPT, CHS, CDX and KHV), South Asians (SAS, 601 individuals from GIH, PJL, BEB, STU and ITU) and sub-Saharan Africans (AFR, 893 individuals from YRI, LWK, MAG, MSL, ESN, ASW and ACB) after LD-pruning and filtering the combined dataset (Table S4.1). Figure S4.1-4 show the results of PCA. In Figure S4.1, we observe that PC1 separates populations related to sub-Saharan African from other worldwide populations. PC2 shows a gradient of individuals of East Asian-related ancestry (in blue) on the one end and individuals of European-related ancestry ( in red) on the other end. Individuals of South Asian-related ancestry (Indians from LASI-DAD and SAS from 1000G) fall in between the European- and East Asian-related ancestry clusters (referred to as ''Indian cline''), which we have previously shown to reflect variable proportions of ancestry from two ancestral groups: the Ancestral North Indians (ANI) who harbor large proportions of ancestry related to West Eurasians, and the Ancestral South Indians (ASI) who are distantly related to West Eurasians[31,32]. Figures 1 and S4.2-4 show the results of the PCA with LASI-DAD, EUR and EAS described in the main text. We find the population structure in India is related to state of birth (Figure S4.2), linguistic affiliation (Figure S4.3) and caste affiliation (Figure S4.4).

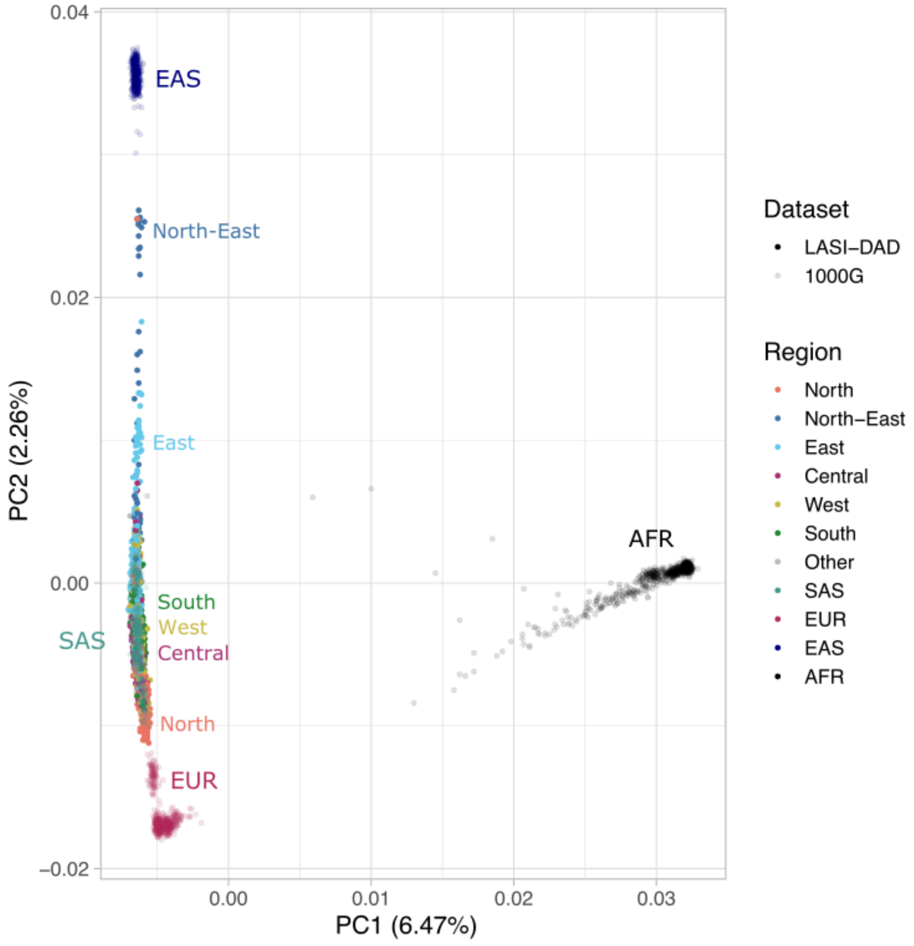

**Figure S4.1: Principal Component Analysis of LASI-DAD and 1000G.** We show the PCA of 2,620 Indians from LASI-DAD samples and individuals of African (AFR), East Asian (EAS), European (EUR) and South Asian (SAS) populations in 1000G. Colors represent the superpopulation for 1000G individuals and the birth region for LASI-DAD individuals.

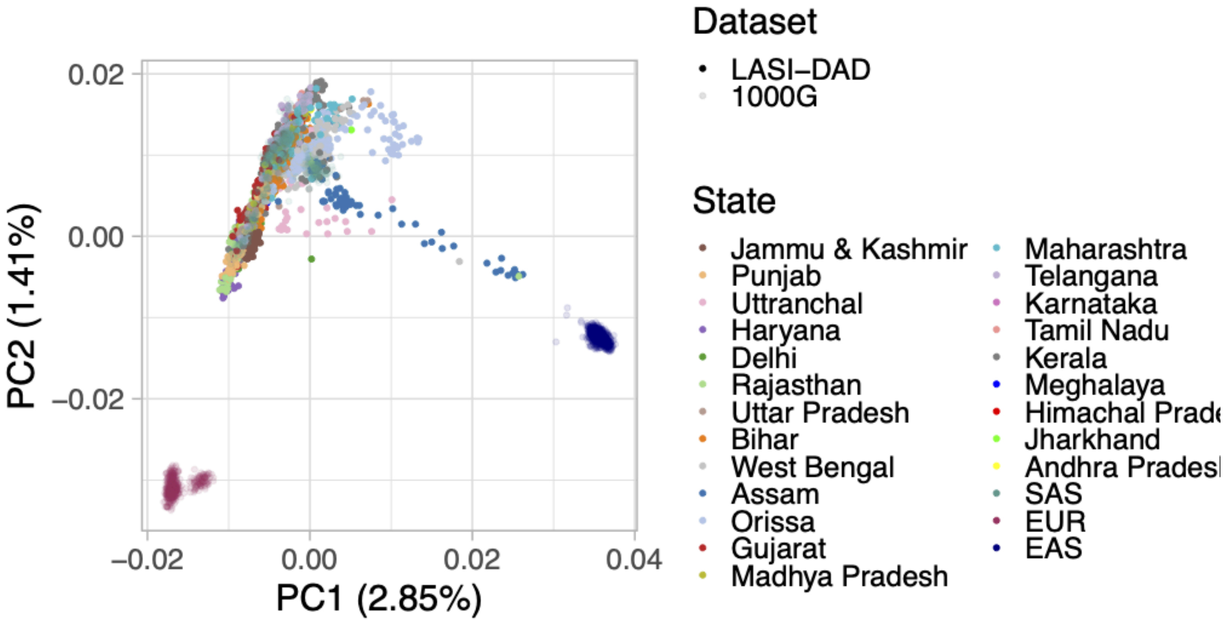

**Figure S4.2: Principal Component Analysis of LASI-DAD and 1000G.** We show the PCA of 2,620 Indians from LASI-DAD samples and individuals of East Asian (EAS), European (EUR) and South Asian (SAS) populations in 1000G. Colors represent the superpopulation for 1000G individuals and the state of birth in India for LASI-DAD individuals.

A

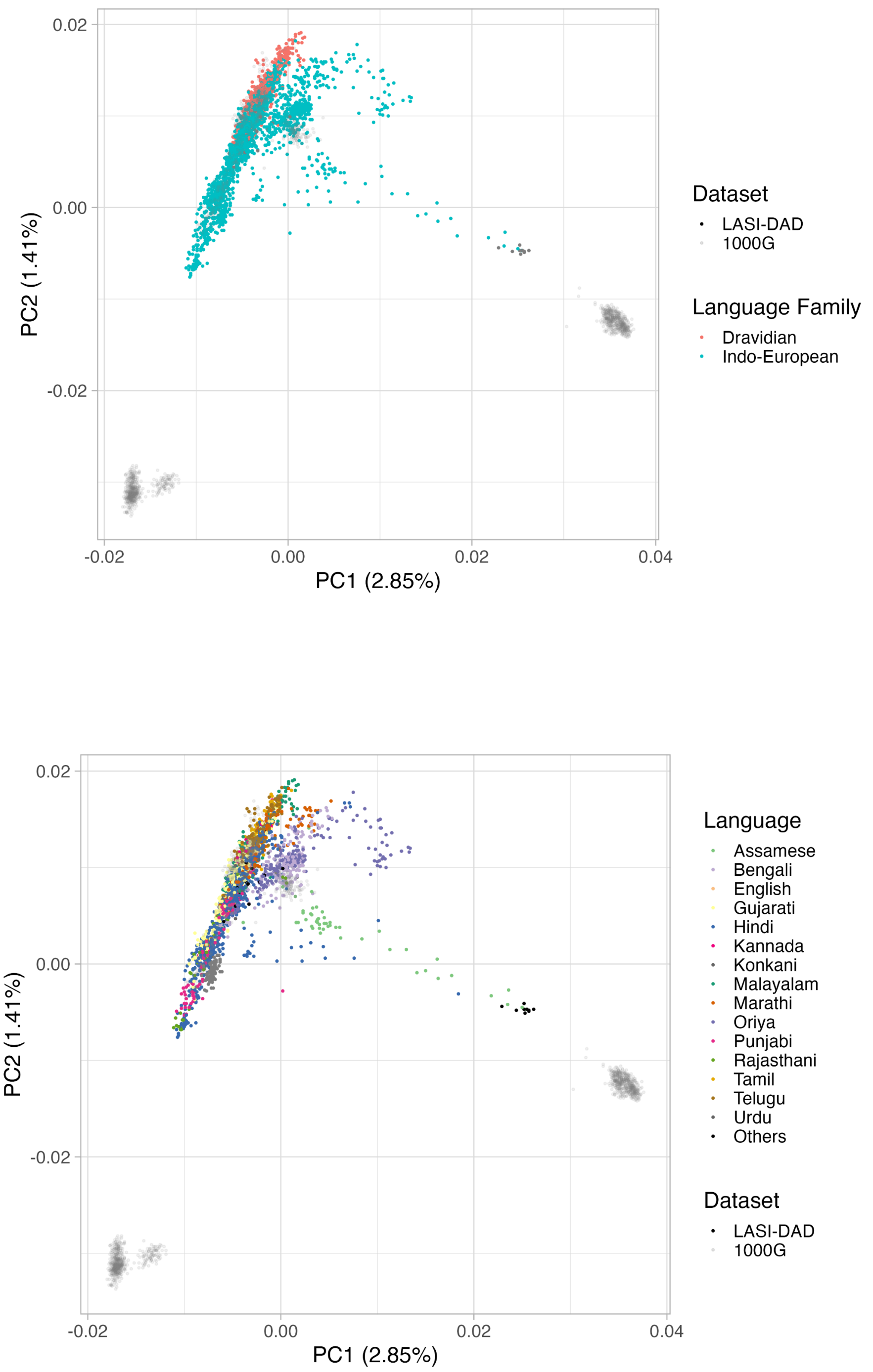

**Figure S4.3: Principal Component Analysis of LASI-DAD and 1000G.** We show the PCA of 2,620 Indians from LASI-DAD samples and individuals of East Asian (EAS), European (EUR) and South Asian (SAS) populations in 1000G. Colors represent the superpopulation for 1000G individuals. For LASI-DAD, we show (A) the language family of the mother tongue, and (B) the mother tongue for LASI-DAD individuals.

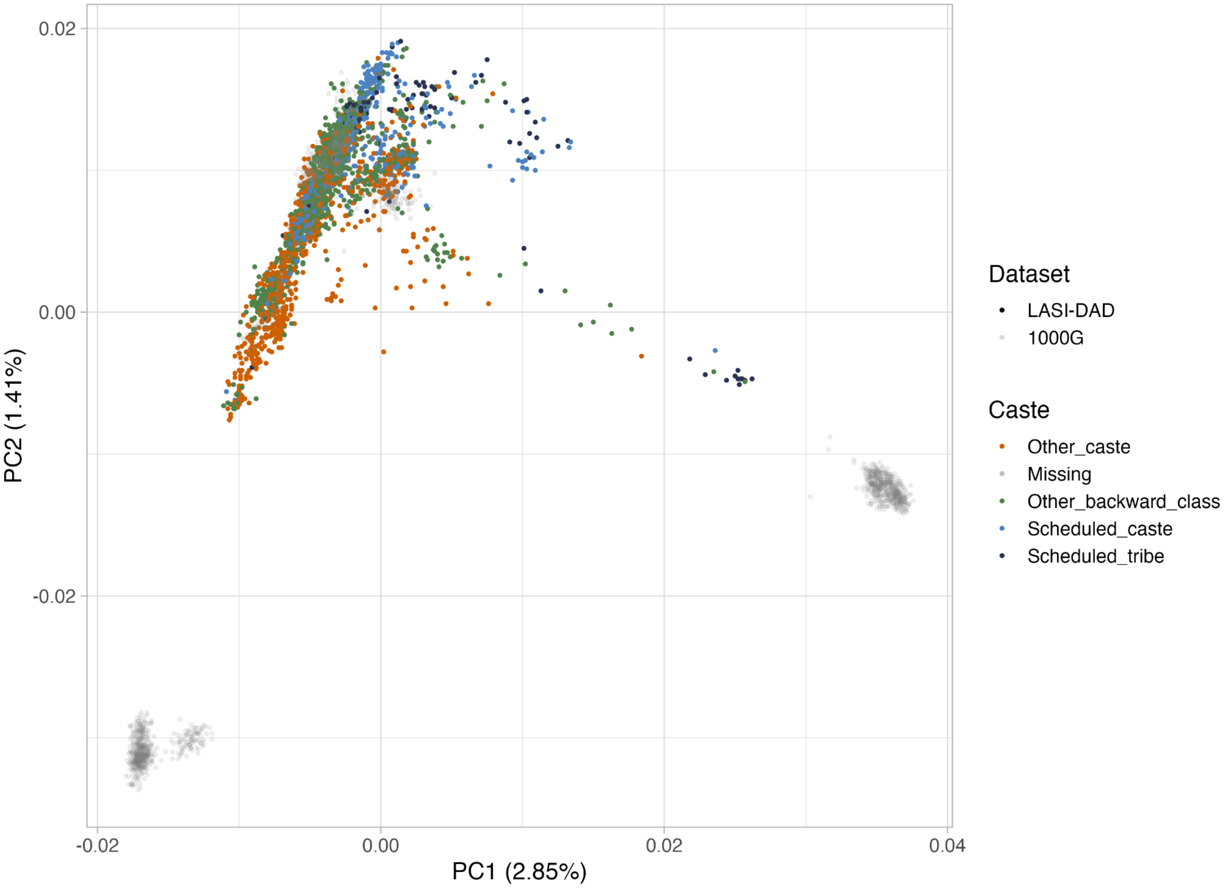

**Figure S4.4: Principal Component Analysis of LASI-DAD and 1000G.** We show the PCA of 2,620 Indians from LASI-DAD samples and individuals of East Asian (EAS), European (EUR) and South Asian (SAS) populations in 1000G. Colors represent the superpopulation for 1000G individuals and the caste affiliation for LASI-DAD individuals.

***ADMIXTURE***

We performed unsupervised clustering using ADMIXTURE[28] with the merged dataset of LASI-DAD and 1000G (including EAS, EUR, SAS and AFR), similar to the PCA analysis. We varied the number of clusters (*K*) between 2–6 and performed cross validation by running the method ten times (option: *--cv=10*). We obtained the lowest cross-validation error (*CV*) for *K*= 5 which increased at *K*=6 (we note, *CV* for *K*=4 and *K*=5 are very similar) (Figure S4.5). As seen in PCA, the first two clusters in ADMIXTURE (*K*=2) separate sub-Saharan African-related populations from non-Africans. At *K*=3, we observe clustering based on relatedness to three main continental ancestries including sub-Saharan African-, East Asian- and European-related groups. Indians from LASI-DAD, as well as SAS from 1000G, have variable relatedness to East Asian- and European-related ancestry components. For *K*=4 and *K*=5, we observe South Asian-related components replacing the East Asian-related (for *K*=4) and European-related (for *K*=5) ancestry components, highlighting that the gene flow in India is not directly related to EUR or EAS but instead to ancestral populations like ANI and ASI respectively. At *K*=6, East Asian-related ancestry is separated into two different components which can be found in some South Asian individuals.

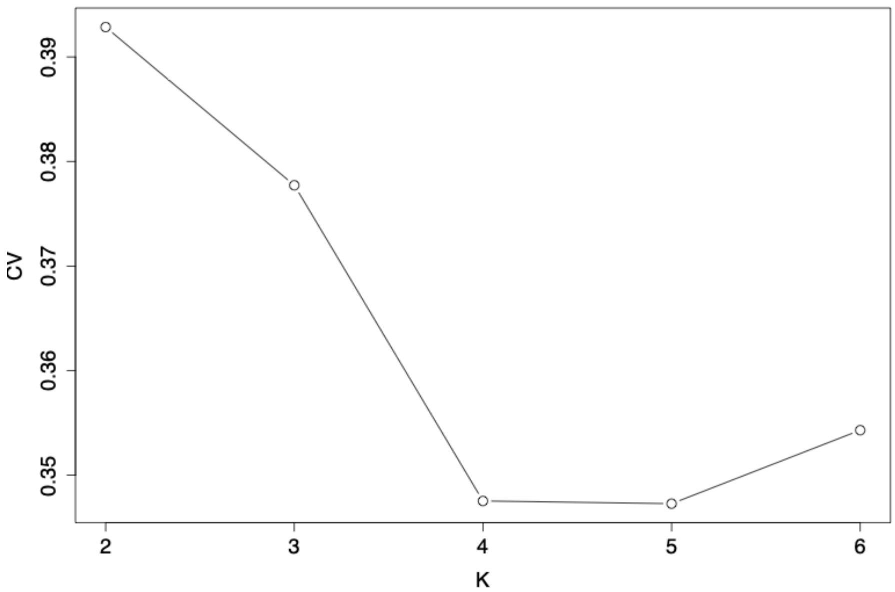

**Figure S4.5: Cross-validation errors from ADMIXTURE analysis.** We performed ADMIXTURE analysis using the merged dataset of LASI-DAD and 1000G (with AFR, EAS, EUR and SAS populations) (Table S4.1). We show the cross-validation errors for the number of clusters ( *K*) between 2-6. The lowest value is obtained for *K*= 5, *CV*=0.34727 (for *K*=4, *CV*= 0.34753).

**K=2**

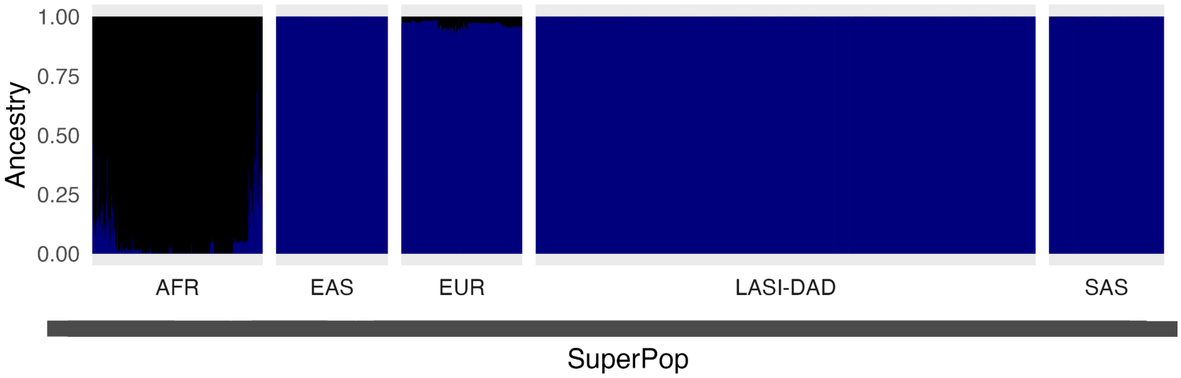

**K=3**

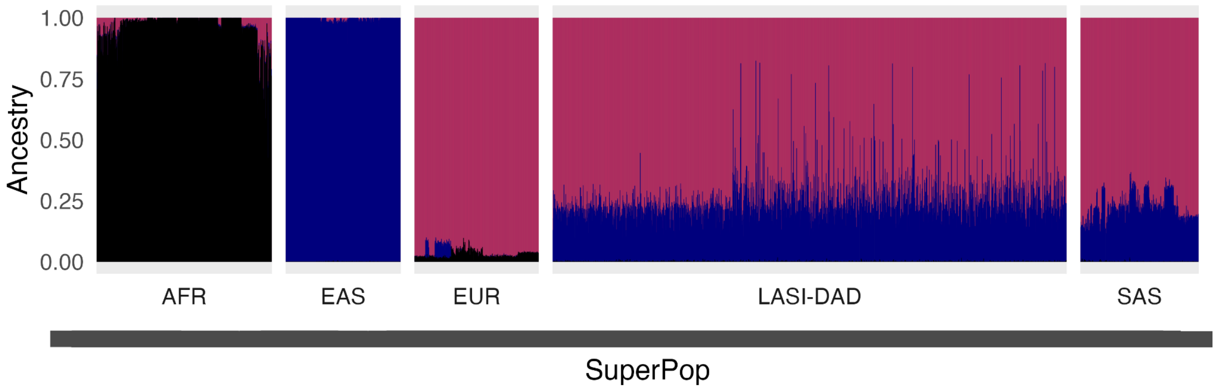

1.00
0.75
0.50
0.25
0.00
Ancestry
AFR
EAS
EUR
LASI-DAD
SAS
SuperPop

**K=4**

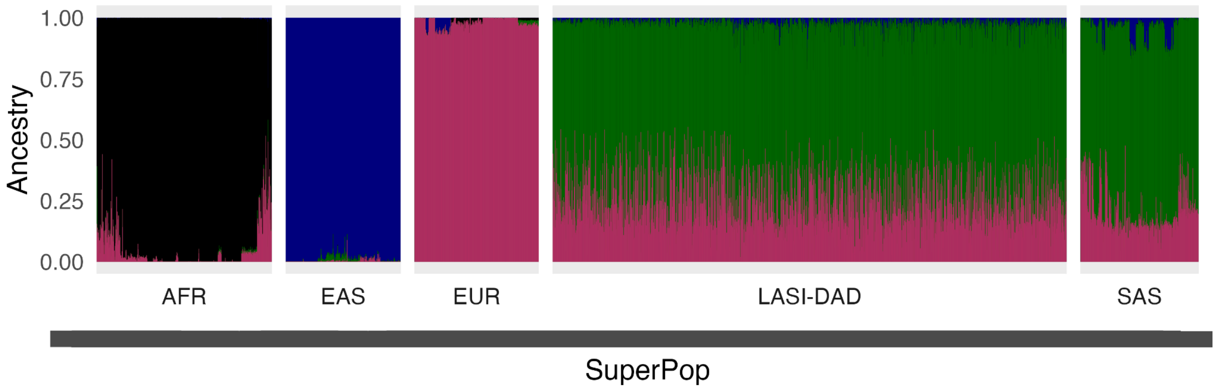

1.00
0.75
0.50
0.25
0.00
Ancestry
AFR
EAS
EUR
LASI-DAD
SAS
SuperPop

**K=5**

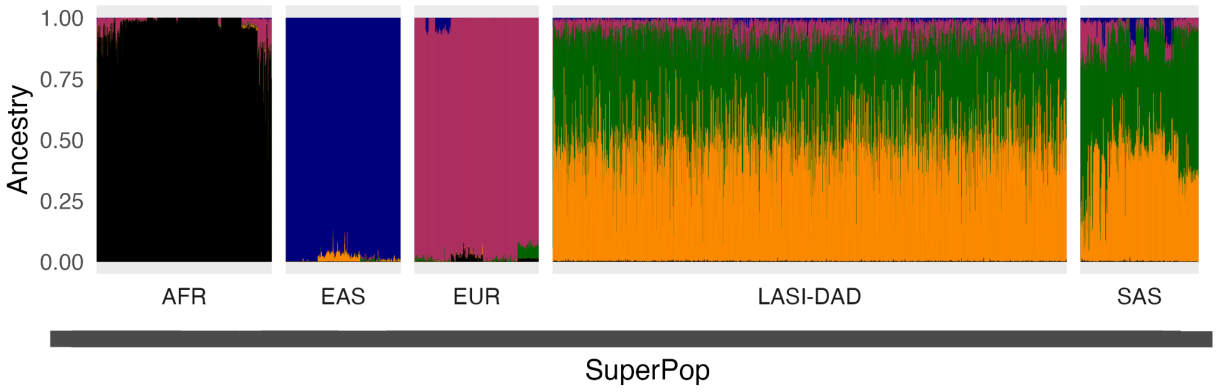

1.00
0.75
0.50
0.25
0.00
Ancestry
AFR
EAS
EUR
LASI-DAD
SAS
SuperPop

**K=6**

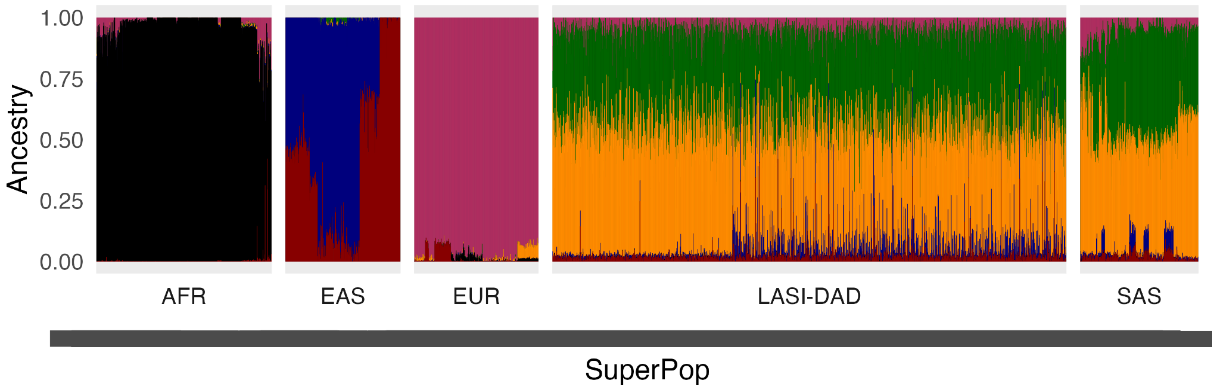

**Figure S4.6. ADMIXTURE analysis of LASI-DAD and 1000G populations.** We performed ADMIXTURE analysis using the merged dataset of LASI-DAD and 1000G (with AFR, EAS, EUR and SAS populations) (Table S4.1). We vary the number of clusters (*K*) between 2-6. Each vertical line represents one individual, the different colors are the ancestry proportions from the different clusters inferred with ADMIXTURE.

### Ancestry composition of India

We applied *qpAdm*[29,33] that compares allele frequency correlations between the population of interest and a set of reference and outgroup populations to formally test the model of population relationships and then infer the ancestry proportions for the best-fitted model. For this analysis, we used the merged dataset including LASI-DAD, AADR (v54)[26] and the Genome Asia[25] (Table S4.1).

#### *Model of ancestry for individuals on the Indian cline*

Following Narasimham et al. 2019, we modeled the individuals on the Indian cline as a mixture of three ancestral populations related to *Ancestral Ancient South Indians* (AASI), ancient Eurasian Steppe pastoralists and ancient Iranian farmers that are represented by the following source populations: the *Andamanese hunter-gatherers* (*AHG*) that are an indigenous group from the Andaman Islands that form a clade with the AASI group, the *Indus Periphery West* that has the highest Iranian farmer-related ancestry among the *Indus Periphery Cline* individuals that has been shown to be a good proxy for the Iranian farmer-related ancestry in India, and the *Central_Steppe_MLBA* that is a group of 34 individuals from the Middle to Late Bronze Age of Steppe pastoralists[34]. This group includes individuals from *Georgievsky Bugor* (n=1), *Kazakh Mys* (n=4), *Kyzl bulak* (n=1), *Oy-Dzhaylau* (n=6), *Shoendykol* (n=3), *Taldysay* (n=1) from Kazakhstan, from Kashkarchi (n=2) in Uzbekistan and form Krasnoyarsk (n=16) in Western Siberia. It has been suggested that the *Central_Steppe_MLBA* cluster was the primary conduit for spreading Yamnaya Steppe pastoralist–derived ancestry to South Asia in the first half of the second millennium BCE[34]. We used the following outgroups: *Ethiopia_4500BP.SG*, *WEHG*, *EEHG*, *IranGanjDareh_N*, *Anatolia_N*, *WSHG*, *ESHG* and *Dai.DG*.

Using *qpAdm*, we find the three-way model fits well for 92.7% of the individuals in LASI-DAD (*p*-value > 0.01). We find qualitatively similar results when the *Indus Periphery Cline* ($n$ = 11) is used as the source for the Iranian farmer-related ancestry (with *AHG-related* and *Central_Steppe_MLBA* as the other two sources). Specifically, the inferred ancestry coefficients are highly correlated for all three ancestry components (Pearson's correlation coefficients: $r^2_{\_Iran\text{-}farmer\text{-}related} > 0.99$, $r^2_{\_AHG\text{-}related} > 0.99$ and $r^2_{\_Steppe\text{-}Pastoralist\text{-}related} > 0.995$), though shifted as *Indus Periphery West* has the lowest proportion of *AHG*-related ancestry among *Indus Periphery Cline* individuals[34] (Fig S4.7). This model with *Indus Periphery Cline*, however, fits only for 81.9% of the individuals highlighting the heterogeneity in ancestry of the *Indus Periphery Cline*.

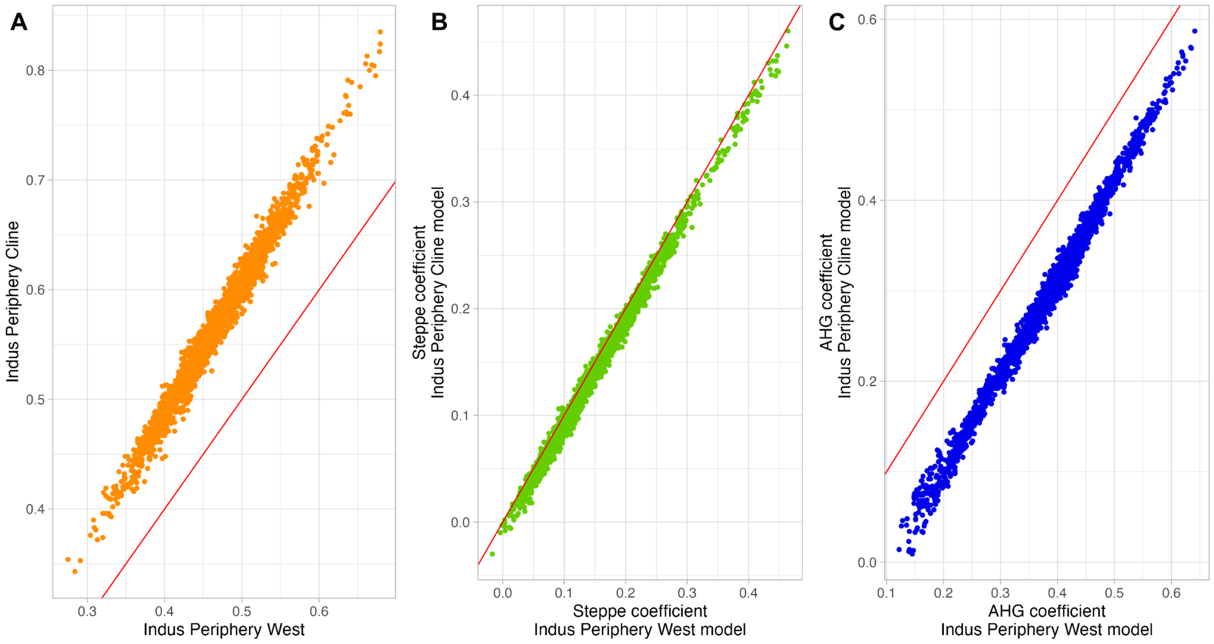

**Figure S4.7. Correlations between coefficients inferred with *Indus Periphery West* and *Indus Periphery Cline* as the Iran farmer-related source for the three-way model applied on Indian cline individuals.** For (A) the Iran farmer-related coefficient, (B) the Steppe pastoralist-related coefficient and (C) the *AHG*-related coefficient.

#### *Source of Iranian farmer-related ancestry in India*

To investigate the Iranian farmer-related ancestry in India, we first searched for Indian individuals that lack Steppe pastoralist-related ancestry (referred to as *ASI* henceforth). To this end, we examined if any individual on the Indian cline is better modeled as a two-way mixture with Iranian farmer-related and *AHG*-related ancestries, compared to the three way model including Steppe pastoralist-related ancestry. We find 22 individuals in LASI-DAD are a good fit for the two-model without Steppe pastoralist-related ancestry (with higher *p*-value for the two-way model compared to the three-way model).

Next, we tested 15 ancient Iranian ancestry groups from the Neolithic to Iron Age as the source for Iranian farmer-related ancestry for the 22 ASI samples, with *AHG-related* ancestry as the other source (Table S4.2). A previous study has shown that the *Indus Periphery Cline* individuals can also be modeled as a two-way mixture with ancestry from Iranian farmer-related and *AHG*-related groups, with the most likely proximal model including *AHG*-related + *Parkhai_Anau_EN* + *Sarazm_EN*[34]. For comparison, we therefore added the *Indus Periphery West* individual and also considered the three way model including two Iranian farmer-related groups (*Parkhai_Anau_EN* + *Sarazm_EN).* We used the following outgroups (*right* populations) for the analysis: *Ethiopia_4500BP.SG, WEHG, EEHG, Anatolia_N, ESHG ,Dai.DG, Iran_GanjDareh_N* and *Russia_Samara_EBA_Yamnaya*. For models involving *Iran_GanjDareh_N* as one of the source populations, we used *Iran_HajjiFiruz_N* as an outgroup population. We report *p*-values for all models for the 23 individuals (22 LASI-DAD + *Indus Periphery West*) in Table S4.2.

**Table S4.2. Testing different sources for the Iranian farmer-related populations in two-way models with Iranian farmer-related + *AHG*-related ancestries in *qpAdm*.** For Indus Periphery West, green are fits according to the acceptance criterion (*p*-value > 0.01) and red shows models that fail, for ASI individuals, we represent the number and percentage of individuals that fit, in green are models with 100% of the individuals that fit and red otherwise.

| Iranian-related populations | *22 ASI count (%)* | *Indus Periphery West qpAdm p-value* |
|---|---|---|
| Aigyrzhal_BA | 0 (0%) | 0.000 |
| BMAC | 12 (54%) | 0.000 |
| Geoksyur_EN | 20 (91%) | 0.012 |
| Hajji_Firuz_BA | 0 (0%) | 0.000 |
| Iran_C_SehGabi | 0 (0%) | 0.000 |
| Iran_C_TepeHissar | 2 (9%) | 0.000 |
| Iran_DinkhaTepe_BA_IA_1 | 0 (0%) | 0.000 |
| Iran_DinkhaTepe_BA_IA_2 | 0 (0%) | 0.000 |
| Iran_GanjDareh_N | 3 (14%) | 0.000 |
| Iran_ShahrISokhta_BA1 | 20 (91%) | 0.000 |
| Namazga_CA | 22 (100%) | 0.231 |
| Parkhai_Anau_EN | 16 (73%) | 0.004 |
| SPGT | 0 (0%) | 0.000 |
| Sarazm_EN | 22 (100%) | 0.004 |
| Turkmenistan_Gonur_BA_1 | 15 (68%) | 0.049 |
| Parkhai_Anau_EN + Sarazm_EN | 22 (100%) | 0.097 |
| Iran_GanjDareh_N + WSHG | 16 (73%) | 0.267 |

We find that there are three models that provide a good fit for the 22 ASI individuals, including *Sarazm_EN, Namazga_CA* and (*Sarazm_EN* + *Parkhai_Anau_EN*) (Table S4.2). Interestingly, we find that for *Indus Periphery West* only the latter two models provide a good fit; *Sarazm_EN* alone does not provide a good fit (Table S4.2). *Parkhai_Anau_EN* and *Sarazm_EN* have similar ancestry profiles including ancestry from groups related to Western Siberian hunter-gatherers (*WSHG*), Anatolian farmers (*Anatolia_N*) and Iranian farmers (*Ganj_Dareh_N*), thus these results seem puzzling.

Next, we explored the source of the Iranian farmer-related groups for the individuals on the Indian cline (*n*=2,126, note we include the 22 ASI individuals in this set). Using *qpAdm*, we examined the fit of the three-way model with *AHG*-related and *Steppe pastoralist*-related groups, varying the source of the *Iranian farmer*-related ancestry between *Sarazm_EN, Namazga_CA* and (*Sarazm_EN* + *Parkhai_Anau_EN*). We also explored the two-way model (*Iranian farmer*-related + *AHG*-related sources) without *Steppe pastoralist*-related ancestry. We used the following outgroups: *Ethiopia_4500BP.SG, WEHG, EEHG, Anatolia_N, ESHG, Dai.DG, Iran_GanjDareh_N* and *Russia_Samara_EBA_Yamnaya*. For each model, we report the number of individuals for which the model fits (*p*-value > 0.01 and there are no negative mixture coefficients) in Table S4.3.

**Table S4.3 Testing different sources for the Iran farmer-related ancestry for individuals on the Indian cline (n=2,126).** We used *qpAdm* with two-way: *AHG*-related + Iranian farmer-related ancestry or three-way: *AHG*-related + Iranian farmer-related ancestry + Steppe pastoralist-related (Central_Steppe_MLBA) ancestries. Source for Iranian farmer-related ancestry shown below. We use the acceptance criterion of *p*-value > 0.01.

| **Iranian farmer-related group** | **Model with *p*-value > 0.01 (count / %)** | **Model with p-value > 0.01 & mixture coefficients > 0 (count / %)** | **Comment** |
|---|---|---|---|
| *Sarazm_EN* (either two-way or three-way model | 2029 / 95.4% | 2029 / 95.4% | |
| *Sarazm_EN* (three-way model) | 2028 / 95.4% | 1942 / 91.3% | 86 individuals have negative coefficients for *Steppe pastoralist*-related ancestry |
| *Sarazm_EN* (two-way model) | 825 / 38.8% | 825 / 38.8% | |
| *Namazga_CA* (either two-way or three-way model) | 1773 / 83.4% | 1773 / 83.4% | |
| *Namazga_CA* (three-way model) | 1773 / 83.4% | 1760 / 82.8% | 13 individuals have negative coefficients for *Steppe pastoralist*-related ancestry |
| *Namazga_CA* | 499 / 23.5% | 499 / 23.5% | |

| | | | |
|---|---|---|---|
| (two-way model) | | | |
| *Sarazm_EN* + *Parkhai_Anau_EN* (either two-way or three-way model) | 1999 / 94.0% | 1497 / 70.4% | In the 502 individuals with negative coefficients:<br>5 for *Sazarm_EN*, 497 for *Parkhai_Anau_EN* , 3 for *Steppe pastoralist*-related ancestry |
| *Sarazm_EN* + *Parkhai_Anau_EN* (three-way model) | 1999 / 94.0% | 1318 / 62.0% | In the 681 individuals with negative coefficients:<br>5 for *Sazarm_EN*, 658 for *Parkhai_Anau_EN*, 30 for *Steppe pastoralist*-related ancestry |
| *Sarazm_EN* + *Parkhai_Anau_EN* (two-way model) | 554 / 26.1% | 301 / 14.2% | 253 individuals have negative coefficients for *Parkhai_Anau_EN* |

Comparing among the sources of Iranian farmer-related ancestry, we find *Sarazm_EN* provides a good fit for the largest fraction of individuals on the Indian cline (>95%) (Table S4.3). In contrast, *Namaza_CA* fails for >15% of the models. Interestingly, (*Sarazm_EN* + *Parkhai_Anau_EN)* provides a good fit (*p*-value > 0.01) for similar fraction of individuals (~94%), but the mixture coefficient of *Parkhai_Anau_EN*-related ancestry are negative for 32.9% of the individuals on the Indian cline. This suggests *Sarazm_EN* is the best proxy for the Iranian farmer-related ancestry for India.

***Model of ancestry for individuals that fall outside the Indian cline***

In PCA, there are 494 individuals that fall outside the Indian cline including 332 individuals from the East, 68 individuals from the North-East, 31 from the West, 30 from the Central, 7 from the North, 2 from the South and 24 from Other*.* To understand their ancestry composition, we performed *qpAdm* analysis. First, we applied the three-way model that provides a good fit for the individuals on the Indian cline including Iranian farmer-related (*Sarazm_EN*), Steppe Pastoralist-related (*Central_Steppe_MLBA*) and *AHG*-related ancestries (referred to as model *a* in Table S4.4*)*. We find this model provides a good fit for ~64% (*n* = 314) individuals and they possibly appear off the cline due to drift (we define a good fit as models with *p*-value > 0.01 with non-negative ancestry coefficients).

The remaining 180 individuals fall in two main clusters in the PCA: one towards the *ASI*-end of the cline and other intermediate to the Indian cline exhibiting clear relatedness to East Asian-related groups in PCA (Fig 1). We separately modeled the individuals in the two clusters (Figure S4.8). As the individuals in the *ASI*-end of Indian cline are mostly from Odisha (38 out of 53) and speak Indo-European and Austroasiatic languages, we performed qpAdm with model (*b*) including Iranian farmer-related (*Sarazm_EN*), *AHG*-related and Austro-asiatic-related (using *Nicobarese*) ancestries. For the second cluster where individuals show East Asian-related ancestry, we tried model (*c*) that includes Iranian farmer-related (*Sarazm_EN*), *AHG*-related and East Asian-related (using Han Chinese from Beijing China (*CHB)*) ancestries. We also tested four-way models with addition of *Central_Steppe_MLBA* if models (*b-c*) failed. We used the following outgroups: 'Ethiopia 4500BP.SG', 'WEHG', 'EEHG', 'Iran GanjDareh N', 'Anatolia N', 'ESHG', 'Dai.DG', 'Russia Samara EBA Yamnaya' and 'Vietnam_BA'. We report the details of the fits in Table S4.4.

Among the 180 individuals, we obtain good fits for 127 individuals, including 51/53 individuals with Austro-asiatic-related and 87/127 individuals with East Asian-related sources, with or without *Central_Steppe_MLBA* (Table S4.4). Notably, there are 91 individuals that can be modeled without Steppe pastoralist-related ancestry, including ~96% of the Austro-asiatic-related individuals (using model *b*). This suggests Iranian farmer-related ancestry likely did not come through Steppe pastoralist-related groups in the late Bronze Age.

**Table S4.4 Best fitted model for the 494 off cline individuals.**

| Best model | *Model a* | *Model b* % (count / ASI-end total) | *Model c* % (count / EAS-related total) | *Model c* + *Steppe pastoralist*-related % (count / EAS-related total) | No fit* % (count / total) |
|---|---|---|---|---|---|
| Number of Individuals | 314 | 96% (51/53) | 31% (40/127) | 37% (47/127) | 9% (42/494) |

*We define **Model *a***: *Sarazm_EN* + *AHG*-related + Steppe pastoralist-related ancestries (same as Indian cline), **Model b**: *Sarazm_EN* + *AHG*-related + Austro-asiatic-related (using *Nicobarese* as proxy), **Model c**: *Sarazm_EN* + *AHG*-related + East Asian-related (using *CHB* as proxy), and **Model c + Steppe pastoralist-related**: *Sarazm_EN* + *AHG*-related + Steppe pastoralist-related + East Asian-related ancestries. No fit describes the set of individuals for whom none of the above four models provide a good fit (*p*-value > 0.01 with non-negative ancestry coefficients in *qpAdm*). This includes 40 individuals in the East Asian-related cluster and two in the Austro-asiatic related cluster.

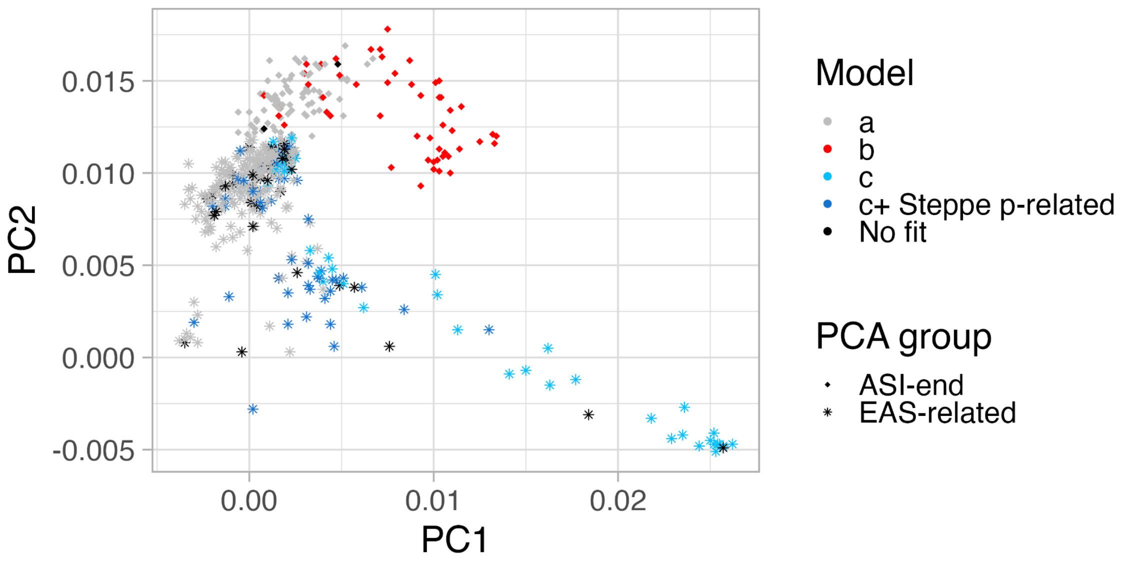

**Figure S4.8 Indian relatedness to Eurasian populations colored per inferred Off cline model.** qpAdm results for the 494 individuals that fall outside the Indian cline in Fig 1. We define **Model *a***: *Sarazm_EN* + *AHG*-related + Steppe pastoralist-related ancestries (same as Indian cline), **Model b**: *Sarazm_EN* + *AHG*-related + Austro-asiatic-related (using *Nicobarese* as proxy), **Model c**: *Sarazm_EN* + *AHG*-related + East Asian-related (using *CHB* as proxy) and **Model c + Steppe pastoralist-related**: Sarazm_EN + AHG-related + Steppe pastoralist-related ancestries + East Asian-related ancestries. Colors represent the best model inferred, first we test model a (gray), then either model b (red) or model c (light blue) / c+ Steppe pastoralist-related (dark blue), in function of the PCA group of the individuals (ASI-end in diamond and EAS-related in star). We define a good fit as models with p-value > 0.01 with non-negative ancestry coefficient.

*Population structure within the Sarazm_EN individuals*

The Sarazm_EN group consists of two samples from an agricultural village site located along the lower Zerafshan River valley in Tajikistan. The samples were dated to 3700–3300 BCE. Interestingly, one of the two *Sarazm_EN* individuals was found with shell bangles that are identical to ones found at sites in Pakistan and India such as Shahi-Tump, Makran and Surkotada, Gujarat (*J. Mark Kenoyer*, personal communication). To study if there was gene flow from *AHG*-related ancestry into *Sarazm_EN* individuals, we applied *qpAdm*. Following [34], we used the model with *Western Siberia hunter-gatherers* (*WSHG*)*, Anatolia_N* and *Ganj_Dareh_N* that was proposed earlier for *Sarazm_EN* (*n* = 2), but explored if addition of *AHG*-related ancestry provides a good fit for either of the two *Sarazm_EN* individuals (*p-value* > 0.01 with non-negative ancestry coefficients in *qpAdm*). We used the following outgroups: 'Ethiopia_4500BP.SG', 'WEHG', 'EEHG', 'ESHG', 'Dai.DG', 'Russia_Ust_Ishim_HG.DG', 'Iran_Mesolithic_BeltCave' and 'Israel_Natufian'.

Interestingly, we find significant variation in the ancestry of the two individuals in this group. One of the individuals,referred to *Sarazm_EN_1* (*I4290*) described above that was discovered with shell bangles showing affiliation with South Asia, has significant amount *AHG*-related ancestry, while a model without *AHG*-related ancestry provides the best fit for *Sarazm_EN_2* (*I4210*) (Table S4.5).

**Table S4.5 Ancestry profile of the two Sarazm_EN individuals.**

| | *p-value* | *Iranian farmer*-related | *WSHG* | *Anatolia_N* | *AHG*-related |
|---|---|---|---|---|---|
| Sarazm_EN_1 (I4290) | 0.300 | 0.639 | 0.132 | 0.070 | 0.159 |
| Sarazm_EN_1 (I4290) | 0.144 | 0.838 | 0.198 | <0 | Not inferred (3-way model) |
| Sarazm_EN_2 (I4910) | 0.0123 | 0.605 | 0.312 | 0.134 | <0 |
| Sarazm_EN_2 (I4910) | 0.023 | 0.543 | 0.278 | 0.179 | Not inferred (3-way model) |

To assess if our findings that *Sarazm_EN* is the most likely source for Iranian farmer-related ancestry in India is not merely due to the *AHG*-related gene flow in one of the individuals, we reran our *qpAdm* analysis with *Sarazm_EN_2* (the individual without *AHG*-related ancestry, Table S4.5). First, we modeled the 22 *ASI* individuals as a mixture of *Sarazm_EN_2* and *AHG*-related ancestry and show that this model fits the data. While *Sarazm_EN* alone does not provide a good fit to *Indus Periphery West* (requiring addition of *Parkhai_Anau_EN* earlier), we find use of *Sarazm_EN_2* provides fits the data (Table S4.6). This suggests that the earlier model potentially failed due to lack of enough Iranian farmer-related ancestry which was supplemented through*Parkai_Anau_EN* that has a very similar ancestry composition to *Sarazm_EN* (without *AHG*-related ancestry).

**Table S4.6 Comparison of *Sarazm_EN* and *Sarazm_EN_2* as sources for the Iranian farmer-related populations in two-way models with Iranian farmer-related + AHG-related ancestries in qpAdm.** Similarly to S4.2, for Indus Periphery West, green are fits according to the acceptance criterion (p-value > 0.01) and red shows models that fail, for ASI individuals, we represent the number and percentage of individuals that fit, in green are models with 100% of the individuals that fit and red otherwise.

| Iranian-related populations | 22 ASI | Indus Periphery West |
|---|---|---|
| *Sarazm_EN* | 22 (100%) | 0.004 |
| *Sarazm_EN_2* | 22 (100%) | 0.012 |

Next, we tested the model with *Sarazm_EN_2* as the proxy of Iran farmer-related ancestry for all individuals on the Indian cline. Using the three-way model with Iranian farmer-related (*Sarazm_EN_2)*, Steppe pastoralist-related (*Central_Steppe_MLBA*) and *AHG*-related ancestries, we find this models fits 96% of the individuals on the Indian cline, which is similar to the model with two *Sararm_EN* individuals (Table S4.3, S4.7). Moreover, *Sarazm_EN_2* also provides a good fit for 94% of the individuals that fall outside the Indian cline (Table S4.8). This is even slightly better than using Sarazm_EN which fits for 91% of the off-cline individuals (Table

S4.4). Together, our analysis shows that Sarazm_EN is the most likely source of Iranian farmer-related ancestry for ANI, ASI, Austroasiatics-related and East Asian-related individuals in India.

**Table S4.7 Testing of *Sarazm_EN_2* for the Iran farmer-related ancestry for individuals on the Indian cline (n=2,126).** As described in Table S4.3, we used qpAdm with two-way: *AHG*-related + *Iranian farmer*-related ancestry or three-way: *AHG*-related + Iranian farmer-related ancestry + *Steppe pastoralist*-related (Central_Steppe_MLBA) ancestries. Source for Iranian farmer-related ancestry shown below. We use the acceptance criterion of p-value > 0.01.

| **Iranian farmer-related group** | **Model with *p*-value > 0.01 (count / %)** | **Model with p-value > 0.01 & mixture coefficients > 0 (count / %)** | **Comment** |
|---|---|---|---|
| *Sarazm_EN_2* (either two-way or three-way model | 2043 / 96.1% | 2043 / 96.1% | |
| *Sarazm_EN_2* (three-way model) | 2042 / 96.0% | 1967 / 92.5% | 75 individuals have negative coefficients for *Steppe pastoralist*-related ancestry |
| *Sarazm_EN _2* (two-way model) | 825 / 38.8% | 825 / 38.8% | |

**Table S4.8 Best fitted model for the 494 off cline individuals using Sarazm_EN_2 as a proxy for *Iran farmer*-related ancestry.**

| Best model | *Model a* | *Model b* % (count / ASI-end total) | *Model c* % (count / EAS-related total) | *Model c + Steppe pastoralist-related* % (count / EAS-related total) | No fit* % (count / total) |
|---|---|---|---|---|---|
| Number of Individuals | 316 | 96% (55/57) | 38% (46/121) | 38% (46/121) | 6% (31/494) |

*We define **Model *a***: *Sarazm_EN_2* + *AHG*-related + Steppe pastoralist-related ancestries (same as Indian cline), **Model b**: *Sarazm_EN_2* + *AHG*-related + Austro-asiatic-related (using *Nicobarese* as proxy), and **Model c**: *Sarazm_EN_2* + *AHG*-related + East Asian-related (using *CHB* as proxy). No fit describes the set of individuals for whom none of the above four models provide a good fit (*p*-value > 0.01 with non-negative ancestry coefficients in *qpAdm*). This includes 29 individuals in the East Asian-related cluster and two in the Austro-asiatic related cluster.

*Estimating the genome-wide ancestry proportions*

Using the three-way model with Iranian farmer-related (*Sarazm_EN*), *AHG-related* and Steppe pastoralist-related (*Central_Steppe_MLBA)* ancestries, we infer the ancestry proportions for individuals on the Indian cline (considering only *p*-value > 0.01 and considering only models with non-negative mixture proportions). The ancestry coefficients are highly variable across India, with 26.6-67.6% deriving from *Sarazm_EN*-related, 0-45.1% from Steppe pastoralist-related and 18.6-68.6% of *AHG*-related groups (Figure S4.9). These estimates are highly correlated with previous results with *Indus Periphery West* as the source for Iran farmer-related ancestry (Pearson's correlation coefficients: $r^2_{\_Iran\text{-}farmer\text{-}related}$ >0.98, $r^2_{\_AHG\text{-}related}$ > 0.995 and $r^2_{\_Steppe\text{-}Pastoralist\text{-}related}$ > 0.992). Notably, mixture proportions vary significantly across India and are associated with the region of birth, linguistic affiliation and caste affiliation (Figure S4.9).

To further understand the patterns of variation in ancestry proportion across India, we plotted the mixture proportions against the PCA results from Figure 1. The *AHG*-related ancestry is most correlated to the PCA loadings or Indian cline (Table S4.9). Interestingly, we find the Iranian farmer-related ancestry is almost uniform across India, while *AHG*-related and Steppe pastoralist-related ancestries seem anticorrelated. The variation in these ancestries follows a North-South gradient with the highest Steppe pastoralist-related ancestry in the North (18.8%, p-value<2e-16) and the lowest in the South (4.2%, p-value<2e-16). The *AHG*-related ancestry has the opposite trend with the highest coefficients in the South (50.0%, p-value<2e-16) and the lowest in the North (34.7%, p-value<2e-16). The individuals on the ASI-end of the cline have mostly Iranian farmer-related and *AHG*-related ancestries, with Steppe pastoralist-related ancestry coefficients close to zero (Fig S4.10). At the opposite end of the cline, individuals have mostly Iranian farmer-related and Steppe pastoralist-related ancestries. We obtain similar results when we use Indus Periphery West instead of Sarazm_EN as the source for Iranian-farmer related ancestry in India (Fig S4.10). This provides insights into the ordering of the two gene flow events in India, indicating the Steppe pastoralist-related mixture occurred in the already mixed groups with *AHG*-related and Iranian farmer-related groups as also shown by ancient DNA studies[34].

**A**

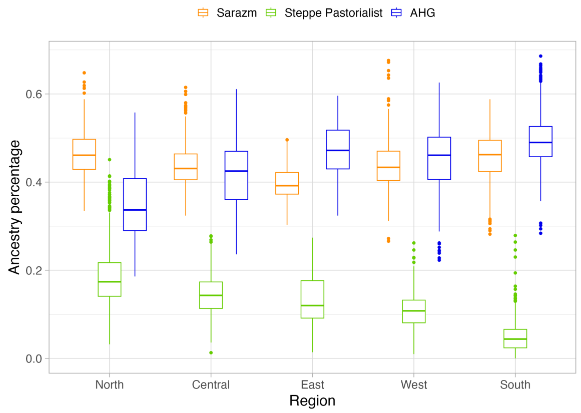

**B**

**C**

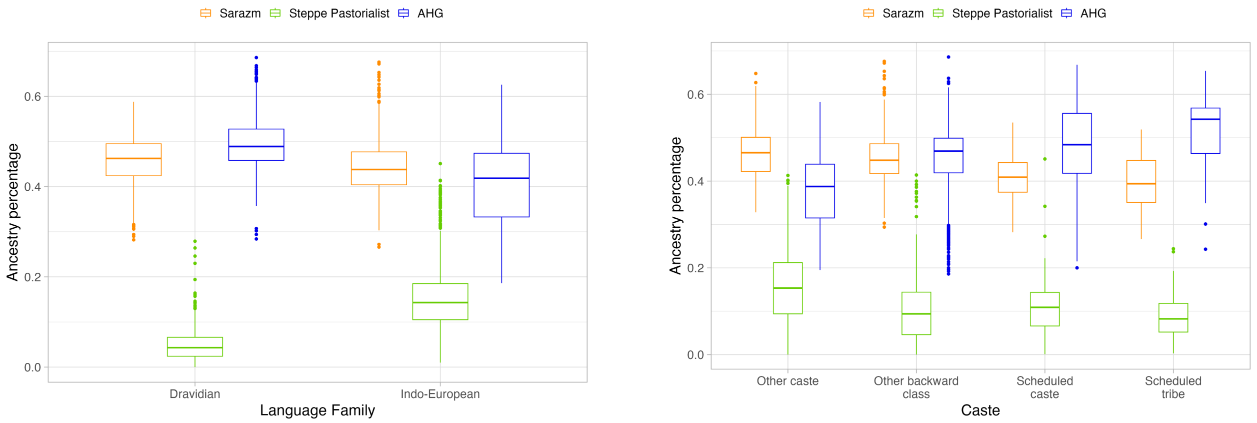

**Figure S4.9 Ancestral population-related coefficients using the revised model.** Inferred coefficients based on *qpAdm* using the three-way model with *Sarazm_EN*, *Central_Steppe_MLBA* and *AHG*-related groups shown by (A) region, (B) language family and (C) caste group. We show only results for 1,942 individuals for whom the three-way model was a good fit (*p*-value > 0.01 and inferred ancestry proportions were non-negative).

**A**

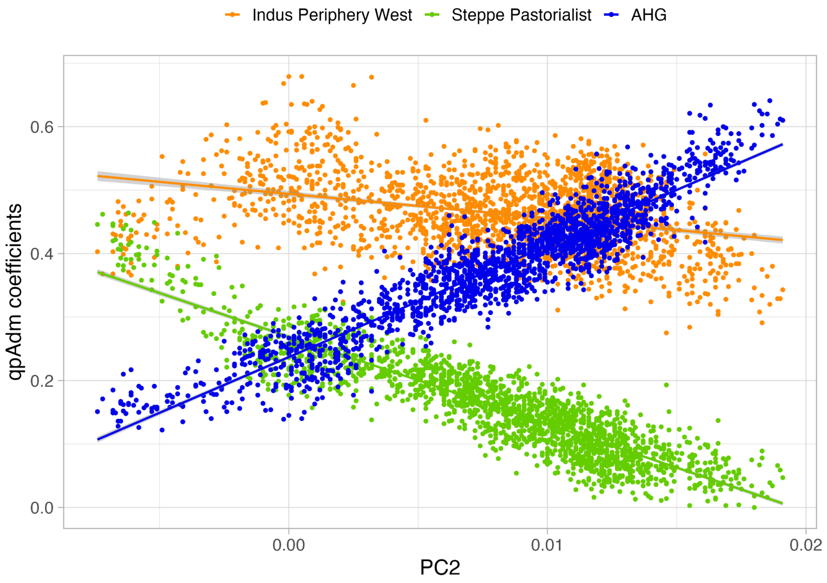

**B**

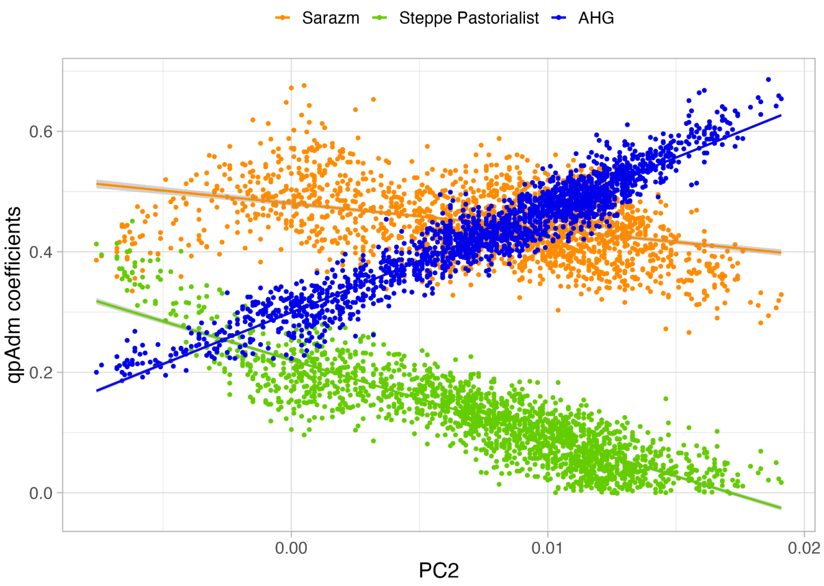

**Figure S4.10. *qpAdm* ancestry coefficients in function of the position on the Indian cline**. We represent the position on the Indian cline by the PC2 value of the PCA in Figure 1. We show inferred ancestry proportions using the three-way model with *AHG*-related, *Central_Steppe_MLBA*-related and Iranian farmer-related group either (A) *Indus Periphery West (n=1,969)* or (B) *Sarazm_EN (n=1,942)*. We performed generalized linear model (*glm*) regression for each coefficient against the PC2 value of each individual, all regressions are significant (*p*-value < *2e-16*).

**Table S4.9 Correlation between ancestry coefficients and position on the Indian cline.** Correlation coefficients between the PC2 value in Figure 1 and ancestry coefficients (using the three-way model with Iranian farmer-related, Steppe pastoralist-related and *AHG*-related). We use either *Indus Periphery West* or *Sarazm_EN* as the source for Iranian farmer-related ancestry. All correlations were computed with a Pearson's t-test and have a p-value < 2.2e-16.

| **Models/ ancestry** | **Indus Periphery -related coefficients** | **Steppe pastoralist -related coefficients** | ***AHG*-related coefficients** |
|---|---|---|---|
| *Indus Periphery West* + *AHG*-related + *Central_Steppe_MLBA* | -0.362 | -0.905 | 0.941 |
| *Sarazm_EN* + *AHG-related* + *Central_Steppe_MLBA* | -0.371 | -0.879 | 0.958 |

**Population structure in West Bengal**

West Bengal is one of the most represented states among the individuals outside the Indian cline including 140 individuals (out of 494 off-cline individuals) (Fig 1, Figure S4.2). Individuals from West Bengal show clear East Asian-relatedness on the PCA. Indeed, 90% of these individuals have non-negative East Asian-related ancestry (p-value>0.01 and non-negative coefficients using *qpAdm* and model c or c + *Steppe pastoralist*-related). To infer the timing of the East Asian-related admixture, we applied ALDER[30]. This method measures the weighted linkage disequilibrium (LD) in the population of interest related to a reference population to infer the proportion and timing of admixture. We ran ALDER in the 'one-reference' mode using *CHB* from the AADR dataset as the reference population to represent the East Asian-related ancestry component. Standard errors were inferred using chromosome jackknife where one chromosome is dropped in each run. We infer the East Asian-related admixture occurred 51 ± 5 generations ago ($z$ = 11.19). This translates to 1,428 years or 522 AD, assuming the human generation time of 28 years[35]. ALDER also infers a lower bound on the East Asian-related ancestry proportion of 6.1 ± 0.6% (Figure S4.11). The date and proportion of East Asian-related admixture in Bangladeshis (who are genetically similar to Bengalis from West Bengal) is consistent with previous findings of 52.2 ± 2 generations ago and 9.3 ± 2.6%[36].

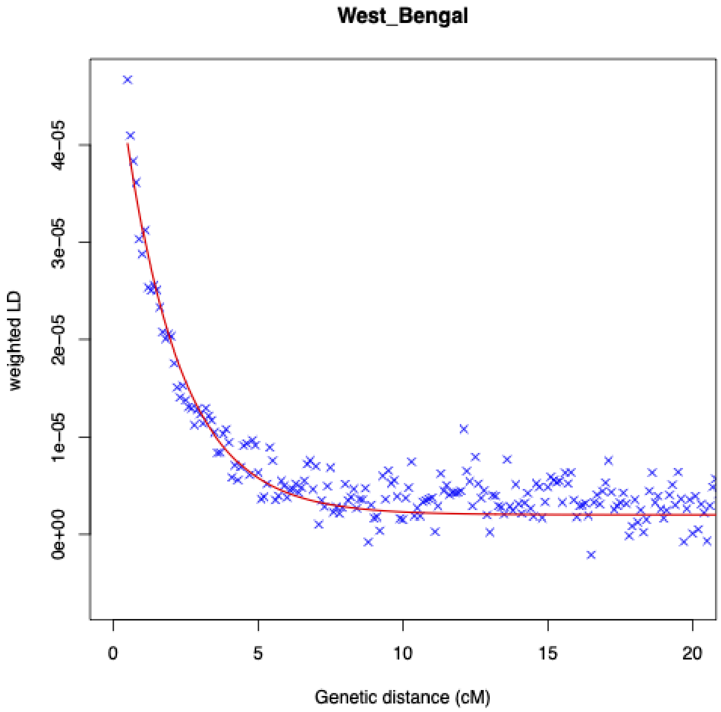

**Figure S4.11: Decay of admixture linkage disequilibrium in West Bengal individuals.** We ran ALDER with West Bengal individuals as the target population using the 'one-reference' model with East Asians (*CHB*) as the reference population. For details, see Methods.

### Supplementary Note 5: Founder events and consanguinity in India

Founder events and consanguineous marriages increase the relatedness of individuals in a group. As a consequence, individuals share large genomic regions inherited identical-by-descent (IBD) from a common ancestor. A special case of IBD within an individual—i.e., when two chromosomes of an individual share IBD— leads to homozygous-by-descent (HBD). The total amount (proportion of the genome) and length of IBD and HBD segments is informative about the history of population size changes over time. Intuitively, many and long shared regions of HBD in a population indicate small population size, while few and short segments reflect large population size.

We identified IBD and HBD segments using *hap-IBD*[37] (see Methods). To minimize false positives, we only considered segments with length greater at 2 cM and we filtered out segments that overlapped centromeres (using the GRCh38/hg38 annotation from genome.ucsc.edu/cgi-bin/hgTables). We analyzed the following datasets:

- LASI-DAD phased data without a reference panel (Supplementary Note S3) after excluding first degree related individuals and offspring of trios (2,620 individuals).
- The high coverage phased 1000G dataset[19] after excluding the offspring of trios from the analysis (2,590 individuals).

**Distribution of HBD in India**

In the main text, we report the results for the 8 cM threshold to identify long HBD segments (Figure 2.A). Here, we use 20 cM to define long HBD segments and show the conclusions are robust to the choice of the threshold (Figure S5.1). On average, Indians have a larger fraction of their genome in HBD segments (~29 cM) compared to 1000G EAS (~6 cM), EUR (~6 cM), and AFR (~4 cM) and even in long HBD segments (~1.6 cM) compared to 1000G EAS (~0.04 cM), EUR (~0 cM), and AFR (~0.04 cM). Within India, individuals from South have significantly higher homozygosity, both in terms of the total amount of their genome in HBD segments (on average, ~56 cM in South compared to ~19 cM in other regions, p-value < $10^{-16}$) and the fraction of long HBD segments (6.3% vs. 0.4%, p-value = 0.00913), reflecting the higher prevalence of consanguineous marriages in the South of India[38] (Figure 2A, Figure S5.2). Overall, less than 1% of Indians have HBD segments longer than 20cM (Figure S5.2B), compared to ~0.1% in other regions. We obtain qualitatively similar results with 8cM HBD threshold (Figure S5.2A).

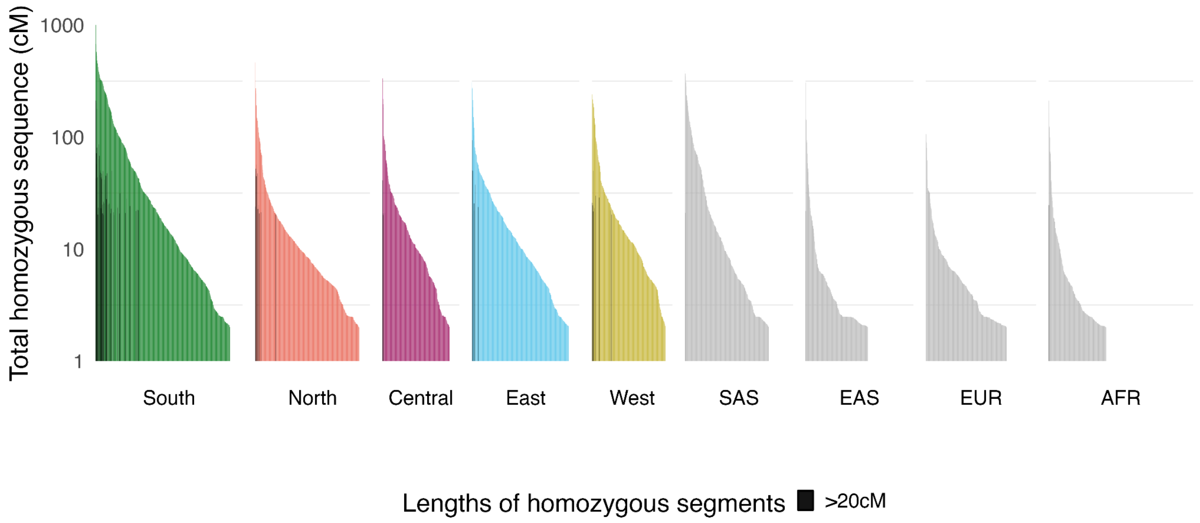

**Figure S5.1: Homozygosity per individual across worldwide populations.** Each vertical bar is an amount of genome (in cM) in HBD segments in each individual's genome. The amount of HBD is stratified by size, greater than 20cM (in darker colors) and less than 20cM (in lighter colors). Colors represent regions in India for LASI-DAD and superpopulation for 1000G individuals.

**A**

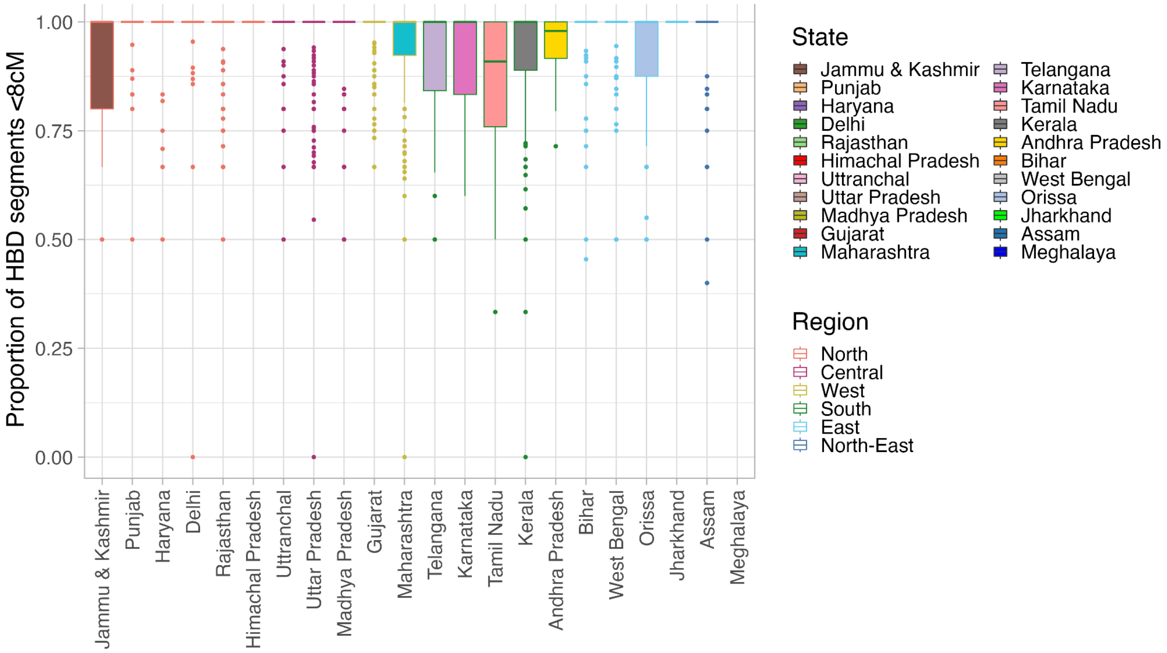

**B**

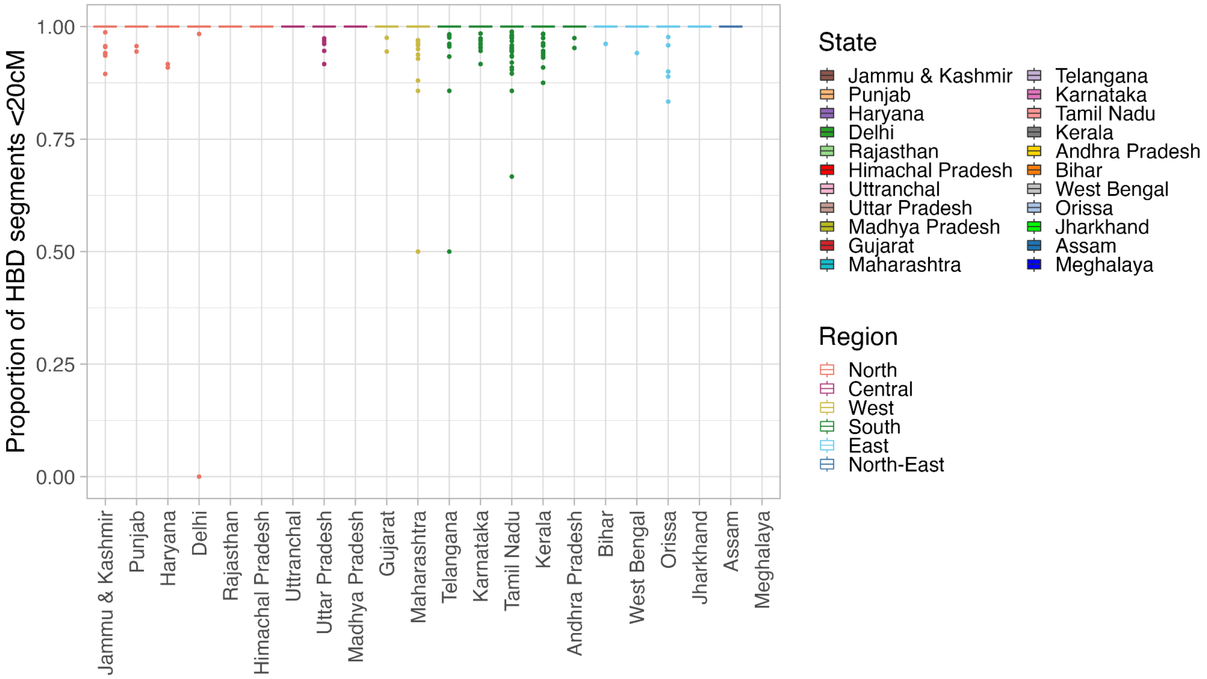

**Figure S5.2: Proportion of long HBD segments per individual.** We grouped individuals by their state of birth and we show the fraction of the genome in HBD segments deriving from long HBD, using two thresholds: (A) Threshold of 8cM and (B) threshold of 20cM. Colors used to fill the box plots reflect the state of birth and the outline reflects the region in India.

### Distribution of IBD sharing in LASI-DAD and 1000G SAS

We find that the overall relatedness is higher among individuals in the LASI-DAD dataset vs. among individuals in the 1000G SAS population (though no systematic difference in structure is observed in PCA (Figure 1B)). Some potential reasons leading to this difference may include: 1) four-fold higher sample size of LASI-DAD, and 2) ascertainment bias of selecting individuals in each study. To investigate these possibilities, we first performed bootstrap resampling where we randomly selected 500 individuals (similar to 1000G SAS) from the LASI-DAD dataset and estimated the mean and the 95% confidence interval across runs (Figure 2B). As expected, measures of the relatedness depend on sample size. With 500 individuals, we find ~24.2% (19.4-28.6%) of the individuals have a third-degree cousin or closer in the dataset. This number is still higher than in 1000G SAS (~14.2%). However, the fraction of individuals that share IBD equivalent to a 2nd and 1st degree cousin is qualitatively similar in 1000G SAS and down-sampled LASI-DAD datasets (Table S5.1).

**Table S5.1 Percentage of individuals having a kth-degree cousin or closer in the dataset.** For each of the AFR, EAS, EUR and SAS individuals in 1000G and the 2,620 Indian samples from LASI-DAD (full dataset and bootstrapped to 500 individuals). We present the 95% confidence interval in parenthesis for the bootstrap analysis.

| Dataset | IBD equivalent to **1st degree** cousin or closer (*>847.75cM*) | IBD equivalent to **2nd degree** cousin or closer (*>221.94cM*) | IBD equivalent to **3rd degree** cousin or closer (*>52.98cM*) | IBD equivalent to **4th degree** cousin or closer (*>13.25cM*) | IBD equivalent to **5th degree** cousin or closer (*>3.31 cM*) |
|---|---|---|---|---|---|
| 1000G AFR (*n=686*) | 0% | 3.8% | 17.2% | 57.3% | 100% |
| 1000G EAS (*n=512*) | 1.9% | 2.7% | 8.8% | 87.5% | 100% |
| 1000G EUR (*n=525*) | 0.4% | 2.3% | 8.8% | 94.3% | 100% |
| 1000G SAS (*n=514*) | 0.6% | 3.7% | 14.2% | 97.4% | 100% |
| LASI-DAD (*n=2,620*) | 3.1% | 13.2% | 51.0% | 100% | 100% |
| LASI-DAD bootstrap* (*n=500)* | 0.4% (0-1.6%) | 3.0% (1.2- 5.2%) | 24.2% (19.4-28.6%) | 100% (100-100%) | 100% (100-100%) |
| LASI-DAD cross-SSU comparison (*n=365-699)*[§] | 0-0.5% | 1.0-3.4% | 16.4-35.0% | 100-100% | 100-100% |

Note: * shows 95% CI in brackets based on bootstrap resampling; § shows the range across regions in India with the smallest sample size in Central (*n*=365) and largest in South (*n*=699), we do not include North-East because of its small sample size (*n=71*).

Another confounder for the comparison is the selection of samples in each study. In LASI-DAD, we used a stratified random sampling strategy. We first selected states and computed sampling weights to represent the population at the national level. Then we selected Sampling Secondary Units (SSUs) (villages/urban census blocks) from selected states and within each SSUs, we randomly selected individuals. To understand the impact of this complex sampling strategy on IBD sharing in India, we inferred the closest genetic relative accounting for the SSU, allowing only cross-SSU comparisons (and not within SSU) (Figure S5.4). Specifically, for each individual in LASI-DAD, we inferred the closest genetic relative only allowing comparisons of pairs of individuals from different SSUs. In this case, we inferred that 31.6% of the individuals share IBD equivalent to a 3rd cousin or closer (from 16.4% if only considering Central individuals to 49.3% if only considering North-East individuals or 34.8% if only considering West individuals), which is more than in SAS 1000G and for some values, above the 95% confidence interval obtained after bootstrapping the whole dataset to a smaller sample size (*n*=500) (Table S5.1). After accounting for the sample size and ascertainment scheme, we continue to find a significant shift in LASI-DAD compared to 1000G SAS, indicating limitations of the sampling of South Asian variation in 1000G.

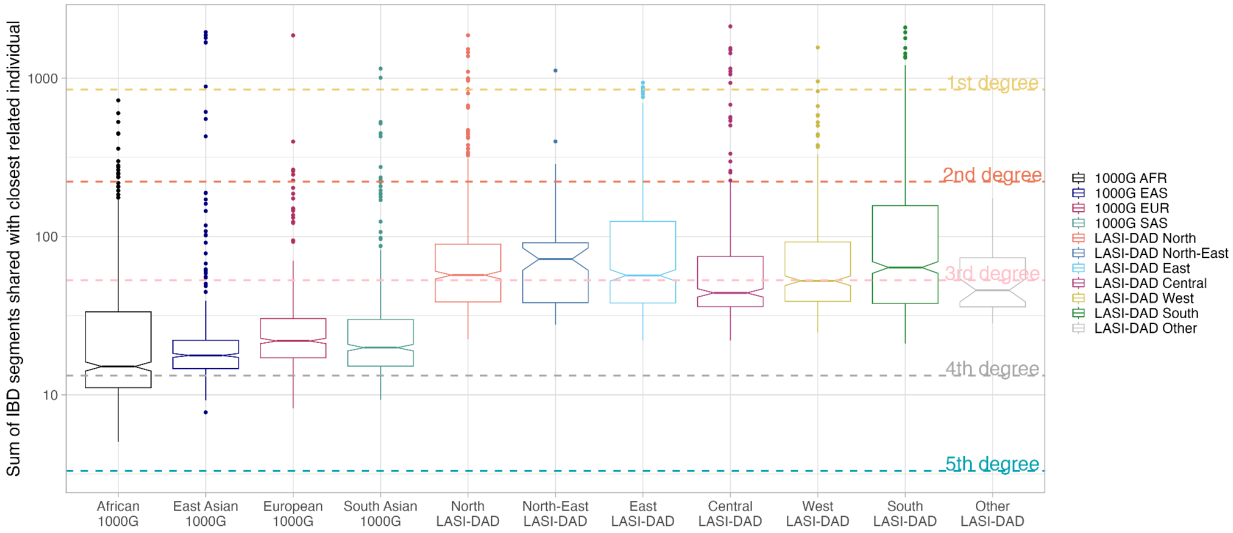

**Figure S5.3: IBD sharing between each individual and its inferred closest related individual.** For each of the 2,620 Indian samples from LASI-DAD and AFR, EAS, EUR and SAS individuals in 1000G, we detected the individual sharing the largest total amount (in cM) of genome IBD, referred to as 'closest related individual'. On the Y-axis we report the total shared genome (in cM). The horizontal dashed lines indicate the expected value of the total IBD sharing for kth degree cousins (see description), in brown k=1, purple k=2, red k=3, green k=4 and in yellow k=5.

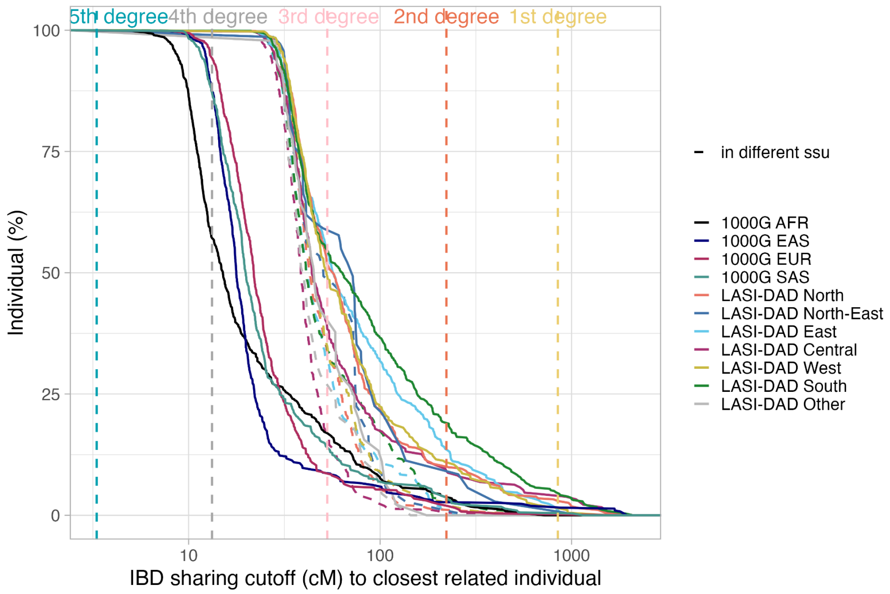

**Figure S5.4: Impact of sampling strategy on the genetic relatedness inferred between individuals.** For each of the 2,620 Indian samples from LASI-DAD and AFR, EAS, EUR and SAS individuals in 1000G, we inferred the total amount of IBD (in cM) shared among individuals and inferred the closest genetic relative for each individual with expected IBD sharing equivalent to a k-th degree cousin or closer. For each value *x* of total IBD shared genome (in cM) on the *X*-axis, we report the percentage of individuals (*Y*-axis) that share *x* or more with their closest individual. For LASI-DAD individuals, we also detect the closest individuals sampled in a different SSUs (dashed lines). The vertical dashed lines indicate the expected value of the total IBD sharing for *k*th degree cousins (see description), *k*=1 (in yellow), *k*=2 (in orange), *k*=3 (in pink), *k*=4 (in gray) and *k*=5 (in blue).

#### Functional impact of founder events in LASI-DAD

We compare the number of missense and pLoF variants in LASI-DAD and gnomAD (Table S5.2) and assess its relative burden of HBD per individual in LASI-DAD (Figure S5.5).

**Table S5.2 Missense and pLoF variants in LASI-DAD dataset compared to gnomAD database.**

| Category | Total count | pLoF | Missense | Nb singletons | Nb homozygote ALT/ALT (only pLoF) | In ClinVar* (only pathogenic) |
|---|---|---|---|---|---|---|
| Total | 406,304 | 20,319 | 385,985 | 253,166 | 27,120 (499) | 17,538 (745) |
| South Asian specific | 210,126 | 13,721 | 196,405 | 160,858 | 2,413 (106) | 2,307 (240) |
| LASI-DAD specific | 160,856 | 11,575 | 149,281 | 140,488 | 1,013 (53) | 1,115 (163) |
| $f_{LASI\text{-}DAD} > 5\%$ | 14,021 | 260 | 13,761 | 0 | 10,013 (121) | 1,891 (0) |

| | | | | | | |
|---|---|---|---|---|---|---|
| $f_{LASI\text{-}DAD}$ > 0.1% | 63,242 | 1,839 | 61,403 | 0 | 24,548 (399) | 6,788 (34) |
| $f_{LASI\text{-}DAD}$ > 0.1% & South Asian specific | 9,560 | 304 | 8,755 | 0 | 1,053 (36) | 391 (10) |
| $f_{LASI\text{-}DAD}$ > 0.1% & LASI-DAD specific | 501 | 20 | 481 | 0 | 55 (5) | 11 (1) |

*review status of at least 2 stars, we define as pathogenic variants with ClinVar status : Pathogenic and/or Likely pathogenic

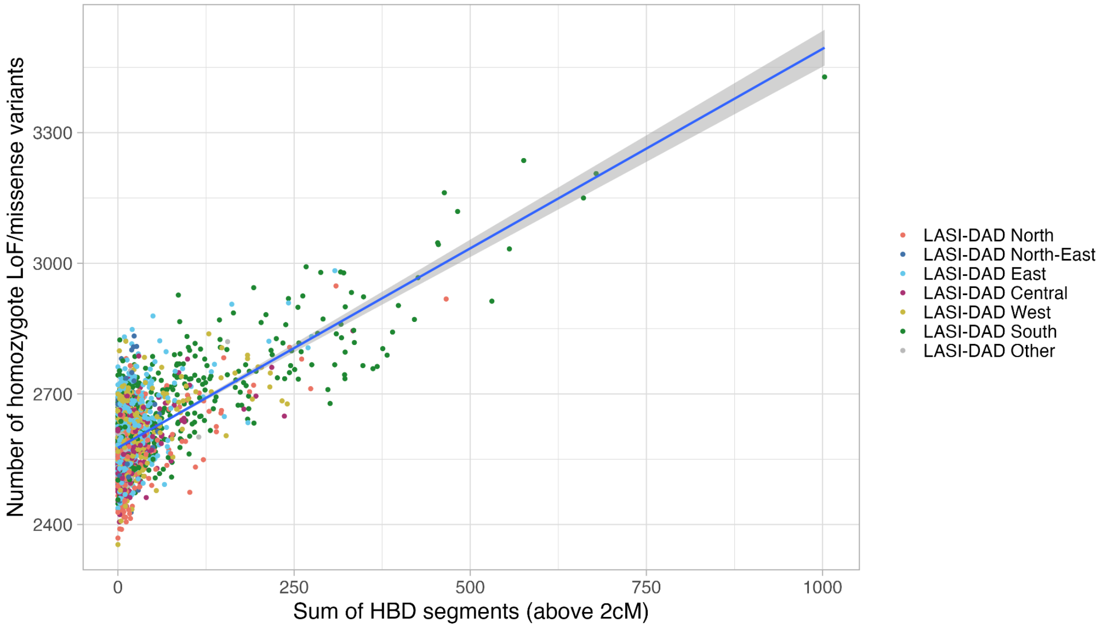

**Figure S5.5: Relationship between the number of homozygous derived missense/pLoFs and the total sum of HBD segments per individual.** Individuals are colored by region of birth. We fit a regression using generalized linear model (*glm*) and obtain the following fit: $y = 2576 + 0.916*x$.

### Supplementary Note 6: Inference of archaic ancestry

Previous studies have shown that South Asians have archaic ancestry from both Neanderthals and Denisovans[39]. To learn about the genomic landscape and regional variation in archaic ancestry in Indians, we applied a previously published hidden Markov model called hmmix, which does not rely on archaic reference genomes[40]. In turn, it uses an outgroup, such as sub-Saharan Africans who have negligible amount of archaic ancestry[41,42]. Briefly, we applied *hmmix*[40] to LASI-DAD and other worldwide datasets (shown below) to search for archaic ancestry segments in non-African individuals. We further compared archaic segments previously published for 27,566 individuals from Iceland that were inferred using *hmmix*[43]. Below we show the sample composition with sample sizes across geographic regions for each dataset.

**Table S6.1 Datasets used for analysis of archaic ancestry in non-Africans.** Number of individuals in each superpopulation and dataset. For South Asians, we show regional distribution of individuals.

| Superpopulation | 1000 Genomes (1000G) | Human Genome Diversity Project (HGDP) | LASI-DAD | deCODE |
|---|---|---|---|---|
| East Asians | 585 | 223 | | |
| Europeans | 633 | 155 | | 27,566 |
| South Asians | 601 | 197 | 2,679 | |
| | 131 Bengali (Bangladesh)<br>103 Gujarati Indian (United States (US)<br>107 Indian Telugu (United Kingdom (UK))<br>146 Punjabi (Pakistan)<br>114 Sri Lankan Tamil (UK) | 24 Balochi (Pakistan)<br>25 Brahui (Pakistan)<br>24 Burusho (Pakistan)<br>19 Hazara (Pakistan)<br>22 Kalash (Pakistan)<br>25 Makrani (Pakistan)<br>24 Pathan (Pakistan)<br>24 Sindhi (Pakistan)<br>10 Uygur (China) | 373 (Central India)<br>530 (East India)<br>555 (North India)<br>73 (North-East India)<br>48 (Other India)<br>715 (South India)<br>385 (West India) | |
| Americans | 490 | 61 | | |
| Middle Easterns | | 161 | | |
| Oceanians | | 28 | | |

Briefly, our pipeline for identifying archaic ancestry in non-Africans had the following steps:

1. First, we removed variants not found in 1000G strict callability mask[19]. This was applied to HGDP, 1000G and LASI-DAD dataset. The callability mask was downloaded from:

http://ftp.1000genomes.ebi.ac.uk/vol1/ftp/data_collections/1000_genomes_project/working/20160622_genome_mask_GRCh38/. The callability mask covers 2,164,834,110 bp on the autosomes corresponding to 75.3% of the total human genome reference sequence on the autosomes of 2,875,001,522 bp. The total amount of uncalled sequence (gaps) is 710,167,412 bp. The mean length of gaps is 184 bp while the median is 53 bp. A total of 385,923,722 bp are located in gaps smaller than 1 kb which is the window size used by hmmix. Therefore, a conservative estimate of the accessible genome is 2,164,834,110 bp while a more realistic estimate of the accessible genome is 2,489,077,800 bp.

2. We polarized all variants with respect to the human ancestral allele and excluded any sites where the ancestral allele was missing. The human ancestral allele information was downloaded from: http://ftp.ensembl.org/pub/current_fasta/ancestral_alleles/homo_sapiens_ancestor_GRCh38.tar.gz
3. We further removed all variants where the derived allele is present in the outgroup. The outgroup consists of African individuals from 1000G and HGDP. We picked 426 individuals from the 1000G[19] including Yoruba in Ibadan, Nigeria (YRI), Mende in Sierra Leone (MSL), Esan in Nigeria (ESN) and 64 Africans from HGDP who have less than 1% west Eurasian admixture[23], including Bantu South Africa, Biaka Pygmy, Mbuti Pygmy, San and Yoruba.
4. Finally, we estimated the local mutation rate in 1 million bp (Mb) windows using all variants present in the outgroup ($n$=490). After removing variants from the outgroup, we retained 140,925 SNPs per individual before phasing (Supplementary Note S3) and 141,042 variants after phasing (Table S6.2). Note, the increase in variants post-phasing is likely because some missing variants were imputed during phasing.

**Table S6.2. Average number of variants per genome (before and after phasing) for different datasets and geographical regions.**

| Region | Dataset | Variants per genome | Phased variants per genome |
|---|---|---|---|
| Americas | 1000G | 128,681 | 121,868 |
| | HGDP | 134,854 | 134,459 |
| South Asia | 1000G | 145,355 | 133,470 |
| | HGDP | 138,678 | 138,263 |
| | LASI-DAD | 140,925 | 141,042 |
| East Asia | 1000G | 146,998 | 133,937 |
| | HGDP | 148,152 | 147,667 |
| Europe | 1000G | 133,412 | 124,526 |

| | | | |
|---|---|---|---|
| | HGDP | 133,820 | 133,499 |
| Middle East | HGDP | 121,504 | 121,075 |
| Oceania | HGDP | 179,142 | 178,829 |

***HMM Training***

We trained hmmix on all individuals separately from each region from each dataset using the Baum-Welch algorithm[40]. We terminated the training when the log likelihood of the new emission, transition and start parameters improved less than 0.001 (in units of the log likelihood) compared to the likelihood of the parameters in the previous iteration. We did this both for unphased data and phased data.

The trained parameters for unphased data are shown in Figure S6.1 and for phased data in Figure S6.2. We note that for phased data there is a noticeable batch effect when using 1000G data due to the lower number of phased variants (Table S6.2). We discuss these further in Supplementary Note S9.

The expected relationship between the human emission parameters for phased on unphased data is:

$$emission_{unphased,\ human\ state} = 2 \cdot emission_{phased,\ human\ state}$$

This is due to the fact that we are counting derived variants on two chromosomes for diploid data. The factor of 2 is valid under the assumption that all SNPs are heterozygous. If all SNPs are homozygous the emission parameters for phased and unphased data will be identical.
For the individuals from India (LASI-DAD) the average emission parameter for the human state is 0.0508 SNPs pr 1000 kb for the unphased data and 0.0261 SNPs pr 1000 kb for the phased data. The ratio of unphased data/phased data is 1.94 in close agreement with the expectation of 2 - indicating that a majority of derived variants are found are heterozygous.

The expected relationship between the archaic emission parameters for phased on unphased data is:

$$emission_{unphased,\ archaic\ state} = emission_{phased,\ human\ state} + emission_{phased,\ archaic\ state}$$

This is valid under the assumption that a all archaic introgressed segments are in a heterozygous state. For the individuals from India (LASI-DAD) the average emission parameter for the archaic state is 0.375 SNPs pr 1000 kb for the unphased data and 0.333 SNPs pr 1000 kb for the phased data. The difference between unphased and phased data is 0.042 SNPs pr 1000 kb in close to the phased human emission parameter of 0.0261 SNPs pr 1000 kb.

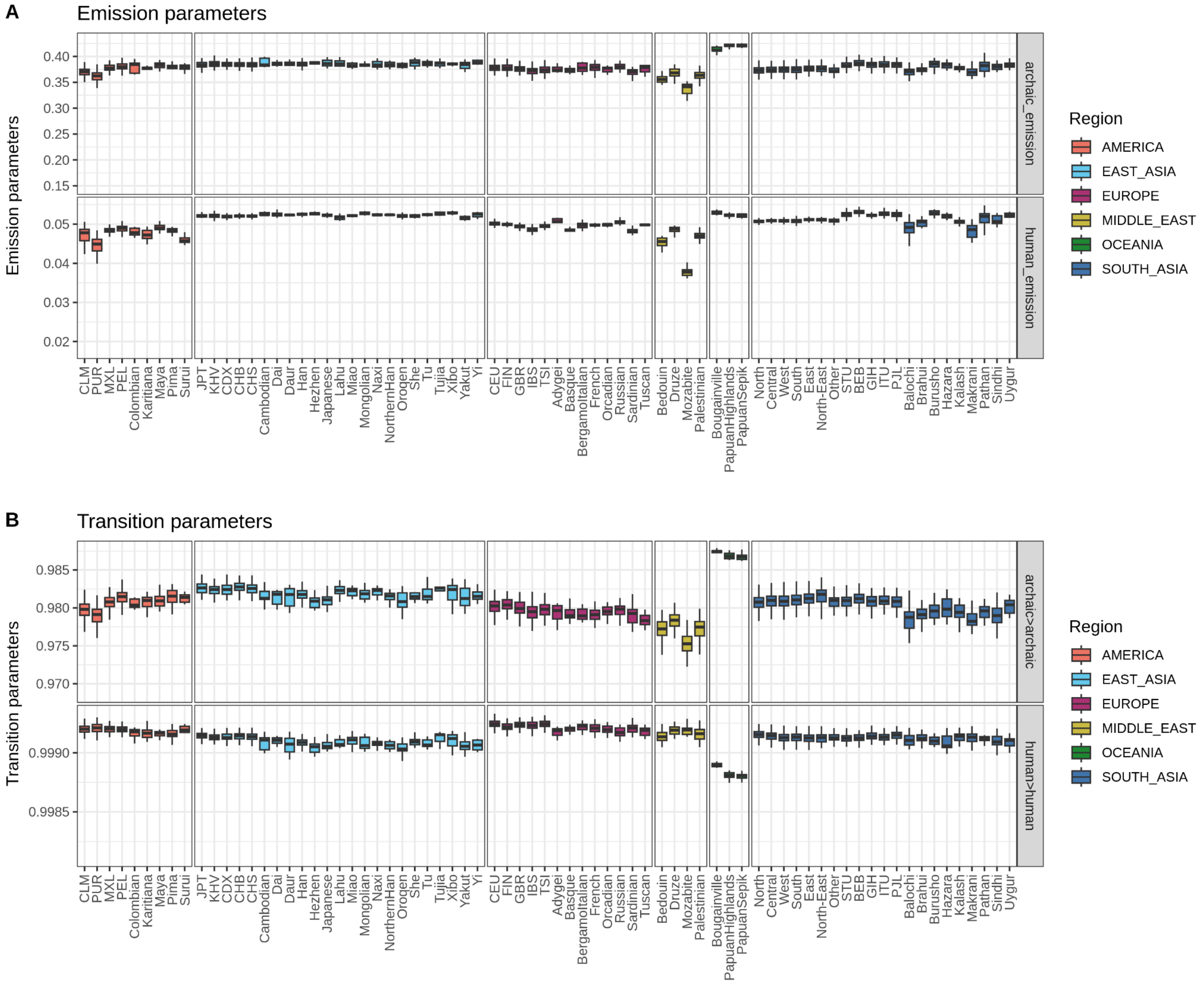

**Figure S6.1. Trained emission and transition parameters for unphased data.** We show the values for the trained parameters on unphased data for each population colored by region in each dataset. The populations are sorted by dataset with individuals from LASI-DAD (North, Central, West, South, East, North-East, Other) first followed by 1000G (three capital letter population names) and the HGDP individuals. We show the archaic and human emission parameters in panel A and the transition parameters for the archaic->archaic state and human->human state in panel B.

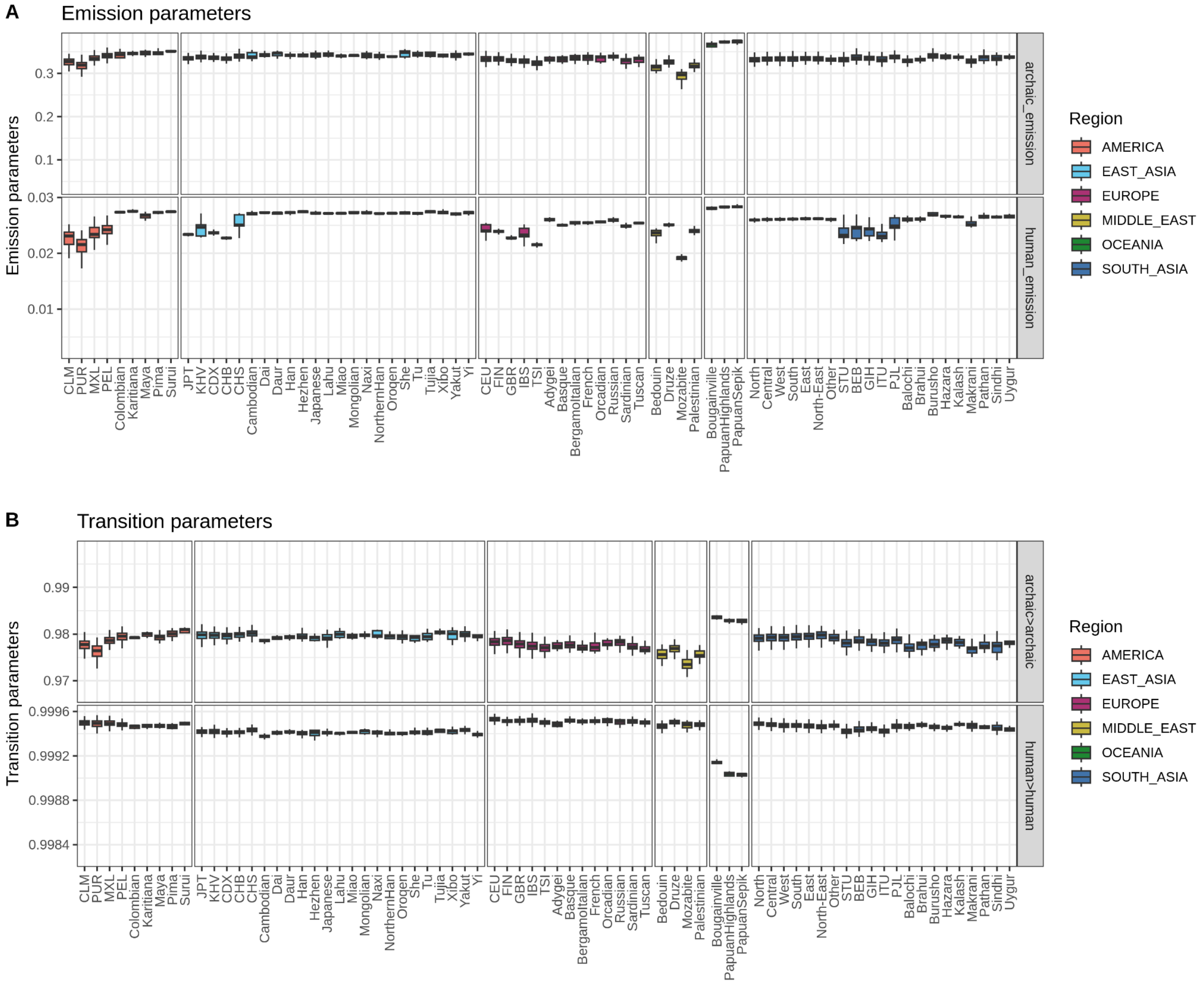

**Figure S6.2. Trained emission and transition parameters for phased data.** We show the values for the trained parameters on phased data for each population colored by region in each dataset. The populations are sorted by dataset with individuals from LASI-DAD (North, Central, West, South, East, North-East, Other) first followed by 1000G (three capital letter population names) and the HGDP individuals. We show the archaic and human emission parameters in panel A and the transition parameters for the archaic->archaic state and human->human state in panel B.

#### *HMM Decoding*

For the decoding step, we used the average of the trained parameters for each region for each dataset separately and applied them to identify archaic segments. We identified archaic regions as consecutive 1000 bp windows with a posterior probability of being in the archaic state which is greater than 0.5. To classify the source of the archaic ancestry, we used the four published high coverage archaic genomes:

Denisova 5 (Altai Neanderthal)[44]
Callable genome:
http://ftp.eva.mpg.de/neandertal/Vindija/FilterBed/Altai/ (1,657,402,507 bp)

Chagyrskaya 8 (Chagyrskaya Neanderthal)[45]
Callable genome:
http://ftp.eva.mpg.de/neandertal/Chagyrskaya/FilterBed/ (1,635,657,520 bp)

Vindija 33.19 (Vindija Neanderthal)[46]
Callable genome:
http://ftp.eva.mpg.de/neandertal/Vindija/FilterBed/Vindija33.19/ (1,654,299,331 bp)

Denisova 3 (Altai Denisovan)[47]
Callable genome:
http://ftp.eva.mpg.de/neandertal/Vindija/FilterBed/Denisova/ (1,648,754,299 bp)

We used two different approaches for classifying archaic segments.

1. *Approach 1:* We classify segments into four categories:

   **Denisovan**: Segments that share more derived alleles with Altai Denisovan than any of the Neanderthals.
   **Neanderthal:** Segments that share more derived alleles with any of the sequenced Neanderthals than Altai Denisovan genome.
   **Both:** Segments that share an equal number of derived variants with Altai Denisovan and the Neanderthal genomes.
   **None**: Segments which do not share any derived variants with any of the sequenced archaic genomes.

This is equivalent to the approach in [43] and has the advantage of also recovering mosaic archaic segments in modern humans[43] i.e., segments where a part of the segment shares more derived alleles with the Altai Denisovan genome and another part shares more derived alleles with the Neanderthal genomes.

2. *Approach 2:* We classified segments into two categories based on the presence of three different site types, polarized by ancestral (=0) or derived (=1) alleles in Neanderthals (*N*) and Denisovans (*D*). ND01 sites where the derived allele is only found in Denisovans, ND10 sites where the derived allele is only found in Neanderthals and ND11 sites where the derived allele is found both in Neanderthals and Denisovans.

   **ND01:** Segments which only contain ND01 + ND11 sites.
   **ND10:** Segments which only contain ND10 + ND11 sites.
   **ND11:** Segments which only contain ND11 sites.
   **ND00:** Segments which do not share any derived variants with Neanderthals or Denisovans.
   **ND11_mosaic:** Segments which contain both ND10 and ND01 sites. This is the same category of sites as mosaic segments[43].

We defined different variant types:

- **DAV:** Derived Archaic variants. These are ND10, ND01 and ND11 sites.
- **linked-DAV**: These are SNPs that are in high linkage disequilibrium (LD) with DAV SNPs (LD is greater than 0.9).
- **unlinked-DAV:** These are SNPs that are in low LD with DAV SNPs (LD is less than 0.9).

**Table S6.3. Number of SNPs per variant types and dataset.** We report the number of ND01, ND10, ND11, DAV SNPs (ND10 + ND01 + ND11), DAV-linked variants, unlinked-DAV variants and number of individuals in each region (*n*).

| region | dataset | ND01 | ND10 | ND11 | DAV | Linked-DAV | unlinked-DAV | *n* |
|---|---|---|---|---|---|---|---|---|
| America | 1000G | 7258 | 165680 | 37952 | 210890 | 83295 | 1183463 | 490 |
| | HGDP | 3588 | 69467 | 16722 | 89777 | 32841 | 197976 | 61 |
| East Asia | 1000G | 23928 | 173015 | 43784 | 240727 | 95166 | 1534590 | 585 |
| | HGDP | 19103 | 167343 | 41136 | 227582 | 95698 | 878633 | 223 |
| Europe | 1000G | 5947 | 171558 | 38559 | 216064 | 82977 | 1335472 | 633 |
| | deCODE* | 6683 | 111336 | 28760 | 146779 | 47193 | 198558 | 27566* |
| | HGDP | 4507 | 150470 | 34059 | 189036 | 72143 | 577911 | 155 |
| South Asia | 1000G | 36676 | 238217 | 59477 | 334370 | 150762 | 1829989 | 601 |
| | HGDP | 18915 | 204835 | 49482 | 273232 | 120859 | 856679 | 197 |
| | LASI-DAD | 55815 | 266839 | 67176 | 389830 | 211568 | 5136368 | 2679 |
| Middle East | HGDP | 3903 | 138888 | 31248 | 174039 | 71157 | 655730 | 161 |
| Oceania | HGDP | 52311 | 81071 | 36348 | 169730 | 132069 | 262513 | 28 |

Note: *For deCODE Europeans, only variants not found in *repeatmasked* regions of the genome are considered.

**Inferred archaic ancestry per genome**

We first calculated the total amount of archaic sequence present in each non-African genome and compared it to the reference archaic genomes for phased and unphased data. We classified segments for the LASI-DAD dataset using both approach 1 in Table S6.4 and approach 2 in Table S6.6. We also show the results for HGDP and 1000G datasets merged by region in Table S6.5 for approach 1 and in Table S6.7 for approach 2. We note that the amount of archaic ancestry sequence differs between the phased and unphased analysis due to the homozygous introgressed archaic segments being counted twice when using phased data. If we merged the archaic segments called on each haploid genome and overlaid it with the segments called in the diploid genome, we find that 95.3% of the archaic sequences, which share at least one derived allele with any archaic reference genome, overlap showing a high concordance when using phased and unphased data. We used the phased version of the datasets for all downstream analyses.

From previous work, it has been shown that a high proportion of low posterior probability segments do not share any derived alleles with the sequenced archaic genomes, and are likely false positives[43]. At posterior probability cutoff from 0.8 to 0.9, we find >80% of archaic segments that overlap with Neanderthal and Denisovan genomes. At a lower posterior probability of 0.5, we retain 30% or greater proportion of archaic segments without any shared variants with archaic genomes (‘none’), suggesting that these variants could be enriched for false positives.

**Table S6.4. Average amount of archaic ancestry in MB and percent inferred using approach 1 for LASI-DAD dataset.** For phased and unphased data using posterior probability cutoffs at 0.5, 0.8 and 0.9. We assumed that the callable genome is 2 * 2,489,077,800 bp.

| Segment type | Region | Unphased | | | Phased | | |
|---|---|---|---|---|---|---|---|
| | | 0.5 | 0.8 | 0.9 | 0.5 | 0.8 | 0.9 |
| Denisova | North | 5.4 (0.1%) | 5.2 (0.1%) | 4.3 (0.1%) | 5.9 (0.1%) | 5.7 (0.1%) | 5 (0.1%) |
| | Central | 6.4 (0.1%) | 6.1 (0.1%) | 5.2 (0.1%) | 6.9 (0.1%) | 6.7 (0.1%) | 5.9 (0.1%) |
| | Other | 6.6 (0.1%) | 6.3 (0.1%) | 5.3 (0.1%) | 7 (0.1%) | 6.8 (0.1%) | 5.9 (0.1%) |
| | West | 7 (0.1%) | 6.7 (0.1%) | 5.7 (0.1%) | 7.5 (0.2%) | 7.3 (0.1%) | 6.5 (0.1%) |
| | East | 7.4 (0.1%) | 7.1 (0.1%) | 6 (0.1%) | 8 (0.2%) | 7.7 (0.2%) | 6.9 (0.1%) |
| | North-East | 7.2 (0.1%) | 7 (0.1%) | 5.8 (0.1%) | 7.8 (0.2%) | 7.6 (0.2%) | 6.7 (0.1%) |
| | South | 7.4 (0.1%) | 7.1 (0.1%) | 6 (0.1%) | 8 (0.2%) | 7.8 (0.2%) | 6.9 (0.1%) |
| Neanderthal | North | 68 (1.4%) | 65.5 (1.3%) | 56.8 (1.1%) | 72.7 (1.5%) | 70.6 (1.4%) | 63.8 (1.3%) |
| | Central | 69.8 (1.4%) | 67.2 (1.3%) | 58.3 (1.2%) | 74.6 (1.5%) | 72.6 (1.5%) | 65.5 (1.3%) |
| | Other | 69.7 (1.4%) | 66.9 (1.3%) | 57.8 (1.2%) | 74.8 (1.5%) | 72.6 (1.5%) | 65.3 (1.3%) |
| | West | 70.6 (1.4%) | 68 (1.4%) | 58.9 (1.2%) | 75.6 (1.5%) | 73.5 (1.5%) | 66.4 (1.3%) |
| | East | 73.1 (1.5%) | 70.3 (1.4%) | 61.1 (1.2%) | 78.2 (1.6%) | 76 (1.5%) | 68.7 (1.4%) |
| | North-East | 75.6 (1.5%) | 72.7 (1.5%) | 63.1 (1.3%) | 80.9 (1.6%) | 78.6 (1.6%) | 71 (1.4%) |
| | South | 70.6 (1.4%) | 68 (1.4%) | 59.1 (1.2%) | 76 (1.5%) | 73.9 (1.5%) | 66.8 (1.3%) |
| Both | North | 1.3 (<0.1%) | 1.1 (<0.1%) | 0.7 (<0.1%) | 1.5 (<0.1%) | 1.4 (<0.1%) | 1 (<0.1%) |
| | Central | 1.3 (<0.1%) | 1.2 (<0.1%) | 0.8 (<0.1%) | 1.6 (<0.1%) | 1.4 (<0.1%) | 1 (<0.1%) |
| | Other | 1.3 (<0.1%) | 1.1 (<0.1%) | 0.7 (<0.1%) | 1.5 (<0.1%) | 1.4 (<0.1%) | 1 (<0.1%) |
| | West | 1.4 (<0.1%) | 1.2 (<0.1%) | 0.8 (<0.1%) | 1.6 (<0.1%) | 1.5 (<0.1%) | 1 (<0.1%) |
| | East | 1.4 (<0.1%) | 1.2 (<0.1%) | 0.8 (<0.1%) | 1.7 (<0.1%) | 1.5 (<0.1%) | 1.1 (<0.1%) |
| | North-East | 1.4 (<0.1%) | 1.2 (<0.1%) | 0.8 (<0.1%) | 1.6 (<0.1%) | 1.4 (<0.1%) | 1 (<0.1%) |
| | South | 1.3 (<0.1%) | 1.2 (<0.1%) | 0.7 (<0.1%) | 1.6 (<0.1%) | 1.5 (<0.1%) | 1 (<0.1%) |
| none | North | 22.8 (0.5%) | 14.5 (0.3%) | 6.7 (0.1%) | 29.9 (0.6%) | 20.6 (0.4%) | 10.9 (0.2%) |
| | Central | 23 (0.5%) | 14.6 (0.3%) | 6.7 (0.1%) | 30.2 (0.6%) | 20.8 (0.4%) | 11 (0.2%) |
| | Other | 23.2 (0.5%) | 14.7 (0.3%) | 6.9 (0.1%) | 30.5 (0.6%) | 21 (0.4%) | 11.1 (0.2%) |
| | West | 23.1 (0.5%) | 14.6 (0.3%) | 6.7 (0.1%) | 30.3 (0.6%) | 20.9 (0.4%) | 11 (0.2%) |
| | East | 23.3 (0.5%) | 14.7 (0.3%) | 6.8 (0.1%) | 30.7 (0.6%) | 21.1 (0.4%) | 11.1 (0.2%) |
| | North-East | 23.1 (0.5%) | 14.7 (0.3%) | 6.9 (0.1%) | 30.8 (0.6%) | 21.2 (0.4%) | 11.2 (0.2%) |
| | South | 23 (0.5%) | 14.6 (0.3%) | 6.7 (0.1%) | 30.5 (0.6%) | 21.1 (0.4%) | 11.2 (0.2%) |
| All | North | 97.5 (2.0%) | 86.2 (1.7%) | 68.5 (1.4%) | 110 (2.2%) | 98.3 (2.0%) | 80.6 (1.6%) |
| | Central | 100 (2.0%) | 89 (1.8%) | 70.9 (1.4%) | 113 (2.3%) | 102 (2.0%) | 83.5 (1.7%) |
| | Other | 101 (2.0%) | 89.1 (1.8%) | 70.8 (1.4%) | 114 (2.3%) | 102 (2.0%) | 83.4 (1.7%) |
| | West | 102 (2.0%) | 90.5 (1.8%) | 72.1 (1.4%) | 115 (2.3%) | 103 (2.1%) | 85 (1.7%) |
| | East | 105 (2.1%) | 93.4 (1.9%) | 74.7 (1.5%) | 118 (2.4%) | 106 (2.1%) | 87.8 (1.8%) |
| | North-East | 107 (2.1%) | 95.5 (1.9%) | 76.6 (1.5%) | 121 (2.4%) | 109 (2.2%) | 89.9 (1.8%) |
| | South | 102 (2.0%) | 90.8 (1.8%) | 72.6 (1.5%) | 116 (2.3%) | 104 (2.1%) | 85.9 (1.7%) |

**Table S6.5. Average amount of archaic ancestry in MB and percent inferred using strategy 1 for HGDP and 1000G datasets** For phased and unphased data using posterior probability cutoffs at 0.5, 0.8 and 0.9. We assumed that the callable genome is 2 * 2,489,077,800 bp.

| Segment type | Region | Unphased | | | Phased | | |
|---|---|---|---|---|---|---|---|
| | | 0.5 | 0.8 | 0.9 | 0.5 | 0.8 | 0.9 |
| Denisova | Oceania | 91.8 (1.8%) | 90.1 (1.8%) | 82.6 (1.7%) | 106 (2.1%) | 104 (2.1%) | 94.5 (1.9%) |
| | East Asia | 5.9 (0.1%) | 5.7 (0.1%) | 4.8 (0.1%) | 6.3 (0.1%) | 6.1 (0.1%) | 5.4 (0.1%) |
| | America | 2.9 (0.1%) | 2.8 (0.1%) | 2.3 (<0.1%) | 3.1 (0.1%) | 3 (0.1%) | 2.6 (0.1%) |
| | Europe | 1.6 (<0.1%) | 1.4 (<0.1%) | 1.1 (<0.1%) | 1.7 (<0.1%) | 1.6 (<0.1%) | 1.3 (<0.1%) |
| | Middle East | 1.4 (<0.1%) | 1.3 (<0.1%) | 0.9 (<0.1%) | 1.5 (<0.1%) | 1.4 (<0.1%) | 1.1 (<0.1%) |
| | South Asia | 6.1 (0.1%) | 5.8 (0.1%) | 4.9 (0.1%) | 6.5 (0.1%) | 6.3 (0.1%) | 5.5 (0.1%) |
| Neanderthal | Oceania | 89.1 (1.8%) | 86.9 (1.7%) | 78.6 (1.6%) | 105 (2.1%) | 103 (2.1%) | 93.6 (1.9%) |
| | East Asia | 81.8 (1.6%) | 79 (1.6%) | 69.6 (1.4%) | 89.8 (1.8%) | 87.5 (1.8%) | 79.5 (1.6%) |
| | America | 63.7 (1.3%) | 61.6 (1.2%) | 54.4 (1.1%) | 69.1 (1.4%) | 67.3 (1.4%) | 61.3 (1.2%) |
| | Europe | 60 (1.2%) | 57.8 (1.2%) | 50.7 (1.0%) | 65 (1.3%) | 63.3 (1.3%) | 57.4 (1.2%) |
| | Middle East | 50.2 (1.0%) | 48.4 (1.0%) | 42.2 (0.8%) | 53.5 (1.1%) | 51.9 (1.0%) | 46.8 (0.9%) |
| | South Asia | 69 (1.4%) | 66.5 (1.3%) | 57.9 (1.2%) | 74.1 (1.5%) | 72 (1.4%) | 65.2 (1.3%) |
| Both | Oceania | 4.1 (0.1%) | 3.6 (0.1%) | 2.7 (0.1%) | 5.9 (0.1%) | 5.3 (0.1%) | 3.9 (0.1%) |
| | East Asia | 1.3 (<0.1%) | 1.1 (<0.1%) | 0.7 (<0.1%) | 1.5 (<0.1%) | 1.4 (<0.1%) | 1 (<0.1%) |
| | America | 0.9 (<0.1%) | 0.8 (<0.1%) | 0.5 (<0.1%) | 1.1 (<0.1%) | 1 (<0.1%) | 0.6 (<0.1%) |
| | Europe | 1 (<0.1%) | 0.9 (<0.1%) | 0.6 (<0.1%) | 1.2 (<0.1%) | 1.1 (<0.1%) | 0.8 (<0.1%) |
| | Middle East | 1 (<0.1%) | 0.9 (<0.1%) | 0.6 (<0.1%) | 1.1 (<0.1%) | 1 (<0.1%) | 0.7 (<0.1%) |
| | South Asia | 1.4 (<0.1%) | 1.2 (<0.1%) | 0.8 (<0.1%) | 1.6 (<0.1%) | 1.5 (<0.1%) | 1 (<0.1%) |
| none | Oceania | 21.8 (0.4%) | 14.1 (0.3%) | 7 (0.1%) | 36.3 (0.7%) | 24.1 (0.5%) | 12.4 (0.2%) |
| | East Asia | 23.3 (0.5%) | 15 (0.3%) | 7.2 (0.1%) | 31.4 (0.6%) | 21.9 (0.4%) | 11.9 (0.2%) |
| | America | 22.9 (0.5%) | 14.5 (0.3%) | 6.7 (0.1%) | 26.6 (0.5%) | 18.4 (0.4%) | 9.7 (0.2%) |
| | Europe | 22.1 (0.4%) | 14.1 (0.3%) | 6.6 (0.1%) | 30.2 (0.6%) | 20.9 (0.4%) | 11.2 (0.2%) |
| | Middle East | 26.6 (0.5%) | 16.5 (0.3%) | 7.4 (0.1%) | 30.2 (0.6%) | 20.6 (0.4%) | 10.9 (0.2%) |
| | South Asia | 23.4 (0.5%) | 14.9 (0.3%) | 6.8 (0.1%) | 31.2 (0.6%) | 21.5 (0.4%) | 11.4 (0.2%) |
| All | Oceania | 207 (4.2%) | 195 (3.9%) | 171 (3.4%) | 254 (5.1%) | 236 (4.7%) | 204 (4.1%) |
| | East Asia | 112 (2.2%) | 101 (2.0%) | 82.3 (1.7%) | 129 (2.6%) | 117 (2.4%) | 97.8 (2.0%) |
| | America | 90.4 (1.8%) | 79.6 (1.6%) | 64 (1.3%) | 99.9 (2.0%) | 89.6 (1.8%) | 74.3 (1.5%) |
| | Europe | 84.6 (1.7%) | 74.3 (1.5%) | 58.9 (1.2%) | 98.2 (2.0%) | 86.9 (1.7%) | 70.7 (1.4%) |
| | Middle East | 79.2 (1.6%) | 67 (1.3%) | 51.2 (1.0%) | 86.3 (1.7%) | 74.8 (1.5%) | 59.5 (1.2%) |
| | South Asia | 99.8 (2.0%) | 88.3 (1.8%) | 70.4 (1.4%) | 113 (2.3%) | 101 (2.0%) | 83.2 (1.7%) |

**Table S6.6 Average amount of archaic ancestry inferred using approach 2 for LASI-DAD dataset.** For phased and unphased data using different posterior probability cutoffs. We assumed that the callable genome is 2 * 2,489,077,800 bp.

| Segment type | Region | Unphased | | | Phased | | |
|---|---|---|---|---|---|---|---|
| | | 0.5 | 0.8 | 0.9 | 0.5 | 0.8 | 0.9 |
| ND01 | North | 4 (0.1%) | 3.8 (0.1%) | 3 (0.1%) | 4.4 (0.1%) | 4.3 (0.1%) | 3.6 (0.1%) |
| | Central | 4.6 (0.1%) | 4.4 (0.1%) | 3.5 (0.1%) | 5.2 (0.1%) | 5 (0.1%) | 4.2 (0.1%) |
| | Other | 4.9 (0.1%) | 4.6 (0.1%) | 3.8 (0.1%) | 5.4 (0.1%) | 5.2 (0.1%) | 4.4 (0.1%) |
| | West | 5.1 (0.1%) | 4.8 (0.1%) | 3.9 (0.1%) | 5.6 (0.1%) | 5.4 (0.1%) | 4.7 (0.1%) |
| | East | 5.4 (0.1%) | 5.2 (0.1%) | 4.2 (0.1%) | 6 (0.1%) | 5.8 (0.1%) | 5 (0.1%) |
| | North-East | 5.4 (0.1%) | 5.1 (0.1%) | 4.1 (0.1%) | 6.1 (0.1%) | 5.8 (0.1%) | 5 (0.1%) |
| | South | 5.3 (0.1%) | 5.1 (0.1%) | 4.1 (0.1%) | 6 (0.1%) | 5.7 (0.1%) | 5 (0.1%) |
| ND10 | North | 65.9 (1.3%) | 63.3 (1.3%) | 54.8 (1.1%) | 71.2 (1.4%) | 69.1 (1.4%) | 62.3 (1.3%) |
| | Central | 67.4 (1.4%) | 64.8 (1.3%) | 56 (1.1%) | 72.9 (1.5%) | 70.9 (1.4%) | 63.9 (1.3%) |
| | Other | 67.3 (1.4%) | 64.6 (1.3%) | 55.6 (1.1%) | 73 (1.5%) | 70.9 (1.4%) | 63.7 (1.3%) |
| | West | 68.1 (1.4%) | 65.4 (1.3%) | 56.6 (1.1%) | 73.8 (1.5%) | 71.7 (1.4%) | 64.7 (1.3%) |
| | East | 70.2 (1.4%) | 67.5 (1.4%) | 58.5 (1.2%) | 76.2 (1.5%) | 74 (1.5%) | 66.8 (1.3%) |
| | North-East | 73 (1.5%) | 70.2 (1.4%) | 60.8 (1.2%) | 79.1 (1.6%) | 76.8 (1.5%) | 69.3 (1.4%) |
| | South | 67.9 (1.4%) | 65.3 (1.3%) | 56.6 (1.1%) | 74.1 (1.5%) | 72 (1.4%) | 64.9 (1.3%) |
| ND11 | North | 1.1 (<0.1%) | 0.9 (<0.1%) | 0.5 (<0.1%) | 1.3 (<0.1%) | 1.2 (<0.1%) | 0.8 (<0.1%) |
| | Central | 1.1 (<0.1%) | 1 (<0.1%) | 0.6 (<0.1%) | 1.4 (<0.1%) | 1.2 (<0.1%) | 0.8 (<0.1%) |
| | Other | 1.1 (<0.1%) | 1 (<0.1%) | 0.6 (<0.1%) | 1.3 (<0.1%) | 1.2 (<0.1%) | 0.8 (<0.1%) |
| | West | 1.1 (<0.1%) | 1 (<0.1%) | 0.6 (<0.1%) | 1.4 (<0.1%) | 1.2 (<0.1%) | 0.8 (<0.1%) |
| | East | 1.2 (<0.1%) | 1 (<0.1%) | 0.6 (<0.1%) | 1.4 (<0.1%) | 1.3 (<0.1%) | 0.9 (<0.1%) |
| | North-East | 1.1 (<0.1%) | 1 (<0.1%) | 0.6 (<0.1%) | 1.4 (<0.1%) | 1.2 (<0.1%) | 0.8 (<0.1%) |
| | South | 1.1 (<0.1%) | 1 (<0.1%) | 0.6 (<0.1%) | 1.4 (<0.1%) | 1.2 (<0.1%) | 0.8 (<0.1%) |
| ND11 mosaic | North | 3.8 (0.1%) | 3.7 (0.1%) | 3.5 (0.1%) | 3.1 (0.1%) | 3.1 (0.1%) | 3 (0.1%) |
| | Central | 4.4 (0.1%) | 4.3 (0.1%) | 4 (0.1%) | 3.7 (0.1%) | 3.6 (0.1%) | 3.5 (0.1%) |
| | Other | 4.3 (0.1%) | 4.2 (0.1%) | 3.9 (0.1%) | 3.6 (0.1%) | 3.6 (0.1%) | 3.4 (0.1%) |
| | West | 4.7 (0.1%) | 4.7 (0.1%) | 4.3 (0.1%) | 4 (0.1%) | 3.9 (0.1%) | 3.8 (0.1%) |
| | East | 5 (0.1%) | 5 (0.1%) | 4.6 (0.1%) | 4.2 (0.1%) | 4.2 (0.1%) | 4 (0.1%) |
| | North-East | 4.6 (0.1%) | 4.6 (0.1%) | 4.3 (0.1%) | 3.8 (0.1%) | 3.7 (0.1%) | 3.6 (0.1%) |
| | South | 5 (0.1%) | 5 (0.1%) | 4.6 (0.1%) | 4.2 (0.1%) | 4.2 (0.1%) | 4 (0.1%) |
| ND00 | North | 22.8 (0.5%) | 14.5 (0.3%) | 6.7 (0.1%) | 29.9 (0.6%) | 20.6 (0.4%) | 10.9 (0.2%) |
| | Central | 23 (0.5%) | 14.6 (0.3%) | 6.7 (0.1%) | 30.2 (0.6%) | 20.8 (0.4%) | 11 (0.2%) |
| | Other | 23.2 (0.5%) | 14.7 (0.3%) | 6.9 (0.1%) | 30.5 (0.6%) | 21 (0.4%) | 11.1 (0.2%) |
| | West | 23.1 (0.5%) | 14.6 (0.3%) | 6.7 (0.1%) | 30.3 (0.6%) | 20.9 (0.4%) | 11 (0.2%) |
| | East | 23.3 (0.5%) | 14.7 (0.3%) | 6.8 (0.1%) | 30.7 (0.6%) | 21.1 (0.4%) | 11.1 (0.2%) |
| | North-East | 23.1 (0.5%) | 14.7 (0.3%) | 6.9 (0.1%) | 30.8 (0.6%) | 21.2 (0.4%) | 11.2 (0.2%) |
| | South | 23 (0.5%) | 14.6 (0.3%) | 6.7 (0.1%) | 30.5 (0.6%) | 21.1 (0.4%) | 11.2 (0.2%) |
| All | North | 97.5 (2.0%) | 86.2 (1.7%) | 68.5 (1.4%) | 110 (2.2%) | 98.3 (2.0%) | 80.6 (1.6%) |
| | Central | 100 (2.0%) | 89 (1.8%) | 70.9 (1.4%) | 113 (2.3%) | 102 (2.0%) | 83.5 (1.7%) |
| | Other | 101 (2.0%) | 89.1 (1.8%) | 70.8 (1.4%) | 114 (2.3%) | 102 (2.0%) | 83.4 (1.7%) |
| | West | 102 (2.0%) | 90.5 (1.8%) | 72.1 (1.4%) | 115 (2.3%) | 103 (2.1%) | 85 (1.7%) |
| | East | 105 (2.1%) | 93.4 (1.9%) | 74.7 (1.5%) | 118 (2.4%) | 106 (2.1%) | 87.8 (1.8%) |
| | North-East | 107 (2.1%) | 95.5 (1.9%) | 76.6 (1.5%) | 121 (2.4%) | 109 (2.2%) | 89.9 (1.8%) |
| | South | 102 (2.0%) | 90.8 (1.8%) | 72.6 (1.5%) | 116 (2.3%) | 104 (2.1%) | 85.9 (1.7%) |

**Table S6.7 Average amount of archaic ancestry inferred using approach 2 for HGDP and 1000G datasets** For phased and unphased data using different posterior probability cutoffs. We assumed that the callable genome is 2 * 2,489,077,800 bp

| Segment type | Region | Unphased | | | Phased | | |
|---|---|---|---|---|---|---|---|
| | | 0.5 | 0.8 | 0.9 | 0.5 | 0.8 | 0.9 |
| ND01 | Oceania | 59.4 (1.2%) | 57.7 (1.2%) | 51.3 (1.0%) | 77.7 (1.6%) | 75.5 (1.5%) | 67 (1.3%) |
| | East Asia | 4.7 (0.1%) | 4.4 (0.1%) | 3.6 (0.1%) | 5.3 (0.1%) | 5 (0.1%) | 4.4 (0.1%) |
| | America | 2.3 (<0.1%) | 2.1 (<0.1%) | 1.7 (<0.1%) | 2.5 (0.1%) | 2.4 (<0.1%) | 2 (<0.1%) |
| | Europe | 1.3 (<0.1%) | 1.2 (<0.1%) | 0.8 (<0.1%) | 1.5 (<0.1%) | 1.4 (<0.1%) | 1 (<0.1%) |
| | Middle East | 1.2 (<0.1%) | 1.1 (<0.1%) | 0.8 (<0.1%) | 1.3 (<0.1%) | 1.2 (<0.1%) | 0.9 (<0.1%) |
| | South Asia | 4.4 (0.1%) | 4.1 (0.1%) | 3.4 (0.1%) | 4.9 (0.1%) | 4.7 (0.1%) | 4 (0.1%) |
| ND10 | Oceania | 72.6 (1.5%) | 70.5 (1.4%) | 62.9 (1.3%) | 93.1 (1.9%) | 90.6 (1.8%) | 82 (1.6%) |
| | East Asia | 79.2 (1.6%) | 76.4 (1.5%) | 67.2 (1.3%) | 88 (1.8%) | 85.7 (1.7%) | 77.8 (1.6%) |
| | America | 62.1 (1.2%) | 60 (1.2%) | 52.9 (1.1%) | 67.9 (1.4%) | 66.2 (1.3%) | 60.2 (1.2%) |
| | Europe | 58.7 (1.2%) | 56.5 (1.1%) | 49.4 (1.0%) | 64 (1.3%) | 62.3 (1.3%) | 56.5 (1.1%) |
| | Middle East | 49 (1.0%) | 47.2 (0.9%) | 41.1 (0.8%) | 52.5 (1.1%) | 50.9 (1.0%) | 45.9 (0.9%) |
| | South Asia | 66.6 (1.3%) | 64.1 (1.3%) | 55.7 (1.1%) | 72.4 (1.5%) | 70.3 (1.4%) | 63.6 (1.3%) |
| ND11 | Oceania | 2.6 (0.1%) | 2.2 (<0.1%) | 1.5 (<0.1%) | 4.1 (0.1%) | 3.6 (0.1%) | 2.4 (<0.1%) |
| | East Asia | 1.1 (<0.1%) | 0.9 (<0.1%) | 0.5 (<0.1%) | 1.3 (<0.1%) | 1.2 (<0.1%) | 0.8 (<0.1%) |
| | America | 0.8 (<0.1%) | 0.7 (<0.1%) | 0.4 (<0.1%) | 0.9 (<0.1%) | 0.8 (<0.1%) | 0.5 (<0.1%) |
| | Europe | 0.8 (<0.1%) | 0.7 (<0.1%) | 0.4 (<0.1%) | 1 (<0.1%) | 0.9 (<0.1%) | 0.6 (<0.1%) |
| | Middle East | 0.8 (<0.1%) | 0.7 (<0.1%) | 0.4 (<0.1%) | 1 (<0.1%) | 0.9 (<0.1%) | 0.6 (<0.1%) |
| | South Asia | 1.1 (<0.1%) | 1 (<0.1%) | 0.6 (<0.1%) | 1.4 (<0.1%) | 1.3 (<0.1%) | 0.8 (<0.1%) |
| ND11 mosaic | Oceania | 50.4 (1.0%) | 50.2 (1.0%) | 48.3 (1.0%) | 42.4 (0.9%) | 42.2 (0.8%) | 40.6 (0.8%) |
| | East Asia | 4 (0.1%) | 3.9 (0.1%) | 3.7 (0.1%) | 3.1 (0.1%) | 3.1 (0.1%) | 2.9 (0.1%) |
| | America | 2.3 (<0.1%) | 2.3 (<0.1%) | 2.2 (<0.1%) | 1.9 (<0.1%) | 1.9 (<0.1%) | 1.9 (<0.1%) |
| | Europe | 1.8 (<0.1%) | 1.7 (<0.1%) | 1.6 (<0.1%) | 1.5 (<0.1%) | 1.5 (<0.1%) | 1.4 (<0.1%) |
| | Middle East | 1.6 (<0.1%) | 1.5 (<0.1%) | 1.4 (<0.1%) | 1.3 (<0.1%) | 1.3 (<0.1%) | 1.3 (<0.1%) |
| | South Asia | 4.2 (0.1%) | 4.2 (0.1%) | 3.9 (0.1%) | 3.5 (0.1%) | 3.4 (0.1%) | 3.3 (0.1%) |
| ND00 | Oceania | 21.8 (0.4%) | 14.1 (0.3%) | 7 (0.1%) | 36.3 (0.7%) | 24.1 (0.5%) | 12.4 (0.2%) |
| | East Asia | 23.3 (0.5%) | 15.1 (0.3%) | 7.2 (0.1%) | 31.4 (0.6%) | 21.9 (0.4%) | 11.9 (0.2%) |
| | America | 22.9 (0.5%) | 14.5 (0.3%) | 6.8 (0.1%) | 26.6 (0.5%) | 18.4 (0.4%) | 9.7 (0.2%) |
| | Europe | 22.1 (0.4%) | 14.1 (0.3%) | 6.7 (0.1%) | 30.2 (0.6%) | 20.9 (0.4%) | 11.2 (0.2%) |
| | Middle East | 26.6 (0.5%) | 16.5 (0.3%) | 7.5 (0.2%) | 30.2 (0.6%) | 20.6 (0.4%) | 10.9 (0.2%) |
| | South Asia | 23.5 (0.5%) | 14.9 (0.3%) | 6.9 (0.1%) | 31.2 (0.6%) | 21.5 (0.4%) | 11.4 (0.2%) |
| All | Oceania | 207 (4.2%) | 195 (3.9%) | 171 (3.4%) | 254 (5.1%) | 236 (4.7%) | 204 (4.1%) |
| | East Asia | 112 (2.2%) | 101 (2.0%) | 82.3 (1.7%) | 129 (2.6%) | 117 (2.4%) | 97.8 (2.0%) |
| | America | 90.4 (1.8%) | 79.6 (1.6%) | 64 (1.3%) | 99.9 (2.0%) | 89.6 (1.8%) | 74.3 (1.5%) |
| | Europe | 84.6 (1.7%) | 74.3 (1.5%) | 58.9 (1.2%) | 98.2 (2.0%) | 86.9 (1.7%) | 70.7 (1.4%) |
| | Middle East | 79.2 (1.6%) | 67 (1.3%) | 51.2 (1.0%) | 86.3 (1.7%) | 74.8 (1.5%) | 59.5 (1.2%) |
| | South Asia | 99.8 (2.0%) | 88.3 (1.8%) | 70.4 (1.4%) | 113 (2.3%) | 101 (2.0%) | 83.2 (1.7%) |

**No evidence for introgression from unknown archaic (ghost) hominin group in India**

To assess the evidence for ghost archaic introgression in India, we compared the amount of archaic sequence found in individuals from India to what is found in other worldwide populations from 1000G, HGDP and deCODE Genetics using phased data (Figure S6.3). We expect these regions will not share any derived variants with sequenced archaic genomes of Neanderthals and Denisovans (i.e., classified as ‘none’ in Tables S6.4-S6.5). We find similar amounts of archaic ancestry that does not share any alleles with Denisovans and Neanderthals in India compared to other worldwide populations, suggesting that there is no strong evidence for additional unknown archaic introgression in India.

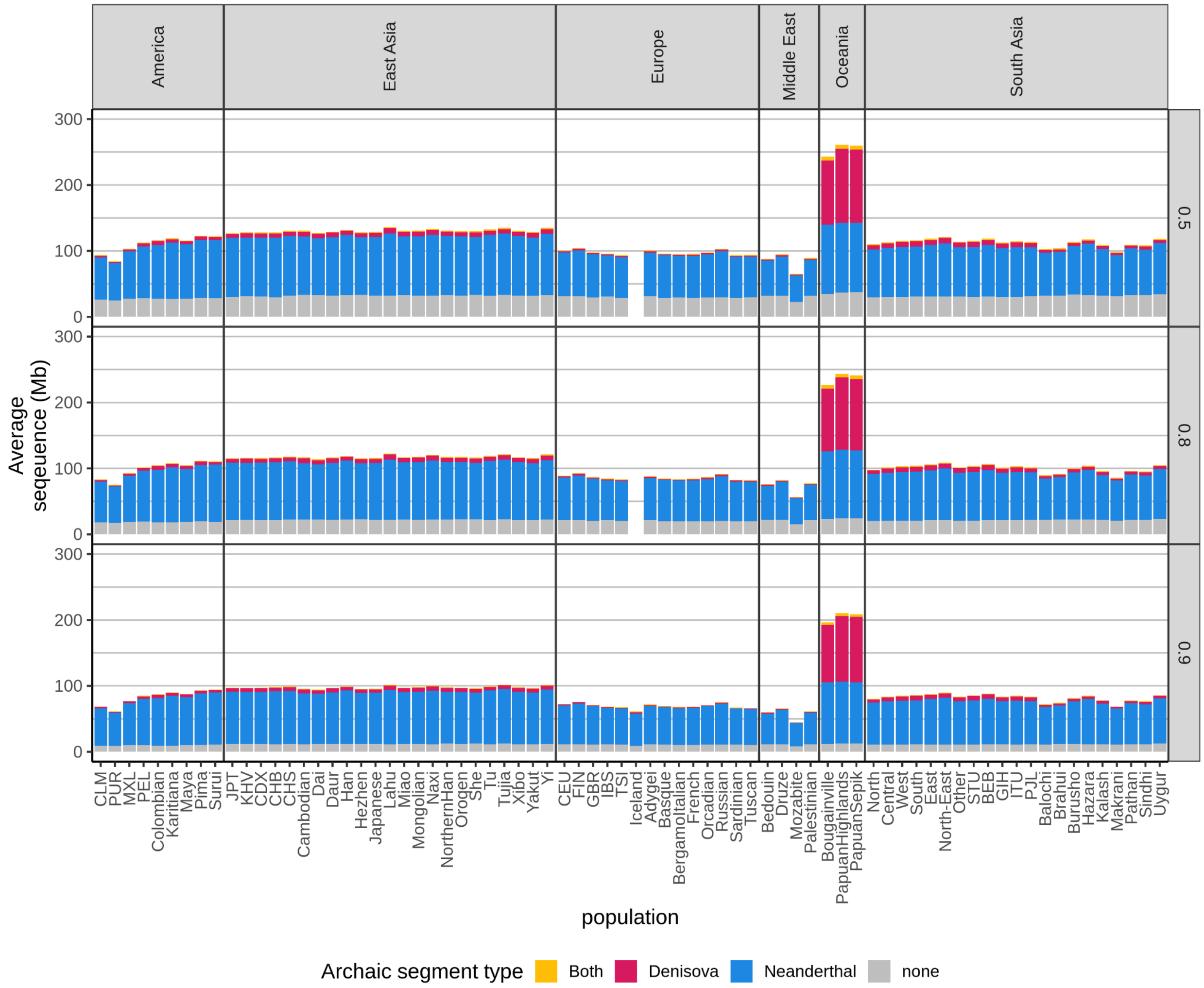

**Figure S6.3. Average of archaic sequence per population.** We show the average amount of archaic sequence at posterior cutoffs of 0.5, 0.8 and 0.9 for each population in Mb. Note data from Icelanders from sequenced at deCODE are only available at a posterior probability cutoff at 0.9.

#### Fossil-free reconstruction of introgressed archaic genomes

Due to recombination, archaic ancestry segments may be present in different genomic regions across individual genomes in a population. We assembled non-overlapping archaic ancestry segments across individuals in a population and compared the cumulative genome length of the introgressing Neanderthal and Denisovan genomes recovered from each population in each dataset (Extended Data Table S2). Specifically, we counted the amount of high-quality archaic sequence (with posterior probability ≥ 0.9 in order to compare to data from deCODE) in modern human genomes classified as Neanderthal or Denisovan ancestry for each population in four datasets (1000G, deCODE, HGDP and LASI-DAD), comparable to previous analysis in Icelanders from deCODE Genetics[43] shown in Figure S6.4.

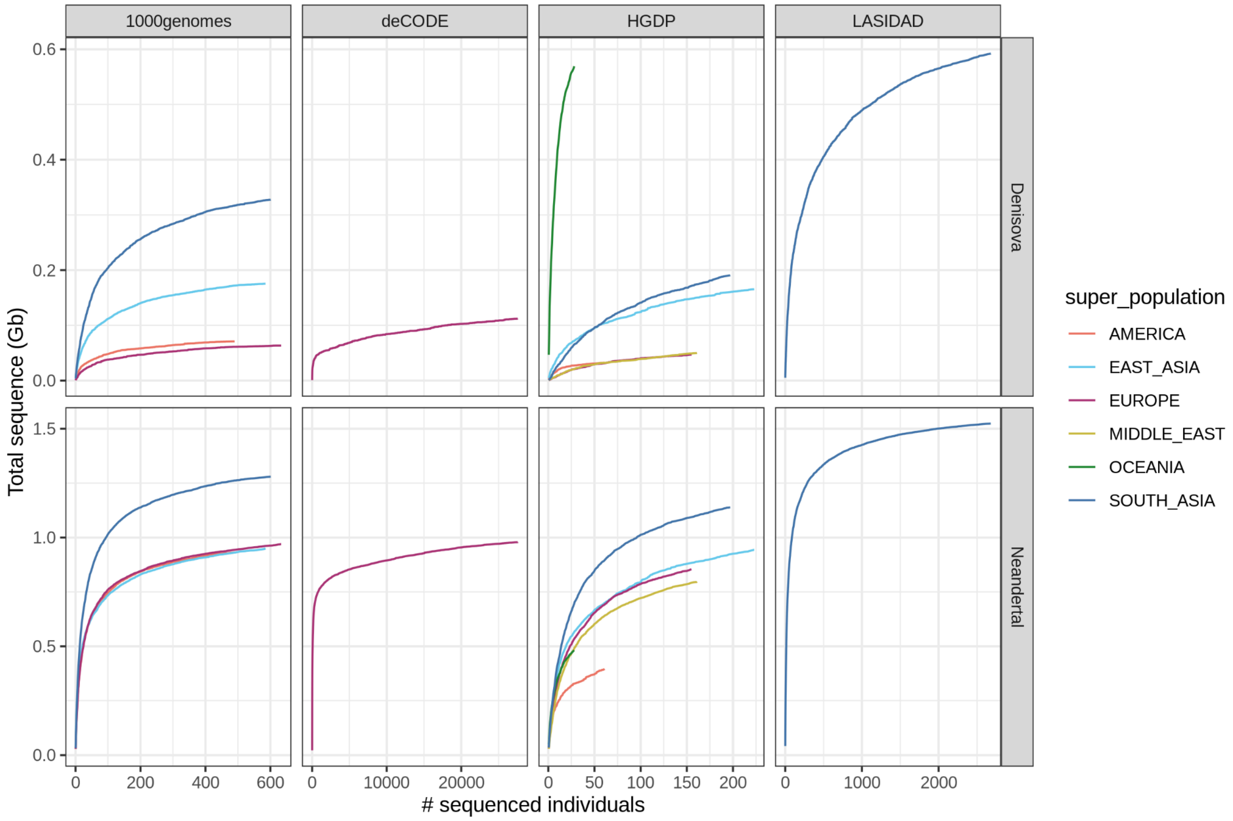

**Figure S6.4. Amount of unique archaic sequence in worldwide populations.** For Denisovan (top) and Neanderthal (bottom) as a function of the analyzed number of individuals in four different datasets (at a posterior cutoff of 0.9).

Excluded individuals from deCODE genetics and using a posterior probability cutoff of 0.8 (the cutoff used throughout this paper) allows us to reconstruct 1,679 Mb of introgressing Neanderthal sequence and 1,080 Mb of introgressing Denisovan sequence (Table S.6.8).

**Table S6.8. Total amount of archaic sequence found in the world.** We show the total amount of archaic sequence that could be constructed for all regions in HGDP, 1000G and LASI-DAD datasets using a posterior probability cutoff of 0.8 in the "Total" column. We then group the individuals by region and downsample to either 28 individuals, 490 individuals or the full dataset and count the length of the reconstructed introgression archaic genome for Neanderthal segments (NEA), ND10 segments (ND10), Denisovan segments (DEN) and ND01 segments (ND01). All values are in megabases (Mb) and the numbers in parentheses denote the 2.5% percentile and 97.5 percentile for 100 bootstrap replicates. No bootstrap resampling was done for the full dataset.

| **Downsample to 28 individuals for all regions** | | | | | | | | |
|---|---|---|---|---|---|---|---|---|
| Segment type | Total | LASI-DAD | South Asia | East Asia | Europe | America | Middle East | Oceania |
| NEA | 1679 | 711 (690-729) | 701 (678-722) | 551 (531-567) | 529 (517-543) | 520 (488-543) | 468 (436-489) | 421 (394-447) |
| ND10 | 1651 | 699 (677-717) | 689 (667-710) | 543 (524-560) | 523 (510-537) | 515 (483-537) | 463 (430-484) | 385 (360-410) |
| DEN | 1080 | 109 (102-118) | 95 (84-103) | 67 (62-73) | 19 (16-22) | 29 (26-32) | 18 (16-21) | 497 (468-525) |
| ND01 | 947 | 84 (78-91) | 74 (66-81) | 56 (51-60) | 16 (14-18) | 24 (21-27) | 16 (14-18) | 409 (380-436) |
| **Downsample to 490 to individuals for all regions - Middle East and Oceania excluded due to low sample size** | | | | | | | | |
| Segment type | Total | LASI-DAD | South Asia | East Asia | Europe | America | Middle East | Oceania |
| NEA | 1679 | 1314 (1304-1325) | 1262 (1253-1269) | 962 (949-979) | 941 (932-951) | 893 (882-901) | - | - |
| ND10 | 1651 | 1298 (1288-1306) | 1246 (1237-1253) | 951 (938-967) | 934 (925-942) | 884 (873-893) | - | - |
| DEN | 1080 | 384 (373-394) | 304 (297-310) | 176 (170-182) | 58 (55-60) | 65 (63-67) | - | - |
| ND01 | 947 | 317 (308-325) | 256 (251-261) | 154 (148-159) | 50 (48-52) | 55 (54-57) | - | - |
| **Full dataset. Number of individuals sampled: LASI-DAD (2679), South Asia (601 + 197), East Asia (585 + 223), Europe (633 + 155), America (490 + 61), Middle East (161) and Oceania (28)** | | | | | | | | |
| Segment type | Total | LASI-DAD | South Asia | East Asia | Europe | America | Middle East | Oceania |
| NEA | 1679 | 1524 | 1354 | 1078 | 1027 | 938 | 797 | 484 |
| ND10 | 1651 | 1504 | 1337 | 1066 | 1020 | 930 | 788 | 444 |
| DEN | 1080 | 591 | 360 | 218 | 74 | 73 | 50 | 568 |
| ND01 | 947 | 511 | 311 | 193 | 65 | 62 | 43 | 476 |

**Sharing of archaic ancestry across worldwide populations**

We calculated the amount of archaic sequence that is shared between all possible combinations of worldwide populations at a posterior probability threshold of 0.8. We find that the highest fraction of Neanderthal ancestry segments are shared across worldwide populations. We observe ~195.9 Mb of the 1,524 Mb Neanderthal sequence (12.8%) is unique to individuals from India, while 301.6 Mb of the 591 Mb of Denisovan sequence (51.0%) is unique to individuals from India. The large proportion of unique Denisovan sequences in South Asian datasets has previously been reported for 1000G populations[48]. We show the sharing of Neanderthal segments as an upset plot in Figure S6.5 and of Denisovan segments in Figure S6.6.

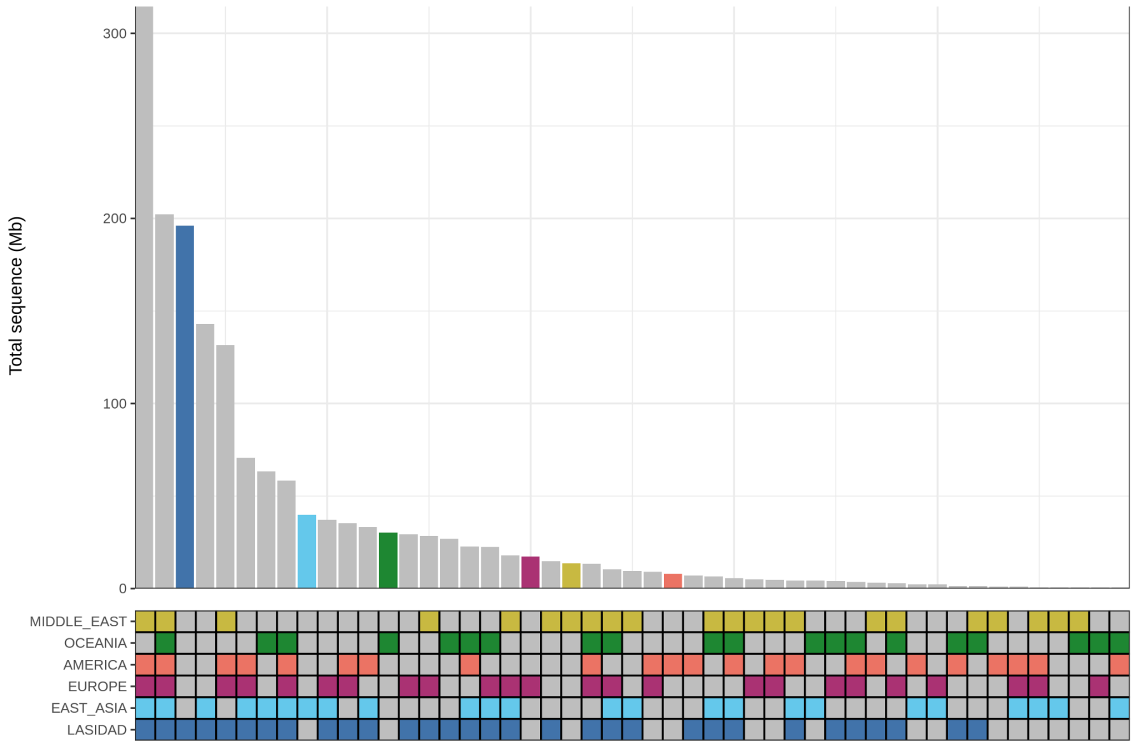

**Figure S6.5. Amount of unique and shared Neanderthal sequence.** The amount of Neanderthal sequence (y-axis) that is shared between any combinations of population (x-axis). Sequence that is unique to one population is colored according to which population it is found in while sequence that is shared is colored in grey.

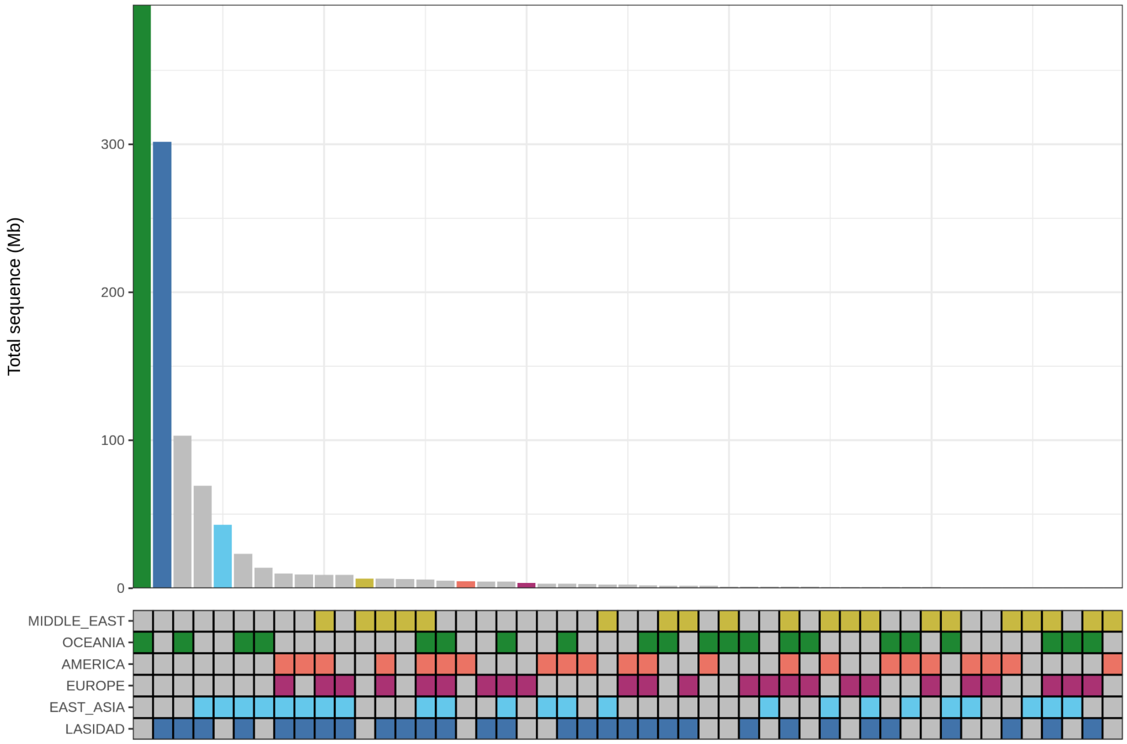

**Figure S6.6. Amount of unique and shared Denisovan sequence.** The amount of Denisovan sequence (y-axis) that is shared between any combinations of population (x-axis). Sequence that is unique to one population is colored according to which population it is found in while sequence that is shared is colored in grey.

The larger amount of unique Denisovan and Neanderthal sequences in India might be due to the larger sample size of 2,679 individuals in LASI-DAD, compared to other populations groups. In order to correct for this effect, we down-sampled all populations in the 1000G project to 490 individuals and compared the results with 490 randomly selected Indians from LASI-DAD. We also down-sampled all HGDP populations to 28 individuals and compared them to 28 randomly selected individuals from LASI-DAD. We show the results for Neanderthal sequence in Figure S6.7 and for Denisovan sequence in Figure S6.8.

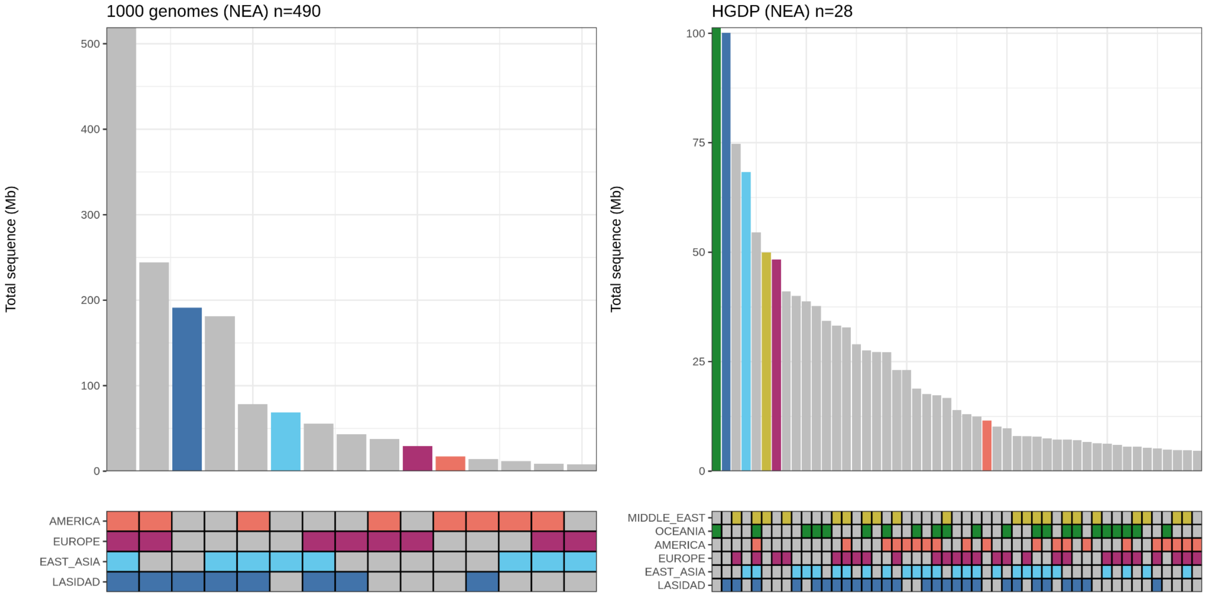

**Figure S6.7. Amount of unique and shared Neanderthal sequence in subsampled populations.** The amount of Neanderthal sequence (y-axis) that is shared between any combinations of population (x-axis). Sequence that is unique to one population is colored according to which population it is found in while sequence that is shared is colored in grey. All populations in 1000G have been down-sampled to 490 individuals and all populations in HGDP have been down-sampled to 28 individuals.

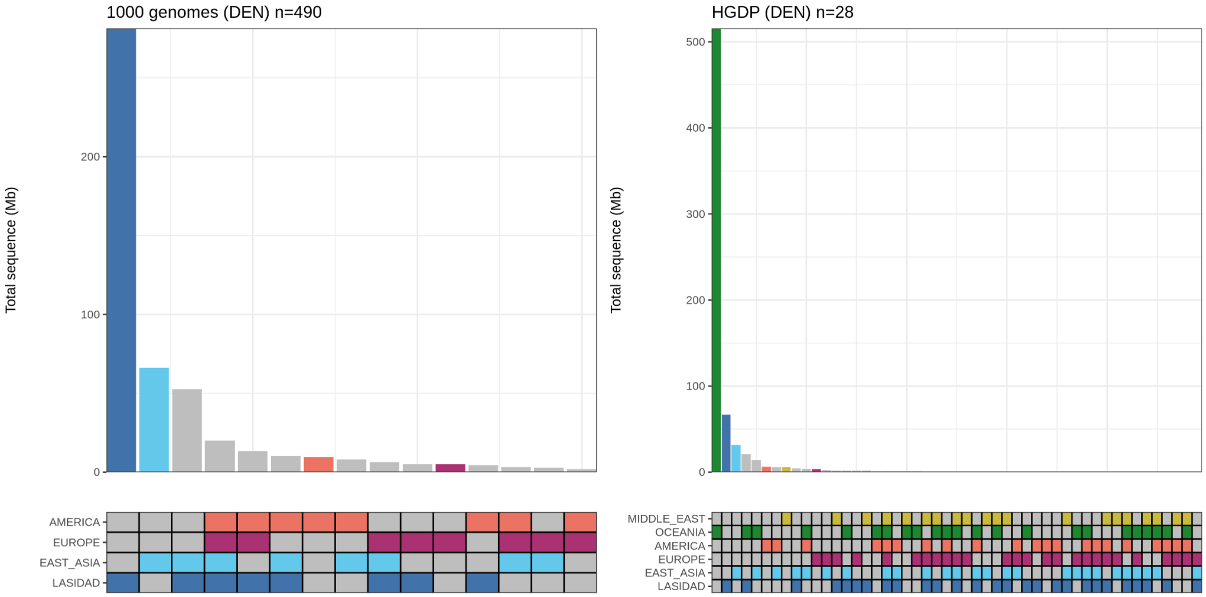

**Figure S6.8. Amount of unique and shared Denisovan sequence in subsampled populations.** The amount of Denisovan sequence (y-axis) that is shared between any combinations of population (x-axis). Sequence that is unique to one population is colored according to which population it is found in while sequence that is shared is colored in grey. All populations in 1000G have been down-sampled to 490 individuals and all populations in HGDP have been down-sampled to 28 individuals.

**Impact of recent demographic history on shaping patterns of archaic ancestry in India**

We examined the variation in archaic ancestry and its association with recent ancestry components for individuals on the Indian cline where the p-value for the *qpAdm* was greater than 0.01 and all ancestry components were positive (n = 1,936). Specifically, we investigated how the proportion of archaic ancestry varies by AHG-related, Iranian-farmer-related and Steppe pastoralist-related ancestry in India (Supplementary Note S4). We found the strongest association between Denisovan ancestry and Andamanese Hunter-gatherer-related ancestry (AHG) (Table S6.9). As for AHG ancestry, correlations with both Denisovan and Neanderthal are significant even when a stringent criteria is used (ND01 or ND10). Correlations with both Denisovan and Neanderthal are significant even when a stringent criteria is used (ND01 or ND10).

**Table S6.9. Correlation between proportion of archaic ancestry and Iranian farmer-related (Sarazm_EN), Steppe pastoralist-related (Central_Steppe_MLBA) and AHG-related (Onge) ancestry in India**. The intercept can be interpreted as the amount of archaic ancestry (in Mb) that each individual would have on average if they had 0% Steppe pastoralist-related, Iranian farmer-related or AHG-related ancestry. The slope can be interpreted as the amount of archaic ancestry (in Mb) for every unit increase in ancestry component coefficient. Both *p*-values and spearman correlation between the fitted model and data are shown.

| Archaic ancestry | Ancestry component | Intercept | Slope | p-value | Correlation |
|---|---|---|---|---|---|
| Denisova | *Sarazm_EN* | 11.44 | -10.31 | 3.61E-57 | 0.123 |
| | Steppe pastoralist | 8.40 | -12.97 | 1.26E-183 | 0.35 |
| | AHG | 1.46 | 12.45 | 3.32E-281 | 0.485 |
| Neanderthal | *Sarazm_EN* | 83.26 | -24.01 | 4.75E-41 | 0.088 |
| | Steppe pastoralist | 75.22 | -22.27 | 2.75E-64 | 0.137 |
| | AHG | 62.29 | 23.70 | 2.27E-114 | 0.234 |
| Both | *Sarazm_EN* | 1.82 | -0.88 | 1.38E-07 | 0.014 |
| | Steppe pastoralist | 1.54 | -0.97 | 5.85E-15 | 0.031 |
| | AHG | 1.01 | 0.97 | 6.04E-22 | 0.046 |
| none | *Sarazm_EN* | 21.64 | -1.83 | 6.43E-04 | 0.005 |
| | Steppe pastoralist | 21.24 | -3.50 | 1.03E-18 | 0.039 |
| | AHG | 19.52 | 3.01 | 1.18E-20 | 0.043 |
| ND01 | *Sarazm_EN* | 8.54 | -7.74 | 2.69E-54 | 0.117 |
| | Steppe pastoralist | 6.20 | -9.29 | 3.47E-154 | 0.303 |
| | AHG | 1.17 | 9.05 | 2.19E-240 | 0.432 |

| | | | | | |
|---|---|---|---|---|---|
| ND10 | *Sarazm_EN* | 80.64 | -22.11 | 3.89E-37 | 0.08 |
| | Steppe pastoralist | 73.16 | -19.84 | 4.96E-54 | 0.116 |
| | AHG | 61.52 | 21.38 | 1.25E-97 | 0.203 |
| ND11 | *Sarazm_EN* | 1.49 | -0.61 | 5.70E-05 | 0.008 |
| | Steppe pastoralist | 1.31 | -0.80 | 9.03E-13 | 0.026 |
| | AHG | 0.89 | 0.76 | 9.69E-17 | 0.035 |
| ND11_mosaic | *Sarazm_EN* | 5.85 | -4.74 | 1.13E-23 | 0.05 |
| | Steppe pastoralist | 4.49 | -6.29 | 1.44E-75 | 0.16 |
| | AHG | 1.17 | 5.94 | 6.61E-104 | 0.215 |
| ND00 | *Sarazm_EN* | 21.64 | -1.83 | 6.68E-04 | 0.005 |
| | Steppe pastoralist | 21.24 | -3.50 | 1.13E-18 | 0.039 |
| | AHG | 19.52 | 3.01 | 1.36E-20 | 0.043 |

### Inferring the number of pulses of Neanderthal and Denisovan gene flow in India

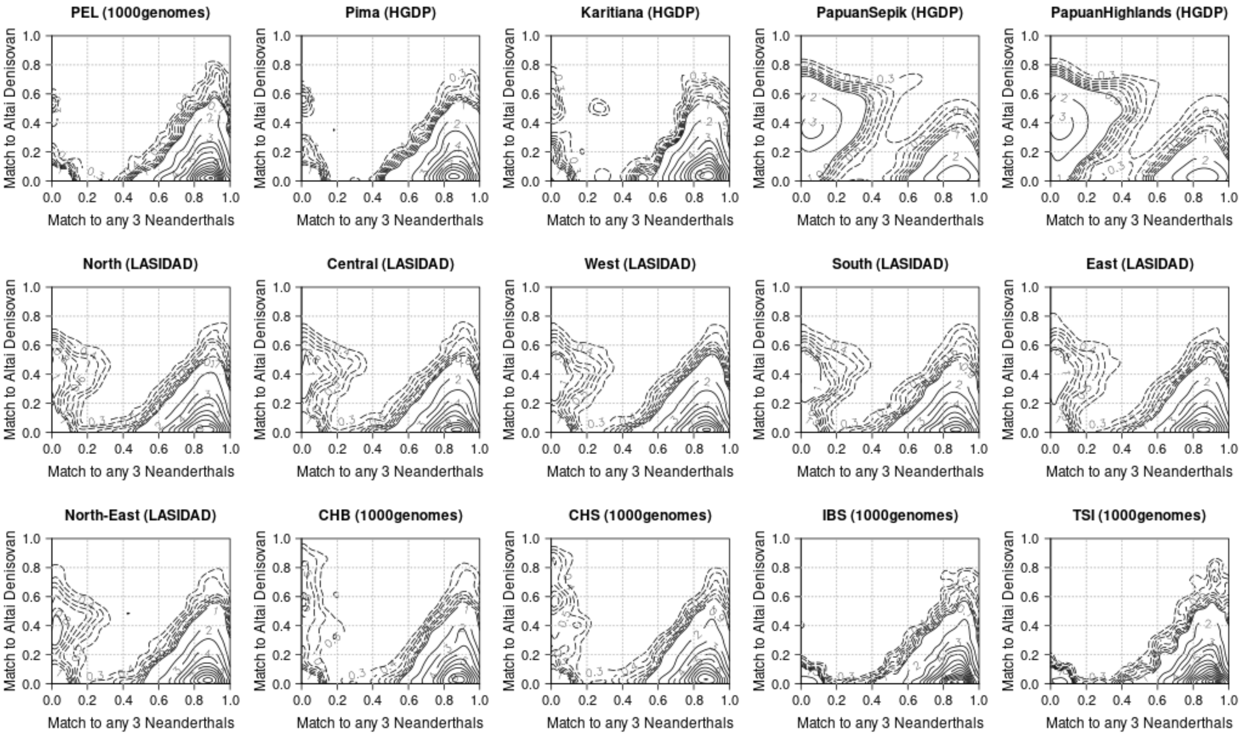

**Figure S6.9. Affinity of archaic ancestry segments in modern humans to sequenced archaic genomes.** The x axis indicates the proportion of derived variants in a given segment that match to any of the 3 sequenced Neanderthals and the y axis indicates the proportion of the derived variants in a given segment that match to the sequenced Denisovan genome.

To test for evidence for multiple pulses of archaic ancestry in Indians and other non-African populations, using the same approach as a previous study[49] (Figure S6.9). To this end, we retained all non-overlapping introgressed segments with a minimum of 30 sites which could be matched to the Denisovan genome and a minimum of 30 sites to any of the three sequenced Neanderthal genomes. In order to minimize the effect of incomplete lineage sorting, we required Neanderthal segments to contain no ND01 sites and all Denisovan segments to contain no ND10 sites. We restricted our set of sites to those which could be assessed in all 4 high coverage archaic genomes i.e., sites that fall within the intersection of the four individual manifesto maps created for. We only count the number of base pairs which passed all filters on the autosomes. The intersection of all four manifesto maps spans 1,402,898,253 bp of the autosomes.

We only considered populations with at least 50 Denisovan or Neanderthal segments in order to detect multiple admixture waves if they are present. This left us with 48 non-African populations.

*Denisovan or Denisovan-related ancestry components*

Following Browning et al. 2018[49], for each set of non-overlapping archaic ancestry segments we fitted a normal distribution (*mu1*, *sd1*) to the Denisovan match rate distribution and kept the parameters which maximize the likelihood. We then fitted a mixture of two normal distributions. We chose the means, standard deviations, and the mixture component *w* of the two normal distributions (*mu1*, *sd1*) and (*mu2*, *sd2*) which maximized the likelihood using a grid search approach. We allowed *mu1* and *mu2* to vary between 0 and 1 with a step size of 0.02. We also allowed *sd1* and *sd2* to vary between 0 and 0.3 with a step size of 0.02. However, we required that 99% of the normal distribution is between 0 and 1 - meaning that the (mean + 3 standard deviations) is less than 1 and the (mean - 3 standard deviations) is greater than 0. We performed a likelihood ratio test with 3 degrees of freedom and corrected for multiple testing using Bonferroni correction for 48 tests (one test per population), leading to a *p-value* threshold for significance of 0.05/48 = 0.001.

**Table S6.10. Populations with evidence of two mixture components of Denisovan ancestry.** Populations sorted by proportion of high affinity Denisovan component *w*. We report the parameters for the two normal distributions (*mu1*, *sd1*) and (*mu2*, *sd2*) along with the *p*-value of the likelihood ratio test. We highlight the two populations from India in orange.

| Region | Population | Proportion of high-affinity Denisovan (w) | High affinity Denisovan | | Low affinity Denisovan | | p value | segments | Significant level |
|---|---|---|---|---|---|---|---|---|---|
| | | | mu1 | mu2 | sd1 | sd2 | | | |
| America | CLM | 0.28 | 0.86 | 0.04 | 0.42 | 0.14 | 1.17E-07 | 55 | ** |
| | PEL | 0.26 | 0.86 | 0.04 | 0.50 | 0.16 | 3.43E-09 | 59 | ** |
| | MXL | 0.18 | 0.86 | 0.04 | 0.50 | 0.16 | 1.16E-06 | 64 | ** |
| South Asia | Hazara | 0.30 | 0.80 | 0.06 | 0.50 | 0.16 | 1.29E-07 | 65 | ** |
| | Burusho | 0.14 | 0.86 | 0.04 | 0.50 | 0.16 | 3.44E-05 | 83 | ** |
| | Sindhi | 0.14 | 0.80 | 0.42 | 0.06 | 0.14 | 1.24E-02 | 83 | * |
| | North-East | 0.08 | 0.86 | 0.04 | 0.50 | 0.16 | 3.74E-06 | 272 | ** |
| | BEB | 0.04 | 0.86 | 0.50 | 0.04 | 0.16 | 1.76E-03 | 307 | * |
| | South | 0.04 | 0.84 | 0.04 | 0.46 | 0.14 | 7.96E-06 | 552 | ** |
| | Pathan | 0.04 | 0.92 | 0.50 | 0.02 | 0.16 | 9.63E-03 | 80 | * |
| | East | 0.02 | 0.84 | 0.50 | 0.02 | 0.16 | 1.24E-02 | 548 | * |
| | West | 0.34 | 0.58 | 0.42 | 0.14 | 0.14 | 5.65E-03 | 458 | * |
| | North | 0.38 | 0.58 | 0.42 | 0.14 | 0.14 | 7.87E-02 | 424 | n.s. |
| | GIH | 0.02 | 0.86 | 0.46 | 0.04 | 0.14 | 2.18E-01 | 182 | n.s. |
| | ITU | 0.02 | 0.92 | 0.50 | 0.02 | 0.16 | 6.44E-01 | 242 | n.s. |
| | PJL | 0.02 | 0.86 | 0.50 | 0.02 | 0.16 | 6.04E-02 | 228 | n.s. |
| | STU | 0.02 | 0.92 | 0.50 | 0.02 | 0.16 | 2.36E-01 | 252 | n.s. |
| | Central | 0.02 | 0.86 | 0.50 | 0.04 | 0.16 | 6.92E-02 | 427 | n.s. |
| East Asia | NorthernHan | 0.46 | 0.80 | 0.06 | 0.42 | 0.14 | 2.12E-10 | 52 | ** |
| | Miao | 0.36 | 0.80 | 0.06 | 0.50 | 0.16 | 2.37E-09 | 63 | ** |
| | Han | 0.36 | 0.80 | 0.06 | 0.50 | 0.16 | 0.00E+00 | 114 | ** |
| | Japanese | 0.36 | 0.80 | 0.06 | 0.42 | 0.14 | 5.19E-13 | 97 | ** |
| | Yi | 0.32 | 0.80 | 0.06 | 0.44 | 0.14 | 1.90E-07 | 69 | ** |
| | JPT | 0.30 | 0.80 | 0.06 | 0.50 | 0.16 | 9.99E-16 | 135 | ** |
| | Xibo | 0.30 | 0.80 | 0.06 | 0.50 | 0.16 | 9.77E-06 | 56 | ** |

| | | | | | | | | |
|---|---|---|---|---|---|---|---|---|
| | Tujia | 0.28 | 0.80 | 0.06 | 0.50 | 0.16 | 2.53E-07 | 68 | ** |
| | CHB | 0.26 | 0.86 | 0.04 | 0.50 | 0.16 | 0.00E+00 | 171 | ** |
| | Dai | 0.26 | 0.86 | 0.04 | 0.50 | 0.16 | 8.17E-10 | 64 | ** |
| | KHV | 0.24 | 0.80 | 0.06 | 0.50 | 0.16 | 1.11E-16 | 190 | ** |
| | Daur | 0.24 | 0.86 | 0.04 | 0.50 | 0.16 | 1.87E-09 | 61 | ** |
| | Yakut | 0.24 | 0.86 | 0.04 | 0.50 | 0.16 | 3.48E-12 | 91 | ** |
| | She | 0.24 | 0.86 | 0.04 | 0.50 | 0.16 | 1.13E-06 | 58 | ** |
| | CDX | 0.22 | 0.86 | 0.04 | 0.50 | 0.16 | 0.00E+00 | 154 | ** |
| | CHS | 0.22 | 0.86 | 0.04 | 0.50 | 0.16 | 0.00E+00 | 183 | ** |
| | Oroqen | 0.20 | 0.86 | 0.04 | 0.50 | 0.16 | 1.92E-07 | 61 | ** |
| | Tu | 0.18 | 0.86 | 0.04 | 0.50 | 0.16 | 1.79E-05 | 56 | ** |
| | Hezhen | 0.18 | 0.84 | 0.04 | 0.50 | 0.16 | 7.60E-05 | 55 | ** |
| | Lahu | 0.14 | 0.90 | 0.02 | 0.50 | 0.16 | 5.05E-06 | 55 | ** |
| | Mongolian | 0.14 | 0.92 | 0.02 | 0.50 | 0.16 | 2.25E-06 | 62 | ** |
| | Cambodian | 0.12 | 0.90 | 0.02 | 0.50 | 0.16 | 4.08E-06 | 54 | ** |
| Europe | FIN | 0.18 | 0.84 | 0.04 | 0.50 | 0.16 | 2.46E-05 | 54 | ** |
| Oceania | Bougainville | 0.50 | 0.50 | 0.42 | 0.16 | 0.14 | 4.15E-03 | 593 | * |
| | PapuanSepik | 0.28 | 0.60 | 0.12 | 0.42 | 0.14 | 9.30E-05 | 558 | ** |
| | PapuanHighlands | 0.26 | 0.62 | 0.12 | 0.42 | 0.14 | 4.32E-04 | 566 | ** |

Significant level. n.s = not significant, * = marginally statistically significant p-value < 0.05, ** = significant after multiple testing (Bonferroni correction)

A previous study found evidence of a high affinity Denisovan component study (*mu1*=0.84) and a low affinity Denisovan component (*mu2*=0.49)[49]. We also find evidence of a high affinity Denisovan component (*mu1*=0.84) and a low affinity Denisovan component (*mu2*=0.49) in multiple populations (Table S6.10). We recovered the high affinity Denisovan component in East Asian, American, European and South Asian populations. In India, this component is present in the North-East and South of India. The former is likely explained by the presence of recent East Asian ancestry in North-eastern groups in India (Supplementary Note S4). There is also some evidence for two pulses of Denisovan ancestry in East India and Central India, however, the *p-values* for likelihood ratio test are not significant after correcting for multiple hypothesis testing (*p*-value = 0.012 in East India, and *p*-value=0.069 in Central India). Furthermore, we find evidence of multiple Denisovan pulses in Oceanic populations. Papuan from Sepik and the Highlands have two Denisovan components with *mu1*=0.61, *mu2*=0.42 and a proportion of the higher affinity Denisovan component of 0.26 in line with previous studies[50]. We show a histogram of Denisovan match rates for three populations that have evidence for two pulses of Denisovan ancestry, with one component that is closely related to the sequenced Denisovan genome (Figure S6.10).

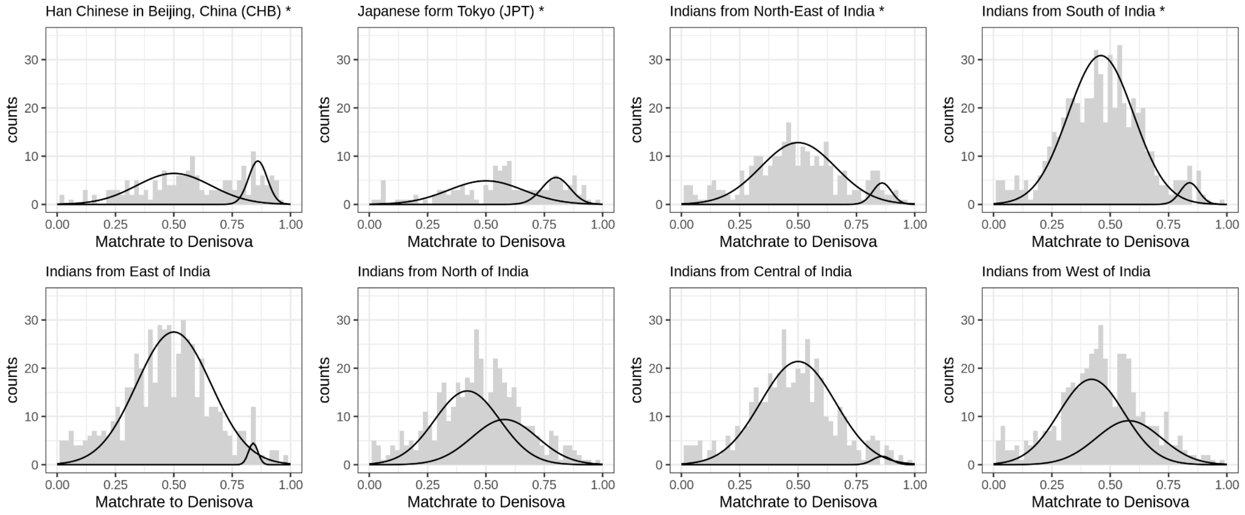

**Figure S6.10. Distributions of Denisovan match rates.** Histogram of Denisovan match rates for ND01 segments for Han Chinese (n=171 segments), Japanese (n=135 segments) North-East Indians (n=272 segments), South Indians (n=552 segments), East Indians (n=563), North Indians (n=433 segments), Central Indians (n=438 segments) and West Indians (n=471 segments). The normal distribution for the best fitting parameters for the high affinity Denisovan component (*mu1, sd1*) and low affinity Denisovan component (*mu2, sd2*) are shown for each. Only populations denoted with a * show significant statistical support.

*Neanderthal or Neanderthal-related ancestry components*

For each region we calculated the mean match rate to the three Neanderthals. We find that the introgressing Neanderthal is more closely related to the Vindija Neanderthal across all non-africans (Table S6.11). This is also consistent when looking at introgressed Neanderthal segments per population (Figure S6.11).There is no detectable difference in different affinities to Neanderthal genomes in any region we investigated - similar to what have previously been reported[49].

**Table S6.11. Match rate to sequenced Neanderthals.**

| Neanderthal | Region | Match rate | | |
|---|---|---|---|---|
| | | **Mean** | **2.5 percentile** | **97.5 percentile** |
| Altai | America | 0.75 | 0.37 | 0.97 |
| | East Asia | 0.75 | 0.33 | 0.95 |
| | Europe | 0.75 | 0.34 | 0.96 |
| | Middle East | 0.75 | 0.37 | 0.96 |
| | Oceania | 0.74 | 0.35 | 0.95 |
| | South Asia | 0.74 | 0.35 | 0.95 |
| Chagyrskaya | America | 0.81 | 0.43 | 1.00 |
| | East Asia | 0.81 | 0.43 | 1.00 |
| | Europe | 0.81 | 0.42 | 1.00 |
| | Middle East | 0.81 | 0.42 | 1.00 |

| | | | | |
|---|---|---|---|---|
| | Oceania | 0.81 | 0.42 | 1.00 |
| | South Asia | 0.81 | 0.41 | 0.98 |
| Vindija | America | 0.85 | 0.48 | 1.00 |
| | East Asia | 0.86 | 0.51 | 1.00 |
| | Europe | 0.85 | 0.46 | 1.00 |
| | Middle East | 0.85 | 0.43 | 1.00 |
| | Oceania | 0.85 | 0.42 | 1.00 |
| | South Asia | 0.85 | 0.48 | 1.00 |

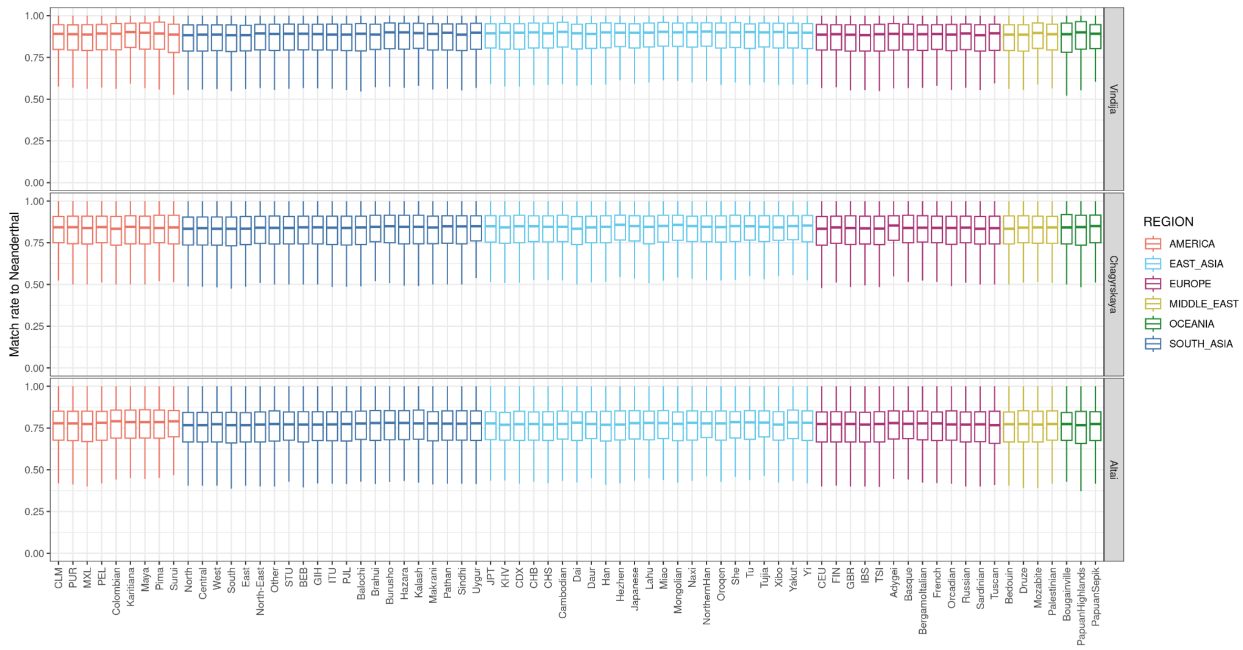

**Figure S6.11. Match rate for Neanderthal segments per population.** Each boxplot represents the distribution of Neanderthal max rates to the Altai, Chagyrskaya and Vindija Neanderthal. The box represents the first to the third quantile of the data with the black bar representing the median value. The whiskers indicate the first and third quantile -/+ 1.5 * Inter quantile range (IQR). The boxplots are colored by region.

Unlike the Denisovan match rate distribution - the Neanderthal match rate distribution is not well approximated by a normal distribution when we require that 99% of the density of the normal distribution is between 0-1 (Figure S6.12). We were thus unable to fit normal distributions to the data.

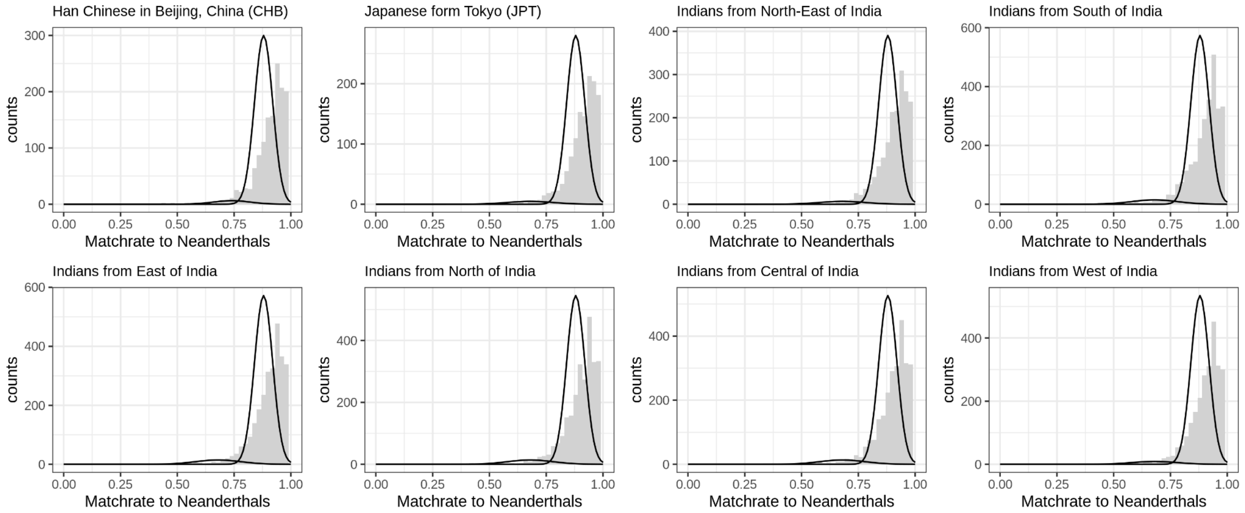

**Figure S6.12. Match rate for Neanderthal segments.** Histogram of Neanderthal match rates for ND01 segments for Han Chinese (n=1566 segments), Japanese (n=1463 segments) North-East Indians (n=2038 segments), South Indians (n=3059 segments), East Indians (n=3051), North Indians (n=2908 segments), Central Indians (n=2807 segments) and West Indians (n=2791 segments). The normal distribution for the best fitting parameters for the high affinity Neanderthal component (*mu1, sd1*) and low affinity Neanderthal component (*mu2, sd2*) are shown for each. Only populations denoted with a * show significant statistical support.

**Software availability**

Tutorial for hmmix version: 0.6.7
https://github.com/LauritsSkov/Introgression-detection

Python installation and tutorial:
https://pypi.org/project/hmmix/#description

### Supplementary Note 7: Genome-wide distribution of archaic ancestry in India

In this section, we investigate the genome-wide distribution of archaic ancestry in India. We identify regions enriched for Neanderthal or Denisovan ancestry in India and compare these results to other world-wide populations to identify regions that have uniquely increased in frequency in India. Finally, we search for regions that are depleted for archaic ancestry referred to as "archaic deserts" and compare them to previous reports.

#### Regions with high frequency of archaic ancestry segments

To detect regions that are enriched for archaic ancestry in Indians, we searched for regions with 'significantly high' archaic ancestry (defined as regions with inferred archaic frequency > mean plus 2*standard deviations). For Neanderthals, our threshold for significantly high archaic ancestry translates to regions with frequency >7.32% (average genome-wide ancestry = 1.5%) and for Denisovans, the threshold is >1.52% (average genome-wide ancestry = 0.1%) (Table S6.4-5). We then computed the mean archaic frequency for windows of 40,000 bp (or 40 Kb as in [51]) with a mean callability of >50% per window (i.e., where at least 50% of the bases are included in the accessible regions (Supplementary Note S6)). We found a total of 117.28 million bp (Mb) enriched for Neanderthal ancestry and 61.52 Mb enriched for Denisovan ancestry, represented in Figure S7.1.

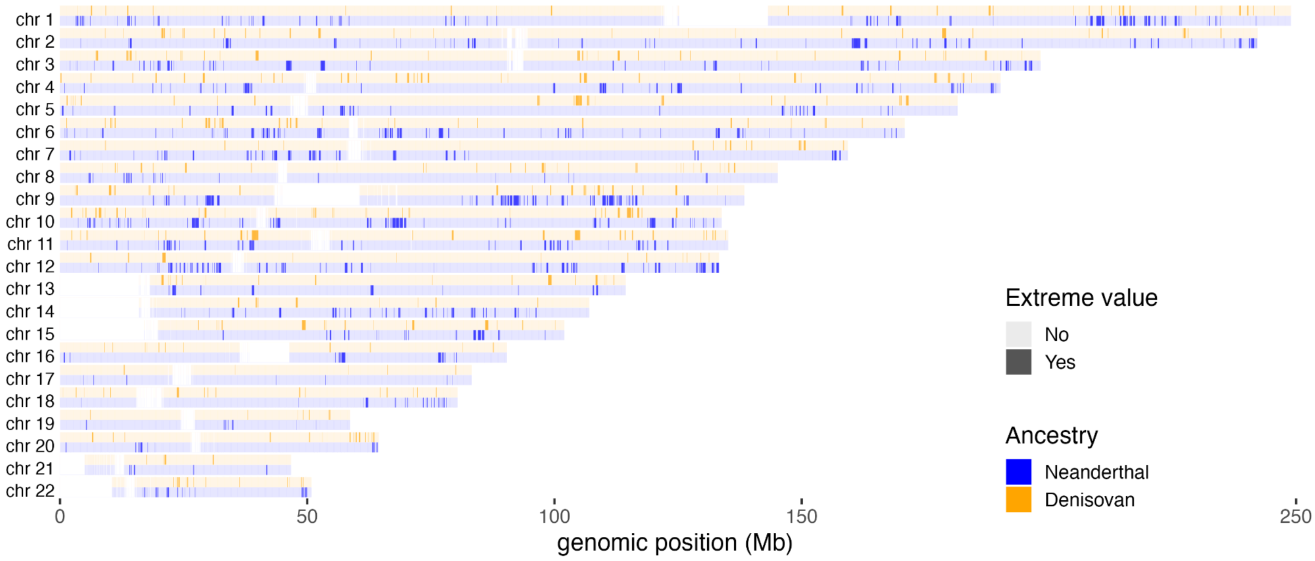

**Figure S7.1: Distribution of archaic ancestry regions across the genome.** We computed the mean archaic frequency along the genome of LASI-DAD individuals and considered segments with an archaic frequency higher than the mean + two standard deviations as enriched. We detected 117.28 Mb enriched in Neanderthal ancestry (in blue) and 61.52 Mb enriched in Denisovan ancestry (in orange).

**Table S7.1: Top five regions of highest frequency of archaic ancestry in Indians.**

| Archaic source | Position (hg38) | Frequency in LASI-DAD (%) | Genes | Reported in |
|---|---|---|---|---|
| Denisovan | chr1:147280000-147360000 | 24.8 | CHD1L | [51] |
| Denisovan | chr13:51720000-51760000 | 21.3 | WDFY2 | [51] |
| Denisovan | chr10:8240000-8280000 | 19.9 | LINC00708 | – |
| Denisovan | chr9:220000000-22040000 | 17.7 | CDKN2B; CDKN2B-AS1 | – |
| Denisovan | chr20:63520000-63560000 | 17.2 | FNDC11; HELZ2; PPDPF; PTK6; SRMS | HELZ2 [51] |
| Neanderthal | chr9:94520000-94560000 | 39.6 | PCAT7; FBP2 | FBP2 [51–53] |
| Neanderthal | chr16:76840000-77000000 | 38.2 | MIR4719 | [54] |
| Neanderthal | chr6:52280000-52320000 | 36.9 | MCM3 | [54] |
| Neanderthal | chr3:45880000-46120000 | 36.2 | CCR9; CXCR6; FYCO1; LZTFL1; XCR1 | FYCO1[51] |
| Neanderthal | chr5:56720000-56760000 | 32.7 | – | – |

#### *Gene Ontology (GO) analysis*

To assess the functional impact of archaic ancestry, we identified the genes and pathways that overlap the regions enriched for archaic ancestry by intersecting the high frequency archaic ancestry regions with the RefSeq database[55]. We identified 1,390 genes enriched for Neanderthal ancestry and 750 genes enriched for Denisovan ancestry (Extended Data Table S3). In Table S7.1, we show the top five genes for both archaic sources in Indians. Many of our most significant genes were identified in previous surveys of worldwide populations[51–54]. Next, we identified the pathways that are enriched for archaic ancestry using GeneSCF[56] exploring all functional categories including biological process, molecular function and cellular components, with the option '-db=GO_all' (GO database release date of 2023-01-05 that includes 1018 genes enriched for Neanderthal and 539 genes for Denisovan ancestry). We accounted for multiple hypothesis testing using the Benjamini and Hochberg method with a false discovery rate (FDR) threshold of 5%. We find 14 biological processes enriched for Neanderthal ancestry and 22 for Denisovan ancestry (Extended Data Table S4). Some of the processes— such as chemokine receptor activity and cellular response to zinc ion— were previously reported for enrichment of Neanderthal ancestry in East Asians, Europeans, Melanesians, and South Asians[54]. Chemokine receptors are important in inflammatory response and the innate immune system[57]. Several Neanderthal-enriched pathways are related to *Metallothionein 1* genes that have a broader role in inflammatory diseases and can shape innate and adaptive immunity[58]. We find Denisovan ancestry is enriched in genes associated with immune response, including clusters of differentiation 1 genes (CD1) and Human Leukocyte Antigens (HLA)-related genes, as previously reported[59].

#### Regions with Neanderthal-specific or Denisovan-specific variants

To identify regions enriched for Neanderthal-specific or Denisovan-specific variants, we examined the density of archaic SNPs that are present only in one archaic group and shared with modern humans: focusing on either Neanderthal-specific (ND10) or Denisovan-specific (ND01) variants (note: 0 refers to ancestral allele, 1 = derived allele). We find a total of 292,129 ND10 and 80,247 ND01 SNPs shared with Indians (where Indians have at least one derived allele). Next, we counted the number of ND10 and ND01 sites in 40 Kb windows across the genome. The average number of ND10 SNPs per window is 5.9 (ranging between 0-56, with the 99.99% tail of 49 SNPs) (Figure S7.2). In contrast, the average number of ND01 sites per window is lower: 2.6 (ranging between 0-103, with 99.99% tail of 51 SNPs). Notably, a region of chr 6 (chr6:32,360,000-32,400,000) contains 103 ND01 sites in a 40 Kb region. This region is part of the major histocompatibility complexes (MHC) that contain HLA genes such as BTNL2 that exhibit the strongest enrichment[60].

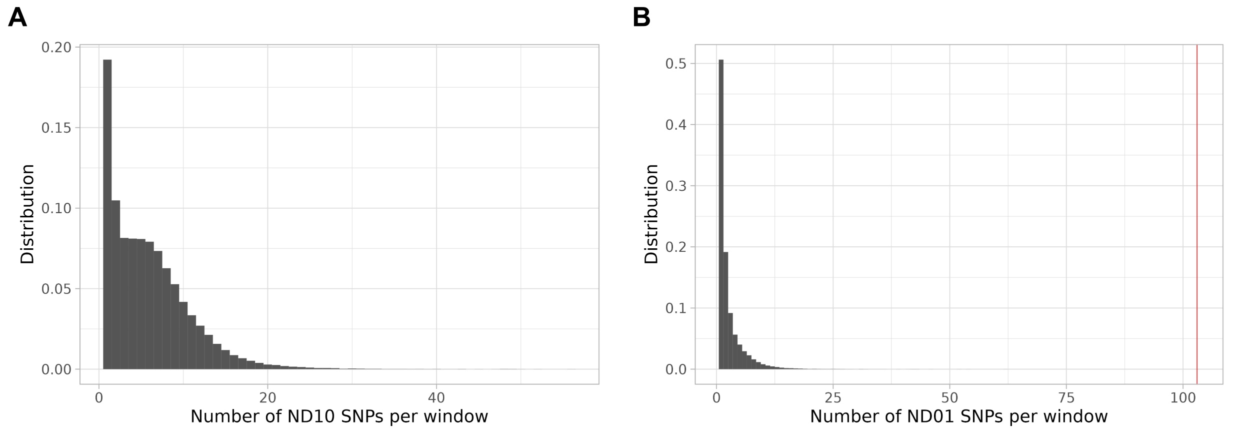

**Figure S7.2 Density of archaic specific SNPs in Indians.** Distributions of the number of ND10 (A) and ND01 (B) SNPs seen in Indian individuals per window of 40kb, conditioned on the presence of at least one ND10 or ND01 SNPs in the window. In red, the number of ND01 SNPs present on chr6:32,360,000-32,400,000.

*BTNL2*

A striking signal of enrichment of Denisovan ancestry in Indians is seen on chromosome 6. This region is part of the HLA complex (HLA-C) and plays a critical role in immune function[61,62]. At this locus, the frequency of Denisovan ancestry is ~9.4% in Indians, which is nearly 100-fold higher than the genome-wide average of ~0.1% (Table S6.4-5). More interestingly, there 103 ND01 variants in a 40 kb region. This region is flanked by several windows with high numbers of ND01 sites, including a gene BTNL2 with 78 ND01 sites in a 13.2 kb (on average, we observe 2.6 ND01 sites per 40Kb). Denisovan ancestry is also enriched in East Asians (~11.82%, >99% of the Denisovan frequency in EAS) but as expected at low (< 0.1%) frequency in other world-wide populations (Table S7.2). A recent scan of selection in modern humans identified BTNL2 as a candidate for balancing selection across 7 (of 12) geographic worldwide regions[63].

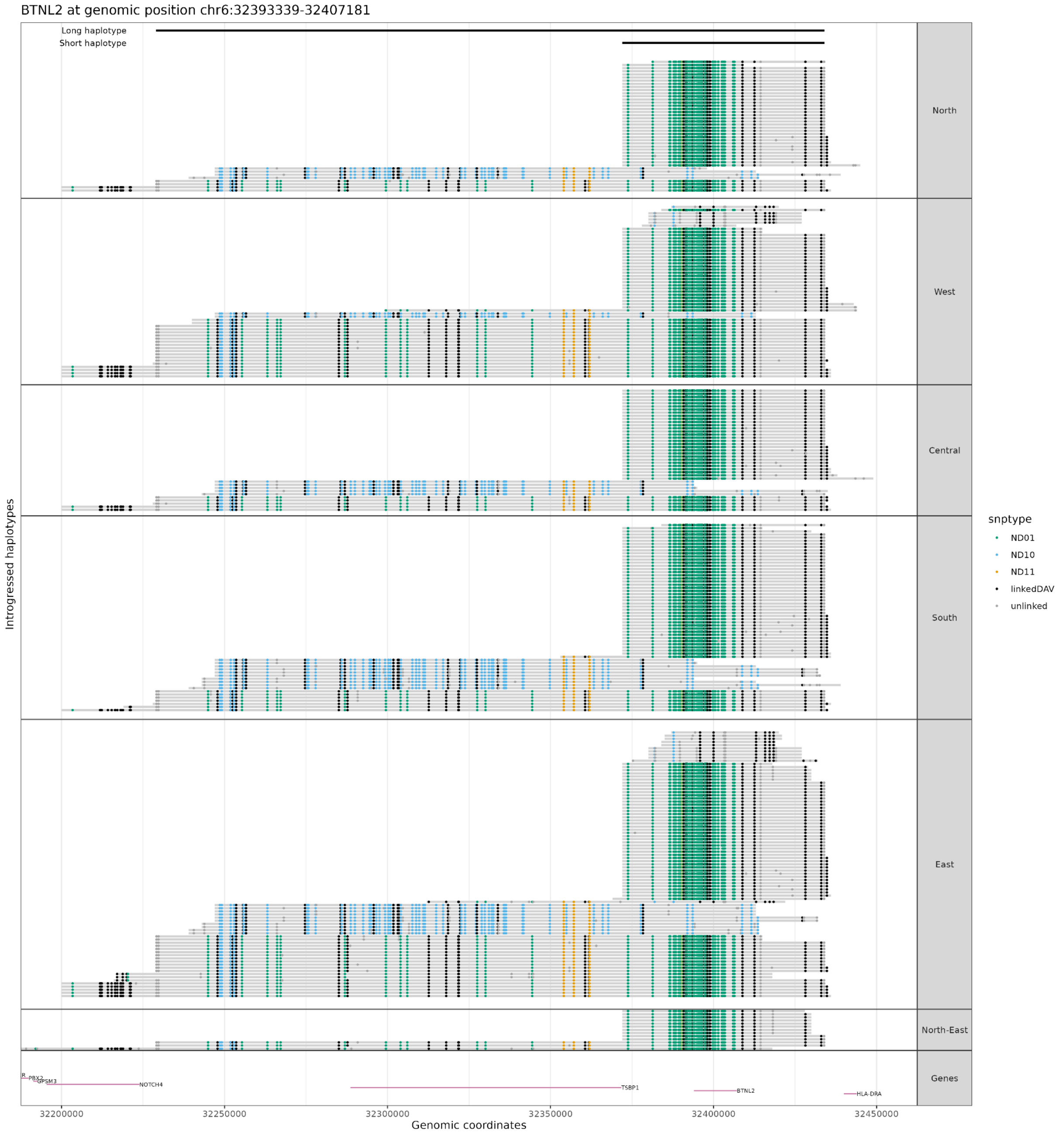

**Figure S7.3. Archaic ancestry haplotypes overlapping BTNL2 gene in Indian individuals.** The x-axis represents the genomic positions, the y-axis individuals who carried archaic haplotypes. In blue mutation Neanderthal-specific, in green Denisovan-specific and in orange mutations both present in Neanderthal and Denisovan. We denote the genomic location of the short and long haplotype.

At the BTNL2 locus, there are two main Denisovan haplotypes in India (Figure S7.3). A 'short' haplotype, between 55–75kb with an average length of 63.2 kb and containing an average of 116.1 ND01 variants, represents 71% of the haplotypes seen in India (Table S7.2). Another haplotype above 150 kb (referred to as 'long' haplotype) with 126.7 ND01 variants and is observed in ~23% of the individuals. In Europeans, only the short haplotype is observed. In East Asians, both haplotypes are present with the long haplotype seen in higher proportions than the short haplotype (Table S7.2). There is also marked variation across regions in India. The frequency of Denisovan ancestry in this region is significantly higher in East (12.9%, z-score=2.74) and North-East (17.1%, z-score=2.24) consistent with higher East Asian-related ancestry among these individuals; and significantly lower in the South (7.1%, z-score=-2.11) (Figure S7.4). The proportion of long haplotypes is significantly lower in the North (12.8%, z-score=-2.26) and significantly higher in the West of India (35.2%, *z*-score=2.57) (Figure S7.4).

**Table S7.2. Frequency and characteristics of the Denisovan haplotype overlapping BTNL2 genes in India, East Asia and Europe.**

| **Superpopulation (DATASET)** | **Denisovan haplotype percentage (Number)** | **Short haplotype** | | | **Long haplotype** | | |
|---|---|---|---|---|---|---|---|
| | | **Percentage (Number)** | **Length** | **#ND01** | **Percentage (Number)** | **Length** | **#ND01** |
| India (LASI-DAD) | 9.44 (506) | 70.9 (359) | 63.15kb | 116.1 | 23.7 (120) | 209.7kb | 126.7 |
| East Asia (1000G) | 11.82 (138) | 26.8 (37) | 63.46kb | 115.5 | 41.3 (57) | 211.6kb | 117.8 |
| Europe (1000G) | 0.395 (5) | 100 (5) | 65.6kb | 117.6 | 0 (0) | 0 | 0 |
| Oceania (HGDP) | 0 (0) | - | - | - | - | - | - |
| Middle East (HGDP) | 1.55 (5) | 100 (5) | 66kb | 117 | 0 (0) | 0 | 0 |
| America (1000G) | 1.22 (12) | 91.7 (11) | 65kb | 118.6 | 0 (0) | 0 | 0 |

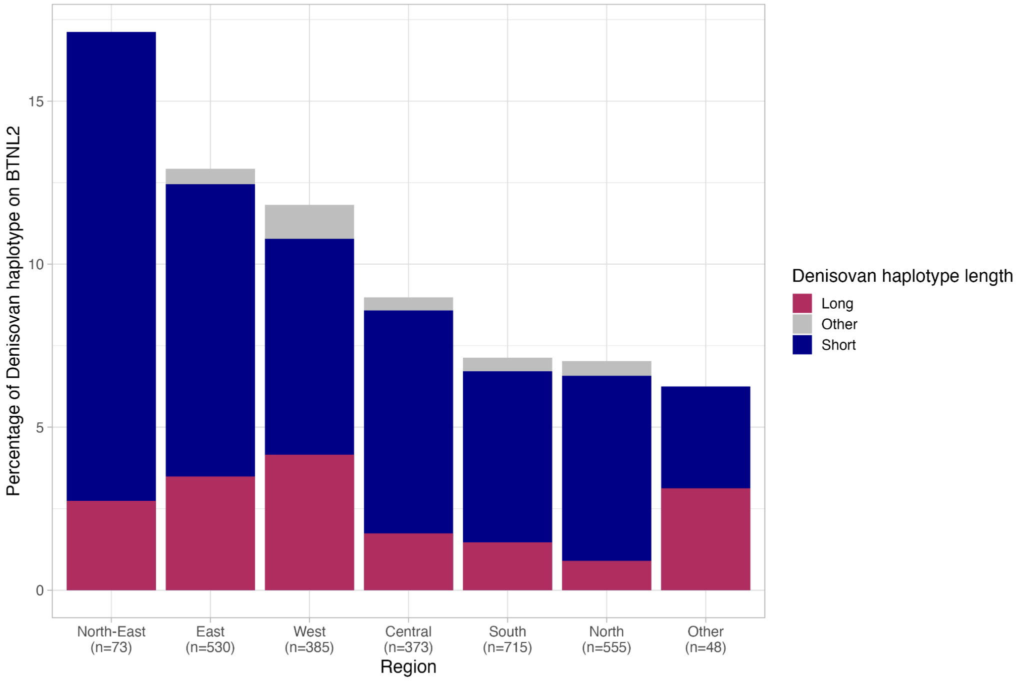

**Figure S7.4 BTNL2 Denisovan haplotype distribution in India.** Percentage of Denisovan haplotype on BTNL2 per region in India, considering both short (55-75kb; in blue) and long (>150kb, in red) haplotypes. Haplotypes with length below 55 kb or between 75 kb and 150 kb are in the 'Other' category (in gray). Denisovan haplotype frequencies are higher in the East (12.9%) and North-East (17.1%) and lower in the South (7.1%). The proportion of long haplotypes (>150Kb) is lower in the North (12.8%) and higher in the West (35.2%).

Next, we estimated the probability that a shared haplotype between Indians and Denisovans, of length at least 55–209 kb, could occur due to incomplete ancestral lineage sorting alone. Following Huerta-Sanchez 2014[64], we estimated the expected length ($L$) of the shared haplotype assuming a Gamma distribution with shape parameter of 2, and rate parameter lambda = $1/L$ where $L=1/(r \times t)$, with $r$ the recombination rate in the region and $t$ the time in generations. Assuming a recombination rate of $0.44\times10^{-8}$/bp/generation inferred for the MHC region[65], a generation time $g$ = 28 years[35] and divergence time of modern humans and Denisovans of 520,000 years (range: 520 to 630 ka[46]), we infer the expected length of the shared Denisovan haplotype would be $L = 1/(0.44 \times 10-8 \times (520{,}000/28)) = 12{,}238$ bp. In turn, the probability of maintaining the Denisovan haplotypes in modern humans by incomplete lineage sorting can be inferred as 1 − GammaCDF(209000, shape = 2,rate = 1/L). For the long haplotype (209 Kb), this probability is $< 10^{-6}$. Even for the shorter haplotype (63 Kb), we can significantly rule out incomplete lineage sorting ($p$-value = 0.027). These results support that the haplotypes in the MHC region in Indians are likely to be inherited through gene flow from Denisovan or Denisovan-related populations.

#### Population Branch Statistics Analysis

In order to detect Indian-specific high frequency archaic regions, we computed the population branch statistic (PBS)[66]. The PBS score measures the increase in allele frequency at a given locus in the population of interest, since its divergence from the two reference populations. Following Witt et al. 2023[53] , we ran the PBS test with archaic allele frequencies rather than genotype allele frequencies to identify signals of population-specific enrichment of archaic ancestry. Specifically, we estimated the frequency of archaic ancestry (Neanderthals and Denisovans) for each window of 1000 bp in three populations—Indians from LASI-DAD, EAS and EUR from 1000G. We excluded any regions with callability of < 50%.

For each pair of population, we computed the population frequency differentiation ($F_{ST}$) using archaic allele frequencies as:

$$F_{ST} = \frac{\bar{p}(1-\bar{p}) - \sum_i p_i(1-p_i)/2}{\bar{p}(1-\bar{p})}, \quad (1)$$

with $p_i$ the frequency in population $i$ and $\bar{p}$ the mean frequency between the two studied populations.

In order to obtain estimates of the population divergence time $T$ in units scaled by population size, we used the transformation as ([66]):

$$T = -log(1 - F_{ST}). \quad (2)$$

We computed PBS scores as:

$$\mathrm{PBS} = \frac{T^{\mathrm{LASIDAD-EUR}} + T^{\mathrm{LASIDAD-EAS}} - T^{\mathrm{EUR-EAS}}}{2}. \quad (3)$$

To assess the significance of the observed PBS score, we retained regions that fall in the 99% tail of the distribution of archaic frequency across the genome, considering Neanderthal and Denisovan ancestry separately, for sites with non-zero archaic ancestry in at least one of the three populations (Indians, EUR or EAS) (Figure S7.5). A significant PBS score includes both regions of high frequency in Indians, and regions that are depleted in East Asians or Europeans. As our focus is on the former, we applied two additional filters: (a) retain significantly high frequency archaic regions in India (which are in the 95% percentile of the genome-wide distribution) and (b) archaic allele frequency in Indians is greater than EAS and EUR.

After filtering, we find the amount of Indian-specific enriched archaic DNA as:

- Neanderthal: 10.7 Mb, which represents 9.1% of the total enriched Neanderthal;
- Denisovan: 5.5 Mb, which represents 8.9% of the total enriched Denisovan.

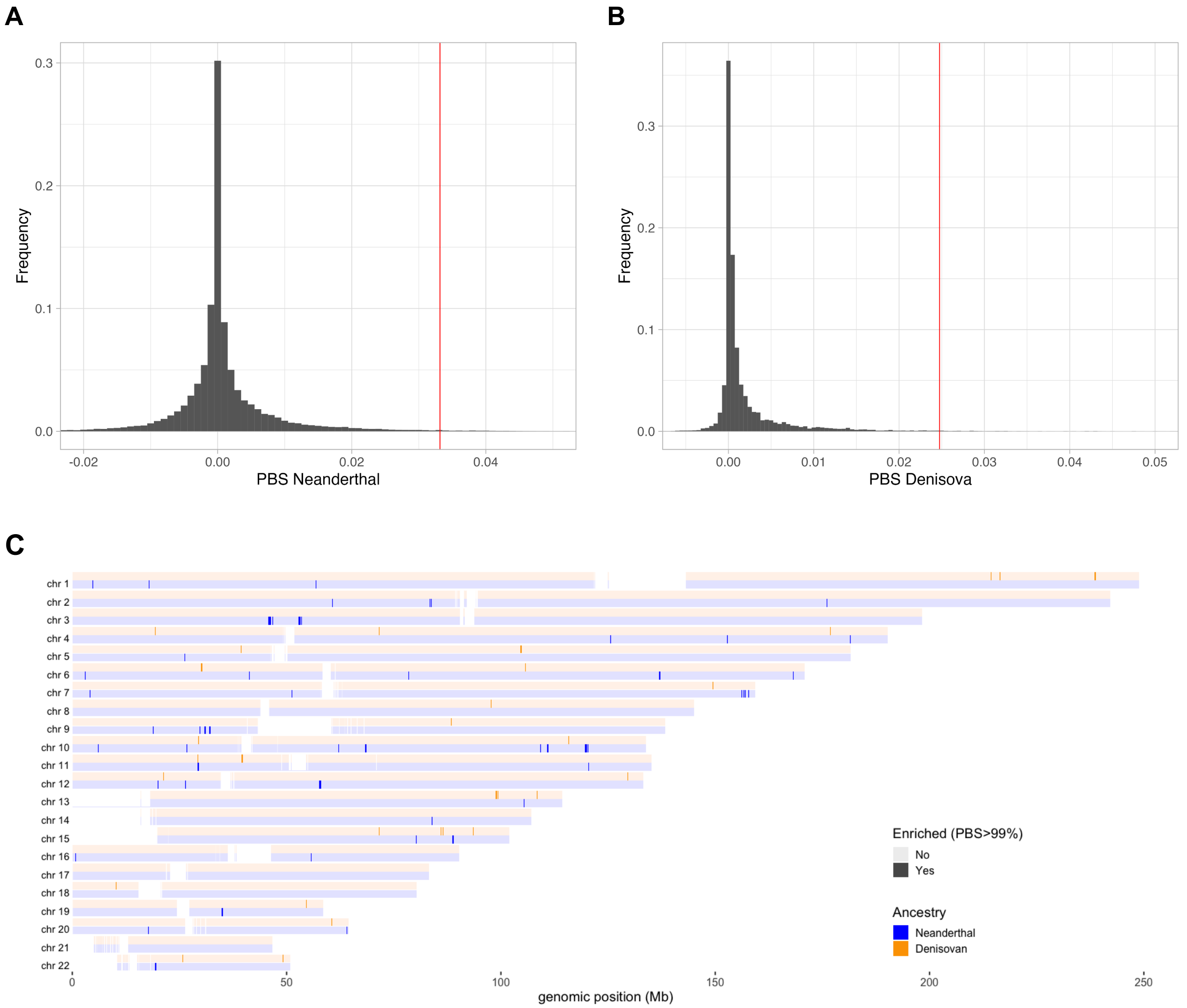

**Figure S7.5: Probabilistic Branch Statistic in Indians compared to European and East-Asians from 1000G.** PBS distributions in Indians for (A) Neanderthal and (B) Denisovan. We computed PBS between Indians from LASI-DAD and Europeans and East-Asians from 1000G using Neanderthal/Denisovan frequencies on 1kb windows. We report the values when at least one population has Neanderthal/Denisovan ancestry inferred on the 1kb window. In red the 99% percentile of the distributions, which is the cutoff for the analysis. (C) We report as enriched, genomic windows with PBS above the cutoff for Neadenrthal (in blue) and Denisovan (in orange).

*GO enrichment analysis of Indian-specific regions of archaic ancestry*

To identify the genes overlapping the Indian-specific archaic regions, we intersected the putative candidate regions (with significant PBS scores) with the RefSeq database (version 52). We obtained a list of 235 genes enriched for Neanderthal ancestry and 84 genes enriched for Denisovan ancestry in Indians (Extended Data Table S3). A subset of these genes, 174 and 64 genes enriched for Neanderthal and Denisovan ancestry respectively, are present in the GO reference list. Using GeneSCF[56] and the full GO database, we find 16 biological processes enriched in Neadenthal ancestry and 20 processes enriched in Denisovan ancestry (Extended Data Table S4). Neanderthal genes are enriched in chemokine receptor activity, similar to the high frequency archaic regions analysis (Extended Data Table S4). Denisovan genes are enriched in processes related to the solute carrier organic anion transporter (SLCO1) gene family that plays a major role for hepatic drug uptake and can be related to hyperbilirubinemia[67,68]; and in the TRIM family protein genes that places a role in innate immunity and resistance to pathogens[69].

Among the most significant candidate regions of Neanderthal ancestry is a gene cluster on chromosome 3 which has been previously associated to COVID susceptibility[70] ($PBS_{Neanderthal}$ > 0.118, in the top 0.015% highest PBS value). We report the distribution of Neanderthal introgressed segments overlapping this region in Table S7.3 and Figure S7.6.

**Table S7.3 Frequency of the Neanderthal haplotype overlapping COVID risk variants in India, East Asia, Europe and per region in India.**

| Superpopulation (DATASET) | Neanderthal haplotype frequency overlapping core haplotype (Number) | Neanderthal haplotype frequency overlapping long haplotype (Number) |
|---|---|---|
| East Asia (1000G) | 0.4% (5) | 0.3% (4) |
| Europe (1000G) | 7.5% (95) | 5.8% (73) |
| India (LASI-DAD) | 29.5% (1579) | 19.2% (1030) |
| Central India (LASI-DAD) | 29.2% (218) | 19.4% (145) |
| East India (LASI-DAD) | 34.8% (369) | 23.2% (246) |
| North India (LASI-DAD) | 26.2% (291) | 18.5% (205) |
| North East India (LASI-DAD) | 20.5% (30) | 13.0% (19) |
| South India (LASI-DAD) | 28.7% (410) | 18.4% (263) |
| West India (LASI-DAD) | 30.3% (233) | 16.9% (130) |
| Other (LASI-DAD) | 29.2% (28) | 22.9% (22) |

**A**

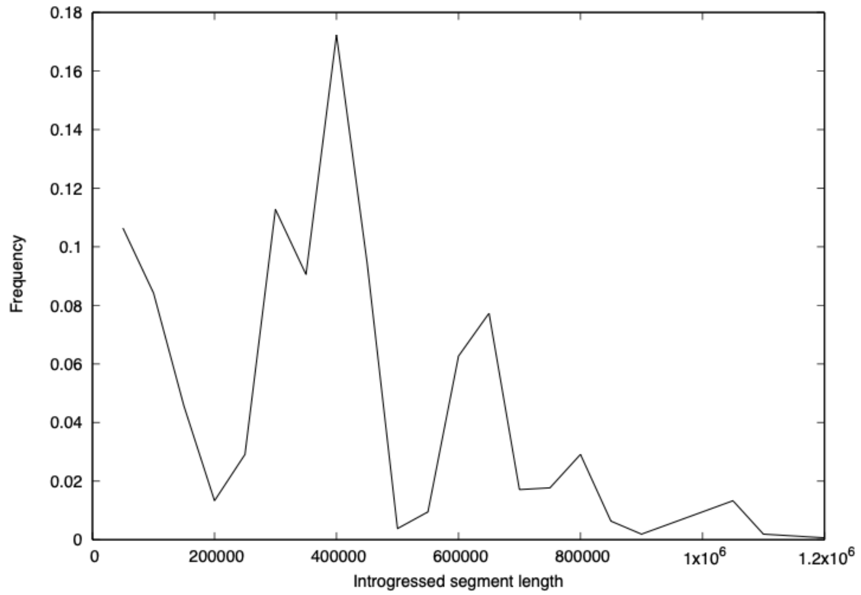

**B**

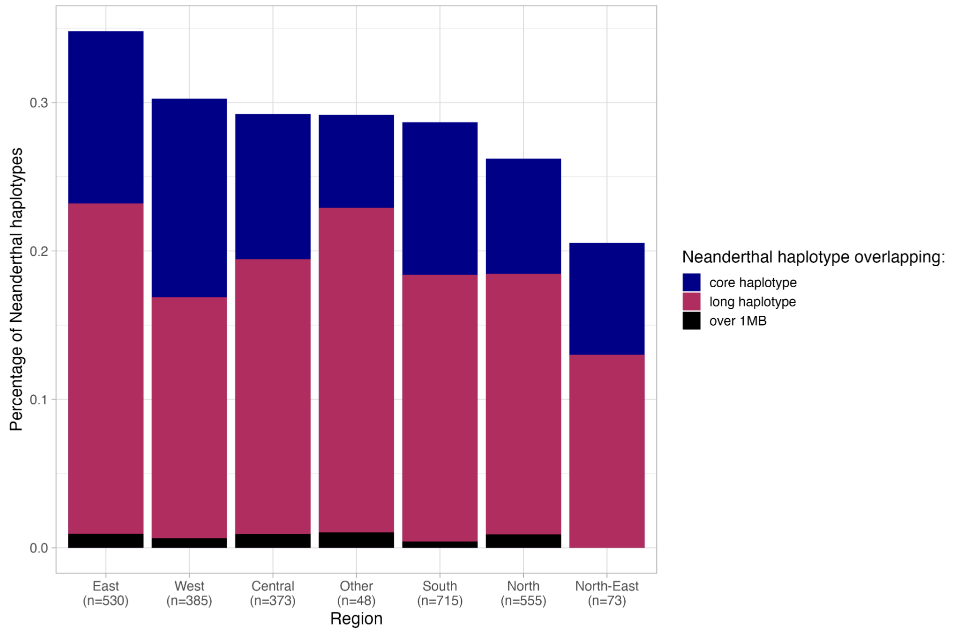

**Figure S7.6 Neanderthal haplotypes overlapping the COVID risk variants.** (A) Length distribution of the Neanderthal haplotypes overlapping positions the core haplotype: chr3:45,818,159-45,867,532 (in hg38). (B) Percentage of Neanderthal haplotype overlapping the core haplotype, the long haplotype : chr3:45,801,823-46,135,604 (in hg38) and the core haplotype with a segment longer than 1MB, per region in India. Segment lengths are in bp.

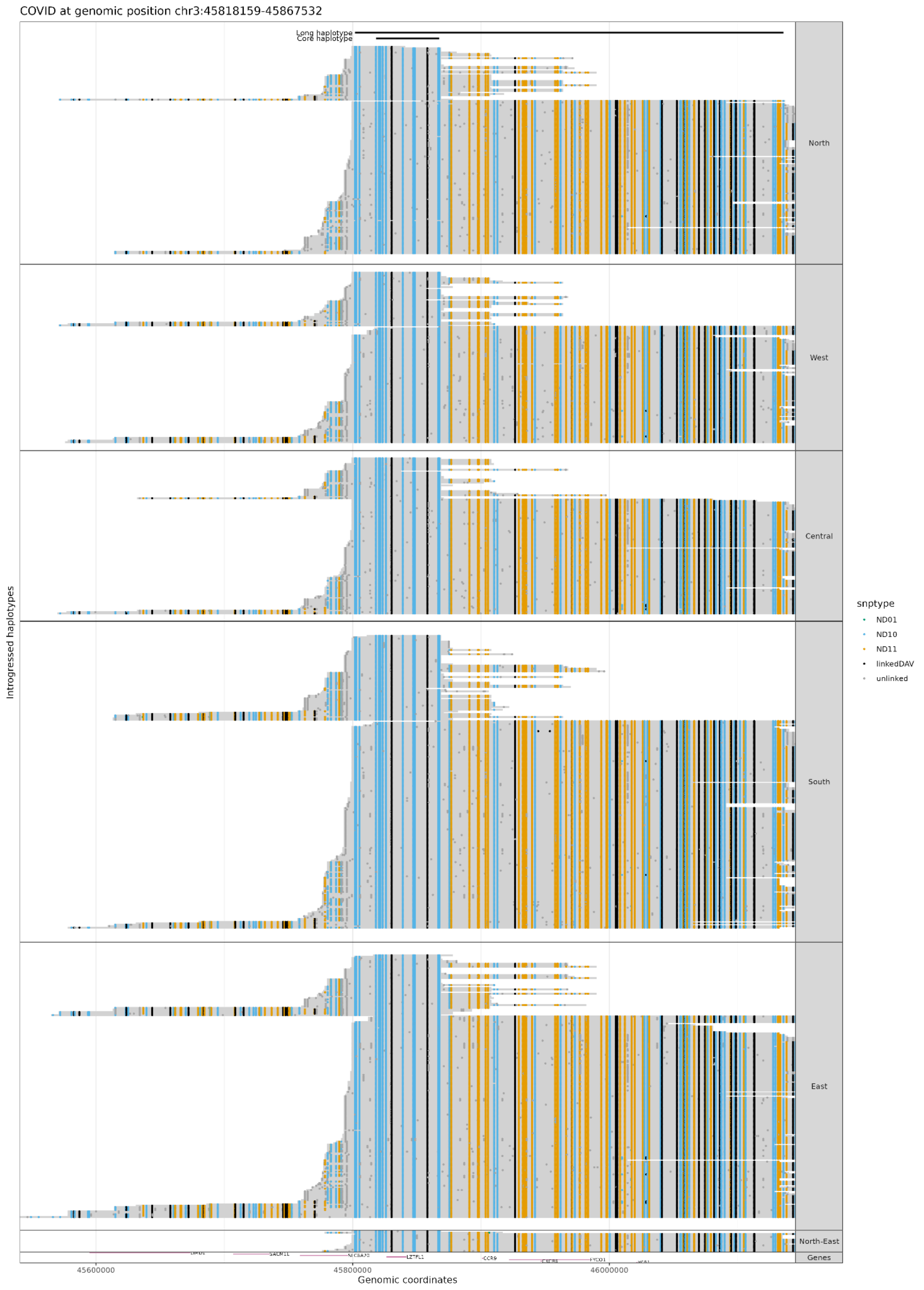

COVID at genomic position chr3:45818159-45867532
Long haplotype
Core haplotype
North
West
Central
South
East
North-East
Genes
Introgressed haplotypes
LZTFL1
CCR9
FYCO1
45600000
45800000
46000000
Genomic coordinates
snptype
ND01
ND10
ND11
linkedDAV
unlinked

**Figure S7.7 Archaic haplotypes overlapping the region overlapping the covid risk variant.** The x-axis represents the genomic positions, the y-axis individuals who carried archaic haplotypes. In blue mutation Neanderthal-specific (ND10), in orange mutations both present in Neanderthal and Denisovan (ND11) and in black mutations linked to an archaic SNP. We show the position of the core and long haplotype.

**Regions depleted for archaic ancestry ('archaic ancestry deserts')**

It has previously been shown that some regions of the genome are devoid of archaic ancestry in modern humans, referred to as 'archaic ancestry deserts'[41,43,52,71]. Given the large sample size of individuals from diverse populations from India, we revisited the evidence of deserts in modern humans as well as examined if the boundaries of the inferred archaic deserts in Indians are similar to previous findings.

We defined archaic deserts as regions of least 10 Mb long with an average frequency of introgressed archaic ancestry to be less than 0.1% for Neanderthals and 0.01% for Denisovans. For reference, the average genome-wide Neanderthal ancestry proportion is 1.3% and the average Denisovan ancestry proportion is 0.12% at a posterior probability cutoff at 0.5 in Indians. Thus, our cutoffs roughly correspond to a tenth of the genome-wide average frequency. We required that the number of called bases in the region is greater than 70% (based on the accessible regions described earlier). To minimize under-calling of archaic ancestry and so, over-calling deserts, we considered all Neanderthal and Denisovan segments with a posterior probability greater than 0.5. We merged individuals from the same region and called deserts for each region; Oceania (n=28), East Asia (n=808), Europe (n=788), South Asia (HGDP and 1000G, n=798), South Asia (LASI-DAD, n = 2,679) and Americas (n=551).

We identified 6 Neanderthal deserts including five that were previously reported in Europeans and other populations (Table S7.4). The deserts found in LASI-DAD span 87.1 Mb where 60.02 Mb overlaps with previously identified deserts.

We show the inferred deserts along with previously reported deserts in Figure S7.8. The location of previously identified Neanderthal deserts remain the same for five deserts (Table S7.4).

**Table S7.4. Chromosome positions (in hg38) for six Neanderthal ancestry deserts in India.**

| chromosome | start | end | Length (Mb) | Frequency (%) | Reported in |
|---|---|---|---|---|---|
| 3 | 76800000 | 90400000 | 13.6 | 0.01 | [41,43,52,71] |
| 5 | 83000000 | 94300000 | 11.3 | 0.008 | [43] |
| 7 | 106700000 | 128200000 | 21.5 | 0.015 | [41,43,52,71] |
| 8 | 52400000 | 65400000 | 13.0 | 0.010 | [41,43,71] |
| 8 | 107800000 | 121500000 | 13.7 | 0.019 | novel |
| 18 | 32300000 | 46300000 | 14.0 | 0.021 | [43,71] |

In addition to the newly identified desert in individuals from India (LASI-DAD) and South Asia (1000G + HGDP) we also find 21 deserts in East Asia, 20 in Oceania, 6 in the Middle East, 7 in America and 5 in Europe . We show all deserts (including known) in Table S7.4.

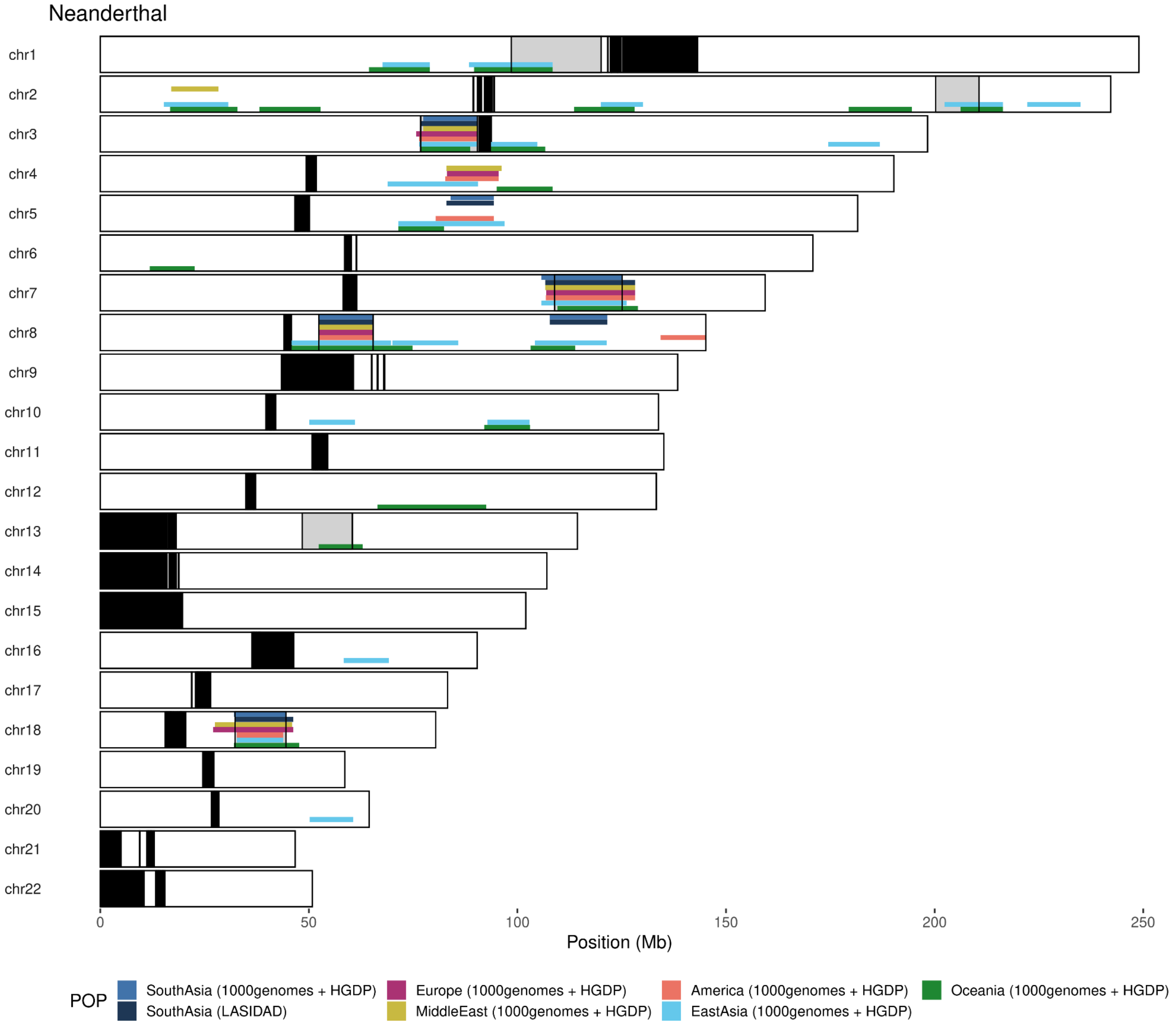

**Figure S7.8. Location of archaic ancestry deserts (regions with <0.1% Neanderthal ancestry over 10 Mb) called for each region.** We show the position of centromeres and telomeres in black and the location of previously identified deserts in grey boxes with black outlines.

We identified 13 Denisovan deserts in Indians (Table S7.5). Only one overlaps with previously reported Neanderthal deserts (Table S7.4). Given the low genome-wide proportion of Denisovan ancestry in Indians, we are likely over-calling some Denisovan deserts. This is also apparent from the cumulative distribution of recovery of Denisovan ancestry segments from Indians which has not yet plateaued (Figure S6.4). We show the locations of Denisovan deserts in Figure S7.9 and Table S7.5.

**Table S7.5. Chromosome positions (in hg38) for 13 Denisovan deserts.**

| chrom | start | end | Length (Mb) | Frequency (%) | Reported in |
|---|---|---|---|---|---|
| 1 | 108400000 | 118800000 | 10.4 | 0.004 | |
| 2 | 98400000 | 108700000 | 10.3 | 0.003 | |
| 2 | 147700000 | 158000000 | 10.3 | 0.002 | |
| 3 | 77000000 | 90400000 | 13.4 | 0.001 | 41,43,52,71 |
| 3 | 93700000 | 104600000 | 10.9 | 0.001 | |
| 3 | 130600000 | 143600000 | 13 | 0.002 | |
| 5 | 58800000 | 69500000 | 10.7 | 0 | |
| 7 | 81700000 | 97900000 | 16.2 | 0.001 | |
| 10 | 96400000 | 106800000 | 10.4 | 0.002 | |
| 12 | 78900000 | 93800000 | 14.9 | 0.002 | |
| 13 | 52300000 | 63000000 | 10.7 | 0.001 | |
| 14 | 53400000 | 64500000 | 11.1 | 0.003 | |
| 14 | 76300000 | 90500000 | 14.2 | 0.003 | |

We identify the same 13 deserts in South Asia (1000G + HGDP), 31 deserts in East Asia, 17 in Oceania, 65 in the Middle East, 64 in America and 74 in Europe (Extended Data Table S4).

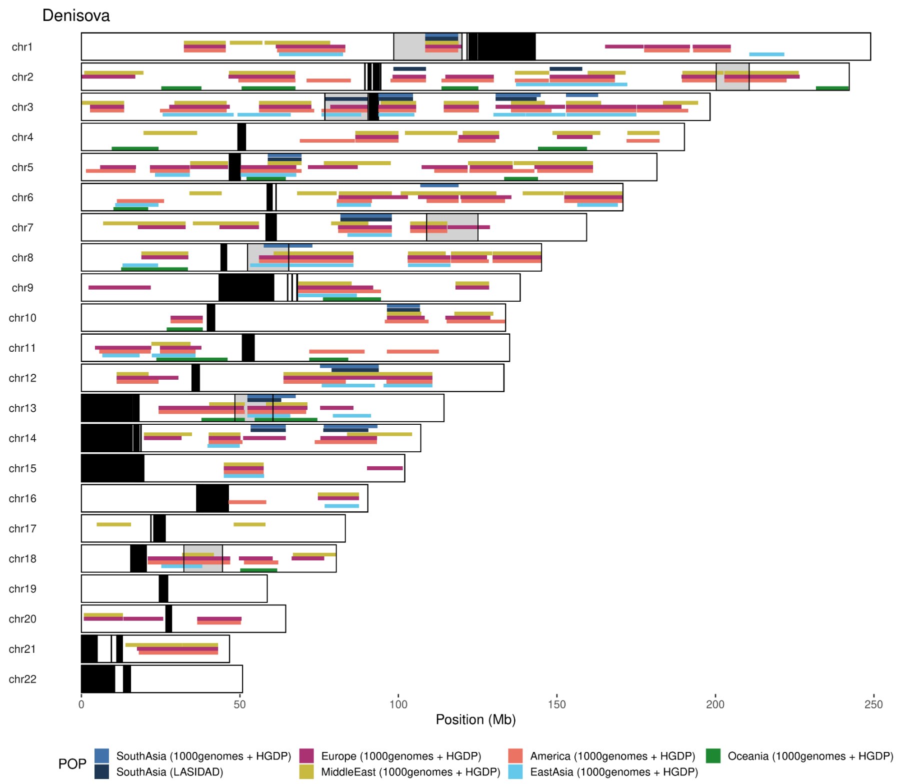

**Figure S7.9. Location of archaic ancestry deserts (regions with <0.01% Denisovan ancestry over 10 Mb) called for each region.** We show the position of centromeres and telomeres in black and the location of previously identified deserts in grey boxes with black outlines.

### Supplementary Note 8: Simulations to evaluate the genomic distribution of archaic ancestry

Archaic ancestry varies among populations, as well across the genome (Supplementary Note 6-7). While natural selection may have played a role in increasing the frequency for beneficial or adaptive variants in different regions or populations, an alternative hypothesis is that demographic history, in particular founder events, could have contributed to these patterns. Previous studies[72–74] and our analysis shows that many South Asian groups have experienced strong founder events in their recent past. Founder events lead to increased genetic drift which causes rapid fixation (or loss) of variants in a population[75]. Thus, we investigate the impact of founder events on the frequency of archaic introgressed variants.

To this end, we performed simulations where we varied the strength of the bottleneck in a population of interest and assessed its impact on the distribution and number of candidate regions exhibiting significantly high archaic ancestry or 'outlier' regions that are often considered as 'candidate regions for adaptive introgression'. Specifically, we measured two summary statistics: (1) the 95% tail of the distribution of the frequency of archaic ancestry across the genome, and (2) the number of candidate regions with archaic frequency of >20%.

#### *Simulation Scenarios*

We performed coalescent simulations for demographic scenarios mimicking the history of out of Africa (OOA) bottleneck and archaic gene flow of 1.3%, into the common ancestor of non-Africans using *msprime*[76] (Figure S8.1). For each simulation, we generated a genome sequence corresponding to the human chromosome 14 with a length of 106,880,170 bp. We used the recombination rate based on the HapMap recombination map for chromosome 14[24]. We considered three different scenarios: '*Constant*', '*Single bottleneck*' and '*Two population bottlenecks*'; illustrated in Figure S8.1 with the similar parameters for all scenarios.

For each simulation, we sampled 500 chromosomes from the 'South Asia' population. We used tree sequences from *msprime* to identify the location of outlier regions, their lengths and their frequencies in each individual. As we have access to the tree sequence information, we obtained the exact introgressed segment lengths without introducing mutations. We generated 50 independent simulations of chromosome 14 and performed bootstrap resampling (with replacement) to estimate the mean and standard deviation for each summary statistic.

The *msprime* command for the constant scenario:

```
demography= msprime.Demography()
#initializing an msprime "demography"
# This is the "trunk" population, here the South Asian population:
demography.add_population(
    name="South Asia",
    initial_size=N_{South Asia},
    initially_active=True)
##Neanderthal population
demography.add_population(
    name="N",
    description="Introgressing-Neanderthal population",
    initial_size=N_{N},
    initially_active=True)

# Add neanderthal gene flow for one generation (pulse model)
demography.add_migration_rate_change(time = t_{N-SA}, rate = a_{N}, dest = "N", source
= "South Asia")
demography.add_migration_rate_change(time = t_{N-SA}+1, rate = 0, dest = "N", source
= "South Asia")

# Add split out of Africa
demography.add_population_parameters_change(
    time=t_{OOA},population="South Asia", initial_size=N_{AH})
demography.add_population_parameters_change(
    time=t_{SA},population="South Asia", initial_size=N_{OOA})
# Add split with Neanderthal
demography.add_population_split(
    time=t_{N}, derived=["N"], ancestral="South Asia" )

demography.sort_events()
```

Model parameters:

```
# Times provided in years, then converted in generations.
gen_time = 29.0 #generation time
# Population sizes
N_{N} = 1_500
N_{South Asia} = 10_000
N_{OOA} = 2_000
N_{AH} = 7_000
# Split Times - converting years BP into generations BP
t_{N} = 575_000 / gen_time
t_{OOA} = 60_000 / gen_time
t_{SA} = 50_000 / gen_time
# Admixture times and proportions from archaic humans
a_{N} = 0.013 #proportion
t_{N-SA} = t_{SA} = 50_000 / gen_time #time
```

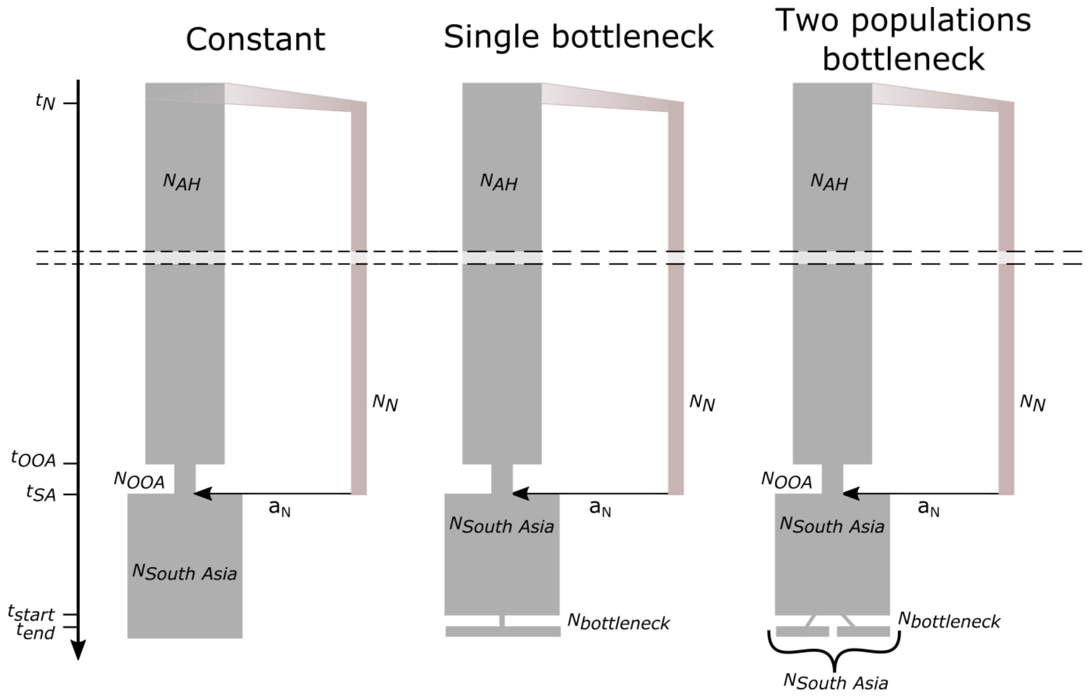

**Figure S8.1. Three scenarios simulated using msprime.** With $N_N$ population size of Neanderthal, $N_{South\ Asia}$ population size of South Asia, $N_{OOA}$ population size for the Out-Of-Africa (00A) bottleneck, $N_{AH}$ population size at split time between Humans and Neanderthal and $N_{bottleneck}$ population size during the bottleneck. $t_N$ split time between human and Neanderthal lineages, $t_{OOA}$ OOA bottleneck timing, $t_{SA}$ end of OOA bottleneck and time of Neanderthal introgression and $a_N$ introgression coefficient.

We performed simulations with varying strengths and timing of bottlenecks in the South Asian population. In the 'Single bottleneck' scenario, bottleneck time varies between 100 and 30 generations ago ($t_{start}$= *100–30*) and lasts for 10 generations ($t_{end}$ = $t_{start}$*-10*). We also varied the population size during the bottleneck ($N_{bottleneck}$ = *[5000–10]*). At the end of the bottleneck, the population size returned to its original value of $N_{South\_Asia}$ = *10,000*.

Bottleneck parameters:

$N_{bottleneck}$: 5000 - 1000 - 100 - 90 - 70 - 50 - 30 - 10

$t_{start}$: 100 - 30 generations

Our LASI-DAD dataset is very heterogeneous and contains individuals with diverse histories of founder events. To mimic this scenario, we generated data for two founder groups with recent bottlenecks and then combined individuals from these groups together into a single population ('*Two populations bottleneck*' Figure S8.1). This has the effect that the frequency of archaic segments will be computed on the mixture of the two populations. The sizes of the populations during the bottleneck are similar in both groups and equal to $N_{bottleneck}$=*100*. We also considered two different timing of bottlenecks ($t_{start}$=100-30 generations), bottlenecks last 10 generations ($t_{end}$=$t_{start}$-10). At the end of the bottleneck, each of the two population sizes returned to *N* = ½*

$N_{South\_Asia}$, so the overall South Asian population size is similar in the scenarios with only one bottleneck ($N_{South\_Asia}*1/2 + N_{South\_Asia}*1/2$ vs. $N_{South\ Asia}$) (Figure S8.1).

**Distribution of archaic ancestry in simulations**

We observe marked impact of the population bottlenecks on the frequency of archaic ancestry and the number of outlier regions detected in simulations. With an extreme bottleneck (e.g., $N_{bottleneck} = 10$), we find that many archaic segments get either lost or fixed, generating a bimodal distribution with modes at *0* and *1*. Even for less extreme bottlenecks (i.e., $N_{bottleneck}=100–1000$), the distributions of archaic frequency differ compared to the constant population size model (Figure S8.2). For instance, we find there are significantly more archaic segments at high frequencies (>20%), though there is also a simultaneous loss of archaic segments overall (Figure S8.3.A). Moreover, the 95% threshold to assess outliers depends on the population history and varies as the function of the strength of the bottleneck. Under our constant population size simulation, we infer the threshold to call outliers would be 6.9% (compared to 1.3% genome-wide). In contrast, this threshold is lower for populations with strong bottlenecks, e.g., ~5.8% under the single bottleneck scenario with $N_{bottleneck}$=100 and $t_{start}=100$ or nearly zero for extreme bottlenecks (with $N_{bottleneck}= 30–10$) as a large fraction of archaic variants are lost due to drift (Figure S8.3.B). Although when archaic segments are retained (frequency > 0), there are significantly more high frequency archaic segments compared to the constant scenario (Figure S8.3.A). The pattern is quite different under the two populations bottleneck scenario. In this case, we find the number of high frequency archaic segments still increases significantly compared to the constant scenario, but the genome-wide threshold is similar to the constant population size scenario (Figure S8.3.C). We note, our simulation scenario has no selection and so the majority of these results are due to drift alone.

In summary, our simulations show that population bottlenecks can lead to an increase in high frequency archaic regions, often interpreted as candidates of adaptive introgression, due to drift alone. In addition, population bottlenecks also impact the loss and depletion of archaic variants both at low and intermediate frequencies, impacting the threshold for identifying adaptive variants. Thus, for populations with a history of bottlenecks, caution is warranted in interpreting high frequency archaic variants as candidates of adaptive introgression.

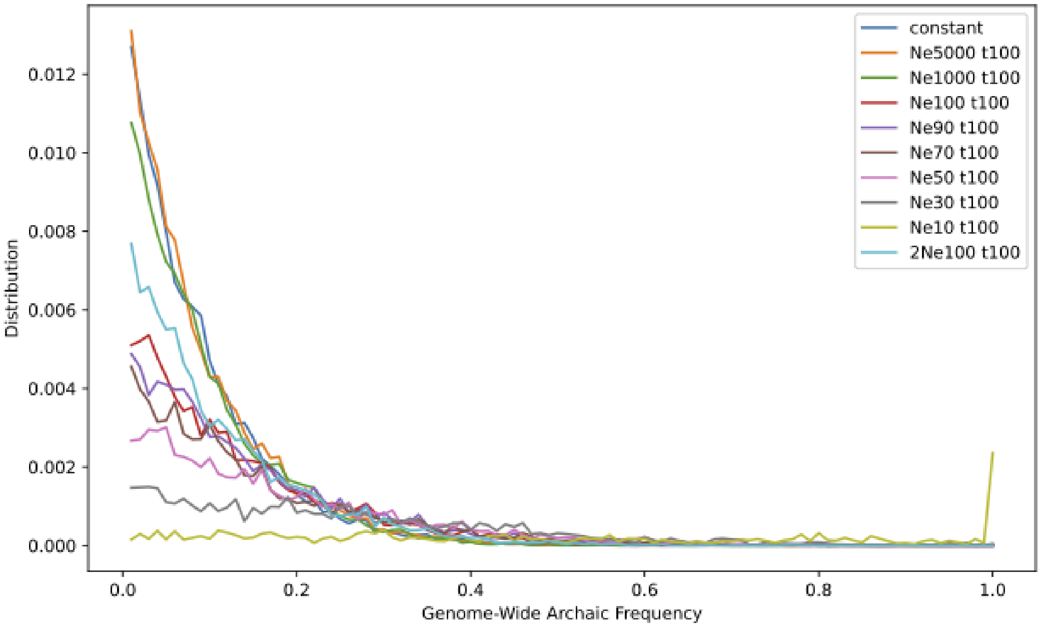

**Figure S8.2. Genome-wide archaic frequency for the different scenarios.** We fix $t_{start}$=100 and vary different sizes during the bottleneck ($N_{bottleneck}$). This distribution is truncated, we removed the first bin [0,0.01[.

**A**

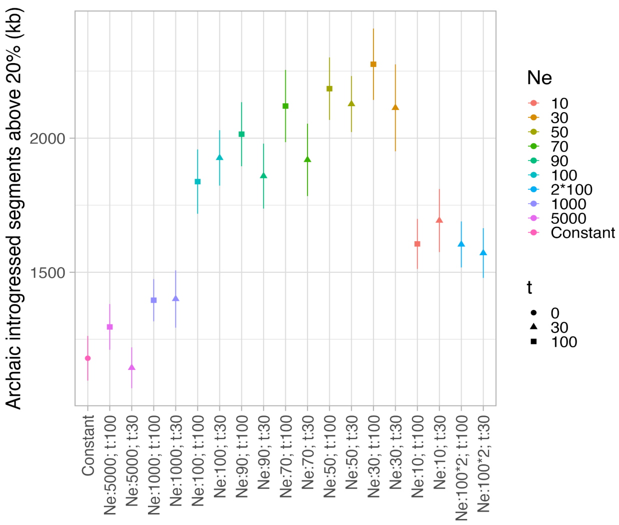

**B**

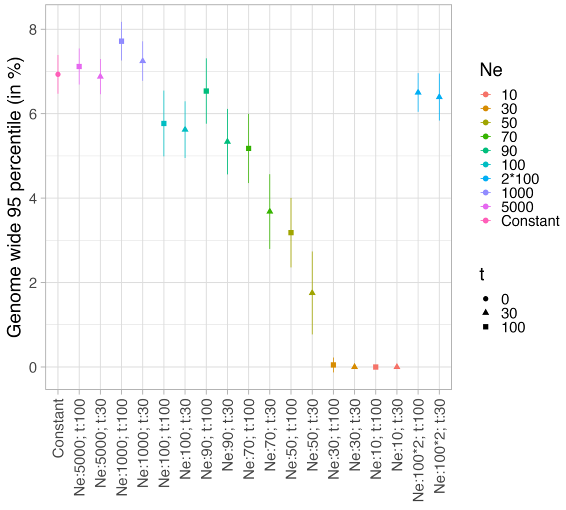

**C**

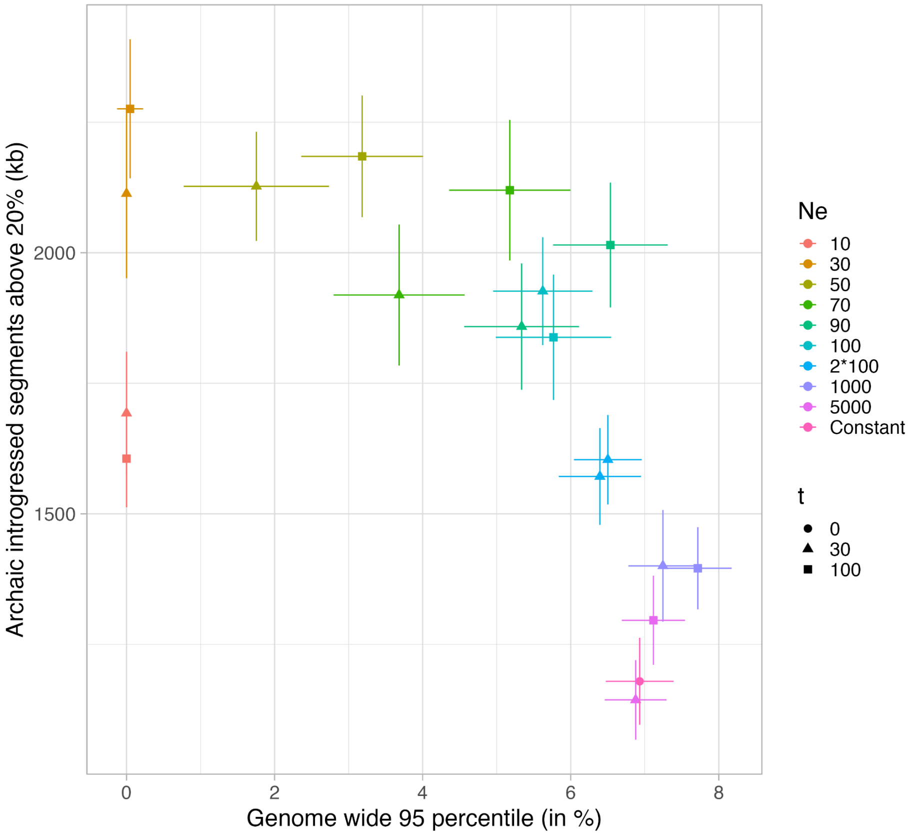

**Figure S8.3. Impact of bottlenecks on allele frequency and amount of introgressed segments at high frequency.** (A) Amount of introgressed archaic segments with a frequency higher than 20%, (B) Genome-wide 95-percentile of archaic frequency, in function of the different simulated scenarios, (C) Genome-wide 95-percentile of archaic frequency in function of the amount of introgressed archaic segments with a frequency higher than 20%.

#### Distribution of archaic ancestry in worldwide populations

We estimated the mean and standard deviation of the 95th-percentile of archaic frequency and the amount of high frequency archaic segments (above 20%) using jackknife in Indians (from LASI-DAD), Europeans (Icelanders from deCODE) and East Asians (EAS from 1000G). We used Neanderthal and Denisovan introgressed segments inferred in Supplementary Note 6.

Most non-African groups— including Europeans, East Asians and Indians— differ in their recent demographic history, including recent admixture events, population bottlenecks, and expansions[11]. Consequently, we find the landscape of both Neanderthal and Denisovan ancestry varies across Eurasians. Despite similar amounts of Neanderthal ancestry in Europeans and Indians, Indians have considerably fewer high frequency archaic variants (>20%) compared to Europeans (27,566 Icelanders from deCODE) or East Asians (EAS from 1000G) (Figure S8.4). East Asians have both the highest 95-percentile threshold and largest number of high frequency archaic segments (Figure S8.4.A). For Denisovans, we find that

Indians have both the highest 95-percentile threshold and the fewest regions of high frequency archaic variants (Figure S8.4.B). Europeans have the lowest amount of Denisovan ancestry in Eurasia, and consequently, the 95-percentile threshold is close to zero (Figure S8.4.B). East Asians have less Denisovan ancestry than Indians (Figure S8.4.B) but more is shared across individuals within the population than Indians.

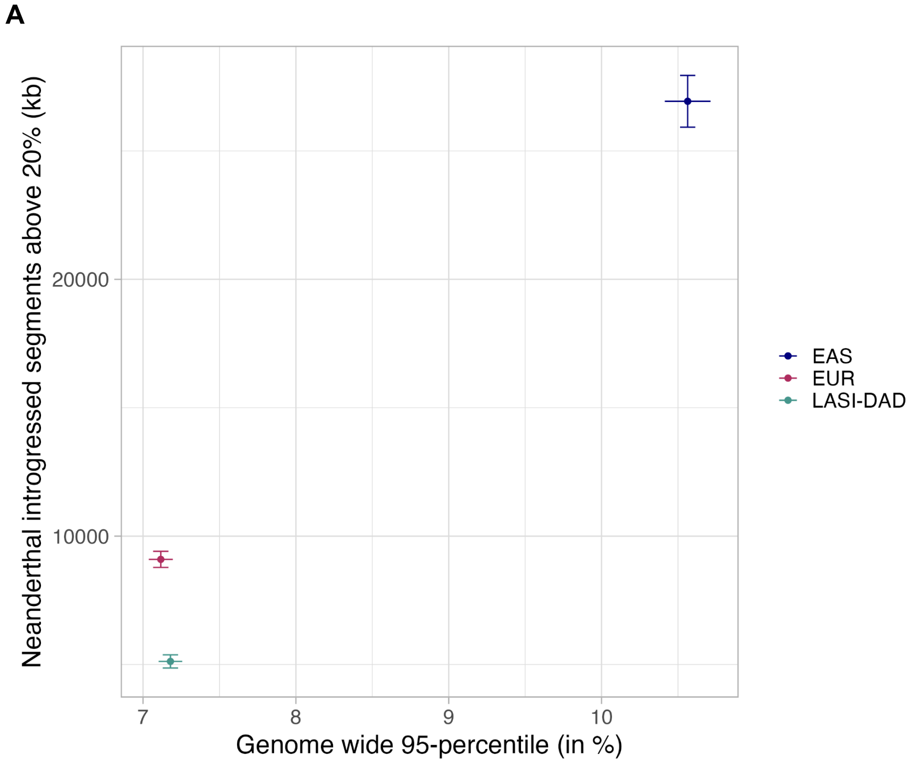

**B**

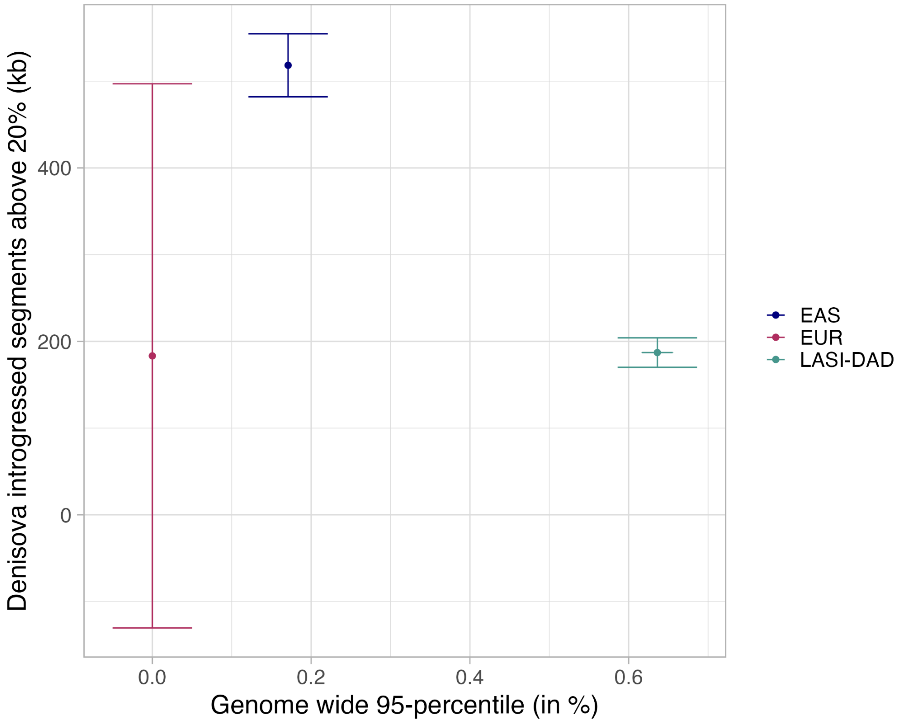

**Figure S8.4. Genome-wide 95-percentile of archaic frequency in function of the amount of introgressed archaic segments with a frequency higher than 20%.** Indians (from LASI-DAD), Europeans and East Asians (from 1000G), for A. Neanderthal and B. Denisova.

### Supplementary Note 9: Revising the Timing the Out of Africa migration to India

A central question in human evolutionary history is understanding when the ancestors of present-day non-African populations left Africa. Previous studies have argued that patterns of genetic variation in present-day individuals are consistent with a single major migration out of Africa that occurred between 40-72 thousand years ago (kya)[23,77–82]. However, archaeological studies have suggested that there is evidence of early human occupation in India since 74 kya[83] where humans likely migrated along the Southern (or coastal) route of dispersal from East Africa to the Near East, across the Red Sea into India, and then finally to Asia and Australasia. In support of this support, one recent study of genetic variation in Papuans suggests that around 3% of their ancestry traces back to an earlier migration out of Africa that occurred ~120 kya[63,84].

Our HMM classifies the genome into two states: a 'modern human (or human)' and an 'archaic' state. We use sub-African Africans as an outgroup and exclude any mutations shared with the outgroup. The emission parameter represents the average number of mutations per base pair (bp) in the given state. In turn, the emission parameter $emission_{i\,|\,human/archaic}$ corresponds to the average minimum coalescence time between the tested individual *i* and any individual from the outgroup[40].

$$emission_{i\,|\,human} = \mu \cdot L \cdot T_i$$

Where $\mu$ is the human mutation rate, $L$ is the genomic window size and $T_i$ is the average minimum coalescence time between the test individual and the outgroup.

For instance, the emission parameter for the modern human $emission_{i\,|\,human}$ state provides insights about the time to the most recent common ancestor for a non-African and Africans or in other words, the time of the out of Africa migration. Further, by comparing this estimate across diverse non-Africans, we can assess if the data support a single major dispersal out of Africa or multiple events.

*Model for estimating emission parameter for modern human state*

The emission parameter for the modern human state provides a count of the number of mutations since the separation from Africans. Theoretically, the emission probability is related to the human mutation rate and minimum coalescence time between the ingroup (i.e., test individual *i*) and the outgroup, in this case Africans.

However the emission parameter can be affected by multiple factors such as 1) Recent African gene flow, 2) Remnant archaic ancestry, and 3) Bioinformatic effects. We will address each of these in turn.

1. Recent African gene flow

Recent studies have shown that there is evidence for recent sub-Saharan African-related ancestry in many European and Middle Eastern groups, as well as there is back gene flow from Europe-related groups into Africa in the past 5,000 years[30,85,86]. Such gene flow will lead to increased sharing between the ingroup and outgroup, in turn leading to a larger fraction of variants being removed from the analysis and consequently, lower emission rates.

To assess the evidence for gene flow between populations related to the outgroup (sub-Saharan Africans) and ingroup (non-Africans), we performed ADMIXTURE[28] in unsupervised mode with $K$=2. We ran ADMIXTURE using 1,150,639 autosomal variants present on the 1240K AADR array[26]. We successfully converted the genomic positions of 99.95% of the variants from the human genome build hg19 to hg38 (532 SNPs failed conversion) using the *liftover* tool. For each individual, we estimated the amount of sub-Saharan African-related ancestry. Consistent with previous reports, we find significant evidence of sub-Saharan African-related ancestry in Middle Easterns, Southern Europeans[11,30,86], groups from the Americas[11,30,86] and unexpectedly, in Oceanians (Figure S9.1).

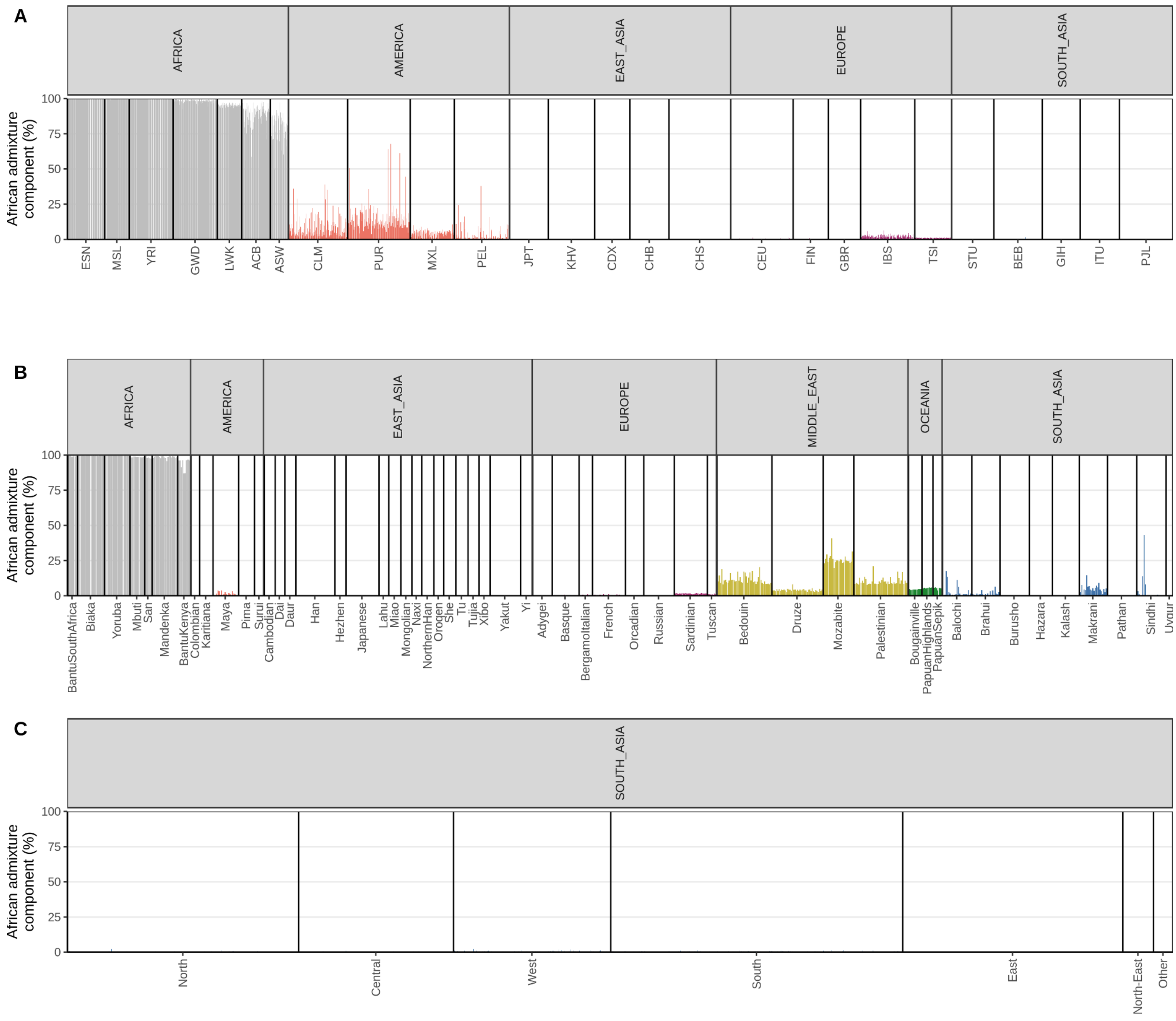

**Figure S9.1. Admixture proportion from ancestry component most common in Africa**. The height of each bar show the admixture proportion of the ancestry component most common in Africa for individuals from 1000G **A)**, HGDP **B)** and in LASI-DAD **C).** The individuals are grouped by region and population and colored by region.

Using the same dataset, we also ran ALDER[30] to infer the minimum ancestry proportion from sub-Saharan African-related populations into non-Africans (*POP*) using the one-reference setup with YRI as the outgroup with the following parameters:

genotypename: 1240K.geno
snpname: 1240K.snp
indivname: 1240K.ind
admixpop: *POP*

refpops: YRI
maxdis: 1.0
seed: 77
runmode: 1
zdipcorrmode: YES
chithresh: 0.0
binsize: .001

ALDER provides a lower bound on the ancestry proportion for each non-African population , while ADMIXTURE reports ancestry contributions per individual. Thus, it is difficult to make direct comparisons between the two methods. However, the median ancestry proportions per population are largely consistent across the two methods: ALDER minimum ancestry proportion and ADMIXTURE median ancestry estimate for individuals in a population is significantly correlated (*spearman rho* = 0.49, *p*-value = 6.712e-06). For the following analysis, we removed all individuals who had more than 1% ancestry related to sub-Saharan Africans based on the ADMIXTURE results (*n*=1881).

2. Amount of archaic introgression

A second confounder in our analysis is the spill over of archaic ancestry to the modern human state. Like most archaic ancestry inference methods, *hmmix* erros towards reducing false positive rates. In turn, it has a considerable false negative rate of ~20% for identifying archaic ancestry regions[40]. Moreover, short regions inherited through gene flow cannot be distinguished from regions of incomplete sorting. Thus, these regions or mutations are then assigned to the modern human state (referred to as 'spill over'). As the emission parameter for the archaic state is more than ten-fold higher than the human state (given the deep divergence between modern humans and archaic groups), this spillover has a significant impact on the estimated modern human emission parameter. Indeed, we observe strong correlation in the emission rates for both modern human and archaic states (Table S9.2). Therefore, we use the total amount of high confidence archaic ancestry (total amount of archaic sequence where segments have a posterior probability greater than 0.9 and share at least one derived variant with either Neanderthals or Denisovans) as a covariate in our analysis. For each individual *i*: $Archaic\ ancestry_i$ is to account for the remnant effects of archaic spill over to the modern human emission rate.

3. Bioinformatic effects

To compare the results across other worldwide populations, we merged the emission rates for LASI-DAD data with individuals from 1000G and HGDP. As observed by other groups, we noticed some batch effects due to different bioinformatic pipelines and filters applied for genotyping and phasing of different datasets, leading to over-calling or under-calling of variants across datasets. Specifically, we find significant variation in phasing drop-out rate across individuals in the 1000G dataset (Table S6.2 and Figure S6.1). We calculate the phasing drop-out rate as (the number of phased variants) / (the number of unphased variants). The phasing dropout rate is highly variable across individuals and groups in the 1000G dataset, which in turn leads to underestimation of the emission parameter for the 1000G populations.

The phasing dropout rate is fairly uniform across individuals in HGDP, regardless of the geographic region of origin with an average value of 0.3% (Figure S.9.2). It is 0% in LASI-DAD due to the fact that we are not using a reference panel for phasing. To minimize the impact of bias due to phasing on downstream analyses, we exclude 1000G from this analysis and focus on HGDP and LASI-DAD only. For each individual, we introduce a variable:

$$Bioinformatic\ effects_i = 1 - \frac{\#\ phased\ variants}{\#\ unphased\ variants}$$

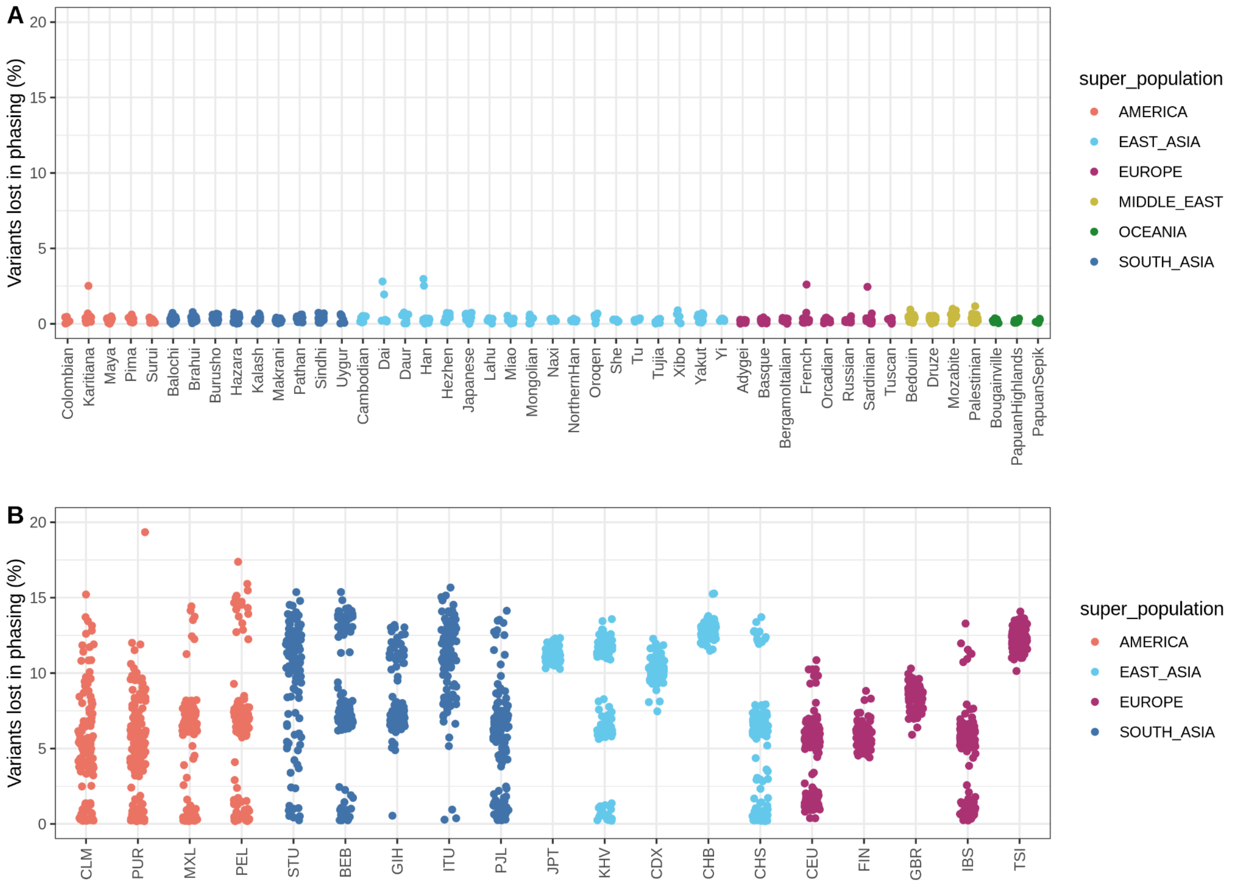

**Figure S9.2. Phasing-drop out rate is 1000G and HGDP individuals.** We calculate the percentage of variants lost in phasing as #phased SNPs / #unphased SNPs. The points are colored by region with HGDP dataset in panel A and 1000G in panel B.

For LASI-DAD, we only retain biallelic sites in the filtered dataset. With a sample size of ~2,700 high coverage genomes , we expect many recurrent mutations[87]. To obtain a minimum bound on the rate of missed recurrent mutations, we extracted all the SNPs from the 1240K array and assessed if they contained multi-allelic variants before filtering (Supplementary Note 2 - VCF STEP 2 Quality Checks). We chose this set of SNPs because they contain common variants

across world-wide populations and thus can be reliably sequenced. We find that 2.35% of sites show evidence for recurrent mutations.
Limiting our analysis to biallelic sites will underestimate the modern human state emission parameter and hence the coalescent time to sub-Saharan Africans for LASI-DAD individuals. To account for this, we add a correction for LASI-DAD individuals which is based on our inference of the depletion of recurrent mutations:

$$Bioinformatic\ effects_i = 1 - 0.0235$$

This also allows us to more reliably compare the results with HGDP which utilizes experimental phasing and includes multi-allelic sites[23].

The corrected emission parameter $corrected\ emission_{i\,|\,human}$ for individual *i* accounts for the confounding variables of $Archaic\ ancestry_i$ and $Bioinformatic\ effects_i$ in the following manner:

$$emission_{i\,|\,human} = corrected\ emission_{i\,|\,human} + \beta_1 * Archaic\ ancestry_i + \beta_2 * Bioinformatic\ effects_i$$

where, $\beta_1$ captures the relationship between archaic ancestry and emission parameter and $\beta_2$ captures the relationship between bioinformatic effects for all individuals (Table S9.1).

The corrected emission parameter $corrected\ emission_{i\,|\,human}$ for each individual in each super-population in each dataset (HGDP and LASI-DAD) is then inferred as:

$$corrected\ emission_{i\,|\,human} = emission_{i\,|\,human} - (\beta_1 * Archaic\ ancestry_i + \beta_2 * Bioinformatic\ effects_i)$$

**Table S9.1. Modeling of the emissions from the modern human state for each dataset.**

| Parameters | Coefficient name | Estimate | P value |
|---|---|---|---|
| Archaic ancestry<br>(change per 1 bp of archaic ancestry) | $\beta_1$ | 3.37e-11 | <2e-16 |
| Bioinformatic effects | $\beta_2$ | -2.78e-04 | <2e-16 |

We then infer the minimum coalescent time time between the ingroup and outgroup using the corrected emission parameter $corrected\ emission_{i\,|\,human}$ using the formulation described earlier.

$$corrected\_emission_{i\,|\,human} = \mu \cdot L \cdot T_i$$

Where $\mu$ is the human mutation rate, $L$ is the genomic window size and $T_i$ is the average minimum coalescence time between the test individual and the outgroup.

Recent studies have shown that the human mutation rate is highly uncertain and may have evolved over the course of human evolution[88]. To capture the uncertainty in the estimates of the mutation rate, we used a mean mutation rate of 0.45 x $10^{-9}$ per bp per year, with a range of 0.4-0.5 x $10^{-9}$ per bp per year (Figure S9.3).

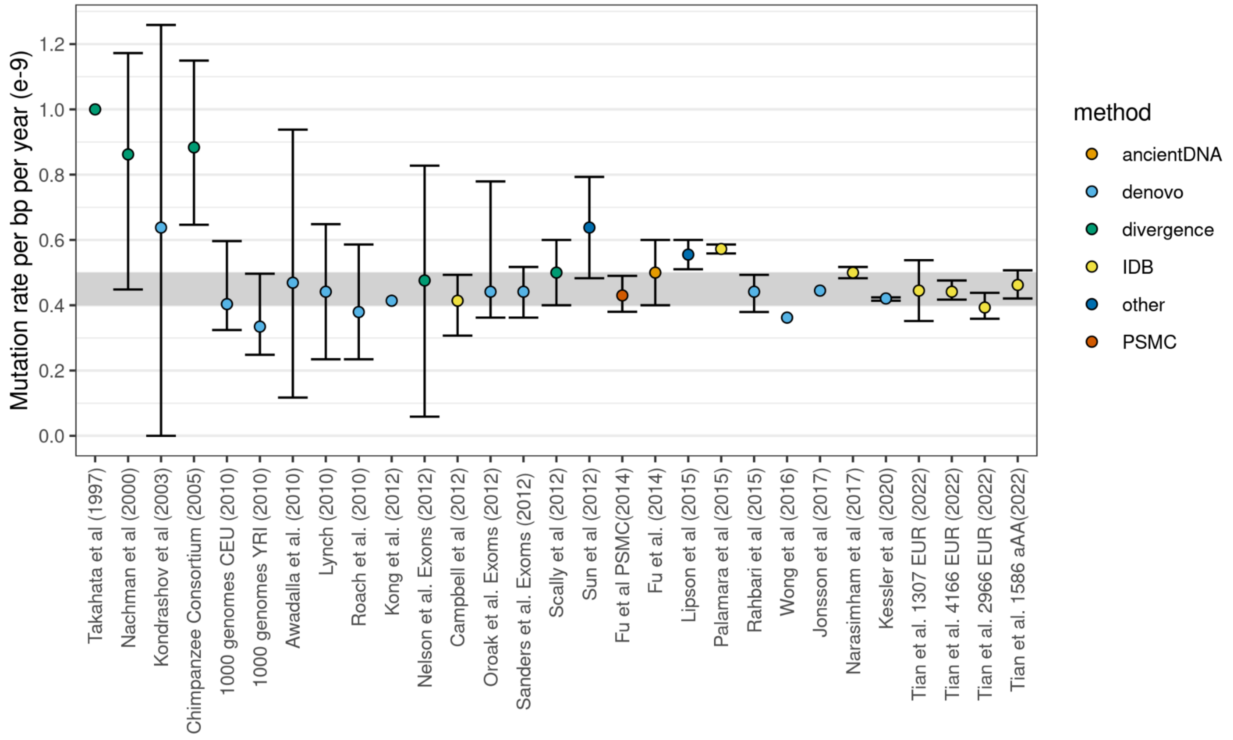

**Figure S9.3. We show 26 previous estimates of the mutation rate in modern humans.** The citations are listed on the *X*-axis and sorted by year of publication. They are colored according to the method used to estimate mutation rate (shown in the legend). The dot denotes the mean estimate and the error bars represent the 95% confidence intervals of the mean estimate from the publications. The grey horizontal bar indicates the range of mutation rates we consider for dating the OOA event.

We show the average minimum coalescent time of various non-African groups to sub-Saharan Africans in Figure S9.4 and Table S9.2**.**

**Table S9.2 Estimated average minimum coalescent time with the outgroup for different regions from HGDP and LASI-DAD datasets.** We used a window size *L* of 1000 bp and a mutation rate ($\mu$) ranging from (0.4 - 0.5) x $10^{-9}$ mutations per bp per year[13]. We report the mean and 95% confidence interval of the coalescence times (in years) for each non-African population with the outgroup. We also calculate the mean emissions jointly for all non-Africans.

| dataset | population | $\mu$ = 0.4 x $10^{-9}$ per bp per year | | $\mu$ = 0.45 x $10^{-9}$ per bp per year | | $\mu$ = 0.5 x $10^{-9}$ per bp per year | |
|---|---|---|---|---|---|---|---|
| | | mean | 95% percentile range | mean | 95% percentile range | mean | 95% percentile range |
| HGDP | America | 48874 | 47894 - 50758 | 54983 | 53881 - 57103 | 61092 | 59868 - 63448 |
| HGDP | East Asia | 48228 | 47308 - 49332 | 54256 | 53221 - 55499 | 60284 | 59134 - 61666 |
| HGDP | Europe | 46708 | 45396 - 48244 | 52547 | 51071 - 54275 | 58386 | 56746 - 60306 |
| HGDP | South Asia | 48192 | 47175 - 49728 | 54216 | 53072 - 55944 | 60240 | 58969 - 62160 |
| LASI-DAD | South Asia | 47940 | 47280 - 48572 | 53932 | 53190 - 54644 | 59924 | 59100 - 60716 |
| LASI-DAD + HGDP | All non-africans | 47940 | 46916 - 48816 | 53932 | 52781 - 54918 | 59925 | 58645 - 61020 |

Although the 95% percentile intervals overlap we do find statistically significant differences between the following populations. For the estimated minimum coalescence time with Africa, 99.4% of individuals from LASI-DAD fall within the 95% percentile range for all non-Africans ( 52,781 - 54,918 years). The same is true for populations in HGDP, with 81.8% of South Asians, 89% of East Asians, 52% of Americans and 37.1% of Europeans falling within the CI for all non-Africans. This lower overlap in Europeans could reflect additional ancestry from a population that diverged more recently from the outgroup, possibly related to the basal eurasian group as suggested in some studies[89].

We note, however, that simulations show that the inferred coalescent time is slightly underestimated by *hmmix*[40]. e.g., for demography of non-Africans, a difference of ~5% was observed between the true and inferred modern human emission rate.

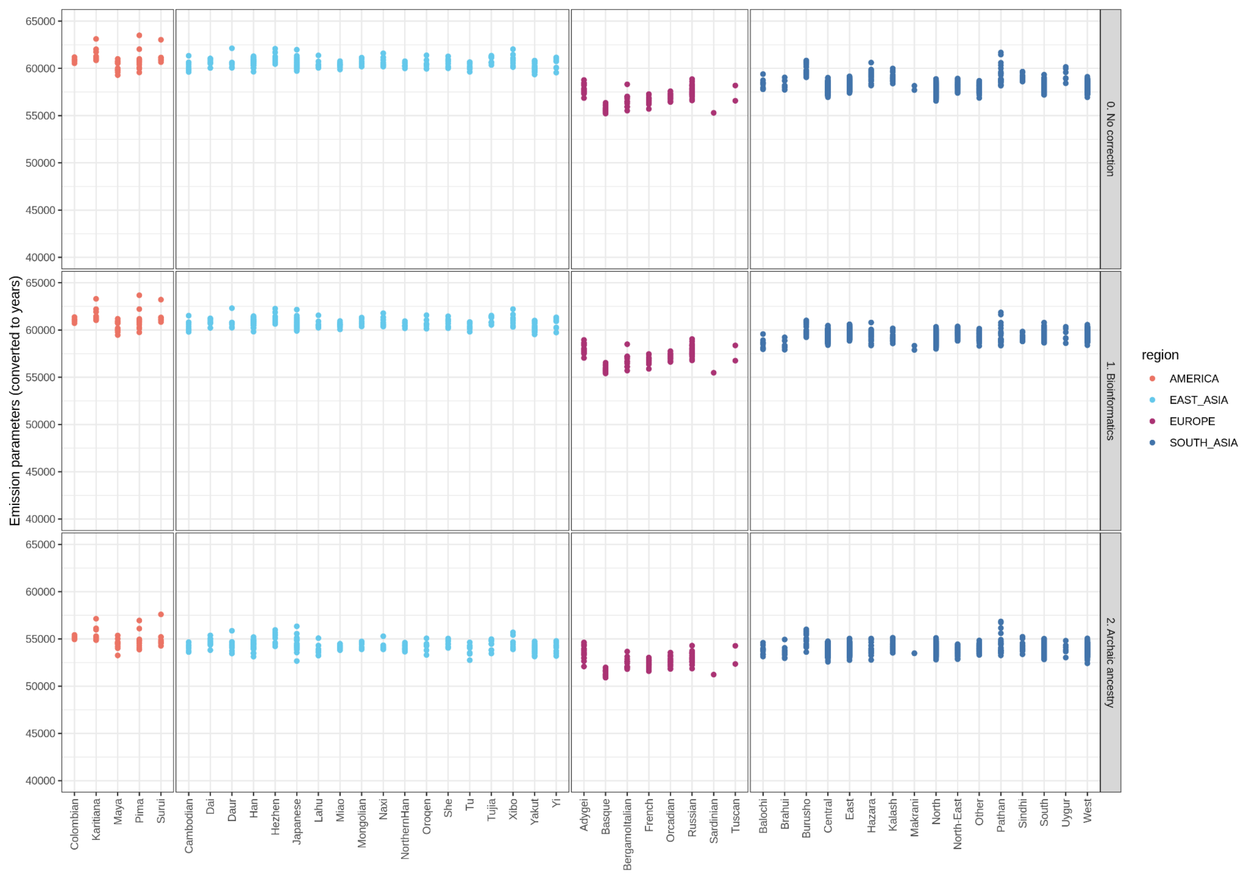

**Figure S9.4. Minimum coalescence time with Africans using emission parameters for populations from HGDP and LASI-DAD dataset.** We show uncorrected emission values (top panel), emission values corrected for bioinformatic effects (middle panel), and emissions values corrected for bioinformatic effects and archaic ancestry (bottom panel). We use a mutation rate of 0.45 x $10^{-9}$ per base pair per year to convert the emission parameters into years.

**Power to detect multiple waves out of Africa**

In order to assess if the corrected emission parameters in India shows evidence of ancestries from multiple out of Africa migrations, we performed simulations. We simulated 2.48 Gb of total sequence ($L$) and modeled the observed mutations deriving from a mixture of two poisson processes with means ($\lambda_1$, $\lambda_2$), where the proportion of the genome coming from $\lambda_2$ is $p$ and the rest from $\lambda_1$. Thus, the emission parameter for the modern human state for each individual can be modeled as:

$$emission_{human} = (1 - p) * \lambda_1 + p * \lambda_2$$

where, $\lambda_1$ represents the expected number of mutations per bp per year through the major wave out of Africa migration that occurred at $T_1$ years ago ($\lambda_1 = L \cdot \mu \cdot T_1$) and $\lambda_2$ represents the expected number of mutations due to the minor wave out of Africa that occurred at $T_2$ years ago ($\lambda_2 = L \cdot \mu \cdot T_2$).

We allow the timing of the minor wave out of Africa ($T_2$) to vary between 0-130,000 years ago and the proportion of the minor wave ($p$) between 0 and 25%. We set the human mutation rate to 0.45 x $10^{-9}$ per bp per year.

In Figure S9.5, we show the range of estimates for parameters that are consistent with the data. By consistent we mean that the simulated values fall between the minimum and maximum value in India (53,190 and 54,644 years respectively) of the emission parameters in India (Table S9.2).

Archaeological evidence suggests the timing of the Southern Dispersal to India was ~75,000 years ago, as there is evidence of human occupation in India before and after the eruption of the Toba that occurred 75.0 ± 0.9 kya[90]. Assuming ancestry from an earlier minor wave out of Africa at 75,000 years ago, our data is consistent with contribution of 0-3% ancestry from this earlier wave.

In conclusion, the populations of South Asian, East Asian and Americans have qualitatively similar human emission parameters. These correspond to an estimated minimum coalescence time of around 53,932 years ago. For India, our results are consistent with the majority of the ancestry deriving from a single major pulse, with minimal (0-3%) contribution from an earlier wave pre-Toba eruption.

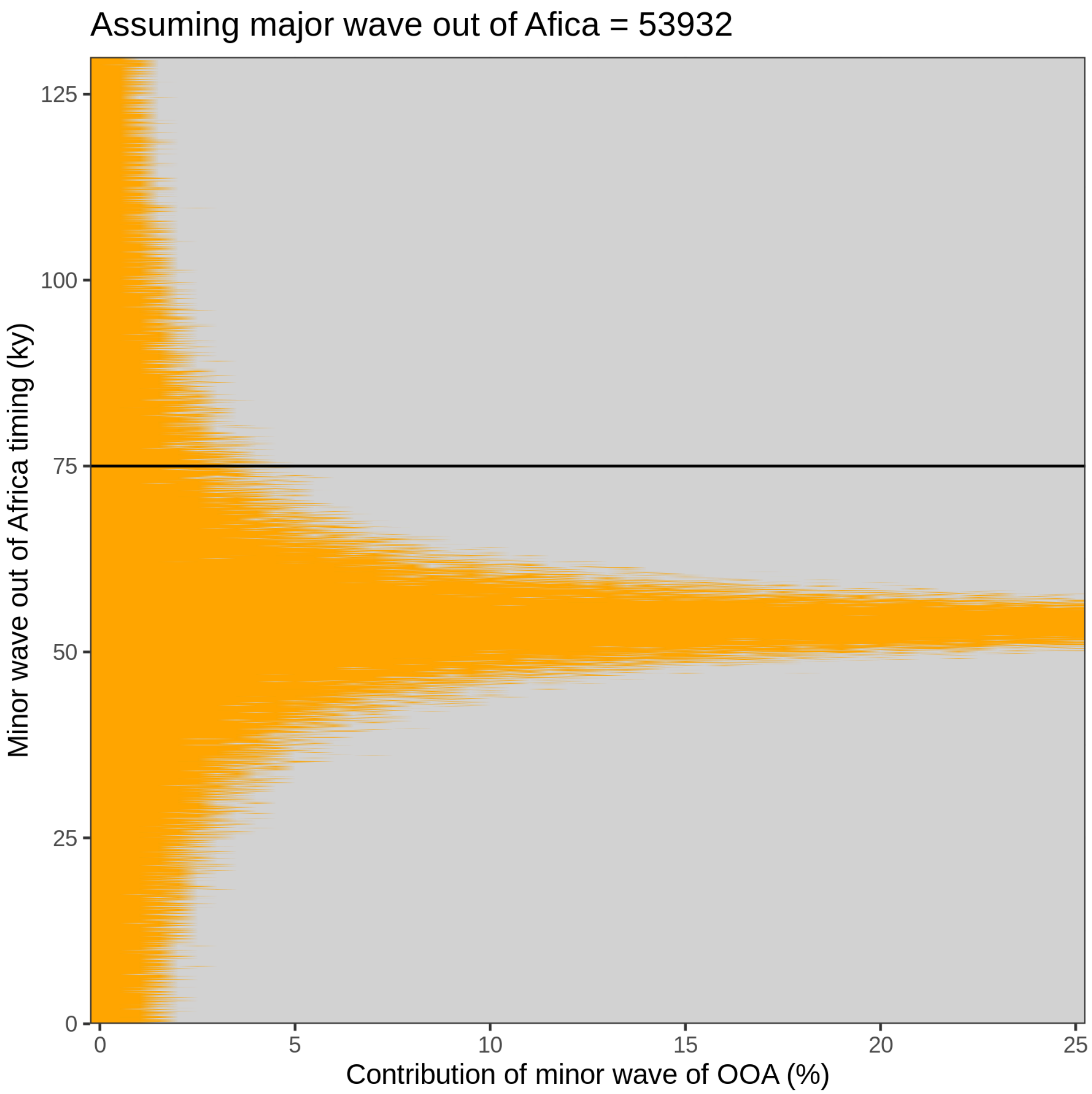

**Figure S9.5 Estimates of the contributions and timing of the minor wave out of Africa that are consistent with LASI-DAD.** The horizontal line indicates a suggested initial out of Africa wave at 75 kya (putative estimate of the timing of the Southern Dispersal out of Africa to India). The vertical line indicates the maximum contribution of the minor wave that is consistent with the corrected modern human emission rate in LASI-DAD.
